# Biomolecular Condensates Dictate the Folding Landscape of Proteins

**DOI:** 10.64898/2026.01.12.699095

**Authors:** Nathaniel Hess, Jerelle A. Joseph

## Abstract

Protein structure is exquisitely sensitive to the surrounding chemical environment, and many proteins encounter complex environments within cells. Importantly, numerous proteins organize into biomolecular condensates—dense macromolecular assemblies with distinct physicochemical properties. This raises a fundamental question: how does condensation reshape protein structure and dynamics? Here, we investigate how protein folding landscapes are altered inside condensates, using the protein α-helix as a model folded domain. Atomistic simulations show that free energy surfaces within condensates differ markedly from those in dilute solution or in the presence of inert crowders. We then use Bayesian optimization to develop a chemically specific, near-atomistic model for quantification of α-helical folding and apply it to characterize diverse helices, including α-helical domains from the disease-associated proteins TDP43, Annexin A11, and the Androgen Receptor, within condensates of varying physicochemical properties. Our results support a framework in which multivalent interactions drive unfolding while crowding promotes folding, and protein conformational ensembles inside condensates emerge from this balance. Additionally, we show that folding transitions are kinetically frustrated inside condensates because they are coupled to the timescale of contact rearrangement with co-condensate proteins. As such, folding landscapes within condensates are dually sequence-dependent, informed by both the sequence of the folded domain and co-condensate proteins. Together, our work has implications for understanding condensate-mediated proteinopathies, targeting aberrant condensates, and designing condensates to program protein function across scales.

## I. INTRODUCTION

Biological environments within cells have been evolutionarily tuned over millions of years to optimize their conditions for regulating the biomacromolecules essential for life. Two of these environments, the cytosol and the nucleoplasm, have tightly controlled properties including their molecular crowding [1, 2] and pH [3]. However, the picture of a relatively homogeneous bulk cytosol and nucleoplasm has been complicated by research demonstrating that macromolecules, like proteins and nucleic acids, can form distinct membraneless compartments within cells [4, 5]. These compartments, referred to as biomolecular condensates, can have chemical environments that are dramatically different from the underlying conditions present in the bulk cellular milieu, including variations in pH by over a unit from standard physiological conditions [6, 7], a dielectric constant as much as ten-fold lower than that of water [8], and substantial differences in small molecule partitioning [9–11]. Much still remains to be determined in regard to how the distinct chemical environments within biomolecular condensates impact the biophysical characteristics of their molecular constituents.

Understanding how condensates affect molecular biophysics is especially crucial for structured proteins and structured protein domains within multidomain proteins (i.e., proteins that contain both structured and disordered regions), which are essential to condensate form and function [12, 13]. Prior work has reported striking alterations in domain conformation within biomolecular condensates, including modulating the folding equilibrium of immature SOD1 in condensates formed of CAPRIN1 low-complexity domain [14] as well as the unfolding of PAB1’s and TIA1’s RNA recognition motifs upon condensing [15, 16]. Studies involving enzymatic folded domains underscore these findings; for example, in the conformational switching of the Dcp1/Dcp2 enzyme complex with the Edc3 activator protein in P-bodies [17] and the increase in reaction rate efficiency for BTL2 enzyme in Laf1 condensates [18]. In many of these studies [14, 15, 18] it is proposed that interactions with co-condensate proteins—and chiefly intrinsically disordered regions (IDRs)—can alter structured protein domain conformation. Ultimately, the tight coupling between protein structure and function makes it imperative to elucidate the biophysical principles by which disparate chemical environments of biomolecular condensates modulate folded protein structures.

The protein α-helix is an ideal model system to investigate how biomolecular condensates alter protein folding landscapes. α-helices are readily modulated by chemical environments, with dramatic changes in helical fraction demonstrated for helical peptides in alcohols [19] and hydrophobic micellar solvents [20]. Critically, α-helix domains are key players in condensate function and dysfunction. These include the α-helix domain in the C-terminal region of TDP43, which modulates TDP43’s condensation and function [21], and the α-helix domains in the N-terminal region of the Androgen Receptor (AR) protein, which are integral to its role as a transcription factor and are hypothesized to mediate its oncogenic behavior [22]. Additionally, the α-helix domains of TDP43 and ANXA11 can seed amyloidlike fibrils and protein aggregates in frontotemporal lobar degeneration [23], Alzheimer’s disease [24] and ALS [25–27]. Interestingly, a proposed pathway for protein aggregation involves biomolecular condensates [28, 29]. Therefore, investigating the conformational behavior of α-helix domains within condensed environments can provide critical insights on cellular proteinopathies and folded domain biophysics within biomolecular condensates.

Here, we leverage well-tempered metadynamics and temperature replica exchange molecular dynamics (MD) at atomistic resolution to show that a polyglutamine-based helix–forming peptide undergoes a substantial shift in its conformational ensemble within a condensate, relative to both dilute solution and inert crowding conditions. Thereafter, we develop a chemically specific residue-resolution model, Mpipi-Helix, to probe the folding landscapes of model helical peptides within condensates at longer time and length scales. Additionally, we characterize the biophysical behavior of α-helix domains within the IDRs of TDP43, ANXA11, and AR, simulating them in condensates composed of their physiological interaction partners. We further take advantage of milliseconds of aggregate simulation time for these systems to build detailed Markov State Models (MSMs) and interrogate the folding landscape of α-helices in condensates at a near-atomistic resolution.

Our work predicts that multivalent interactions between folded domains and co-condensate proteins favor unfolding, counteracting the entropic crowding forces that typically stabilize compact structures. Strikingly, we find a coupling between protein-folding timescales and contact lifetimes with co-condensate partners, a relationship that universally frustrates folding kinetics within condensates. Therefore, our results point to a “dually sequence-dependent” framework for protein folding in condensates—one shaped by both the sequence of the folded domain and that of its co-condensate partners.

## II. RESULTS

### A. Crowding Alone Does Not Explain a Condensate’s Effect on Protein Folding Thermodynamics

We began our investigation with atomistic simulations of a polyglutamine helical peptide construct (AQ). In particular, we asked how the thermodynamic helix–coil (folded– unfolded) equilibrium of the AQ peptide would be altered if it were incorporated into a condensate compared to its behavior in a dilute or crowded environment (Fig. 1a). To this end, we simulated the AQ peptide in a dilute solution comprised of water and salt, a condensate formed by proteins of the sequence NYQQYN, and a crowded environment containing polyethylene glycol (PEG) polymers with an identical volume fraction as NYQQYN in the condensate. Our condensate-forming sequence was chosen so that it contained multiple polar groups, thereby ensuring the condensate proteins would be fluid and amenable to sampling. We further leveraged enhanced sampling approaches to facilitate convergence of our measurements, with well-tempered metadynamics [30] for the dilute condition and temperature replica exchange [31] for the two dense environments.

**Fig. 1.**
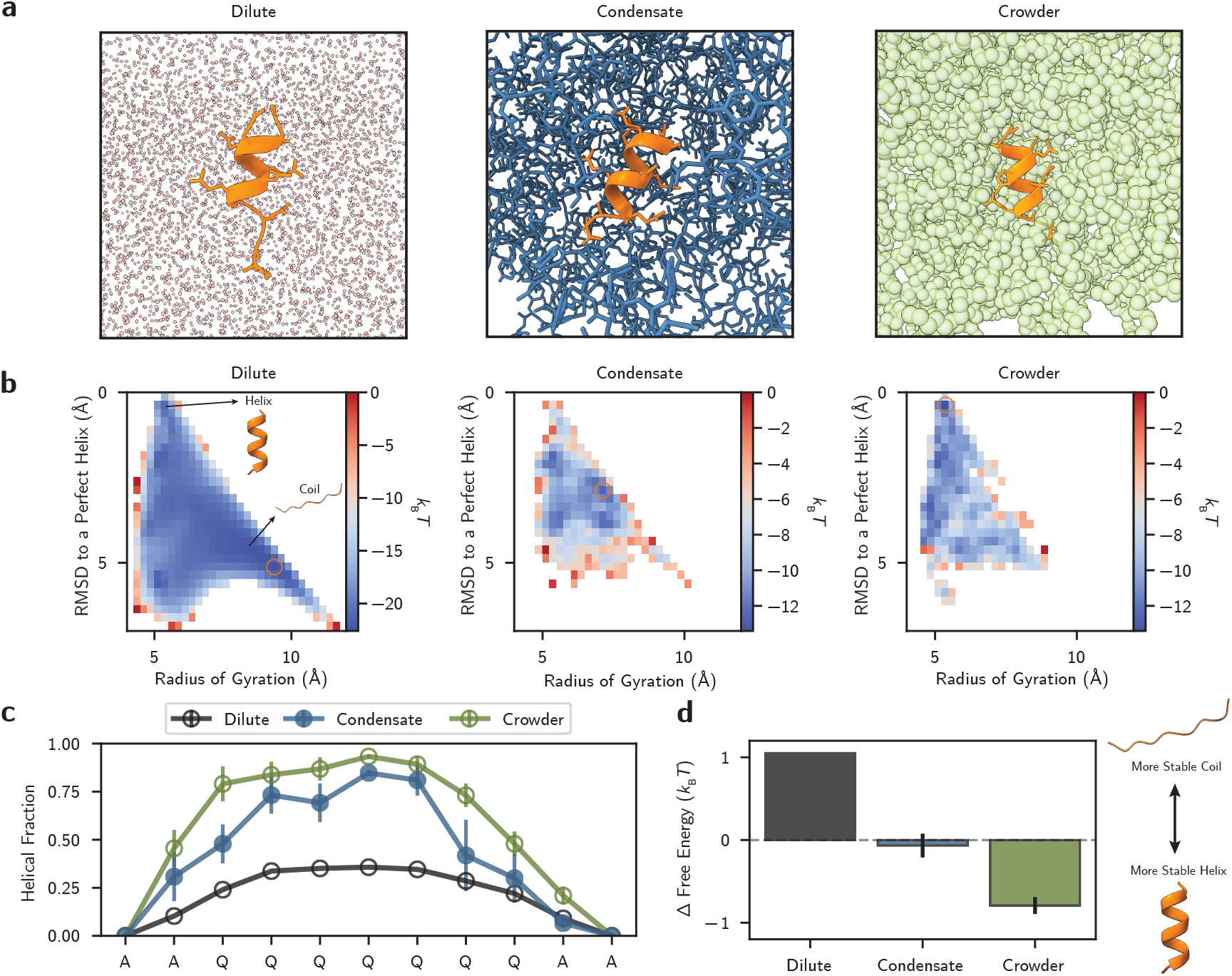
Crowding alone does not explain protein folding thermodynamics within a condensate. **a**, Atomistic simulations of the AAQQQQQQQAA (AQ) helical peptide simulated in dilute (left), NYQQYN condensate (middle), and PEG crowder (right) conditions. **b**, Free energy surfaces of the AQ peptide with radius of gyration and root-mean-squared deviation between the AQ peptide and a folded helix as order parameters. Orange circles indicate free energy minima. **c**, Per-residue helical fractions of the AQ peptide from backbone dihedral angles analysis. **d**, Estimated free energy differences between the folded helix and unfolded coil states of the AQ peptide. Errors for **c**,**d** are measured using block averaging over three equally-sized blocks.

We calculated the free energy surface (FES) of the AQ peptide in each environment against the order parameters of root-mean-squared deviation (RMSD) to a perfect helix and radius of gyration (Fig. 1b). The FES shows that in the dilute environment AQ has a narrow helical basin at low RMSDs and a broad unfolded region at high RMSDs. Intriguingly, however, within the condensate AQ has a substantially altered FES. In particular, the stability of the helical basin is enhanced relative to the coil state. This is reflected by the position of the global free energy minimum (orange circles in Fig. 1b), which shifts from *R*_g_ = 9.38 Å, RMSD= 5.12 Å in the dilute solution to *R*_g_ = 7.12 Å, RMSD= 2.88 Å in the condensate. Thus, the most stable structure for the AQ peptide in the condensate is a partially folded helix, compared to the dilute solution where the unfolded coil is favored. Additionally, in the crowded environment, the helix is stabilized even more strongly relative to the coil state compared to the dilute solution or condensate.

This increase in folding for the AQ peptide is further evidenced by measurements of the per-residue helical fractions (Fig. 1c). This quantity reflects whether the backbone dihedral angles (*ϕ, ψ*) for each amino acid in the peptide are in a helical configuration and thus provides a coarse picture of the helix–coil equilibrium for a given protein. Using common classifications for (*ϕ, ψ*) angles [32, 33], we report the per-residue helical fractions in each of the three chemical environments. As expected based on the FESs, we find an increase in helical fraction for the AQ peptide going from the dilute, to condensate, to crowded conditions. Using a two state model for helical propensity, we estimated the free energy difference between the helix and coil states of the AQ peptide in each environment (Fig. 1d; Methods). We find a substantial increase in helix stability in both the condensate and crowded environments compared to the dilute environment, with a helix–coil free energy difference of −0.07 *k*_B_*T* and −0.79 *k*_B_*T* that favors the folded state compared to the dilute environment (1.05 *k*_B_*T*).

A plausible explanation for the change in helical content within the dense environments is that the high density of neighboring molecules (NYQQYN in the condensate and PEG in the crowded environment) crowds the AQ peptide. This crowding would provide an entropic driving force for protein folding and bias proteins towards more compact structures [34, 35]. Further, higher volume fractions of ‘crowders’ should result in a greater propensity for AQ to fold into a helix. However, in our simulations, the volume fractions for PEG and NYQQYN are approximately the same (45.45% ± 0.15% for PEG compared to 44.74% ± 0.15% for NYQQYN). Therefore, crowding only partially explains the change in folding of the AQ peptide within dense environments. We hypothesize that, alongside crowding, sequence-dependent interactions with co-condensate proteins tune the conformational landscape of folded domains inside condensates. We systematically probe this hypothesis next.

### B. Multivalent Interactions with Co-Condensate Proteins Drive Protein Unfolding

Probing the folding landscapes of α-helices inside condensates at an atomistic resolution is computationally demanding even with enhanced sampling approaches. This is exemplified by the systems in Fig. 1 requiring over 60 µs of aggregate simulation time on *>* 25,000 atoms. Therefore, to enable the exploration of longer length and time scales for sequence diverse α-helices and condensates, we developed a chemically-specific coarse-grained model (Mpipi-Helix; see Methods and SI). Here, each amino acid is represented as a single interaction site and the model is trained to dynamically capture α-helix domain folding and unfolding in a sequence-dependent manner. To this end, we designed a novel combined Gaussian potential that encodes both the thermodynamic stability (i.e., sequence-dependent well depths) and the kinetic barriers associated with α-helix formation (Extended Data Fig. 1). In brief, Mpipi-Helix was parameterized through a Bayesian optimization approach informed by experimental data and atomistic simulations (Extended Data Fig. 1, 2). We validated our model against NMR measurements and atomistic data, confirming that Mpipi-Helix accurately captures α-helix folding landscapes (Extended Data Fig. 3).

We leveraged Mpipi-Helix to systematically probe the folding landscapes of model helical peptides in different condensates (Fig. 2a,b). We also characterized the peptides in dilute and crowded environments, developing an excluded volume polymer model to represent polyethylene glycol for the latter (Fig. 2b). We then simulated each peptide–environment pairing in Mpipi-Helix and built MSMs to interrogate the α-helix folding landscapes (Fig. 2c).

**Fig. 2.**
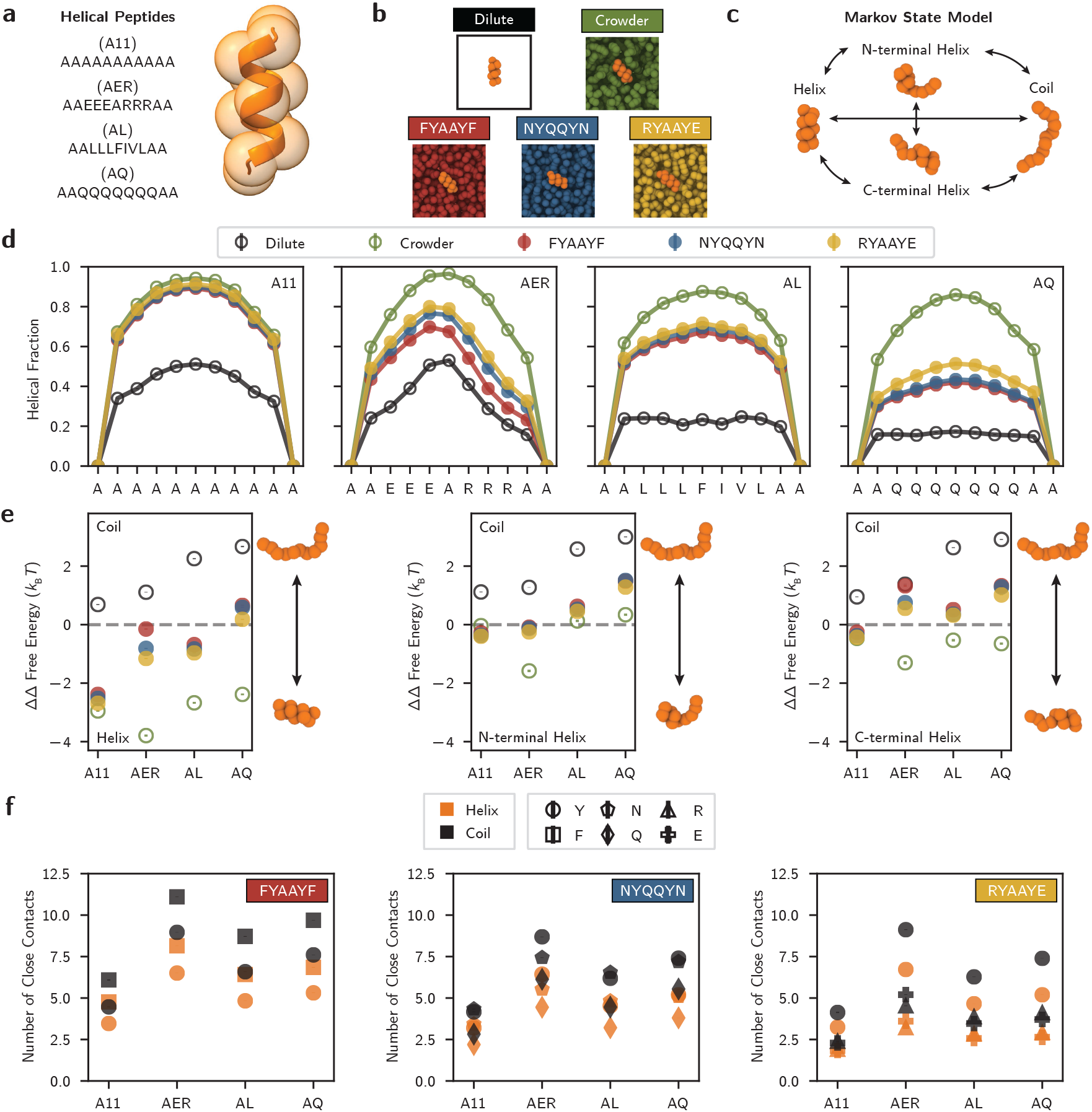
Condensates control protein folding thermodynamics through multivalent interactions and crowding. **a**, Four model helical peptides are simulated: an all alanine peptide (A11), a charged peptide (AER), an aliphatic peptide (AL), and a polar peptide (AQ). **b**, These peptides are simulated in five chemical environments, three condensates (FYAAYF, NYQQYN, and RYAAYE) as well as a dilute solution and generic crowding condition. **c**, Markov state models are built with the PyEMMA software [86] and coarse-grained into four interpretable metastable states using PCCA+ spectral clustering [36]. **d**, Helical fractions for each peptide– environment pairing. Standard errors are calculated via 300 2.5 µs replicates for each pairing and are smaller than the line width. **e**, Relative free energy differences between the (left) helix and coil states, (middle) N-terminal helix and coil states, and (right) C-terminal helix and coil states. Standard error are taken from 100 different instantiations of the MSMs combined with Bayesian uncertainty within the models. **f**, Number of close contacts between the helical peptide in the folded helix state (orange) versus the unfolded coil state (black) and co-condensate protein residues. Error is calculated as in **d**.

For each peptide, the helical fraction in the condensates lies between the values observed in dilute conditions (lower bound) and the crowded environment (upper bound) (Fig. 2d). Moreover, there are large variations in helical fraction both between the helical peptides and for the same helical peptide across different condensate environments. For example, A11’s helical fraction is almost identical to the crowded environment whereas the AQ peptide has helical contents that are relatively more closely aligned with the dilute helical fractions. Further, the AER and AQ peptides exhibit substantial variation (~9%) in their helical fractions across the condensates, whereas A11 and AL display comparatively smaller variation (≤4%). There is also a consistent ordering of helical fractions within the condensate environments that is conserved across all helical peptides, with FYAAYF exhibiting the lowest helical content, RYAAYE the greatest, and NYQQYN falling in between. Overall, there are clear qualitative differences in the folding equilibrium of the helical peptides within the various condensates.

Next, we predicted the free energy differences between the metastable states of the helical peptides in our MSMs using PCCA+ analysis [36] (Fig. 2e). As expected, the dilute environment favors an unfolded state whereas the crowded condition greatly increases the relative stability of the helix. However, our results depict striking differences in the relative free energies of the helical peptides within condensates. Although the helix–coil stability for A11 and AER are similar in both the dilute and crowded environments, we observe substantial differences in their average helix–coil stability inside the condensates suggesting that condensates can amplify differences in the conformational preferences of peptides. Moreover, while the helical state is favored over the coil state by approximately 4*k*_B_*T* for AER in the crowded environment, within the FYAAYF condensate AER shows virtually no preference for the helix or coil state, further exemplifying the ability of condensates to tune peptide conformational ensembles. Indeed, the AQ peptide favors the unfolded state in all three condensates (Fig. 2e left panel). Additionally, the FYAAYF condensate favors unfolding the most, which holds across all three metastable states (Fig. 2e) and is shown unambiguously by examining the relative free energies of the coil state (Extended Data Fig. 4a right panel). The degree to which FYAAYF unfolds a helix is, however, peptide specific; for example, only AER exhibits a notable difference (*>* 1*k*_B_*T*) in N versus Cterminal helix stabilities (Fig. 2e right two panels). Taken together, the evidence suggests that condensates regulate α-helix folding thermodynamics.

At least two non-exclusive hypotheses exist for what is causing these observed differences in α-helix folding thermodynamics in the various condensates: differences in the crowding of the co-condensate proteins and differences in the multivalent interactions that underlie the condensates. To interrogate the former, we measured the density and volume fraction of each condensate (Extended Data Fig. 4b,c). However, both properties are similar across condensates; for example, the volume fractions of FYAAYF, NYQQYN, and RYAAYE are 0.530, 0.547, and 0.529, respectively. Further, the modest difference in volume fraction between the NYQQYN and RYAAYE condensates does not align with crowding-based expectations; despite its higher volume fraction, NYQQYN induces greater peptide unfolding than RYAAYE. Thus, although crowding explains the overall increase in folding (i.e., from the dilute to the condensate to the crowded environment), it cannot account for the differences in folding observed from one condensate to another.

Another plausible explanation is that differences in the multivalent interactions between the helical peptides and co-condensate proteins govern the α-helix folding equilibrium. To test this hypothesis, we computed protein–protein contacts within the condensate. Specifically, we calculated the number of close contacts between the helical peptides and co-condensate proteins, using the metastable states in our MSMs to delineate when the peptides were folded or unfolded. We found that close contacts are universally enriched when the helical peptides are unfolded (black symbols in Fig. 2f, Extended Data Fig. 4e,f). This increase in contacts ranges from 25% for unfolded A11 interacting with charged residues in the RYAAYE condensate, to about 36% for unfolded AER in FYAAYF interacting with phenyalanine residues. We further calculated the difference in pairwise interaction enthalpies between the helical peptide and co-condensate proteins in the folded and unfolded states (Extended Data Fig. 4d). The rank ordering of pairwise enthalpy differences for each helical peptide in the condensates precisely mirrors the rank ordering of the condensate’s ability to promote protein unfolding. Further, the average pairwise enthalpy difference for each helical peptide matches the ordering of how strongly their folds are destabilized in condensates: A11 shows the smallest enthalpy difference (−1.15 ± 0.02 *k*_B_*T*) and is least destabilized, whereas AER shows the largest difference (−5.05 ± 0.02 *k*_B_*T*) and is, overall, the most destabilized in condensates. Together, these results suggest that condensates exert an enthalpic driving force to unfold α-helices, sustained by multivalent interactions between the unfolded peptide and co-condensate proteins.

Overall, the evidence presented here demonstrates that condensates can strongly modulate the conformational ensembles of folded protein domains. In particular, the data show that within condensates there is a competition between crowding, which favors compaction, and multivalent interactions with co-condensate proteins, which provide an enthalpic driving force to unfold. This competition explains why the FYAAYF condensate, with its strongly interacting tyrosine and phenylalanine residues, unfolds the helical peptides the most. It also explains the variations in the observed thermodynamics of the helical peptides within condensates; A11 only weakly interacts with the co-condensate proteins and thus exhibits folding most similar to the crowded condition, AER and AQ interact the most strongly with co-condensate proteins and unfold the most, and AL resides between the two. Additionally, the observation that the C-terminal helix of AER is highly destabilized within condensates is also explainable within this framework, as the arginine residues in that region of the peptide have strong cation–π interactions with aromatic residues in the co-condensate proteins. Thus, the data support a framework in which multivalent interactions counterbalance crowding within condensates to tune the folding thermodynamics of α-helices, in a manner shaped by the specific chemistries of cross-interactions between folded domains and co-condensate proteins.

### C. Condensates Frustrate Protein Folding Kinetics

Another critical aspect of a protein’s folding landscape is its kinetic barriers, which inform transition times between metastable conformations. Accordingly, we utilized our MSMs to characterize the kinetics of α-helix folding transitions within condensates, and compared these with the corresponding behavior in the dilute and crowded environments. Concretely, we computed the mean first passage time (MFPT) for four transitions: helix to coil, coil to helix, N to C-terminal helix, and C to N-terminal helix (Fig. 3a,b). As anticipated, we observe faster folding times and slower unfolding times in the crowded environment compared to the dilute solution (Fig. 3a), with the dilute folding times closely matching experimentally measured helix nucleation dynamics in alanine-based peptides [37–39]. Additionally, the transition times for the helical peptides within condensates are arrayed between the dilute and crowded conditions, with the lone exception of the folding time for AER in FYAAYF being ~40% longer than that of the dilute solution. In fact, FYAAYF has transition times that most closely resemble those in the dilute solution, with RYAAYE being relatively more like the crowded condition, and NYQQYN in between the two. This trend reflects the rank ordering of how each condensate tunes α-helix folding thermodynamics, with FYAAYF favoring unfolded states the most out of all three condensates. We also observe an outsized impact of the C-terminal domain of AER which has over twice as long of an N to C-terminal as C to N-terminal helix folding time in condensates, which aligns with the C-terminal helix’s lower stability within condensates (Fig 2e). Additionally, we find substantial variations within the same helical peptide across condensate environments; for example, both AQ and AER exhibit higher folding MFPTs (~23% and ~59%, respectively) in FYAAYF compared with RYAAYE condensates. Altogether, the MFPTs indicate that kinetic transitions between conformational states are altered within condensates.

**Fig. 3.**
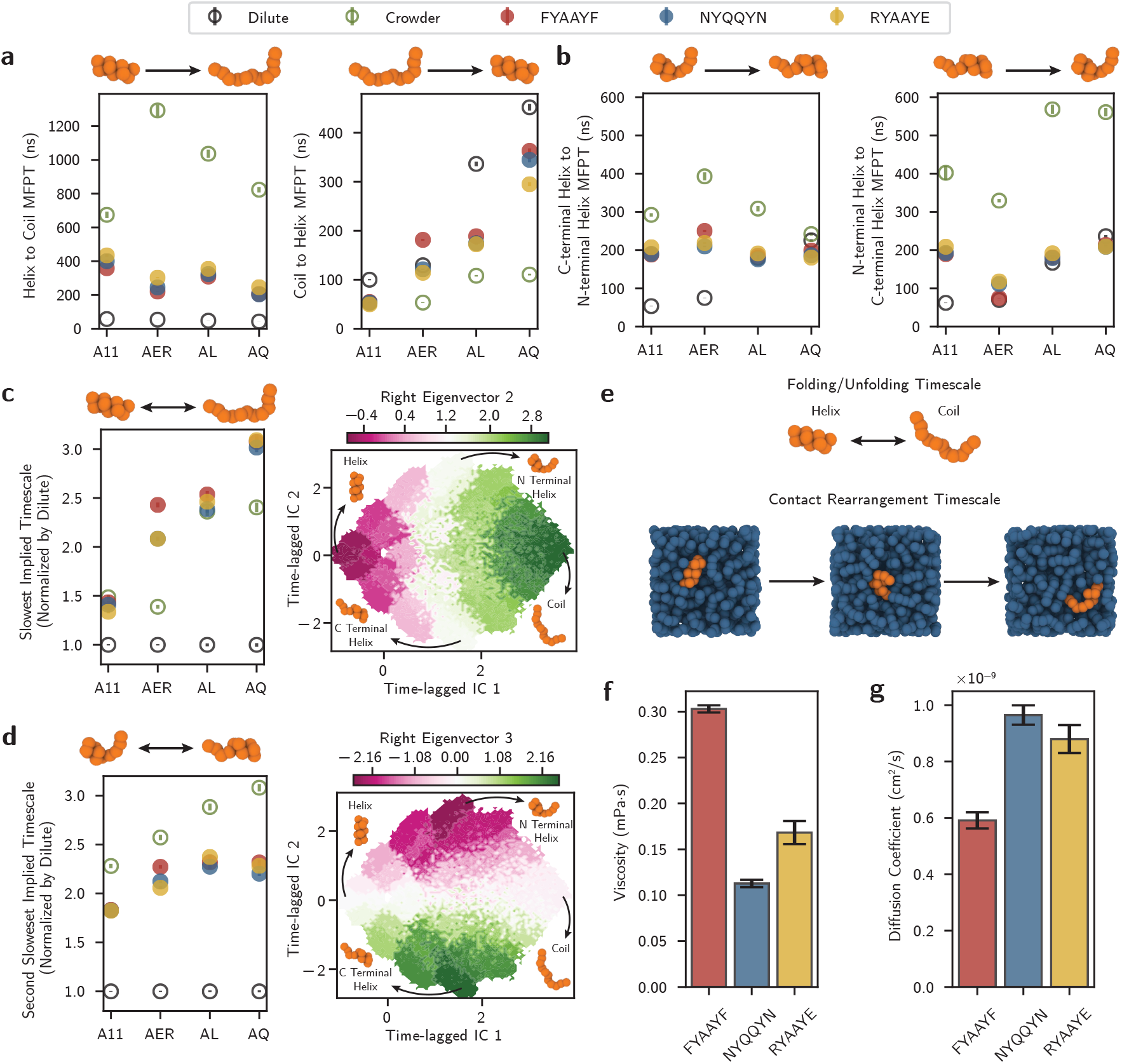
Condensates frustrate protein folding kinetics through coupling conformational transitions to contact rearrangements with co-condensate proteins. **a**, Mean first passage time (MFPT) between (left) helix to coil and (right) coil to helix. Standard errors are from 100 different instantiations of the MSMs combined with Bayesian uncertainty within the models. **b**, MFPT between (left) N to C-terminal helix and (right) C to N-terminal helix. Error is calculated as in **a. c**, The slowest implied kinetic relaxation timescale (left), corresponding to the transition between the helix and coil states (right). Errors calculated as in **a**. The right eigenvector plot is from an MSM applied to the A11–FYAAYF condensate pairing. **d**, The second slowest implied kinetic relaxation timescale (left), corresponding to the transition between the N and C-terminal helix. Errors are calculated as in **a** and the right eigenvector plot as in **c**. Values in **c** and **d** were normalized by the dilute solution mean. **e**, Both folding and contact rearrangements occur simultaneously in condensates. **f**, Viscosity calculated by the Green–Kubo relation for each condensate. Standard error is calculated across 5 independent simulation replicates. **g**, Diffusion coefficients of the co-condensate proteins within each condensate. Error is calculated as in **f**.

Our MSMs also provide access to the relaxation timescales of the slowest kinetic modes for α-helix folding. These timescales provide a robust classifier of the overall ruggedness of the α-helix free energy landscape, where longer decay times indicate higher kinetic barriers between metastable states. For the AER and AQ peptides, the characteristic decay time of the slowest kinetic mode of the folding process is greatly increased in condensates (Fig. 3c, left panel), exceeding even that observed in the crowded environment. For example, there is over a 50% increase in the relaxation timescale of AER in the condensates over that observed in dilute and crowded conditions. The right eigenvector of this decay process, which reflects the kinetic mode of the folding–unfolding transition (Fig. 3c, right panel), indicates that there are substantially higher barriers between the folded and unfolded states for AER and AQ within condensates. In contrast, both A11 and AL have decay times in condensates that are comparable with a crowded environment (differences of *<*6.5%), which likely reflect their weaker interactions with the co-condensate proteins. Further, the strongest interacting co-condensate protein, FYAAYF, has equal or longer relaxation times than the other condensates, which supports a hypothesis that multivalent interactions with co-condensate proteins contribute to the observed variability in folding timescales. However, our analysis is complicated by the second kinetic mode, corresponding to a transition between an N and C-terminal helix, which is uniformly faster in condensates than in a generic crowded environment (Fig. 3d). A plausible reason for this observation is that this transition requires both unfolding (which is faster in condensates than a crowded environment) and folding (which is slower in condensates than a crowded environment), causing these competing effects to partially cancel each other. Overall, however, the evidence from the slowest timescale suggests that condensates can have a substantial effect on the folding–unfolding kinetic timescales, increasing them beyond what is experienced in a crowded environment.

One possible explanation for slower folding timescales in condensates is the coupling of folding transitions to rearrangements of contacts with co-condensate proteins (Fig. 3e). It is likely that contacts with co-condensate proteins must be broken or established when a folded domain undergoes conformational changes inside a condensate. This view is supported by our prior observation that upon unfolding, the number of contacts between co-condensate proteins and helical peptides increases markedly (Fig. 2f). Two quantities that are correlated with the timescale for contact rearrangements are viscosity and the diffusion coefficient of the co-condensate proteins (Fig. 3f,g). For example, a condensate with a longer contact rearrangement timescale should exhibit a higher viscosity than one with a shorter rearrangement timescale (and vice-versa for diffusion coefficients). As such, we would expect the ordering of viscosities, based on the relaxation timescales in Fig. 3c,d, to follow the order of *η*_FYAAYF_ *> η*_RYAAYE_ ≈ *η*_NYQQYN_. This is precisely what we observe when we measure the viscosity of each condensate (Fig. 3f). Additionally, the diffusion coefficients follow the inverse rank ordering (Fig. 3g). However, it should be noted that the co-condensate protein rearrangement timescales only partially explain these observations; the true folding timescale within condensates is set by the cross interactions between the folded domain and the co-condensate proteins, which can not be fully captured by the co-condensate protein timescales alone.

Taken together, the evidence from the kinetic analyses suggest that α-helix folding landscapes are frustrated within condensates, prolonging transitions between folded and unfolded metastable states beyond what is explainable by crowding alone. We hypothesize that the timescales for protein folding are coupled to relaxation of contacts with co-condensate proteins. In other words, the multivalent interactions between a folded domain and co-condensate proteins must relax and rearrange before a conformational change can occur in the folded structure. Hence, strong interactions within condensates may result in longer timescales for conformational rearrangements of folded proteins. Ultimately, this effect may dominate if the timescale for contact rearrangements with co-condensate proteins is longer than the timescale of conformational transitions in a folded domain.

### D. The Folding Landscapes of Physiological Proteins are Dually Sequence-Dependent

Finally, we extended our investigation to probe the effects of condensates on α-helical domains within disordered proteins. We chose three disordered regions on condensate-associated proteins that contain at least one α-helix domain: TDP43 [21], ANXA11 [40], and AR [22, 41, 42] (Fig. 4a). Each of these proteins have known physiological partners that they interact with inside condensates (i.e., co-condensate proteins), including FUS, hnRNPA1, and TIA1 which TDP43 and ANXA11 interact with inside of RNA granules [43, 44], and RPB1, MED1, and BRD4 which AR interacts with inside of transcriptionally active condensates [22] (Fig. 4b). All of these proteins form condensates [16, 45–47], which we confirmed via direct coexistence MD simulations. We then simulated the helix-containing IDR of TDP43, ANXA11, and AR to understand how the folding properties of their α-helix domains are modulated inside condensates.

**Fig. 4.**
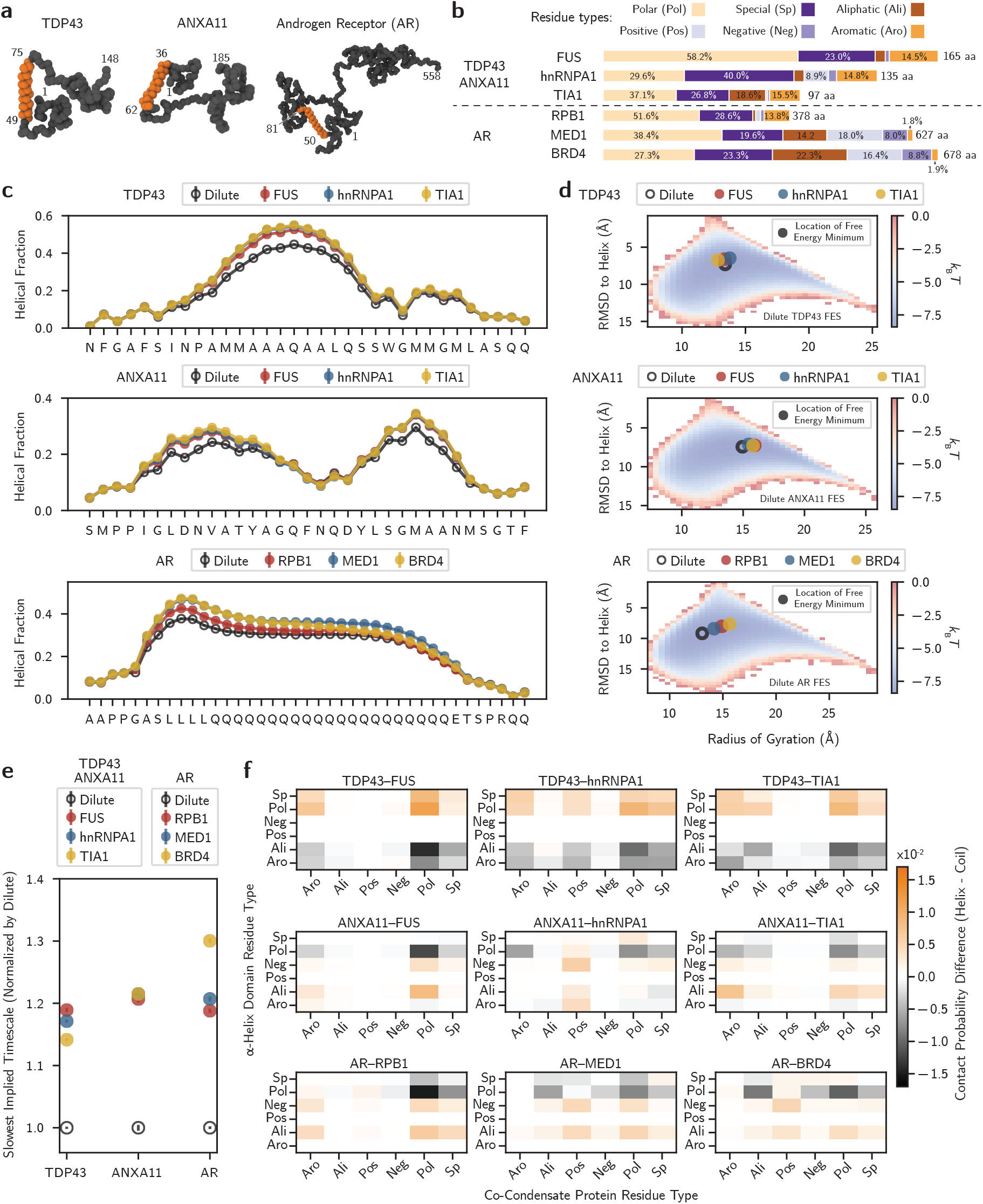
The folding landscapes of proteins are dually sequence-dependent within condensates. **a**, The α-helix domains (orange) within TDP43, ANXA11, and AR are simulated. **b**, Each domain is simulated in a dilute solution and three different condensates formed of physiologically-associated disordered protein domains. **c**, Per-residue helical fractions for each protein–condensate pairing. Standard errors are calculated using 300 2.5 µs replicates for each dilute solution, 200 2.5 µs replicates for TDP43 and ANXA11 in the condensates, and 120 2.5 µs replicates for AR in the condensates. **d**, The free energy surface of the dilute solution of each α-helix overlaid with the free energy minima of the α-helix domains in condensates. **e**, The slowest implied kinetic relaxation timescale for the α-helix domains in each environment. This timescale corresponds to a folding transition (see Extended Data Fig. 5c). Standard error is calculated across 25 different instantiations of our MSMs combined with combined with Bayesian uncertainty within the models. All values were normalized by the dilute solution mean. **f**, Average per-residue contact map differences between the helix and coil state for each protein–condensate pairing.

Consistent with the model helical peptides described in Fig. 2, the helical domains in TDP43, ANXA11, and AR demonstrate altered folding thermodynamics inside condensates. For example, the helical fractions for TDP43 and AR increase by as much as 10% for the TDP43–TIA1 and AR– BRD4 systems compared to the dilute condition (Fig. 4c). We also observe a shift in the global free energy minimum of the helical domains within the condensates, where the RMSD to a perfect helix for the lowest free energy conformation decreases (Fig. 4d). Importantly, none of these thermodynamic features are identical across condensates for the same α-helical domain. For example, there is a *>*5.5% decrease in helical fraction within the polyleucine region of AR in AR–RPB1 compared to AR–MED1 and AR–BRD4. This decrease in helical fraction is likely because RPB1 has over 10% more aromatic residues than BRD4 and MED1, which increases the enthalpic driving force for unfolding. Further, the FESs demonstrate that a given co-condensate protein does not equally perturb the conformational ensemble of the α-helix domains. Specifically, despite TDP43 and ANXA11 being colocalized with the same proteins, the ordering of the Wasserstein distances between their condensate and dilute solution FESs are not the same (Extended Data Fig. 5a,b). As such, we conclude that both the co-condensate protein and the α-helix domain sequences inform folding thermodynamics within condensates.

Next, we constructed MSMs to assess the folding kinetics of the α-helix domains inside condensates. As with the model helical peptides, we quantified the first implied timescale corresponding to the slowest decaying relaxation mode of the α-helix domains (Fig. 4e), i.e., folding and unfolding (Extended Data Fig. 5c). We find that there is an increase in kinetic timescale for each system compared to the dilute solution, with the largest increase of *>*25% for the AR–condensate systems and smaller gains of ~22% and ~17% for the ANXA11–condensate and TDP43–condensate systems. Notably, however, TDP43 and ANXA11’s implied timescales are not identical despite being colocalized with the same proteins. This indicates that the timescale for folding within condensates involves sequence-dependent cross interactions between folded domains and the co-condensate proteins. Overall, we observe kinetic frustration for α-helix domains within condensates that is a function of both the folded domain and co-condensate protein sequences.

The data presented above suggests that the folding landscapes of structured domains within condensates are dually sequence-dependent. In other words, protein folding thermodynamics and kinetics are influenced by the cross interactions between folded domains and co-condensate proteins. To more directly quantify how the α-helix domain sequences contributed to these changes in folding landscapes, we examined variations in contacts between the α-helix domains and co-condensate proteins. In particular, we calculated the difference in normalized contact probabilities between the α-helix domains and co-condensate proteins when the α-helix was folded versus unfolded (Fig. 4f). From this analysis, we observed distinct patterns of residue interactions that are enriched in folded and unfolded conformations. For instance, when the domain is folded, certain residues within an α-helix domain form relatively more contacts with co-condensate proteins, implying that interactions involving these residues promote folding. Specifically, interactions involving special (e.g., glycine and proline) and polar residues in TDP43 are relatively enriched in the folded state, whereas ANXA11 and AR have comparatively greater contacts involving their aliphatic and negative residues (orange squares in Fig. 4f). One explanation for this trend is that sites on the α-helix that engage in weak interactions with co-condensate proteins crowd the helix and promote its stabilization. Indeed, many of these residues are helix-breakers (glycines and prolines) or helix-neutral (aspartic acid, serine, and valine) [48], so interactions that crowd them may substantially aid helix formation.

In a similar manner, certain interactions are favored in the unfolded state and these are specific to the α-helix domain involved. In the unfolded state, TDP43 has a relative enrichment of contacts involving its aliphatic residues, whereas ANXA11 and AR preferentially interact with their polar residues (black squares in Fig. 4f). This behavior is explained by the sequence pattern of each domain: TDP43 has aliphatic residues in the middle of its highest helical stretch making these sites integral to its helix formation; ANXA11’s composition is more diverse and slightly more enriched in polar residues within high helical stretches; AR’s polyglutamine domain relies on the polar glutamine residues to form its helix. Overall, our data suggest that weakly interacting residues within structured domains become crowded inside condensates, promoting fold stabilization. In contrast, residues essential for maintaining domain integrity show relatively more interactions in the unfolded state within condensates, in a domain-specific manner. This is analogous to different point residue mutations in structured proteins being unequal in terms of their destabilization to a fold; similarly within condensates the residues on a folded domain that interact with co-condensate proteins and destabilize structure the most are not equal across disparate sequences.

In total, our data suggest that protein folding inside condensates depends on the sequence of both the structured protein domain and co-condensate proteins. Specifically, the exact residues which are stabilized or destabilized on a folded domain are unique to its own sequence. Similarly, co-condensate protein sequences shape how strongly a condensate perturbs the conformational ensemble of a folded domain and frustrates its conformational rearrangements. Therefore, within condensates, protein folding landscapes are dually sequence-dependent.

## III. DISCUSSION

Protein structures are profoundly influenced by their surrounding chemical environments, and biomolecular condensates constitute distinct physicochemical milieus that are densely enriched in proteins. Here, we present evidence for a biophysical framework that explains how biomolecular condensates tune the conformational ensembles and kinetic transitions of structured protein domains. We leverage a multiscale simulation approach that complements atomistic enhanced sampling simulations with a novel residue-resolution coarse-grained model to probe the folding behavior of α-helix domains inside condensates. Our work uncovers three emergent principles governing protein folding landscapes within condensates. First, multivalent interactions between structured domains and co-condensate proteins generate an enthalpic driving force for protein unfolding that directly competes with the crowding-induced entropic stabilization of compact folds (Fig. 5a). Condensates can therefore tune folding thermodynamics and conformational ensembles by balancing the generic crowding effect of weakly interacting residues (e.g., prolines and glycines) with stronger interactions (e.g., π–π, cation–π, charged interactions) between a structured domain and co-condensate proteins. Second, kinetic transitions within protein structural ensembles are uniformly frustrated inside condensates due to a coupling between conformational changes and the rearrangement of contacts with co-condensate proteins (Fig. 5b). Although the α-helix folding timescale that we characterize here may not generalize to conformational transitions of other protein structures, our data strongly suggest that when the rearrangement of contacts with co-condensate proteins is slower than the timescale of conformational changes, condensates can substantially prolong transitions between metastable states. Finally, the effects of condensates on folding thermodynamics and kinetics are dually sequence-dependent as they are driven by the cross-interactions between a folded domain and co-condensate proteins (Fig. 5c). As such, the fraction and patterning of multivalent and weakly interacting amino acids in the co-condensate proteins can determine protein folding thermodynamics and kinetics. Likewise, the residues which are stabilized or destabilized within a given folded domain are specific to that domain’s sequence. Together, these findings demonstrate how condensates can dictate the folding landscape of proteins.

**Fig. 5.**
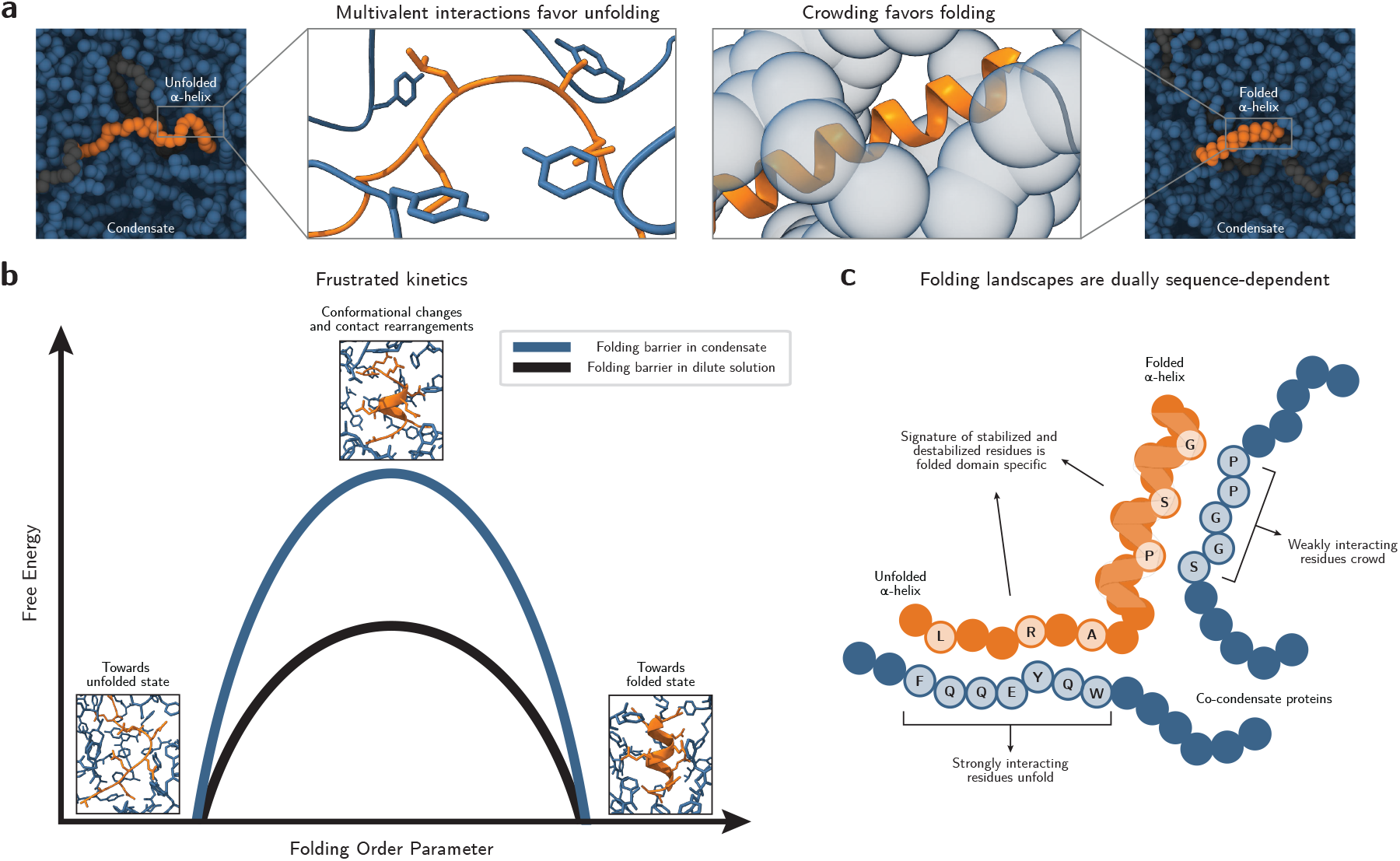
Condensates dictate the folding landscape of proteins. **a**, Multivalent interactions between folded domains and co-condensate proteins tend to unfold proteins and counteract the generic crowding provided from a dense environment. **b**, Condensates frustrate the folding kinetics of structured domains by coupling folding transitions to rearrangements of contacts with co-condensate protein residues. **c**, The folding landscapes of folded domains within condensates are dually sequence-dependent as they are informed by both the sequence of the structured domain and the co-condensate proteins.

Our results contextualize previous studies on protein and RNA structure inside condensates. Specifically, the Kay lab recently reported the unfolding of the immature SOD1 protein within CAPRIN1 condensates and provided evidence using NMR measurements that the CAPRIN1 low complexity domain acts as a proteinaceous solvent of SOD1 molecules [14]. Our observation that unfolded domains form increased contacts with co-condensate proteins is consistent with the notion of a proteinaceous solvent; the increased solvent accessible surface area generated upon unfolding is solvated by co-condensate proteins and is enthalpically stabilized via compatible chemistries. Further, the Sosnick lab [15] has demonstrated that the interactions which mediate condensate formation can lead to protein unfolding, as exemplified through the sequential activation and unfolding of the PAB1 RNA-recognition motifs. Additionally, the Barducci group used MD simulations to predict that positively charged condensates can melt an RNA tetraloop by acting as an RNA-philic solvent [49], consistent with previous results demonstrating the destabilization of RNA base-pairing in biomolecular condensates [50–52]. Our findings align with these prior works and provide evidence for the generality of condensate-mediated tuning of molecular folding landscapes. Future investigations into this effect promise to enable the engineering of systems with spatiotemporal control over structural ensembles through biomolecular condensates. Such studies will also clarify how condensates function within cells to influence aspects of folded-domain biophysics, for example binding affinities, which are a function of protein conformation and thus directly affected by the physicochemical environment within condensates.

Additionally, our framework (Fig. 5) provides insight on how the folding landscapes of disease-associated domains can be altered in biomolecular condensates. Concretely, we demonstrate that the folding landscapes of disease-associated α-helix domains of the TDP43, ANXA11, and AR proteins are functions of condensate chemical environments. This observation suggests a couple possible mechanisms by which condensates shape the behavior of disease-associated proteins. First, condensates may act as chaperones of protein structure (e.g., α-helix domains) by enriching residues that favor folded states, particularly through prion-like domains with high proline and glycine content. Second, condensates whose compositions change as they age and become enriched in strongly interacting sites (e.g., via post-translational modifications) may promote both domain unfolding and higher kinetic barriers along folding pathways, owing to longer contact relaxation timescales with co-condensate proteins. This may drive these domains into regions of the energy landscape where high kinetic barriers hinder refolding, leaving them in unfolded states that are more susceptible to aggregation. Separately, higher kinetic barriers within condensates may lead to the stabilization of metastable conformations in disease-associated proteins, introducing additional complexities for small-molecule drug targeting. Ultimately, elucidating the structures that proteins adopt within condensates will be essential for understanding condensate-related proteinopathies and for enabling precise therapeutic targeting [22, 42, 53].

Overall, our study provides a holistic framework that explains how the physicochemical environment inside biomolecular condensates tunes protein folding landscapes, putting forward a new role for condensates in directly shaping protein structure and dynamics.

## IV. METHODS

### A. Atomistic Simulations

#### 1. Dilute AQ Well-Tempered Metadynamics

The well-tempered metadynamics simulation for AQ peptide was performed using a combined force field, taking the structural parameters for AQ from the ff19sb force field [54] and combining that with a99sb-disp [55] in TIP4P-D water and salt. Simulations were performed in GRO-MACS v2024.3 [56] and PLUMED v2.9 [57]. Initial configurations (i.e., extended structures) were constructed using PyMOL [58]. Simulation inputs were generated using CHARMM-GUI [59] with 150 mM NaCl salt. Each system was first minimized using 5000 steepest descent steps with a force tolerance of 1000 kJ/mol/nm and positional restraints on the heavy atoms of the protein. A subsequent equilibration step was performed for 100 ns at a timestep of 2 fs at 298.15 K with a v-rescale thermostat and a 1 ps damping constant. Cutoffs for van der Waals and Coulomb interactions were set to 0.9 nm. Long-range electrostatics were calculated using a PME mesh grid. Bonds involving hydrogen were constrained with the LINCS algorithm [60]. Production was performed under the same simulation conditions but using well-tempered metadynamics [30] with the PLUMED [57] plugin for GROMACS. The biased collective variables were the root-mean-squared deviation to a perfect helix (DRMSD in PLUMED) and the radius of gyration. The bias factor was set to 5, with a hill deposition rate of 10 ps, and hill deposition size of 0.025 kcal/mol. Sampling was performed for 10 µs and convergence of the free energy surfaces was assessed, which showed no significant change after 9.2 µs.

#### 2. Atomistic Simulations of the NYQQYN Condensate

We first performed a direct coexistence simulation of a NYQQYN condensate alone to determine the density of the co-condensate proteins. The system comprised 141 NYQQYN protein chains in an extended configuration, constructed using PyMOL [58], in a 9×9×9 nm^3^ simulation box. We then minimized the system with positional restraints on heavy atoms. We used 5000 steepest descent steps with a force tolerance of 1000 kJ/mol/nm. We then performed an *NPT* equilibration step for 100 ns, using a v-rescale thermostat at 298.15 K and a 1 ps damping constant and a C-rescale barostat at a pressure of 1.01325 bar, a damping constant of 5 ps, and a compressibility of 4.5e^−5^ bar. The cutoff for van der Waals and Coulomb interactions were set to 0.9 nm. Electrostatics were calculated with a PME grid. Bonds involving hydrogen atoms were constrained with the LINCS algorithm [60]. After the *NPT* equilibration, we ran a subsequent *NVT* equilibration for an additional 100 ns. All simulation parameters were the same as in the *NPT* simulation except for the lack of a barostat.

The final configuration from the *NVT* equilibration was used to set up a direct coexistence simulation in the slab geometry (6×6×24 nm^3^), solvating with TIP4P-D water and 150 mM of NaCl after resizing the box. The system was then minimized with positional restraints on NYQQYN heavy atoms. We then performed a 1.5 µs *NVT* simulation using analogous parameters to the above *NVT* simulation. Simulation frames were recorded every nanosecond. We discarded the first 500 ns as equilibration and fit sigmoids to the density profiles from the slab simulation to determine the density of water (0.380 ± 0.007g/cm^3^), sodium and chloride ions (0.0005 ± 0.0001g/cm^3^), and protein (0.839 ± 0.010g/cm^3^) within the condensate.

#### 3. Temperature Replica Exchange of AQ in NYQQYN Condensate

We added the uncapped AQ protein initialized from a helical configuration to a system of 141 capped NYQQYN protein chains in a 10×10×10 nm^3^ box, which was solvated with water and salt according to the densities from our direct coexistence simulation. We then ran a minimization via steepest descent with a force tolerance of 1000 kJ/mol/nm for 5000 steps. After minimization, we performed a compression step in the *NPT* ensemble for 10 ns at a step size of 2 fs. We used a similar compression protocol as in the NYQQYN-only system. After *NPT* compression, the box size was approximately 6x6x6 nm^3^. We then ran a 100 ns equilibration in the *NPT* ensemble, with an identical set up as the above, but with the reference pressure of the barostat set to 0 bar. Additionally, the positional restraints on the NYQQYN co-condensate proteins were removed, but the positional restraints on the AQ peptide were maintained until the start of temperature replica exchange sampling.

Temperature replica exchange (T-REMD) was performed after the *NPT* equilibration. Replicas were set up in a temperature ladder (298.15 300.18 302.22 304.27 306.34 308.43 310.52 312.64 314.76 316.90 319.06 321.23 323.41 325.61 327.83 330.06 332.30 334.56 336.84 339.13 341.43 343.76 346.09 348.45 350.82 353.20 355.61 358.02 360.46 362.91 365.38 367.86 370.36 372.88 375.42 377.97 380.54 383.13 385.74 388.36 391.00 393.66 396.34 399.03 401.75 404.48 407.23 410.00 K). Replicas were simulated in the *NPT* ensemble, at a pressure of 0 bar with no positional restraints. During T-REMD, exchanges between neighboring replicas were attempted every 5 ps and frames were saved every 50 ps. T-REMD was run for 1.35 µs per replica (64.8 µs in aggregate), with the initial 100 ns discarded as equilibration. Statistics presented in the main text were gathered from analyzing the coldest replica (298.15 K).

#### 4. Temperature Replica Exchange of AQ in PEG Crowder

In a similar fashion, we simulated the AQ peptide in PEG crowder. We calculated that 385 PEG chains (of length 6 monomers) would produce the same volume fraction as the NYQQYN proteins in the above T-REMD simulation. We therefore created a system with uncapped AQ in a helical configuration and 385 PEG chains. We solvated it with the same amount of water and salt as in the above AQ– NYQQQYN simulation. Besides the difference in the number of polymer chains, every simulation step was performed identically to what is described above for the AQ–NYQQYN system.

#### 5. Free Energy Surfaces

Free energy surfaces were calculated for the AQ metadynamics simulation in the dilute phase, as well as from the T-REMD simulations of AQ in the NYQQYN condensate and PEG crowder. For the free energy surface calculations, we used the radius of gyration (*R*_G_) of the AQ peptide and the RMSD between the AQ peptide and a perfect helix as order parameters. For the metadynamics simulation, we took the radius of gyration and RMSD from every frame and weighted these values using the final deposited bias. For the T-REMD simulations, we leveraged MBAR [61] and the pyMBAR package to get the statistical weights of each simulation snapshot from the twelve coldest T-REMD replicas at 298.15 K. From there, we used the weights to compute the probability of AQ configurations in each *R*_G_–RMSD bin, with *P*_bin_ ∝ exp(−*F/k*_B_*T*) providing us the corresponding free energy (*F* is the free energy of the bin). We further applied shifting and truncation such that the zero free energy mark was the highest free energy of our systems and excessively high free energies (*>* 15 *k*_B_*T* above the free energy minimum originating from MBAR) were removed.

#### 6. Helix–Coil Classification in Atomistic Simulations

Helix and coil classifications for all atomistic simulations were determined using cutoffs of *ϕ* and *ψ* backbone angles that correspond with the region of the Ramachandran plot for α-helices. We use −30.0 *> ϕ >* −100.0 and −7.0 *> ψ >* −67.0, consistent with previous work [32].Within these cutoffs of *ϕ* and *ψ*, a residue is classified as helical and outside it was considered to be a random coil. The *ϕ* and *ψ* angles were calculated via the rama command in GROMACS.

#### 7. Atomistic Helix–Coil Free Energy Calculations

We calculated the relative free energy difference between the folded helix and unfolded coil states for the atomistic systems using a two state model. Specifically, we took the average helical fraction of the central nine residues of the AQ peptide in each environment and calculated the relative free energy difference as ln 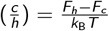. Here, *c* and *h* are the fraction of time spent in the helical and coil states. *F*_*h*_ and *F*_*c*_ are the relative free energies of the helix and coil state, respectively.

#### 8. Atomistic Volume Fraction Calculations

Free volume fraction was computed with GROMACS freevolume command on the coldest temperature replica and applied only to NYQQYN or PEG. One minus the free volume was taken as the volume fraction of these species. The error reported in the main text is the standard deviation of the free volume percentage over the course of the simulation.

### B. Mpipi-Helix Parameterization

#### 1. The Combined Gaussian Potential

In Mpipi-Helix, we used a combined Gaussian potential to simulate the α-helix structure. The combined Gaussian includes both a negative Gaussian energetic well that biases α-helix formation and a broader positive Gaussian that represents the barrier between the helix and coil states (Extended Data Fig. 1a,b). The potential is described in Equation 1.

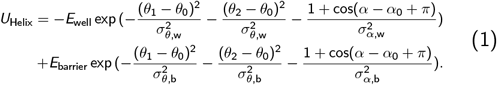

Here, *θ*_0_ and *α*_0_ set the centers of the Gaussians, *σ*_*θ*,w_, *σ*_*α*,w_, *σ*_*θ*,b_, and *σ*_*α*,b_ control the width of the negative and positive Gaussian, *θ*_1_, *θ*_2_, and *α* are the biased bond and dihedral angles, and *E*_well_ and *E*_barrier_ determine the magnitudes of the energetic well and barrier. We have implemented this structural potential—via a modified Wang– Frenkel [62] pairstyle and atom class—in LAMMPS [63] (see SI and the GitHub repository for details).

#### 2. Bioinformatic Data for Determining Gaussian Location and Width

We utilized bioinformatic data to set the location and widths of our Gaussian well and barrier (Extended Data Fig. 1c). The bioinformatic data was collected using the PISCES PDB culling server [64]. We queried for proteins with less than 25% sequence similarity resulting in 7,997 protein structures (exact query: Resolution 0.0–2.0, R-factor 0.25, sequence length 40–10,000, sequence percentage identity ≤ 25.0, chains with breaks excluded, chains with disorder included). A list of the PDB entries gathered from this query can be found in the GitHub repository. Next, we used the DSSP algorithm [65] to assign secondary structure classifications to each amino acid in the PDB structures and selected all α-helical segments of length greater or equal to four amino acids within the dataset. We then extracted the geometry of α-helices at an α-carbon resolution. Plotting the bond angles and dihedral angles of the α-helices informed our choice of *σ*_*θ*,w_ = 0.25 and *σ*_*α*,w_ = 0.4, as there is a significant overlap of the PDB α-helix geometries within a 10% cutoff of the energetic minimum of the helical well depth (Extended Data Fig. 1c). This analysis also informed our decision to place the peak of our helical barrier at the 10% cutoff of the energetic minimum of the helical well depth (Extended Data Fig. 1c). The remaining parameters in Equation 1 were determined through the parameterization procedure discussed below.

#### 3. Barrier Height Parameterization

We aimed to recapitulate the approximately order of magnitude difference in nucleation versus denucleation time for α-helices [37] with our barrier height parameterization. We reproduced this qualitatively by testing a variety of barrier heights with a fixed well depth of 3.0 kcal/mol (Extended Data Fig. 1d). We simulated the A11 peptide in the *NVT* ensemble across a range of helical barrier heights (0.5–5.0 kcal/mol in 0.5 kcal/mol increments). The system was thermostatted with a Langevin thermostat at 298.15 K and a damping constant of 100 ps. All simulations were performed in LAMMPS [63]. For each parameter set, we ran 150 independent simulations, each with an initial 0.5 ns equilibration followed by 10 µs sampling. The timestep was set to 10 fs and frames were saved every 2 ps.

Helix–coil classification was performed on a frame-by-frame basis in the same manner as was done for the Gaussian process training workflow (see below). Thereafter, helix nucleation time was calculated as the time elapsed between segments of three or more consecutive ‘H’ residues appearing in the protein chain. Helix denucleation time was then calculated as the time elapsed between having no groups of three or more consecutive ‘H’ residues in the A11 peptide. Finally, we chose a barrier height of 4.0 kcal/mol as it provides a separation of nucleation and denucleation timescales while maintaining a computationally feasible folding process to explore in MD simulations.

#### 4. Potts Model Training

The underlying supposition in our model parameterization was that there is a correlation between the number of times a sequence of four amino acids appears in helices in known protein sequences and the thermodynamic propensity of that sequence to form a helix. Based on this idea, we built a Potts model that maps any sequence of four amino acids to the statistical likelihood that those amino acids are part of an α-helix (Extended Data Fig. 1e).

The Potts model is described by equations 2 and 3.

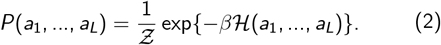

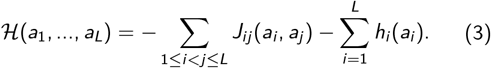

In these equations *a* represents any amino acid, *i* and *j* are any position in the protein sequence, and *L* is the total length of the protein sequence (here, *L* = 4). 𝒵 is the partition function measured over all possible states in the Boltzmann distribution. *J* is a energetic coupling parameter that assign a statistical energy to any amino acid *a*_*i*_ (appearing in position *i*) and any amino acid *a*_*j*_ (appearing in position *j*) that is in principle unique for any given *a*_*i*_ and *a*_*j*_. *h* is a field parameter which is the statistical energy for any amino acid *a*_*k*_ (appearing in position *k*). If the parameters *J*_*ij*_ and *h*_*i*_ are learned accurately, this constitutes a trained Potts model where sampling from the underlying Boltzmann distribution reproduces the single (field) and coupled empirical frequencies of amino acids from the training multiple sequence alignment (MSA). Effectively, this would mean Equation 4 holds.

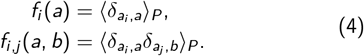

Where *f*_*i*_ (*a*) is the empirical frequency of amino acid *a* being in position *i* and *f*_*ij*_ (*a, b*) is the empirical frequency of amino acid *a* being in position *i* while amino acid *b* is in position *j*. We thus require an MSA to train on that describes length four segments of protein α-helices.

To build this MSA, we collected bioinformatic data using the entire SwissProt Databank [66] with corresponding AlphaFold 2.0 [67] structures. We then clustered the resulting sequences using mmSeqs2 [68] (`createdb, cluster -s 7.5, createsubdb`, and `createtsv` commands). From the mmSeqs2 clustering, we took a set of representative sequences (in total 56,474 structures, see GitHub repository), each of which has a predicted AlphaFold 2.0 structure. By using DSSP [65], we extracted all groups of four consecutive α-helix (‘H’) residues to build an MSA. This analysis produced 5,813,468 sequences, each of length four, corresponding to protein α-helix segments. From this MSA, we normalized for amino acid expression bias (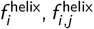 versus 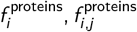 in Equation 5, where 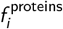 and 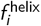 were from the clustered mmSeqs2 sequences) and then calculated empirical frequencies for Potts model training (*f*_*i*_ and *f*_*i,j*_ in Equation 6, Extended Data Fig. 2a).

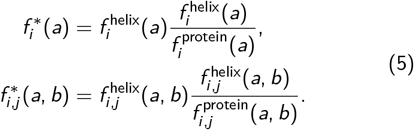

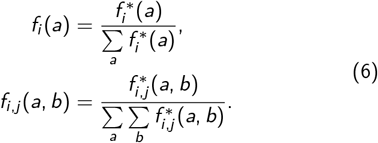

To build our Potts model, we utilized Boltzmann machine learning with a custom-built C++/CUDA code. With the empirical frequencies from the MSA providing our target distribution, we applied direct gradient descent and a basin-hopping approach during training. Equation 7 describes the direct gradient descent for updating Potts model parameters:

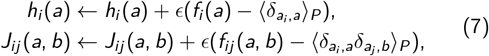

where *ϵ* controls the speed of gradient descent.

The initial values for each training run were set to *h*_*i*_ (*a*) = − log *f*_*i*_ (*a*) and all *J*_*ij*_ (*a, b*) = 0.0. A training iteration constituted 10,240,000 total samples of the Boltzmann distribution built across 100 blocks each with 512 threads running on an NVIDIA A100 GPU. For each iteration, each thread underwent a ‘burning-in’ period with 10,000 mutations to a sequence of four amino acids uniquely tracked by each thread. Mutations were accepted based on the Metropolis criterion. Thereafter, each thread produced 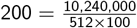 sequences to add to the sampled MSA with 1,000 mutations between each added sequence according to a Markov chain Monte Carlo scheme. After each iteration, model parameters were updated according to Equation 7.

A complete training run for the Potts model consisted of three different phases—optimization phases, search phases, and a final optimization. Optimization phases consisted of 5000 iterations of parameter updates at a temperature of *T* = 1.0, an *ϵ* = 0.25, and a maximum descent (an enforced upper limit of each parameter’s change) equal to 0.1. Between each optimization phase, a search phase (i.e., basin hopping) was undertaken for 100 steps at a temperature of *T* = 25.0, and an *ϵ* = 1.0 with no maximum parameter change enforced. In total, there were ten optimization phases with search phases between each. A final optimization phase was then performed for 5000 steps with *T* = 1.0, *ϵ* = 0.025, and a maximum descent of 0.1. During the training process, the parameter set that provided the greatest agreement with the empirical frequencies was saved. The final parameters for the Potts model were taken as the average of the best parameter sets from ten independent training runs.

#### 5. REST2 Enhanced Sampling Simulations for Potts Model Validation and Gaussian Process Training Data

We simulated peptides at an atomistic resolution to build a dataset for validating our Potts Model and parameterizing Mpipi-Helix well depths. For the former, we aimed to test our hypothesis that sequences with a lower Potts score (i.e., a higher statistical likelihood of being an α-helix) have a higher propensity to form a helix. To this end, we separately simulated 75 peptides each 11 residues in length with the ff19sb force field [54] using replica exchange solute tempering 2 (REST2) [69] in GROMACS [56]. We chose the sequences for these peptides by considering all possible combinations of the 20 canonical amino acids in sequences of four. These sequences were arranged onto a residue size versus hydrophobicity space (using the Urry scale [70]) and clustered using kmeans clustering with 25 cluster centers (Extended Data Fig. 2b). From each cluster we selected three sequences by fitting the distribution of Potts scores to a Gaussian and selecting: the sequence closest to the mean of the distribution, closest to plus two standard deviations from the mean, and closest to minus two standard deviations from the mean (Extended Data Fig. 2c). Each peptide was then constructed from repeating the sequence 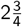 times (e.g., an ABCDABCDABC format). After simulating and removing sequences with poor convergence, we calculated the Potts score and the natural logarithm of the ratio between the number of helical frames versus coil frames for every set of four consecutive amino acids in the peptides. We find a strong linear correlation between the Potts score and the helical propensity of the peptides (Extended Data Fig. 1f). This linear correlation provides evidence that our Potts model is well constructed and can accurately describe differences in the helical characteristics of sequences of four amino acids.

The details of the REST2 simulations are as follows. Inputs for the simulations were generated from CHARMMGUI [59] using PDB files generated from PyMOL [58] that were initialized in an extended conformation. Each system was solvated with OPC water in 150 mM of NaCl and all peptides were uncapped. An initial minimization was performed via 5000 steepest decent steps with a force tolerance of 1000 kJ mol^−1^ nm^−1^ and positional restraints on the heavy atoms of the protein. A 1 ns *NVT* equilibration was then performed with a time step of 1 fs at 300 K using the vrescale thermostat with a time constant of 1 ps. Positional restraints were maintained on the heavy atoms of the protein during this equilibration step. After equilibration, eight temperature replicas were run at 300.00, 322.71, 347.14, 373.42, 401.69, 432.10, 464.81, and 500.00 K. These were simulated in the *NVT* ensemble using a timestep of 2 fs and the v-rescale thermostat with a time constant of 1 ps. PME was used for electrostatics and the cutoffs for van der Waals and Coulomb interactions were set to 0.9 nm. The LINCS constraint algorithm was used for bonds including hydrogen atoms. Sampling was run for 1.55 µs (12.4 aggregate µs), with the first 50 ns used for equilibration. Exchanges between adjacent replicas were performed every 10 ps.

#### 6. Gaussian Process Parameterization of Helical Well Depths

The final step in our parameterization process was to find a relationship between the Potts score of a sequence of four amino acids and a well depth in Mpipi-Helix. Here, we developed a Gaussian process (GP) workflow that iteratively simulated peptides in the Mpipi-Helix force field to replicate the helical content of peptides from the REST2 atomistic dataset and guest–host peptides measured using NMR in Moreau et al. [48] (Extended Data Fig. 1g,h,i). Within the workflow, each peptide was parameterized separately beginning with an initial guess of 5 well depth values aimed at recapitulating the helical propensity from the atomistic simulation or experiment (Extended Data Fig. 1g). These coarse-grained simulations were run in the *NVT* ensemble at 298.15 K using the Langevin thermostat with a damping of 100 ps and a timestep of 10 fs in LAMMPS [63]. The simulations were run with a 1 ns equilibration time and a 1 µs sampling time with simulation frames saved every 0.05 ns. After the simulations were completed, residues in the peptides were assigned helix or coil states on a frame-by-frame basis according to the Mpipi-Helix helix–coil assignment rule (see ‘Helix–Coil Classification’ below).

The helical fractions were further scored using a mean squared error comparison against the target propensity. Thereafter, a GP was trained on all the cumulative data accrued for the peptide to predict scores for the possible well depths (a grid 0.00 to 5.00 at a 0.01 spacing). For GP training, we used an Adam optimizer with a learning rate of 0.1 and 50 training iterations. The GP was defined by a zero mean function and an RBF kernel with a Gaussian prior on the length scale (mean of 0.5 and standard deviation of 0.1). Inputs into the GP were scaled using scikit-learn’s standard scaling function [71].

From the resulting scores, an acquisition function (expected improvement or upper confidence bound) was paired with a negative bias acquisition strategy to choose a new batch of five points to simulate (Extended Data Fig. 1h). For the first 100 batches, an expected improvement acquisition function was used (Equation 8, where *W* is the well depth, *ϕ* is the cumulative distribution function, *ϕ* is the standard normal probability density distribution, *µ*(*W*) is the mean predicted value at *W, σ*(*W*) is the predicted standard deviation at *W, f* (*W* ^+^) is the highest scoring *W* value predicted so far, and EI(*W*) is the expected improvement acquisition function evaluated at *W*). For the next 100 batches an upper confidence bound acquisition function was used (Equation 9, where *ξ* is set to 0.1 and UCB(*W*) is the upper confidence bound acquisition function evaluated at *W*). During batch selection for each acquisition function, well depth points were chosen sequentially, with a negative bias applied to the acquisition function value for all well depths previously chosen points in the batch. This negative bias took the form of a Gaussian with mean centered at the chosen well depth and a random variance selected between 0.005 and 0.015.

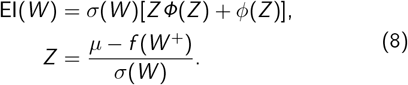

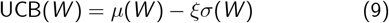

This process terminated after 200 batches (1,000 data points) were completed and a final fitting was performed by the GP to determine the best well depth value (Extended Data Fig. 1i). The only exceptions were for the GTNI, GGKE, and GAAA peptides from the REST2 datasets. These peptides were sampled with an additional 40 batches with the UCB acquisition scheme due to insufficient sampling in the region of the maximum score. In these 40 batches, sampling was restricted to the region of 3.0– 3.21 kcal/mol. For all final predictions, the input data had no scaling, the GP had a constant mean, and an RBF kernel with a length scale enforced to be between 0.2 and 1.0 was used. This prediction was performed using an Adam optimizer with a learning rate of 0.1 and 100 iterations.

All GP-based code in this work was written in the package GPyTorch [72].

#### 7. Helix–Coil Classification

Two methods were used for helix–coil classification in this study. The first method was used exclusively for parameterization of the Mpipi-Helix model within the Gaussian process workflow. This classification followed two steps. First, it was calculated whether a given set of four residues acting under the α-helix potential were inside the peak of the helical barrier (1.33 ≤ *θ* ≤ 1.895 and 0.246 ≤ *α* ≤ 1.558). If this was the case, the central two residues would be classified as helical. Second, Equations 10 and 11 proposed by Tozzini et al. [73] was iteratively solved to determine *ϕ* and *ψ* backbone angles, with *γ*_1_ = 20.7^°^, *γ*_2_ = 14.7^°^, *τ* = 111^°^.

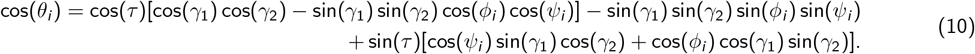

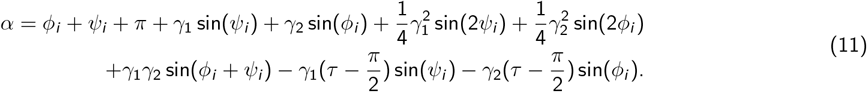

If these equations could be solved self-consistently with the solutions of *α* and *θ* agreeing to within 0.1 radian of their actual values, a given residue would be assigned a helix if −30.0 *> ϕ >* −100.0 and −7.0 *> ψ >* −67.0. The final step was to combine each classifier by taking a residue to be helical if it was classified as such my either method with residues being coil otherwise. We developed and implemented this classification scheme before the cg2all software [74] was published.

The second method leveraged the cg2all backmapping software [74] and was used for all validation and subsequent Mpipi-Helix simulations following parameterization (e.g., all data collected in the manuscript figures outside of Extended Data Fig. 1 and 2). In this scheme, the first step was to take the output LAMMPS trajectory for a simulation and convert it into .dcd format using MDConvert [75]. Second, cg2all [74] was used to backmap into an atomistic trajectory. Finally, MDAnalysis [76, 77] was used to extract the *ϕ* and *ψ* backbone protein angles from the backmapped trajectory. The classifier to determine the helical region from these backbone angles was then −30.0 *> ϕ >* −100.0 and −7.0 *> ψ >* −67.0 for the helical state.

#### 8. Obtaining the Helical Well Depth versus Potts Score Trend

The final Potts score to well depth trend was calculated using a two part linear fitting on the final data from the Bayesian optimization workflow (raw data shown in Extended Data Fig. 2d). First, the intercept was determined using a linear fit on top of the experimental NMR data (Extended Data Fig. 1j), as that data describes helical peptides well in a strong helical context (e.g., the alanine context). However, this data does not perform well at describing peptides with low helical propensity due to alanine amplifying the effects of helix-breaking residues on unfolding [78, 79]. We therefore found the slope of our linear relationship by fitting to the combined REST2 and NMR datasetss, but after shifting the REST2 datasets to lower well depths to maximally overlap with the NMR data (Extended Data Fig. 2e,f). Specifically, we calculated the shift required to maximally overlap the minus two standard deviations dataset from the REST2 peptides on the NMR dataset. Our shifting followed two steps. First, we removed the GAAA sequence from the minus 2 standard deviation group as well as the H, G, and P guest–host constructs from the Moreau group as they were the largest outliers within the two datasets. Second, we performed a shifting whereby we evaluated the KDE overlap between the two sets of points using scipy’s `scipy.stats.gaussian_kde` with a bandwidth of 0.2 and `scipy.integral.simps` functions. The best overlap was selected from shifting offsets tested between 0.0 and 1.0 in an interval of 0.01 which was calculated to be 0.37 kcal/mol. We argue that this shifting was necessary due to the ff19sb force field systematically overpredicting helical propensity when directly compared to the Moreau NMR dataset [54]. Finally, as we were concerned with only sequences that we know will make helices, we removed strong helix-breakers prior to the linear fitting for our trend’s slope (Extended Data Fig. 2g). The final trend is provided in Extended Data Fig. 1j as *W* = 3.45 − 0.04*P* where *W* is the well depth in kcal/mol and *P* is the Potts score. Using this trend, we can now input a sequence of four amino acids into our Potts model and then calculate the well depth to get a sequence-dependent description of the residues in the Mpipi-Helix force field (Extended Data Fig. 1k).

### C. Mpipi-Helix Validation

#### 1. Validation Against Atomistic Simulations

We validated the Mpipi-Helix force field against atomistic simulations and experimental data. We began by simulating alanine-based peptides in the ff19sb [54], ff14sb [80], and CHARMM36m [81] force fields. We ran vanilla MD simulations of guest–host 11-residue peptides of the form A_5_XA_5_, where X represents one of the 20 canonical amino acids. Additionally, we simulated A_21_ and A_4_(A_4_R)_3_A_2_. We further ran well-tempered metadynamics of A11 to enhance convergence of its free energy surface. All simulations were performed in the *NVT* ensemble at 298.15 K.

From the results of these simulations, we see that Mpipi-Helix does an excellent job of recapitulating the thermodynamic features of alanine-based peptides (Extended Data Fig. 3a,b). For example, the prominent features of the FES of the A_11_ peptide are well captured, including the helical free energy basin and the broad coil state (Extended Data Fig. 3a). Matching the size and shape of each of these regions, and critically that of helical well, validates the form of our helical potential, the width of our helical well, and the location of our helical barrier (Extended Data Fig. 1b,c).

Mpipi-Helix also reproduces the trends in residue-specific helical fractions of alanine-based peptides (Extended Data Fig. 3b). This is demonstrated for A_5_EA_5_, A_5_LA_5_, A_5_QA_5_, and A_5_RA_5_ where Mpipi-Helix closely agrees with the rank ordering of helical content for each of the constructs compared to ff19sb (Extended Data Fig. 3b). Further, when all of the A_5_XA_5_ peptides are taken in aggregate and compared with Mpipi-Helix, Mpipi-Helix reproduces variations in helical fractions from the atomistic force fields (Extended Data Fig. 2h–j). Notably, while Mpipi-Helix tends to over-predict the helical content of these alanine-based peptides, this was by design as alanine has been shown to have a comparatively larger response to helix-breaking mutations when compared to other helical residues [78, 79]. Finally, Mpipi-Helix also reproduces residue specific helical fractions for longer peptides despite only being trained on 11-residue peptides. For the A_21_ and A_4_(A_4_R)_3_A_2_ sequences Mpipi-Helix’s predictions closely agrees with the atomistic force fields. This suggests that Mpipi-Helix captures the cooperative effect produced by longer helices which result in higher helical fractions [82].

#### 2. Atomistic Metadynamics Simulations of A11

Metadynamics simulations of A11 in ff19sb [54], ff14sb [80], and CHARMM36m [81] were performed in GROMACS [56]. Input configurations of the uncapped peptide were made using PyMOL [58] in an extended conformation. Simulation inputs were generated using CHARMMGUI [59] at 150 mM NaCl salt. All systems were first minimized using 5000 steepest descent steps with a force tolerance of 1000.0 kJ/mol/nm, and positional restraints on the heavy atoms of the protein. A subsequent equilibration step was performed for 100 ns with a timestep of 2 fs at 298.15 K using a v-rescale thermostat with a 1 ps damping constant. Cutoffs for van der Waals and Coulomb interactions were set to 0.9 nm. Long-range electrostatics were calculated using a PME mesh grid. Bonds involving hydrogen were constrained with the LINCS algorithm [60]. Well-tempered metadynamics [30] were conducted at the same conditions, implemented via the PLUMED [57] plugin for GROMACS. The biased collective variables were the distance root-mean-squared deviation to a perfect helix and the radius of gyration. The bias factor was set to 5, with a hill deposition rate of 10 ps and hill deposition size of 0.025 kcal/mol. Sampling was performed for 5 µs and convergence was assessed using the free energy surfaces which did not significantly change after 4 µs.

#### 3. Atomistic and Mpipi-Helix Simulations of A_5_XA_5_, A_21_, and A_4_(A_4_R)_3_A_2_

Atomistic simulations of A_5_XA_5_ peptides in ff19sb [54], ff14sb [80], and CHARMM36m [81] were run in the *NVT* ensemble at 298 K using GROMACS [56]. Inputs were built with CHARMM-GUI [59] with input PDB files built in PyMOL [58] with peptides started from an extended structure. As before, an initial minimization was conducted via steepest descent with positional restraints on all heavy atoms in the protein. Equilibration was performed with positional restraints for 125 ps at a 1 fs timestep. For the ff19sb simulations, a Nosé–Hoover thermostat was used with a time coupling of 1 ps. For both ff14sb and CHARMM36m, the v-rescale thermostat was used with the same time coupling as ff19sb. In ff19sb and ff14sb, van der Waals and Coulomb cutoffs were set to 0.9 nm with a PME grid for long-ranged electrostatics. CHARMM36m was similar with a 1.2 nm cutoff for van der Waals and Coulomb interactions. Finally, peptides were simulated for either a sampling period of 10 µs (11 length peptides) or 5 µs (21 length peptides), taking three even blocks for the calculation of errors using block averaging. In all simulations, the LINCS algorithm constrained bonds involving hydrogen atoms [60]. The atomistic simulations of A_21_ and A_4_(A_4_R)_3_A_2_ were run in a similar manner at 300 K.

Mpipi-Helix simulations of A_5_XA_5_ were performed at 298.15 K in the *NVT* ensemble using a Langevin thermostat with a damping constant of 100 ps. The aggregate simulation time for each peptide was 750 µs comprised of 300 2.5 microsecond simulations. Simulations were conducted in LAMMPS [63]. Each simulation had an equilibration period of 10 ns with frames saved every 1 ns. Simulations of A_21_ and A_4_(A_4_R)_3_A_2_ were performed in a similar manner but with 3 independent replicates of 10 µs in length for each peptide and frames saved every 1 ns.

#### 4. Validation Against Helical Fractions Predicted from NMR Measurements

We further validated Mpipi-Helix against helical fractions derived from NMR measurements. We extracted chemical shifts of peptides from the Biological Magnetic Resonance Databank (BMRB) and used the program delta2D (d2D) [83] to acquire predicted residue-specific helical fractions. We then used the helical fractions for each peptide to assign helix and coil regions of the protein sequence. Thereafter, we simulated each peptide in our Mpipi-Helix force field (see SI).

Our Mpipi-Helix simulations show near-quantitative agreement with the helical propensities predicted from NMR data (Extended Data Fig. 3c). For example, we capture the C-terminal domain helical domain of TDP43 as well as the effect of a A326P mutation within the helical region. For ANXA11 protein, we obtain excellent agreement with the first helical peak from the NMR measurement and also reproduce a reported second peak from recent published data [84]. For AR, we accurately reproduce the high helical content of the LLLL-polyQ domain.Further, in the rest of the proteins (CREB1, FCP1, GAB1, and HCV-C), Mpipi-Helix performs very well at capturing peak height and shape. This serves as critical validation of our Mpipi-Helix force field as these NMR measurements were performed on complex biological protein sequences. Further, the helical content from the NMR chemical shifts are influenced by the interaction of disordered components of the protein with the helical domain, which is precisely what we measure with our coarse-grained model when we simulate a diverse range of helical domains within chemically-diverse condensates. This suggests our force field is correctly capturing the interactions between disordered and structured domains and can accurately portray the biophysical behavior of α-helix domains within biomolecular condensates.

#### 5. Extracting Helical Fractions from NMR Chemical Shifts

Helical fractions were extracted from BMRB using the delta2D software [83]. This produced predicted helical state populations on a per-residue basis. To apply these to Mpipi-Helix simulations, we developed a smoothing procedure which aided in flattening out any noise in the predicted shifts. We first calculated the smoothed helical fraction of residue *i* in each peptide as the average helical fraction between residue *i* − 2 and *i* + 2. We then added helical caps of 3 additional helical residues to either side of each stretch of helical residues such that every residue within the original helical domain would have maximal helical potentials to describe its helicity. This constituted our helical segments along the protein sequence from the BMRB entry. The rest of the residues were left disordered.

#### 6. Mpipi-Helix Simulations of NMR Measured Peptides

Mpipi-Helix simulations were run in the *NVT* ensemble with a Langevin thermostat using a damping constant of 100 ps at various temperatures corresponding to the NMR measurements. The timestep for the simulations was 10 fs and simulated in LAMMPS [63]. Each peptide was simulated using 3 independent replicates for 10 µs with frames saved every 1 ns. Classification of helix and coil residues was determined using the cg2all [74] procedure.

### D. Parameterization of a Coarse-Grained Model for PEG Crowder

For the crowded environment, we parameterized an excluded volume polymer model to qualitatively represent the polymer polyethylene glycol (PEG). In particular, we represented PEG in a coarse-grained manner with interaction sites localized to the oxygen in the monomer structure. We determined the volume of the PEG monomer using Protein Volume 1.3 [85] and represented pairwise interactions involving PEG beads with the Weeks-Chandler-Anderson (WCA) potential: *ϵ* = 0.586 kcal/mol (approximately *k*_B_*T* at room temperature) and *σ* = 4.137 Å with an arithmetic mixing rule for the *σ* values when interacting with protein residues. The molecular structure of PEG was represented as a disordered polymer using harmonic bonds between adjacent monomers with parameters set to match OPLS simulations of PEG.

### E. Simulations and Analysis of Helical Peptides

#### 1. Simulations in Dilute, Crowded, and Condensate Environments

All coarse-grained simulations were performed via Mpipi-Helix in LAMMPS [63]. Simulations of the helical peptides A11, AER, AL, and AQ in the dilute environment were performed in the *NVT* ensemble using a Langevin thermostat with a 100 ps damping constant. Simulations of the helical peptides within the condensate environments (FYAAYF, NYQQYN, and RYAAYE) were run in the *NPT* ensemble using a Langevin thermostat and a Berendsen barostat at 0 atm, each with a damping constant of 100 ps. Simulations of the helical peptides in crowder were performed likewise but with a pressure of 71.3 atms to match the concentration of the FYAAYF condensate. The timestep for all simulations was set to be 10 fs. For all simulations, an initial minimization using the fire min style from LAMMPS in four incremental minimization steps was performed, incrementing from 0.00000001 to 0.00001 to 0.1 to 10 fs timesteps with a 1e-08 (kcal/mol)/Å force tolerance over 1000 max iterations or 100000 force evaluations. For the *NPT* simulations, an initial 10 ns compression was performed, at 10.0 atms for the condensates and 71.3 atms for the crowder. For the dense environments, a 100 ns equilibration followed. In the dilute environment, a 10 ns equilibration was used. For all simulations, sampling was performed for 2.5 µs with frames of the helix saved every 1 ns and frames of the entire simulation saved every 5 ns. In total, 300 simulations were run for each helical peptide and environment/condensate pair for a total of 750 µs in each condition. Finally, the chain numbers for the simulations were 144 chains, 141 chains, 145 chains, and 144 chains for the FYAAYF, NYQQYN, RYAAYE, and PEG simulations, respectively. For FYAAYF, NYQQYN, and RYAAYE, these chain numbers were selected to create a 60^3^ nm^3^ simulation box as determined from density profiles taken from direct coexistence simulations (see SI).

#### 2. MSM Construction and PCCA+ Metastable States

We constructed Markov state models (MSMs) for the helical peptides in each environment using the PyEMMA software [86]. For our MSMs, we utilized time-lagged independent component analysis on input features, which constituted pairwise distances between non-bonded residues as well as all the bond and dihedral angles along the coarsegrained peptide chain. Thereafter, the MSM was made with 1,000 cluster centers and a lag-time of 2 ns. We also coarse-grained the MSM into four metastable states using PyEMMA’s PCCA+ spectral clustering functionality [36]. Validation of the MSMs were performed using the Chapman-Kolmogorov test and VAMP2 scores on the various hyperparameter choices (see SI).

#### 3. MSM Analysis

All quantities extracted from MSMs were taken as the average of 100 independent instantiations of the MSM with unique random number seeds used during clustering. We leveraged PyEMMA’s bayesian markov model for each iteration and calculated error as the standard error between model constructions combined with the uncertainty within each model. In this manner we calculated the average relative free energies between metastable states from stationary distributions, mean first passage times using PyEMMA’s mfpt command, and implied timescales using sample mean(‘timescales’).

#### 4. Viscosity and Diffusion Coefficient Calculations on Co-Condensate Proteins

Viscosity and diffusion coefficients for the FYAAYF, NYQQYN, and RYAAYE co-condensate proteins were calculated from *NVT* simulations in LAMMPS [63]. In particular, we ran 5 independent simulations for each condensate using the following procedure. First, we replicated a single chain 6 × 6 × 6 times and simulated for 50 nanoseconds using a Nosé–Hoover thermostat and barostat. The thermostat was set to 298.15 K and the barostat was set to 1.0 atms each with a 100 ps damping constant. Following this, another *NPT* relaxation was performed at 0 atms but otherwise in an identical manner. Thereafter, an *NVT* equilibration step followed for 100 ns again using the Nosé– Hoover thermostat with a 1 ps damping constant. Finally, a 2.5 µs sampling period was run with under the same *NVT* conditions, tracking both the mean-squared displacement of the protein chains in the condensates (LAMMPS msd command) and the autocorrelation of the stress tensor components (`LAMMPS ave/correlate/long` command applied to normal and tangential stresses).

Following these simulations, we calculated the viscosity for each independent run using the Green–Kubo formalism for isotropic *NVT* simulations as done in Ref. [87]. Additionally, we also calculated the diffusion coefficient for the co-condensate protein chains using the formula MSD = 6*Dt*. Here, MSD is the mean-squared displacement and *D* is the diffusion coefficient.

#### 5. Pairwise Enthalpy Calculation

We calculated pairwise enthalpies by computing the contact energies between each helical peptide and the co-condensate proteins in our simulations. In particular, we first calculated which frames corresponded to a helix and coil configuration utilizing the metastable states from our MSM. Thereafter, we computed the average pairwise contact energies for each state and calculated their difference within each simulation. The error we report is the standard error across all 300 simulation replicates for each helical peptide–condensate pairing. Note that we assume the pressure–volume contribution to the enthalpy is 0 as the ensemble average pressure, set by the barostat in our system, is 0 atms.

#### 6. Close Contacts and Radial Distribution Function Analysis

Close contacts were analyzed using a custom in-house script. Specifically, we considered the average number of close contacts between the central seven residues in each helical peptide and the residues within co-condensate proteins. A contact was assigned if the pairwise interaction between a helical peptide residue and a co-condensate protein residue was within the *R*_min_ of the pairwise Wang–Frenkel potential. Frames were averaged separately for the helical peptide as coil and helix based on the PCCA+ four metastable states (N-terminal helix and C-terminal helix were not included in either of the averages).

Radial distribution functions were calculated using a custom in-house script. For each of the seven central residues in each helical peptide, a radial distribution function was calculated against each co-condensate residue type. The frames were classified as helix or coil depending on the state of the helical peptide as classified by the PCCA+ metastable states (again ignoring N-terminal and C-terminal helix states). The data from the central seven residues was then collapsed onto a single RDF by normalizing the bin distances using the *R*_min_ of that residue’s pairwise interaction with the co-condensate residue type. Finally, the height of the first peak of the RDF was extracted (except in the case of alanine which had no discernible peaks). In the case that no peak existed for the helix state in the RDFs, the value closest to *r/R*_min_ was taken for comparison with the coil state peak. For both the close contacts and the RDFs, the error was calculated using standard errors over all independent simulation replicates (300 for each pairing).

### F. Simulations and Analysis of TDP43, ANXA11, and AR α-Helix Domains

#### 1. Direct Coexistence Simulations of Co-Condensate Proteins

Coarse-grained direct coexistence simulations in the slab geometry were performed via Mpipi-Helix for the co-condensate proteins in LAMMPS [63]. This includes the proteins FUS, hnRNPA1, TIA1, RPB1, MED1, and BRD4. The chain numbers for each simulation matched those used in the *NPT* condensate simulations (see below). To set up the simulations, two initial *NPT* compression steps with a Langevin thermostat at 200 K under a 100 ps damping constant and a 10 fs timestep were performed. For the first step, a Berendsen barostat at a pressure of 5 atms and a damping of 10 ps was used to compress each system. For FUS, hnRNPA1, and TIA1, this step lasted for 100 ps whereas for RPB1, MED1, and BRD4 this step lasted for 200 ps. Next, for all systems, the barostat was set to 1 atm with a damping of 100 ps for an additional 500 ps. The final simulation frame the compression was used as the initial starting point for the direct coexistence simulation in the slab geometry.

Once the compression was completed, each system was initialized in a slab geometry with the box length in the z dimension extended to be five times that of x and y. For RPB1, MED1, and BRD4, the simulation box was also filled with a 10% volume of PEG-8000 (e.g., PEG chains of 176 monomers in length) where half of the total number of PEG polymers was placed on each side of the condensate. Each condensate was then simulated for 1.25 µs where 250 ns was used as equilibration time. Frames and densities were recorded every 1 ns and used to generate average density profiles for each condensate.

#### 2. Simulations in Dilute and Condensate Environments

Dilute phase *NVT* simulations were run for each of TDP43, ANXA11, and AR using the Langevin thermostat with a damping constant of 100 ps and a timestep of 10 fs at 298.15 K in LAMMPS [63]. Systems were first equilibrated for 50 ns and then sampled for 2.5 µs with frames recorded every nanosecond. In aggregate, 750 µs of simulation time was accrued for each protein via 300 independent simulation replicates.

Condensed phase simulations were performed in the *NPT* ensemble utilizing a Langevin thermostat and a Berendsen barostat, each with a 100 ps damping constant and a 10 fs timestep at 298.15 K in LAMMPS [63]. The pressure for these simulations was 0 atms for TDP43 and ANXA11 in FUS, hnRNPA1, and TIA1 co-condensates, whereas it was set to 2.7 atm for RPB1, 19.6 atm for MED1, and 18.8 atm for BRD4 co-condensates in the AR systems. The non-zero pressures for the AR–condensate systems were necessary as none of the co-condensate proteins phase separated on their own without PEG. We thus determined the pressure that matched the protein’s direct coexistence densities in 10% PEG, as discussed above. Finally, the chain numbers for the simulations were 64 chains, 75 chains, and 104 chains for the FUS, hnRNPA1, and TIA1 co-condensate proteins in the TDP43 and ANXA11 systems. For the AR systems, the chain numbers were 133 chains, 80 chains, and 74 chains for RPB1, MED1, and BRD4 co-condensate proteins. For all systems, the number of chains of co-condensate proteins was set such that there were approximately an order of magnitude more residues in the simulation box from the co-condensate proteins than the α-helix containing protein. All protein sequences are provided in the SI.

For the dense phase simulations, all simulations began with a 100 ns equilibration period followed by 2.5 µs of sampling. We collected 500 aggregate µs consisting of 200 independent simulations for each protein–condensate pairing with TDP43 and ANXA11. For the AR–condensate systems, we collected 300 aggregate µs via 120 independent simulations.

#### 3. Wasserstein Distance Calculations Between Free Energy Surfaces

Wasserstein distances were calculated by mapping the dense phase free energy surfaces of the α-helix domains in the condensates onto the dilute phase free energy surface. The Wasserstein distance was calculated using the emd2 function of the python package Optimal Transport [88]. The visualization of the distance transported in Extended Data Fig. 5a was produced using the `emd` function and plotting the associated transport plan.

#### 4. MSM Construction

MSMs were constructed for each TDP43, ANXA11, and AR condensate pairing as well as in the dilute environment. All models were constructed using PyEMMA [86], leveraging identical input features as the helical peptides described above applied to the α-helix domains within the proteins. For these models, we used a lag time of 5 ns and 1500 cluster centers. The input features, which had time lagged independent component analysis applied to them, were the pairwise distances between non-bonded residues in the αhelical domain, as well as the bond and dihedral angles along the α-helical domain. We validated our MSMs using the Chapman–Komolgorov test and VAMP2 scores of various instantiations of MSM hyperparameters (see SI).

#### 5. Implied Timescale Analysis

The slowest implied timescale was extracted analogously to the method used for the helical peptides above. The only difference was that the timescale was averaged over 25 independent model constructions instead of 100. Additionally, to demonstrate that the slowest timescale was a decay between the folded state and unfolded state, during each MSM construction we collected the average helical content of the structures mapped to the clusters with the highest and lowest values of the second right eigenvector. The error bars in these helical fractions represent the within cluster standard deviation combined with the standard error across independent model constructions.

#### 6. Close Contact Analysis

Close contacts between the α-helical domains and the co-condensate proteins were calculated by differentiating helix and coil residues within the domains on a frame-by-frame basis. Specifically, we considered a residue to be helical if it was in a helical configuration and had neighboring helical residues on either side (e.g., a classification of H<u>H</u>H). This corresponds to the central residue participating in the equivalent of a single hydrogen bond in the helix. Otherwise, the residue was considered to be in a coil configuration. Finally, we calculated separate per-residue averages in the helix and coil states and averaged across all residues of the same types. Errors were then computed across simulation replicates as a standard error.

#### 7. Residue-Type Contact Maps

Equations 12 and 13 were used for contact map construction in the residue-based contact maps shown in Fig. 4:

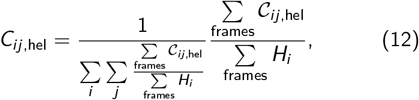

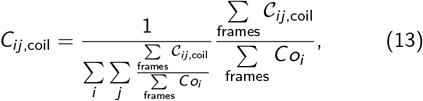

where *C*_*ij*,hel_ is the normalized contact frequency between residue *i* of the α-helix domain and *j* of the co-condensate proteins when *i* is in a helical configuration. Analogously, *C*_*ij*,coil_ is the same but in this case residue *i* is in a coil configuration. 𝒞_*ij*,hel_ and 𝒞_*ij*,coil_ are indicator functions which are 1 if residue *i* is in a helical/coil state and there is a contact between *i* and *j* in the co-condensate proteins. *H*_*i*_ and *Co*_*i*_ are indicator functions that are 1 if residue *i* is in a helical/coil state and 0 otherwise. We classify a contact as being within the *R*_min_ of the pairwise interaction potential between *i* and *j*. To produce the residue type contact maps we group residue–residue interactions based on the following categories: special (G, P, C), polar (S, T, N, Q), negative (D, E), positive (R, H, K), aliphatic (A, V, I, L, M), and aromatic (F, Y, W). Finally, the difference in contacts between the helix and coil states are produced by subtracting the resulting contact maps.

### G. Molecular Renderings

All renderings were made in the ChimeraX [89] and Ovito [90] softwares.

## V. ACKNOWLEDGMENTS

We thank Dr. Alina Emeilianova for GAFF2 PEG parameters for atomistic simulations. We are also grateful to Prof. Michael Webb and Shannon Zhang for discussions and aid in making a coarse-grained model for PEG. We thank Prof. Eric Dufresne, the Dufresne Lab members, Dr. Dilimulati Aierken, Dr. David Kuster, and Virginia Jiang for discussions and feedback on the manuscript, as well as Profs. Dimitrios Fraggedakis, William Jacobs, and Jared Toettcher for insightful discussions. We also thank Philippe Baron for helpful discussions concerning the application of MBAR to calculating free energy surfaces. The authors thank all members of the Joseph Group for their feedback on the work. J.A.J. acknowledges research support from the Chan Zuckerberg Initiative DAF (an advised fund of Silicon Valley Community Foundation; grant 2023-332391), and the National Institute of General Medical Sciences of the National Institutes of Health under Award Number R35GM155259. The content is solely the responsibility of the authors and does not necessarily represent the official views of the National Institutes of Health and other sponsors. The simulations reported on in this work were performed using the Princeton Research Computing resources at Princeton University, which is a consortium of groups led by the Princeton Institute for Computational Science and Engineering (PICSciE) and Office of Information Technology.

## VI. AUTHOR CONTRIBUTIONS

Conceptualization: NH and JAJ; Method development: NH and JAJ; Investigation: NH; Visualization: NH; Funding acquisition: JAJ; Supervision: JAJ; Writing: NH and JAJ.

## VIII. DATA AND CODE AVAILABILITY STATEMENT

All data and code supporting the findings in this study, as well as sample simulation input and output files, are available at the Joseph Group GitHub repository: https://github.com/josephresearch/Condensates_Dictate_Protein_Folding/.

## X. EXTENDED DATA FIGURES

**Extended Data Fig. 1.**
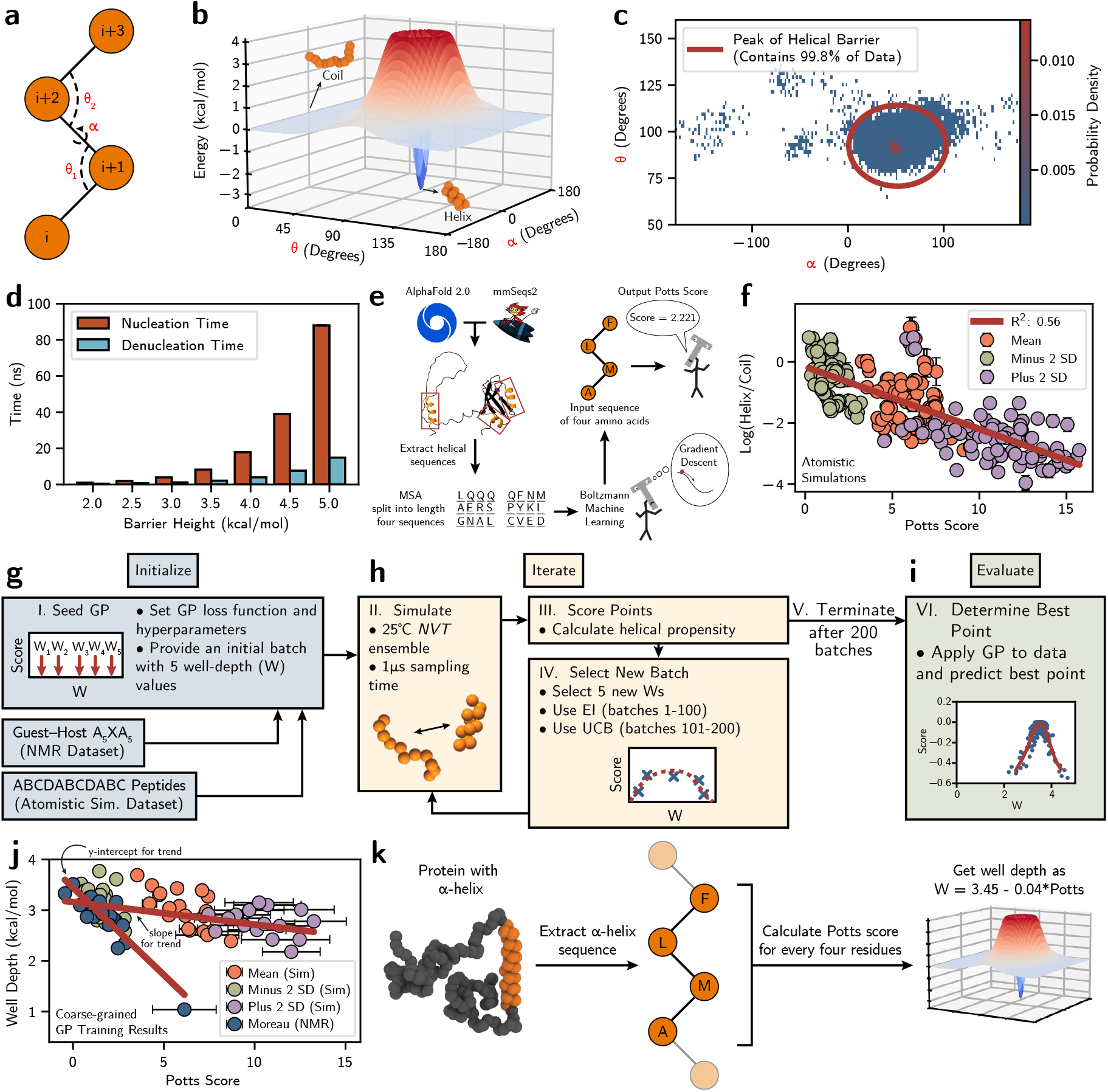
Mpipi-Helix: A structural potential to describe α-helices at a residue resolution parameterized via Bayesian optimization. **a**, The structural potential (Equation 1) operates on the dihedral and two consecutive bond angles of four consecutive amino acids. **b**, The potential is made from two Gaussian potentials centered at the geometry for a perfect helix. The positive Gaussian provides a kinetic barrier while the negative Gaussian biases helix formation. **c**, The width of the well and the location of the helix barrier was determined by overlap with α-carbon coordinates of helices extracted from PDB structures. **d**, The kinetics of the helix–coil transition is set by enforcing a barrier height of 4.0 kcal/mol. Standard errors are calculated across 150 independent 10 µs simulations of an A11 peptide. **e**, A multiple sequence alignment (MSA) is extracted from all helical sequences from predicted AlphaFold 2.0 structures in the SwissProt database that is clustered with mmSeqs2. Boltzmann machine learning with gradient descent is used on the MSA to produce a Potts model. **f**, The natural log of the helix to coil frames for segments of 4 consecutive amino acids within 75 separate 11 length peptides simulated at an atomistic resolution. Block averaging on 3 equally-sized blocks from 1.5 µs of simulation time at the coldest replica (298.15 K) is used for error. **g**,**h**,**i**, A Gaussian process workflow for parameterizing well depths to recapitulate helical propensities from atomistic simulations and NMR measurements [48]. **j**, The final well depth to Potts score trend combines the y-intercept from fitting the NMR experimental datasets with the slope of a fitting to all non helix-breaking sequences. Errors in Potts scores are calculated using the standard deviation of all Potts scores along the peptide sequence. **k**, Mpipi-Helix models α-helices in a sequence-dependent manner.

**Extended Data Fig. 2.**
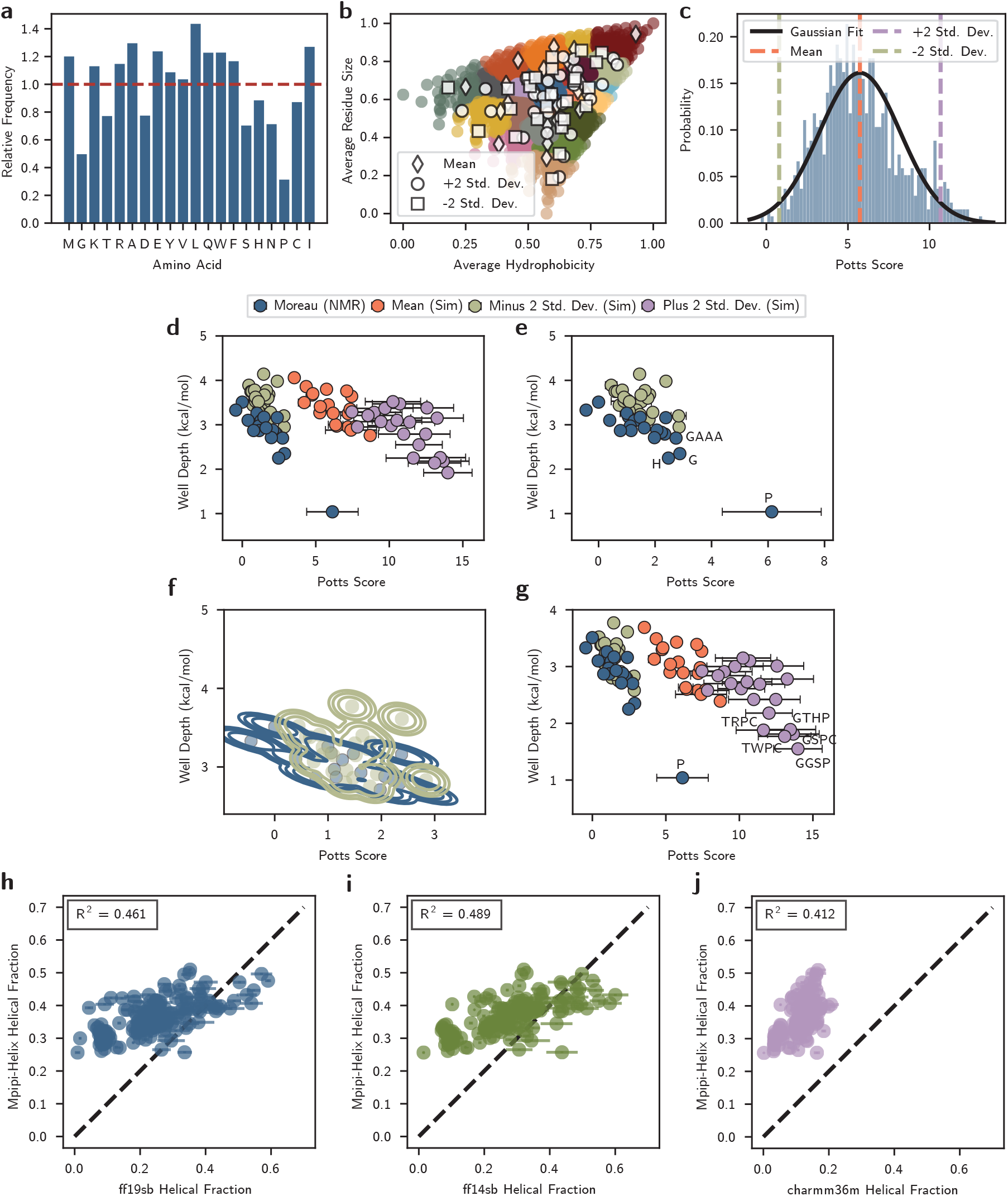
Bioinformatic data for Potts model development to capture sequence-dependent helical propensities used in Mpipi-Helix’s parameterization which recapitulates experimental and atomistic data. **a**, Potts model empirical frequencies are adjusted based on the relative background frequencies with which amino acids appear in α-helices versus in all protein sequences. **b**, All possible protein sequences of length four generated by combinations with replacement of the 20 canonical amino acids plotted against average residue size and average hydrophobicity (Urry hydrophobicity scale [70]). **c**, Mean, minus two standard deviations from the mean, and plus two standard deviations from the mean sequences are extracted by fitting a Gaussian to the Potts score distribution from each cluster in **c. d**, The final well depth and Potts score values for every sequence used in the Bayesian optimization parameterization workflow. Errors in Potts scores are plotted as the standard deviation of the Potts score over each sequence. **e**, Sequences that are ignored from the Moreau and minus two standard deviation dataset for the kernel density shifting process. **f**, Kernel density estimation is used to shift the atomistic simulation data from the minus two standard deviations dataset onto the sequences from the NMR experimental dataset. **g**, The helix-breaking sequences that are removed from the final slope fitting performed in Extended Data Fig. 1j. **h-j**, Parity plots of helical fraction in Mpipi-Helix versus the atomistic **h** ff19sb [54], **i** ff14sb [80], and **j** CHARMM36m [81] force fields.

**Extended Data Fig. 3.**
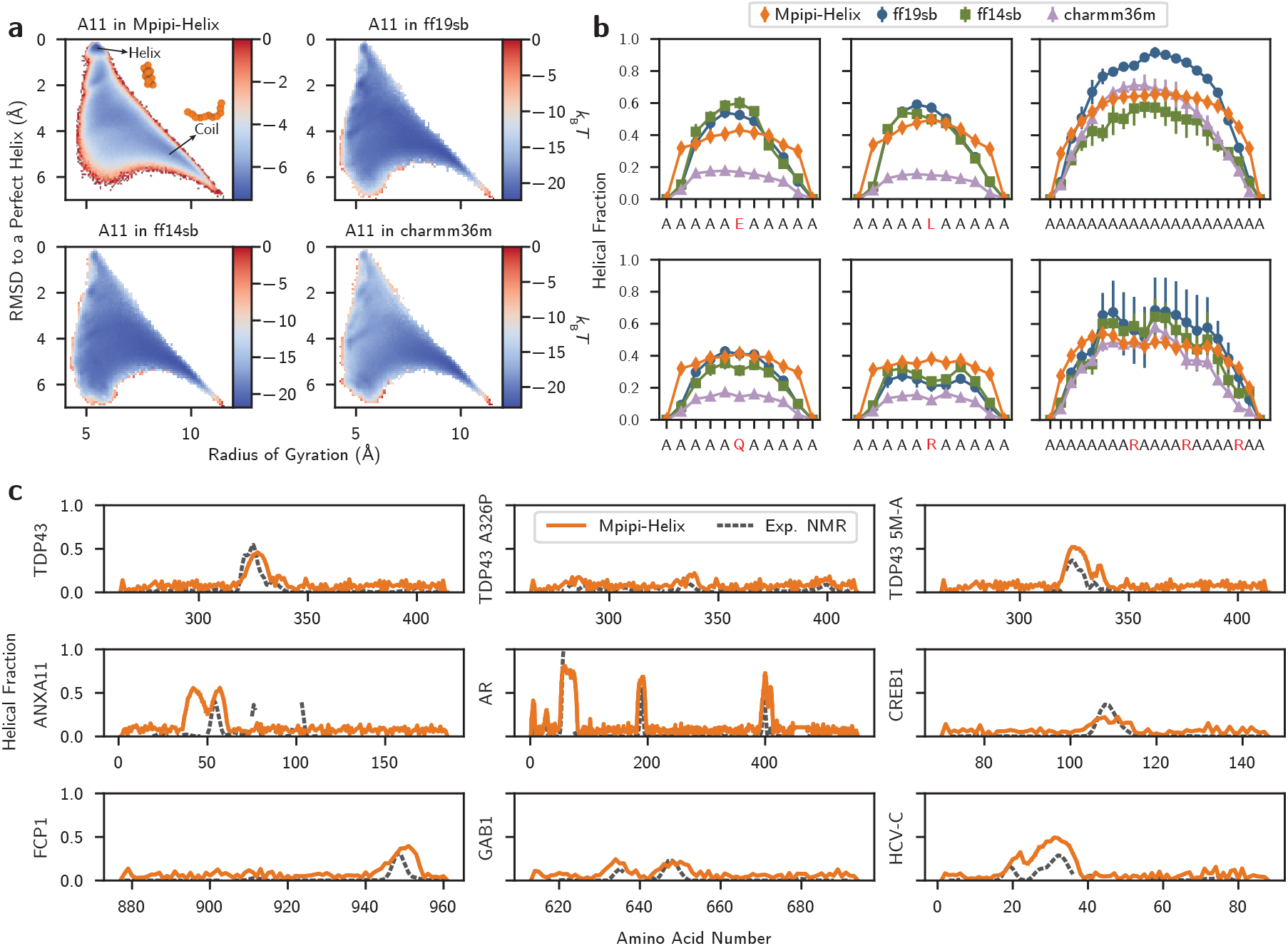
Mpipi-Helix accurately predicts folding of α-helices when compared to atomistic simulations and NMR measurements. **a**, The free energy surface of the A11 peptide in Mpipi-Helix, ff19sb [54], ff14sb [80], and CHARMM36m [81] force fields. **b**, Per-residue helical fractions of alanine-based peptides in Mpipi-Helix, ff19sb [54], ff14sb [80], and CHARMM36m [81] force fields. Errors for the atomistic simulations were determined using block averaging on 3 equally-sized blocks. The standard error across 300 simulation replicates is reported for Mpipi-Helix. **c**, Per-residue helical fractions of disordered proteins with α-helix domains. NMR-based helical fractions were determined using delta2D software [83]. The standard error across 3 simulation replicates are reported for Mpipi-Helix. (BMRB codes: Row 1––26823, 52060, 26828. Row 2–52944, 51479 + 51480, 27648. Row 3–16296, 51019, 15768).

**Extended Data Fig. 4.**
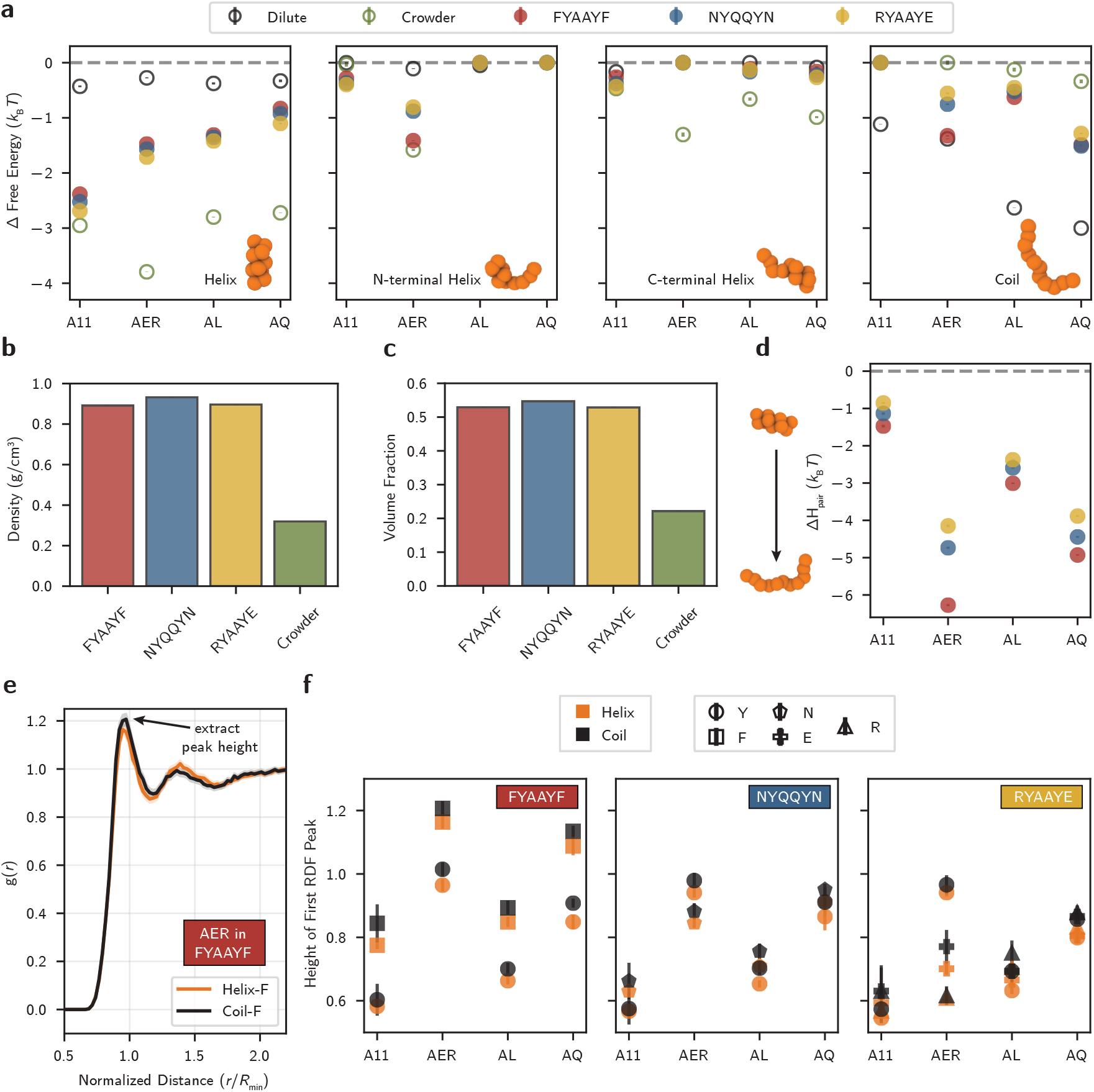
Additional evidence that multivalent interactions and crowding control protein folding landscapes. **a**, Relative free energies of each metastable state from the PCCA+ spectral clustering performed on the MSMs in each condition. Standard errors are taken from 100 different instantiations of our MSMs combined with Bayesian uncertainty within the models. **b**, The density of each condensate and the crowded environment. Standard error is from 5 2.5 µs simulation replicates. **c**, The volume fraction of the co-condensate proteins within each condensate and the crowded environment. Error as in **b. d**, Pairwise contact enthalpy differences for the helical peptide when the peptide is folded versus unfolded. The standard error is taken over all 300 replicates of the 2.5 µs simulations for each pairing. **e**, An example radial distribution function from AER in FYAAYF. Error as in **d. f**, The height of the first peak in the helix (orange) and coil (black) states for every RDF produced from the peptide–condensate pairings. Errors are reported as in **d**.

**Extended Data Fig. 5.**
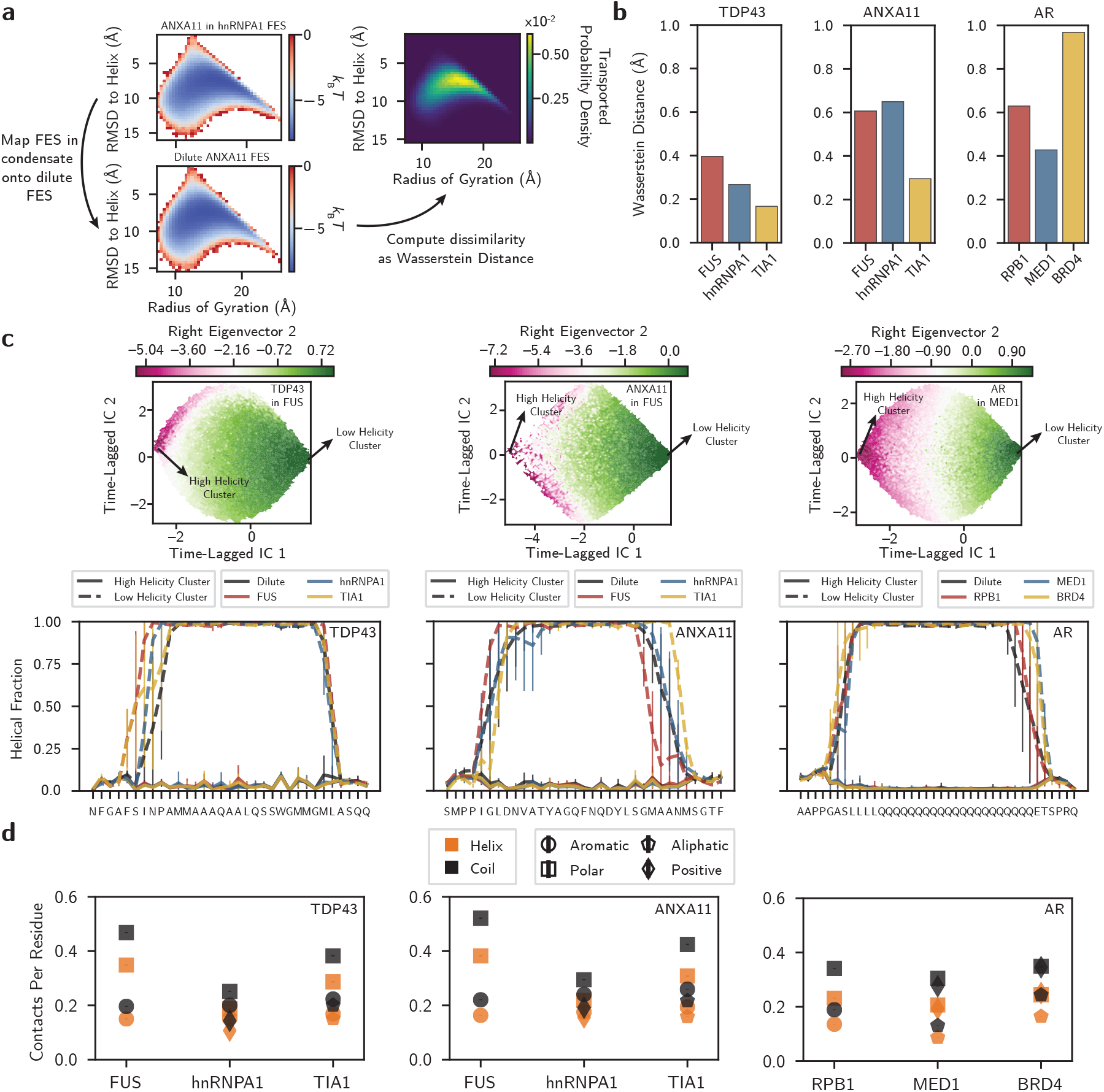
Additional analyses demonstrating that the folding landscape of proteins are dually-sequence dependent in condensates. **a**, The Wasserstein distance as a dissimilarity metric to compare the free energy surface of every α-helix domain in the dilute solution versus the condensates. Calculations are performed with the python package Optimal Transport [88] **b**, The Wasserstein distance for each protein–condensate pairing’s free energy surface compared against the dilute solution free energy surface. **c**, Example MSM data (top row) from TDP43 in FUS, ANXA11 in FUS, and AR in RPB1 depicting that the longest decaying kinetic mode is between the folded and unfolded states for the α-helix domains (bottom row). Standard errors are calculated for per-residue helical fraction using all protein configurations mapped to the highest and lowest cluster over 25 MSM instantiations in each pairing. **d**, Close contact analysis between residues of the *α*-helix domain in a helical (orange) versus a coil state (black) and co-condensate protein residues. The standard error is calculated from all simulation replicates.

