## Supplementary Information for "Biomolecular Condensates Dictate the Folding Landscape of Proteins"

(Dated: January 19, 2026)

#### CONTENTS

I. Atomistic Simulations of the AQ Helical Peptide 2
A. Choice of ff19sb/a99sb-disp Force Field 2
B. Metadynamics Simulations of AQ in Dilute Solution 2
C. NYQQYN Slab Simulations 2
D. Temperature Replica Exchange Simulations (NYQQYN Condensate and PEG Solution) 3

II. Mpipi-Helix Model Development 12
A. Mpipi 12
B. Mpipi-Helix: An $\alpha$-carbon Structural Potential 12
C. Implementing Mpipi-Helix in the LAMMPS Software 14
D. Energy Conservation of Mpipi-Helix 16
E. Potts Model 16
F. Gaussian Process Training 22
G. Helix–Coil Classification in the GP Workflow 28
H. Dataset From Experimental NMR Measurements 38
I. Dataset from REST2 Enhanced Sampling Atomistic Simulations 42

III. Mpipi-Helix Model Validation 56
A. A11 Well-Tempered Metadynamics Simulations 56
B. Validation Against Atomistic Simulations 56
C. Validation Against NMR Measurements and Contemporary Residue-Level Force Fields 58

IV. Helical Peptides in Model Condensates 61
A. Phase Diagrams of Co-Condensate Peptides and Determining Molecular Densities, Co-Condensate Peptide Number, and Crowder Pressure 61
B. PEG Model - Excluded Volume Polymer 65
C. Density and Volume Fraction Calculations 67
D. Markov State Models and PCCA+ Spectral Clustering 67
E. Additional Results of Helical Peptides in Model Condensates 73

V. IDP Helical Domains in Condensates 81
A. Co-Condensate IDP Densities and Pressures 81
B. MSM Validation 84
C. Additional Results for IDP Helical Domains in Condensates 91
D. Thermodynamic and Kinetic Data on the Additional Helices in AR 94

References 96

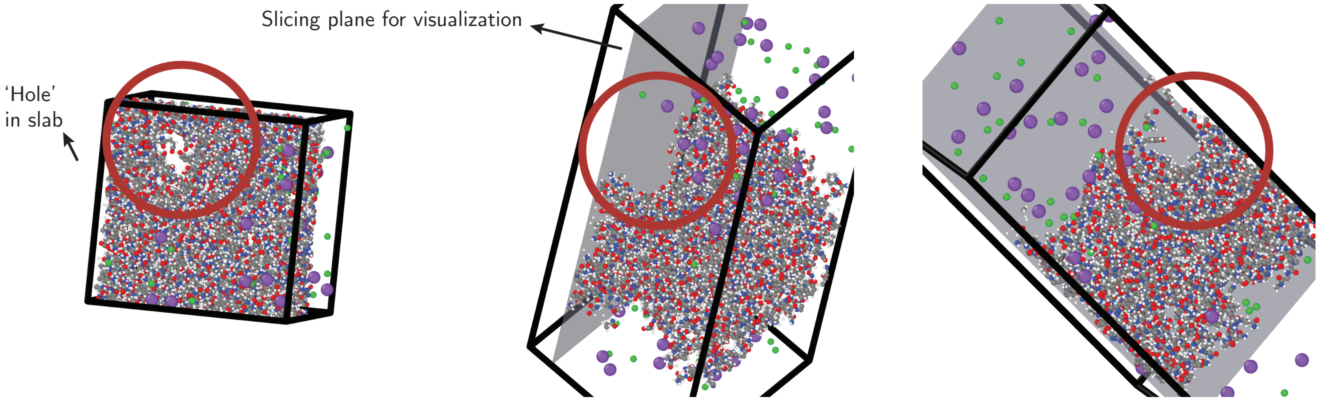


SI Fig. 1. **Direct coexistence simulation of NYQQYN peptides in ff19sb.** An NYQQYN direct coexistence simulation in a slab geometry in the ff19sb [1] force field yields long-lasting holes and gaps, indicating nonphysical behavior for disordered peptides. Water molecules are hidden for visualization purposes.

#### I. ATOMISTIC SIMULATIONS OF THE AQ HELICAL PEPTIDE

##### A. Choice of ff19sb/a99sb-disp Force Field

We initially considered using the ff19sb force field [1] for simulating the AQ peptide, as ff19sb was parameterized to accurately describe $\alpha$-helices. However, when we performed a direct coexistence simulation with NYQQYN in ff19sb we noticed that the proteins appeared to produce a solid aggregate (SI Fig. 1). Specifically, we observed that there were long-lasting (e.g., persisting for the entire duration of the simulation) holes and cavities produced in the slab over the course of the simulation. We speculated that this was because ff19sb was not parameterized or validated on any systems that phase separate and therefore such systems may produce static protein aggregates depending on the protein length and sequence.

Therefore, we considered using a99sb-disp [2] as an alternative force field as it has been used successfully to investigate $\alpha$-helix domains in IDPs [2, 3]. We found that a99sb-disp was able to produce a fluid condensate in direct coexistence simulations (see the 'NYQQYN Direct Coexistence Simulation' section below). However, we also observed that a99sb-disp produced significantly lower helical fractions than ff19sb and did not show substantial differences in helical fraction between an uncapped and capped AQ peptide (SI Fig. 2). We therefore combined the ff19sb and a99sb-disp force fields to obtain the best of each; we used ff19sb's structural parameters for the AQ peptide to provide reasonable helical fractions, but we took every other force field parameter from a99sb-disp (e.g., pairwise potential, structural parameters for NYQQYN, and TIP4P-D water) to ensure that the NYQQYN peptides would not form an aggregate. Finally, we decided to simulate uncapped AQ as we did not plan to parameterize capping groups in Mpipi-Helix (capping refers to an N-terminal acetyl group and a C-terminal N-methyl amide).

The NYQQYN direct coexistence simulation in ff19sb (SI Fig. 1) was performed identically to the simulation of NYQQYN in a99sb-disp as is described in the Methods. Similarly, all dilute solution simulations for SI Fig. 2 were performed using well-tempered metadynamics [4] exactly as is described for the dilute AQ simulation in the Methods.

##### B. Metadynamics Simulations of AQ in Dilute Solution

We demonstrate the convergence of the well-tempered metadynamics sampling of the AQ peptide in dilute solution by analyzing the peptide's free energy surface (FES) at the end of the 10 μs simualtion trajectory. In particular, SI Fig. 3 shows that during the last 800 ns of the simulation trajectory, no significant changes occur in AQ's FES along the biased collective variables (RMSD to a perfect helix and radius of gyration).

##### C. NYQQYN Slab Simulations

We performed direct coexistence simulations of the NYQQYN protein in a99sb-disp utilizing a slab geometry (Methods). The density profiles from the simulation are shown in SI Fig. 4. We used these densities to construct our simulation of AQ in the NYQQYN condensate. As mentioned in the Main Text, the choice of NYQQYN was informed by the need to

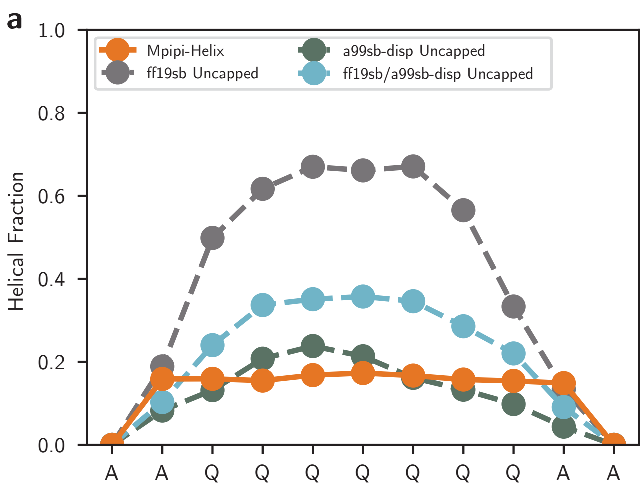


SI Fig. 2. **Helical fractions of the AQ peptide in dilute solution. a,** The helical fractions of the uncapped AQ peptide in Mpipi-Helix and the atomistic force fields a99sb-disp [2], ff19sb [1], and a combined ff19sb/a99sb-disp hybrid. **b,** The helical fractions of the capped AQ peptide in the same force fields as **a**.

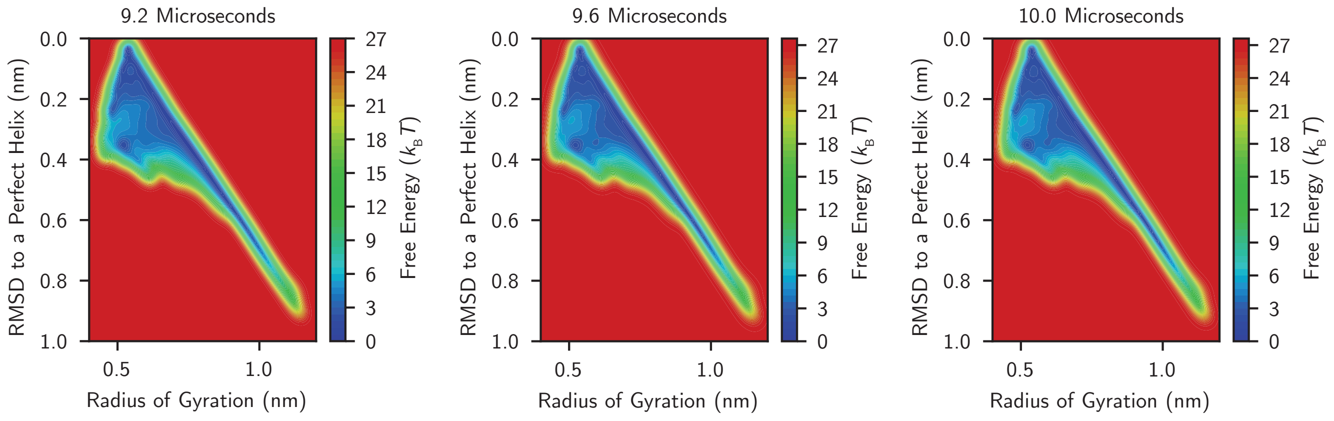


SI Fig. 3. **Convergence of well-tempered metadynamics sampling of AQ peptide in dilute solution.** The FES along the two biased collective variables (RMSD to a perfect helix and radius of gyration) from metadyanmics sampling of the AQ peptide in a dilute solution. Negligible change occurs in the free energy surface between 9.2 µs, 9.6 µs, and 10.0 µs.

have a fluid condensate that was amenable to sampling while still having the requisite multivalent residues (i.e., tyrosine) to enable condensation.

##### D. Temperature Replica Exchange Simulations (NYQQYN Condensate and PEG Solution)

We analyzed the exchange probabilities, helical fractions, and FESs of the AQ peptide in the NYQQYN condensate and PEG solution to determine whether each T-REMD simulation had converged sufficiently to compare their impacts on AQ's folding landscape. We observed that the exchange probabilities were between 20% and 40% for these simulations, indicating efficient sampling of phase space for both the NYQQYN and PEG systems (SI Fig. 5 and 8). However, the helical fractions for the temperature replicas do not strictly adhere to a temperature-ranked ordering where the lowest helical fractions are the replicas with the highest temperatures and vice versa (SI Fig. 7 and 10). This suggests that the T-REMD simulations are not completely converged for the NYQQYN condensate and PEG solution. This is further supported by the fact that the FESs of the cold replicas do not visit all the regions explored by the hottest replicas and the dilute solution (SI Fig. 6 and 9). We anticipate that, given enough time, every region that is explored by the dilute solution should be visited by the coldest replica (298.15 K) in both the PEG solution and the condensate.

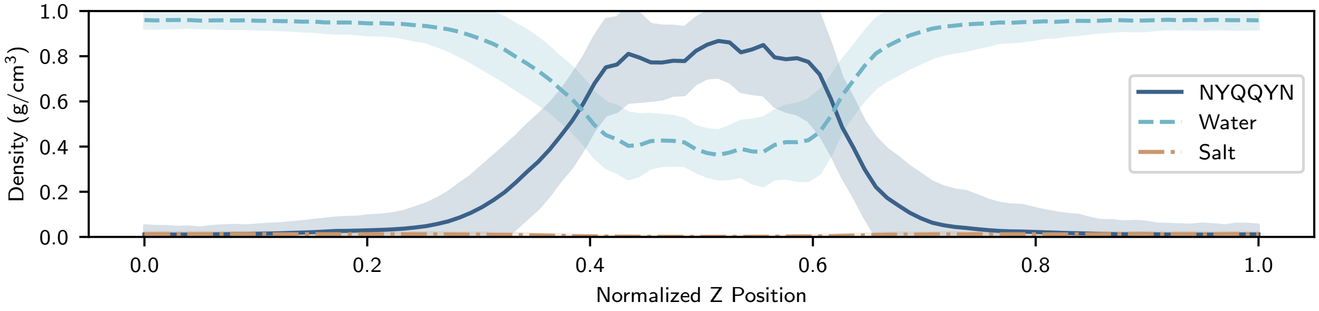


SI Fig. 4. **NYQQYN direct coexistence density profiles in a99sb-disp.** The average density of the NYQQYN protein, TIP4P-D water, and sodium chloride salt from a direct coexistence simulation in a slab geometry in the a99sb-disp force field [2]. The simulation was run for a total of 1.5 µs with 1 µs taken as the sampling period. Error was calculated as the standard deviation of each density across the simulation's sampling period.

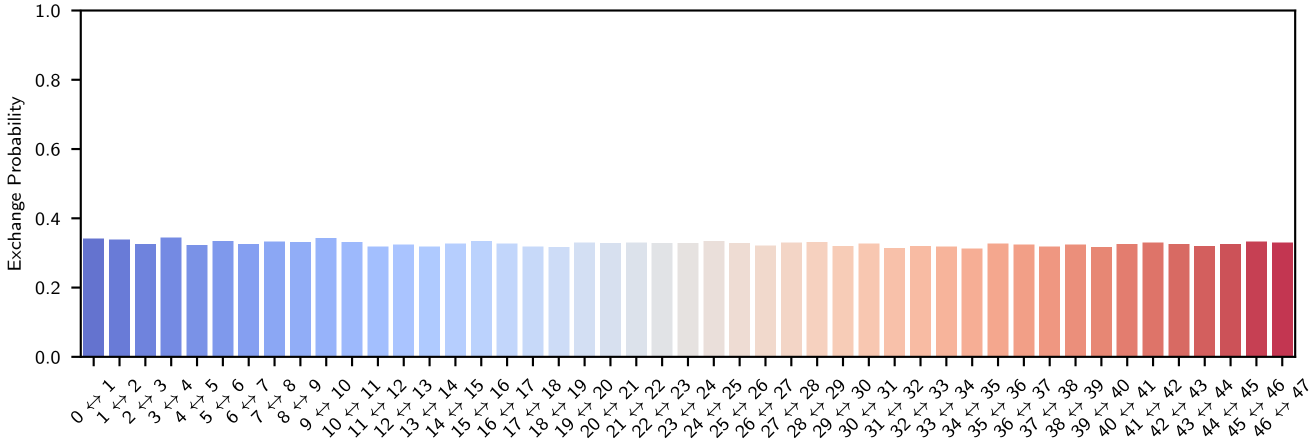


SI Fig. 5. **Replica exchange probabilities for the AQ–NYQQYN T-REMD simulation.** Pairwise exchange probabilities for each temperature replica in the T-REMD simulation of AQ in the NYQQYN condensate using the mixed ff19sb/a99sb-disp force field.

Despite this lack of complete convergence, the data still show qualitative differences between the AQ peptide in the NYQQYN condensate compared to the dilute solution and the solution comprised of PEG crowder. In particular, the difficulty of converging the NYQQYN and PEG T-REMD simulations (despite $>$60 µs of aggregate simulation time) suggests that conformational rearrangements are hindered in the dense environments compared to a dilute solution. Further, the average helical fraction of AQ in the NYQQYN condensate is lower for every temperature replica than the corresponding replica in the PEG solution (SI Fig. 11). This suggests that there are qualitative differences in the thermodynamic behavior of the AQ peptide in each of the environments. Additionally, as NYQQYN and PEG have the same volume fractions in the T-REMD simulations, the data indicate there is some other driving force for the altered conformational landscape of AQ in the NYQQYN condensate besides crowding.

We further recreated Main Text Fig. 1 using the hottest replica (410 K) to emphasize the qualitative difference between the NYQQYN and PEG systems. As demonstrated in SI Fig. 12, at 410 K AQ in NYQQYN is more unfolded than in the PEG solution, providing evidence to support that qualitative differences exist between the two environments.

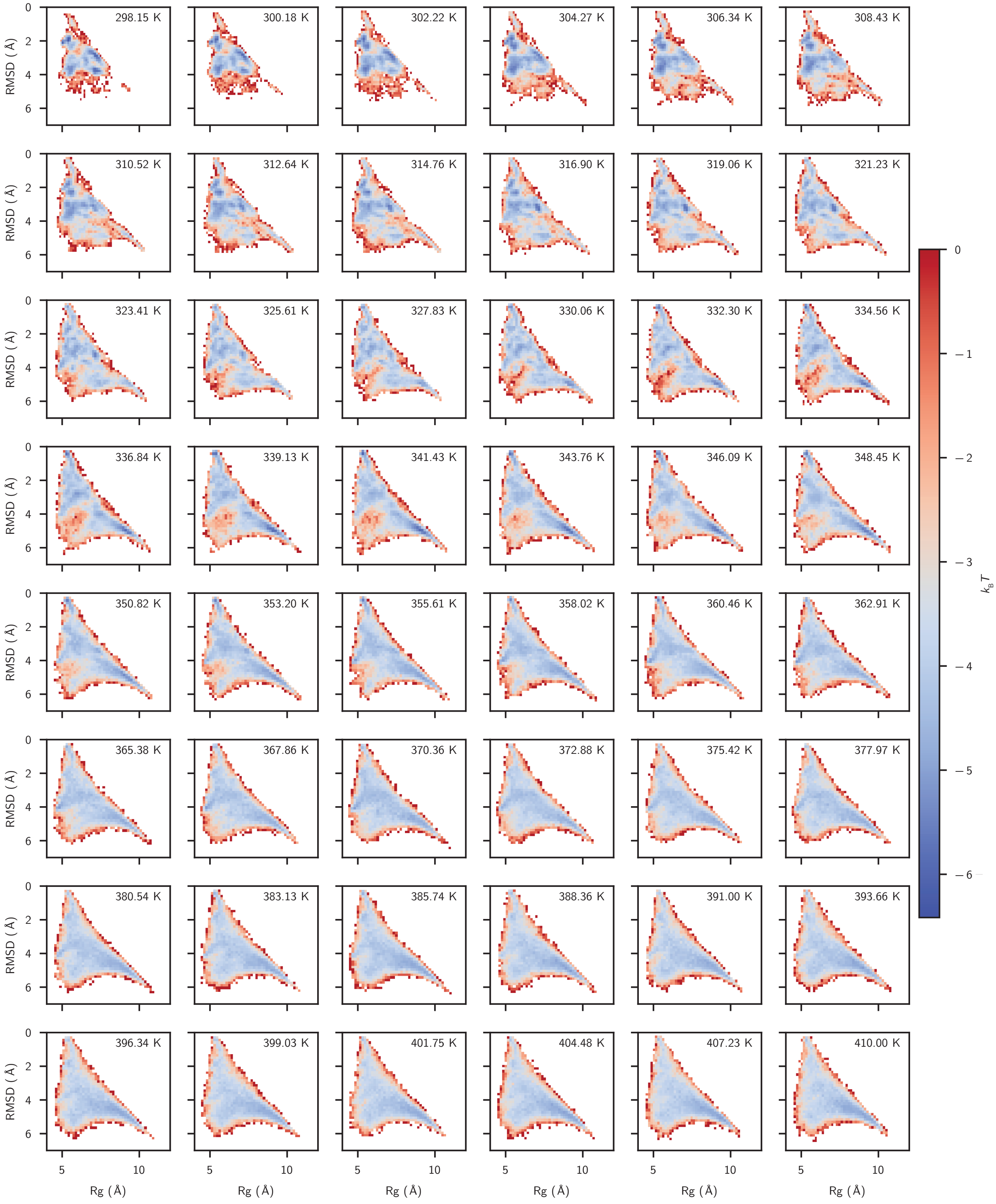


SI Fig. 6. **Free energy surfaces of the AQ peptide in the NYQQYN condensate.** The free energy surface along the RMSD to a perfect helix and radius of gyration order parameters for each temperature replica of the AQ peptide in the NYQQYN condensate computed via T-REMD.

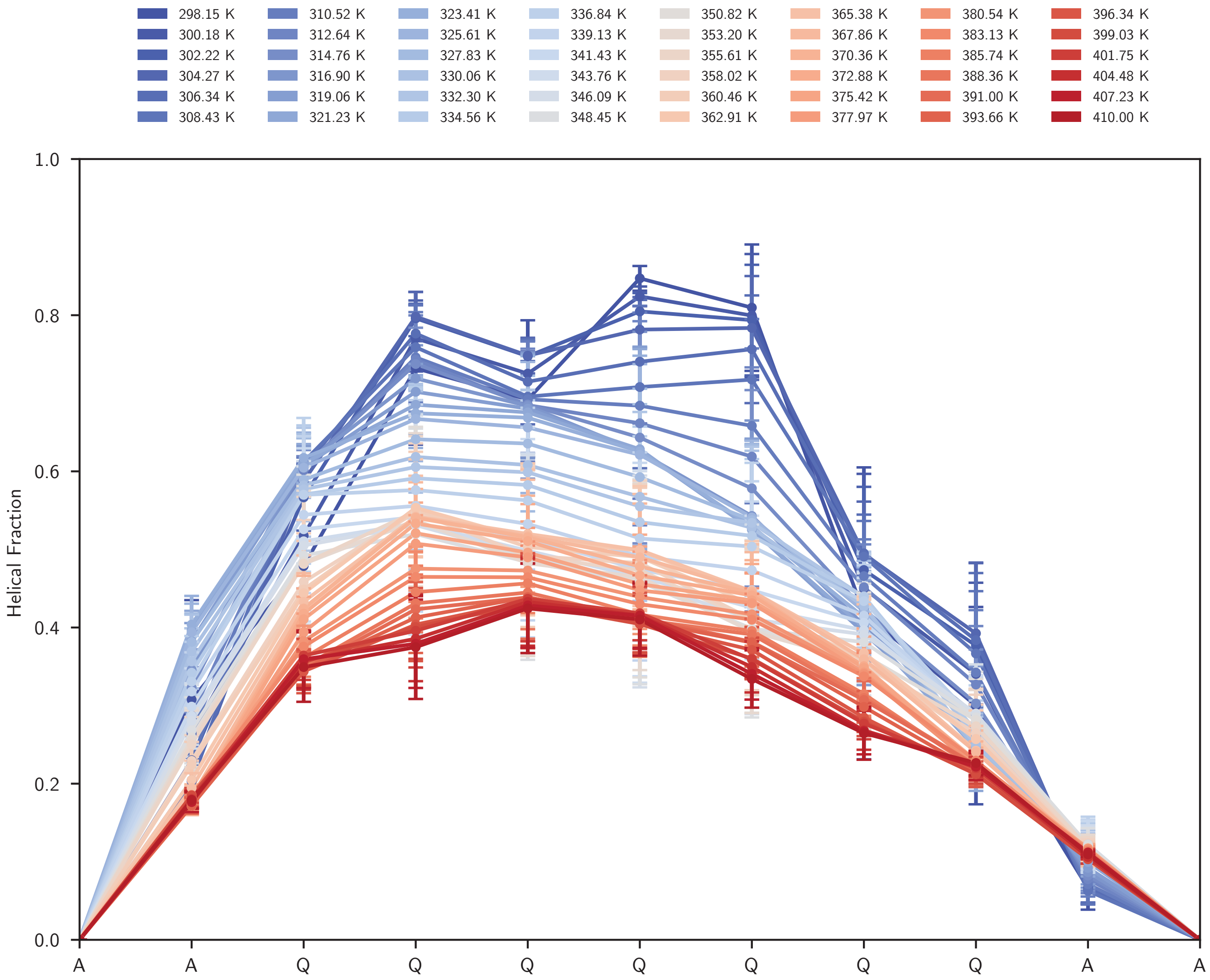


SI Fig. 7. **Helical fractions of AQ peptide in the NYQQYN condensate.** The helical fraction for each temperature replica of the AQ peptide in the NYQQYN condensate computed via T-REMD.

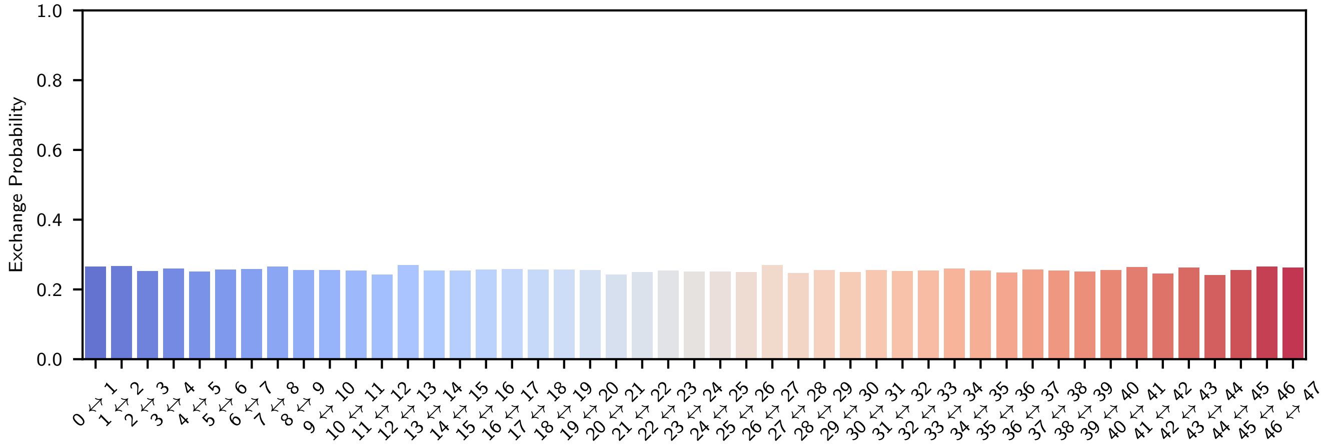


SI Fig. 8. **Replica exchange probabilities for the AQ–PEG T-REMD simulation.** Pairwise exchange probabilities for each temperature replica in the T-REMD simulation of AQ in the PEG solution using the mixed ff19sb/a99sb-disp force field.

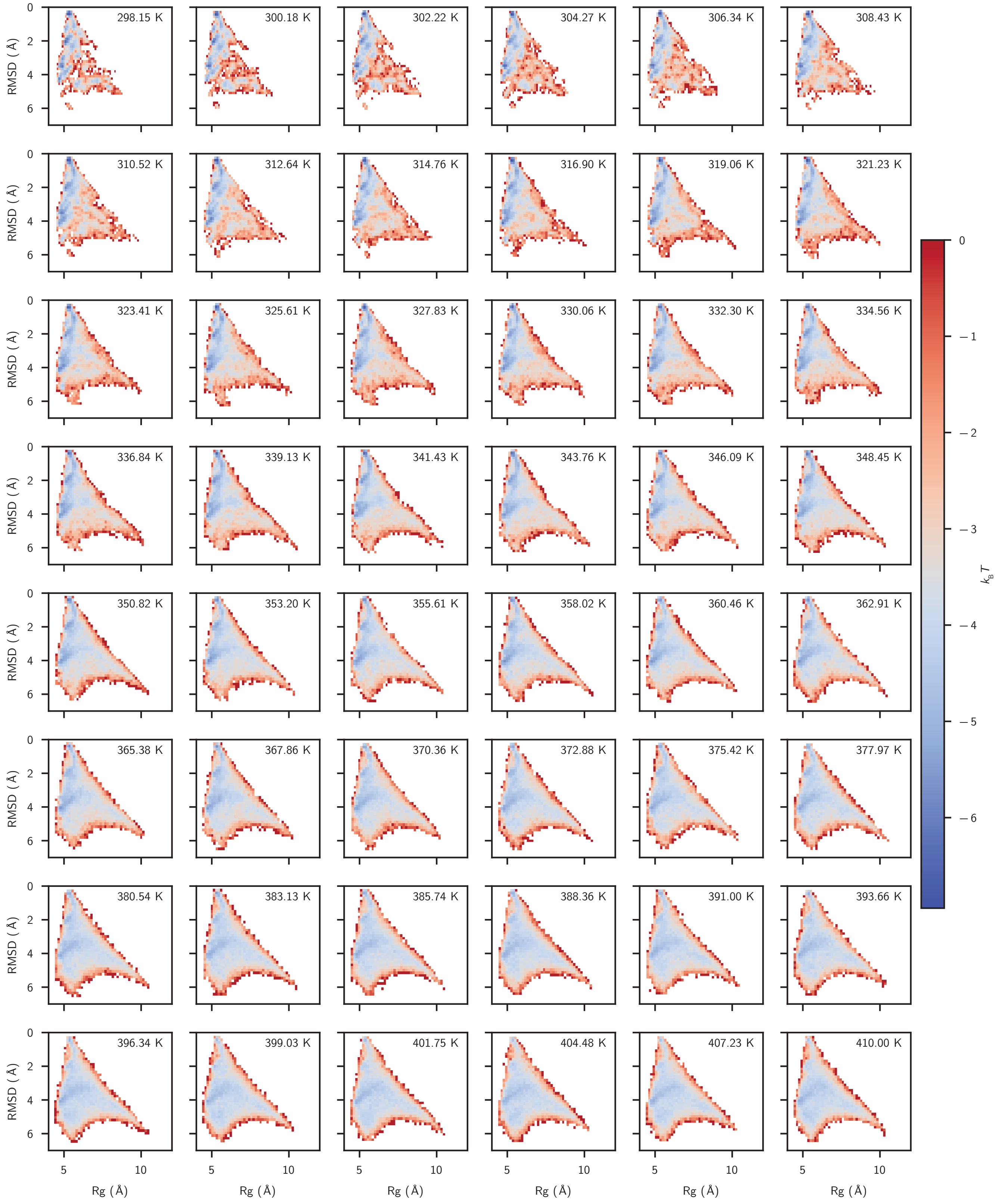


SI Fig. 9. **Free energy surfaces of the AQ peptide in the PEG solution.** The free energy surface along the RMSD to a perfect helix and radius of gyration order parameters for each temperature replica of the AQ peptide in the PEG solution computed via T-REMD.

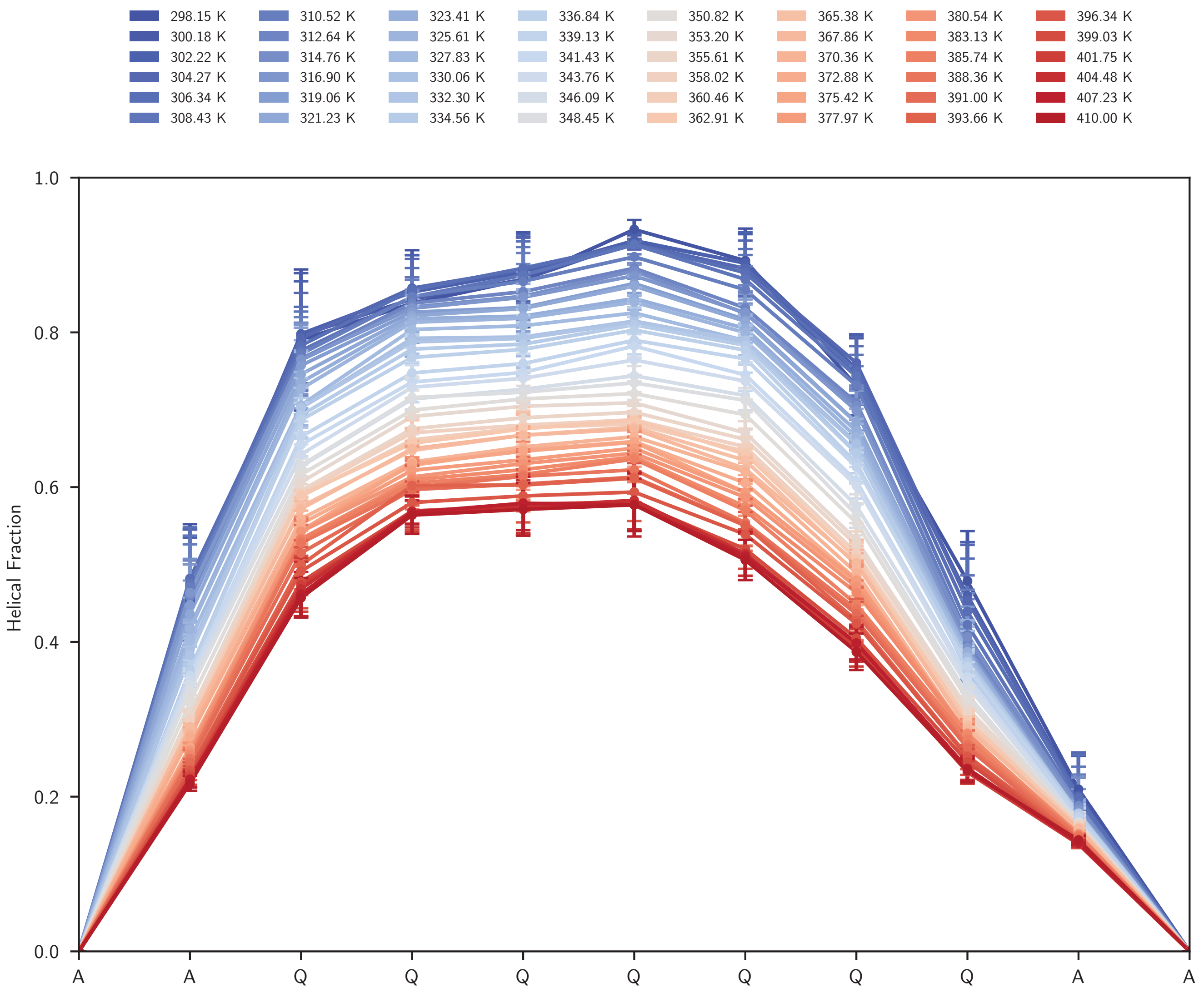


SI Fig. 10. **Helical fractions of AQ peptide in the PEG solution.** The helical fraction for each temperature replica of the AQ peptide in the PEG solution computed via T-REMD.

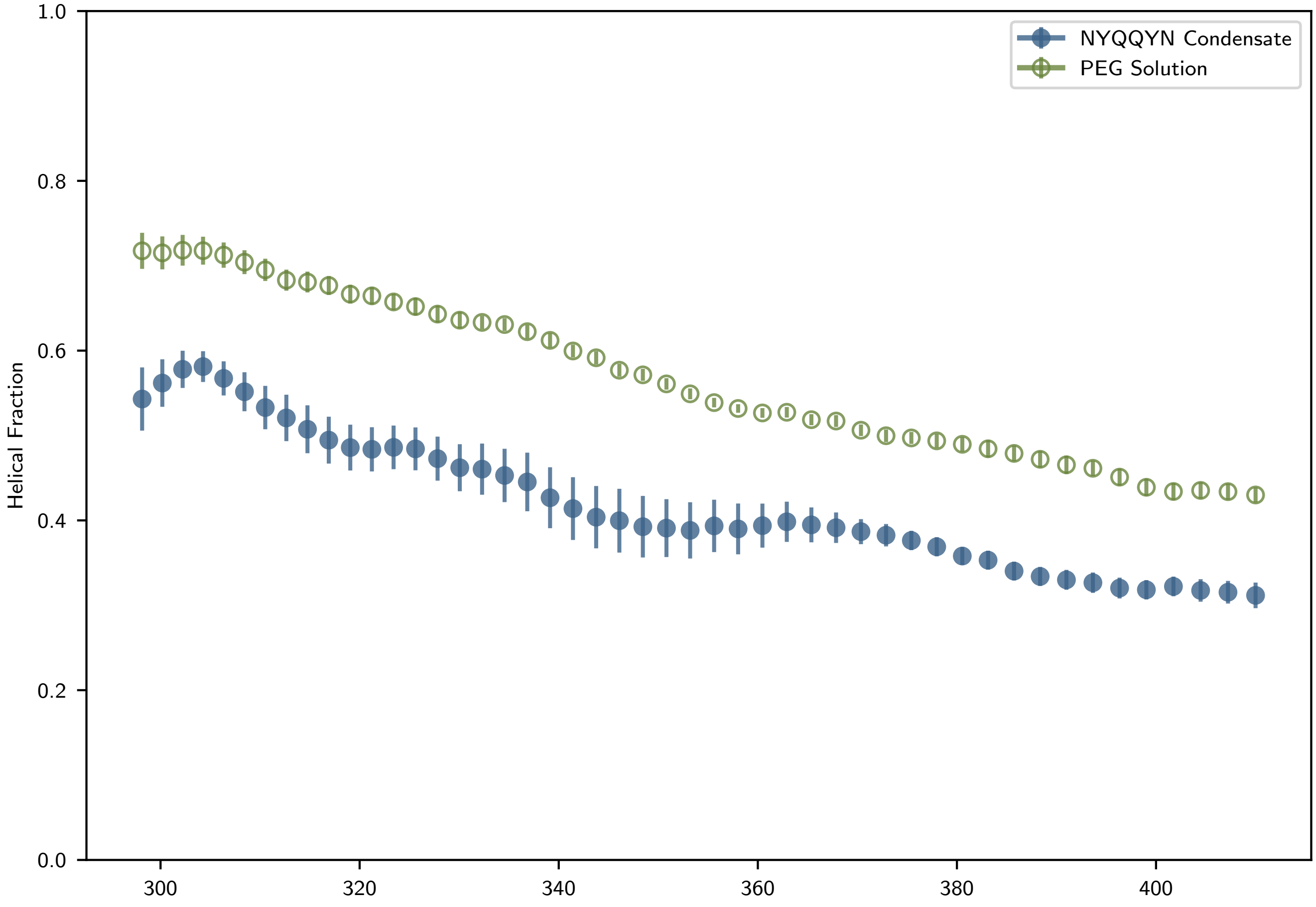


SI Fig. 11. **Average helical fraction of AQ temperature replicas.** The helical fraction for each temperature replica in the NYQQYN condensate (blue symbols) and PEG solution (green symbols). The average helical fraction is calculated from the central nine residues in AQ (i.e., neglecting the uncapped end residues).

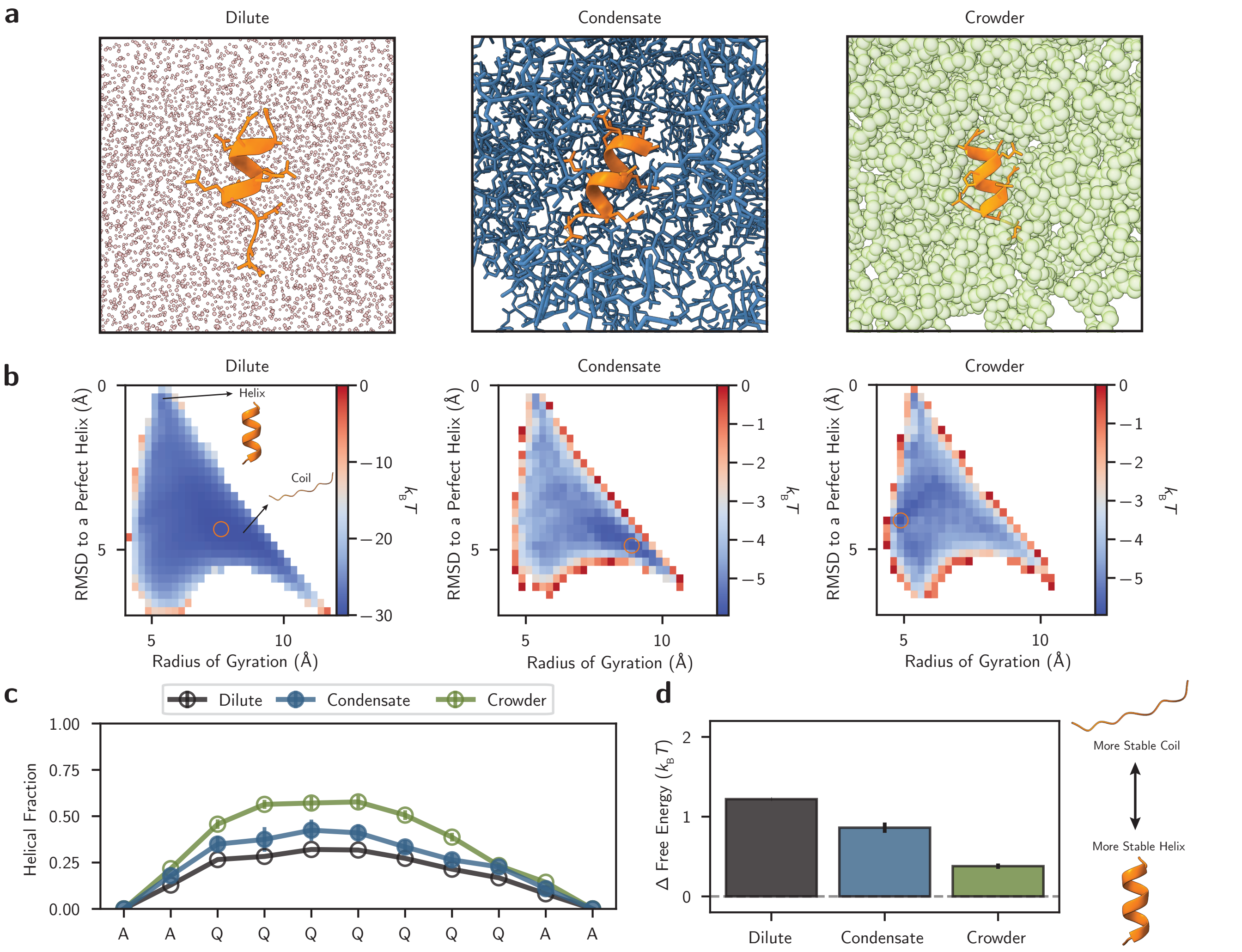


SI Fig. 12. **The hottest replicas from the T-REMD simulations supports that crowding alone does not explain $\alpha$-helix thermodynamics in condensates.** **a**, Atomistic simulations of the AQ helical peptide simulated in dilute (left), NYQQYN condensate (middle), and PEG crowder (right) conditions. **b**, Free energy surfaces of the AQ peptide with radius of gyration and RMSD between the AQ peptide and a folded helix as order parameters. **c**, Per-residue helical fractions of the AQ peptide from backbone dihedral angles analysis. **d**, Estimated free energy differences between the folded helix and unfolded coil states of the AQ peptide. All simulation data taken from the hottest simulation replica (410 K) of the AQ peptide in the NYQQYN condensate and PEG solution. The dilute solution data is taken from a 410 K run of AQ sampled using well-tempered metadynamics for 5.9 µs. Errors in **c**,**d** are calculated via block averaging over three equally-sized blocks.

#### II. MPIPI-HELIX MODEL DEVELOPMENT

This supporting section provides details on the development of the Mpipi-Helix model. All codes associated with this model are provided in the corresponding folder labeled "Mpipi-Helix Model Development" within the GitHub repository associated with this article.

##### A. Mpipi

Mpipi is an implicit solvent residue-resolution force field with each interaction site centered on an amino acid $\alpha$-carbon. Mpipi includes terms for pairwise non-bonded interactions, pairwise electrostatic interactions, and bonded interactions.

The non-bonded interactions are simulated with the Wang–Frenkel potential [5]. The form of the potential is provided in Equations (1) and (2).

$$U_{\mathrm{NB}} = \epsilon\alpha([\frac{\sigma}{r}]^{2\mu} - 1)([\frac{r_{\mathrm{c}}}{r}]^{2\mu} - 1)^{2\nu}. \tag{1}$$

$$\alpha = 2\nu(\frac{r_{\mathrm{c}}}{\sigma})^{2\mu}[\frac{1+2\nu}{2\nu[(\frac{r_{\mathrm{c}}}{\sigma})^{2\mu} - 1]}]^{2\nu+1}. \tag{2}$$

In Equations (1) and (2), $\epsilon$ sets the depth of the attractive well, $\sigma$ dictates where the potential begins to become sharply repulsive, $\mu$ sets how steeply the potential becomes repulsive at $\sigma$, $\nu$ sets the well width, and $r_{\mathrm{c}}$ sets where the potential decays to zero energy. $U_{\mathrm{NB}}$ is the energy of the interaction between the non-bonded amino acids at a distance $r$ apart.

The electrostatic interactions are modeled as a Coulomb potential with Debye–Hückel screening as in Equation (3).

$$U_{\mathrm{C}} = \frac{q_i q_j}{\epsilon r}\exp(-\kappa r) \quad r < r_{\mathrm{c}}. \tag{3}$$

Here $q_i$ and $q_j$ are the charges on particles i and j, $\epsilon$ is the dielectric constant of the solvent, $r$ is the distance between i and j, and $r_{\mathrm{c}}$ is the cutoff distance. The inverse Debye screening length $\kappa$ represents the screening of a polar solvent (here water in 150 mM of sodium chloride; $\kappa^{-1} = 0.795\,\mathrm{nm}$). $U_{\mathrm{C}}$ is the energy of the charged interaction.

The bonds between adjacent residues in Mpipi are modeled with a harmonic bond potential. The form of this potential is provided in Equation (4).

$$U_{\mathrm{B}} = K(r - r_0)^2. \tag{4}$$

Here, $K = 8.03\,\mathrm{J\,mol^{-1}\,pm^{-1}}$ is the bond coefficient dictating the bonds's stiffness, $r_0 = 0.381\,\mathrm{nm}$ is the equilibrium bond distance, $r$ is the distance between the two bonded residues, and $U_{\mathrm{B}}$ is the bond energy.

The overall Hamiltonian for a simulation with Mpipi is

$$U_{\mathrm{Mpipi}} = U_{\mathrm{NB}} + U_{\mathrm{C}} + U_{\mathrm{B}}. \tag{5}$$

Where $U_{\mathrm{NB}}$ and $U_{\mathrm{C}}$ are summed over all non-bonded amino acid interactions and $U_{\mathrm{B}}$ is summed over all bonded residues. All simulations involving Mpipi (and Mpipi-Helix) were performed in the LAMMPS software [6]. For further details on Mpipi, see Ref. [7].

##### B. Mpipi-Helix: An $\alpha$-carbon Structural Potential

We built the Mpipi-Helix $\alpha$-helix potential as an additional structural potential on top of the Mpipi force field. The Hamiltonian for Mpipi-Helix is described in Equation 6.

$$U_{\text{Mpipi-Helix}} = U_{\mathrm{NB}} + U_{\mathrm{C}} + U_{\mathrm{B}} + U_{\mathrm{Helix}}. \tag{6}$$

The $U_{\mathrm{Helix}}$ potential has two features in its energy landscape, an energetic well for helix formation (hereafter referred to as the helical well) and an energetic barrier between the coil and helix state (hereafter referred to as the helical barrier). This $U_{\mathrm{Helix}}$ potential is described in Equation 7.

$$U_{\text{Helix}} = -E_{\text{well}} \exp\left(-\frac{(\theta_1-\theta_0)^2}{\sigma^2_{\theta,\text{w}}} - \frac{(\theta_2-\theta_0)^2}{\sigma^2_{\theta,\text{w}}} - \frac{1+\cos(\alpha-\alpha_0+\pi)}{\sigma^2_{\alpha,\text{w}}}\right) + E_{\text{barrier}} \exp\left(-\frac{(\theta_1-\theta_0)^2}{\sigma^2_{\theta,\text{b}}} - \frac{(\theta_2-\theta_0)^2}{\sigma^2_{\theta,\text{b}}} - \frac{1+\cos(\alpha-\alpha_0+\pi)}{\sigma^2_{\alpha,\text{b}}}\right). \quad (7)$$

Equation 7 operates on three degrees of freedom about four consecutive bonded amino acids, namely the two bond angles $\theta_1$ and $\theta_2$ as well as the dihedral angle $\alpha$ (see Extended Data Fig. 1a). The potential is centered by the variables $\theta_0$ and $\alpha_0$, which set the bias locations in the bond angle and dihedral angle dimensions. $E_{\text{well}}$ sets the depth of the negative Gaussian (the helical well), with the widths in the bond angle and dihedral angle dimensions controlled by $\sigma_{\theta,\text{w}}$ and $\sigma_{\alpha,\text{w}}$. Similarly, $E_{\text{barrier}}$ sets height for the positive Gaussian (the helical barrier), with the widths in the bond angle and dihedral angle dimensions controlled by $\sigma_{\theta,\text{b}}$ and $\sigma_{\alpha,\text{b}}$. See Extended Data Fig. 1b for a depiction of this potential and its features.

The parameters $\theta_0$ and $\alpha_0$ were set based on a geometrically perfect α-helix with $\theta_0 = 92.4°$ and $\alpha_0 = 51.7°$. The negative Gaussian widths ($\sigma_{\theta,\text{w}}$ and $\sigma_{\alpha,\text{w}}$) were set for overlap with α-helix geometries extracted from the PDB. Specifically, we used the PISCES PDB culling server [8] to query for proteins with less than 25% sequence similarity, including entries using X-ray crystallography, cryo-electron microscopy, and NMR (exact query: Resolution 0.0-2.0, R-factor 0.25, sequence length 40-10,000, sequence percentage identity <= 25.0, chains with breaks excluded, chains with disorder included). The results provided 7,997 protein structures; a list of the PDB entries gathered from this query can be found in the GitHub repository. We used the DSSP algorithm [9] to assign secondary structure classifications to each amino acid in the PDB structures and selected α-helical segments within the dataset. With an in-house script, we extracted the geometry of α-helices at an α-carbon resolution. Plotting the bond angles and dihedral angles of the α-helices informed our determination of $\sigma_{\theta,\text{w}} = 0.25$ and $\sigma_{\alpha,\text{w}} = 0.4$, as there is a significant overlap of the PDB α-helix geometries within a 10% cutoff of the energetic minimum of the helical well depth. See the Extended Data Fig. 1c, where the red line indicating the 'peak of helical barrier' also corresponds to the 10% cutoff of the energetic minimum of the helical well depth.

Four more variables are required to be set for Equation 7: $E_{\text{well}}$, $E_{\text{barrier}}$, $\sigma_{\theta,\text{b}}$, and $\sigma_{\alpha,\text{b}}$. To set these values uniquely, we make three choices. The first choice is to locate the peak of the helical barrier at the 10% energy cutoff of the negative well Gaussian (see Extended Data Fig. 1c for a depiction). The second and third choices are specifying our desired helical well depth and desired barrier height. With these stipulations, it is possible to write down four equations that can be uniquely solved for $E_{\text{well}}$, $E_{\text{barrier}}$, $\sigma_{\theta,\text{b}}$, and $\sigma_{\alpha,\text{b}}$. These equations are listed as Equations 8, 9, 10, and 11.

$$0 = \frac{2(\theta_1-\theta_0)}{\sigma^2_{\theta,w}}\left(W + \frac{B + W\exp(w)}{\exp(b)-\exp(w)}\right)\exp(w) - \frac{2(\theta_1-\theta_0)}{\sigma^2_{\theta,b}}\left(\frac{B + W\exp(w)}{\exp(b)-\exp(w)}\right)\exp(b). \quad (8)$$

$$0 = -\frac{\sin(\alpha_1-\alpha_0+\pi)}{\sigma^2_{\alpha,w}}\left(W + \frac{B + W\exp(w)}{\exp(b)-\exp(w)}\right)\exp(w) + \frac{\sin(\alpha_1-\alpha_0+\pi)}{\sigma^2_{\alpha,b}}\left(\frac{B + W\exp(w)}{\exp(b)-\exp(w)}\right)\exp(b). \quad (9)$$

$$E_{\text{barrier}} = \frac{B + W\exp(w)}{\exp(b)-\exp(w)}. \quad (10)$$

$$E_{\text{well}} = W + E_{\text{barrier}}. \quad (11)$$

Where

$$\exp(w) = \exp\left(-\frac{(\theta_1-\theta_0)^2}{\sigma^2_{\theta,w}} - \frac{(1+\cos(\alpha_1-\alpha_0+\pi))}{\sigma^2_{\alpha,w}}\right). \quad (12)$$

$$\exp(b) = \exp\left(-\frac{(\theta_1-\theta_0)^2}{\sigma^2_{\theta,b}} - \frac{(1+\cos(\alpha_1-\alpha_0+\pi))}{\sigma^2_{\alpha,b}}\right). \quad (13)$$

In these equations, $W$ is the desired helical well depth, $B$ is the desired helical barrier height, and $(\theta_1, \alpha_1)$ is any point located at the maximum of the helical barrier which is set uniquely by requiring that the helical barrier is at its maximum at a 10% energy cutoff of the negative Gaussian alone.

With these equations, any desired combination of a helical well depth ($W$) and helical barrier height ($B$) maps to a unique helical potential in Equation 7.

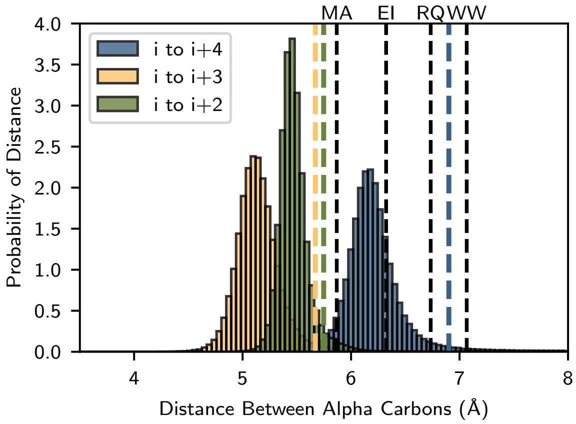


SI Fig. 13. **Intra-helix α-carbon distance distributions.** Histograms of α-carbon distances between residues $i$ to $i+2$, $i$ to $i+3$, and $i$ to $i+4$ within α-helical structures extracted from the PDB. The dashed colored lines indicate the mean plus two standard deviations for each of the distributions. Alongside the distributions, we plot the $\sigma$ Wang–Frenkel potential values for four pairs of amino acids (MA, EI, RQ, and WW).

##### C. Implementing Mpipi-Helix in the LAMMPS Software

Implementing Equation 7 presents a challenge as α-helices are anisotropic structures whereas the Mpipi-Helix force field represents amino acids as isotropic spheres. This is problematic as the intra-helix residue spacing can be significantly smaller than the $\sigma$ of the Wang–Frenkel pair potential used in Mpipi-Helix. To demonstrate this, we extracted the distance distributions between residues $i$ to $i+2$, $i$ to $i+3$, and $i$ to $i+4$ within the α-helices from the PDB dataset taken from the PISCES culling server [8]. These distributions are plotted against several $\sigma$ values from the Mpipi-Helix Wang–Frenkel pairwise potential in SI Fig. 13. Critically, the $\sigma$ values of large residues in Mpipi-Helix are larger than the means of the $i$ to $i+2$, $i$ to $i+3$, and $i$ to $i+4$ distributions which would create artificial steric hindrances for α-helices in the model.

As a demonstration of this problem, we applied our helical potential (Equation 7) to 10,000 random peptide sequences of 11 amino acids in length to determine what percentage of simulation frames the peptides were within the helical well. Each peptide was simulated separately in the $NVT$ ensemble for 50 ns using a Langevin thermostat with a 100 ps damping constant at 300 K and a 10 fs timestep. Frames were recorded every 0.01 ns. For these simulations, we applied a $W = 5.0\,\mathrm{kcal/mol}$ and $B = 1.5\,\mathrm{kcal/mol}$ potential to ensure the simulations had a strong thermodynamic driving force for the helical state. As we were interested in classifying whether our peptides could enter the helical well, we considered a residue to be helical if it was one of the two middle residues in a helical potential currently within the peak of the helical barrier. criteria that considered a residue to be 'helical' if it was one of the central two residues in a helical potential that had a configuration within the peak of the helical barrier. The peptide helical fraction reported in SI Fig 14a shows the average of these helical classifications along the 11 length protein chains. The sequences' helical fractions correlate well with the average residue size of the sequences, where residue size was taken as the average $\sigma$ value from the non-bonded potential for each residue's self-interaction in Mpipi-Helix. This is undesirable as there are sequences which are entirely prevented from forming a helix whatsoever.

Traditionally, this issue is resolved in coarse-grained force fields by eliminating pairwise interactions between residues that are involved in the same structural potential, such as bond angles and dihedral angles (e.g., as is done in Ref. [10, 11]). Then, by having disallowed regions (i.e., high energies) on the energy landscape of these potentials, one can prevent non-physical overlaps of coarse-grained residues. However, we choose to avoid this strategy as it would require parameterizing the features of the energy landscape in the coil state.

Instead, we seek to set the energy of the coil state to be zero without loss of generality. Thus, by parameterizing based on atomistic simulation and experimental NMR data that effectively captures the free energy difference between the helix and coil states, we capture the effect of Mpipi-Helix pairwise potential on the coil state. In other words, as long as we parameterize correctly on accurate data, all aspects of the coil state (in relation to the energetics of the helix state) should be well described by the Mpipi-Helix's full Hamiltonian (including the pairwise potential). In doing so, we sacrifice the ability to describe specific structures in the coil state (e.g., hairpins and β-sheets), but we are happy to make this trade-off given the simplification it provides to the parameterization process. This is in addition to the fact that we are principally concerned with describing the helix–coil transition and we focus on peptides without significant β-sheet content in our

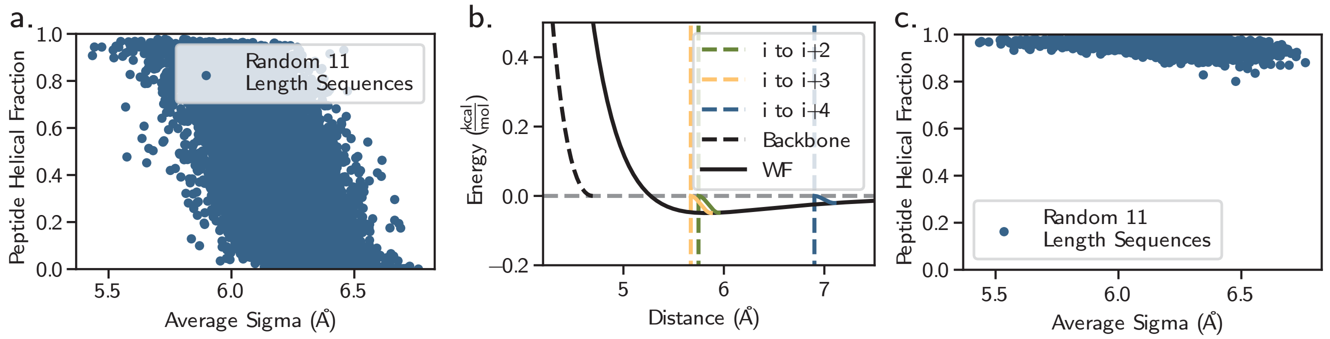


SI Fig. 14. **The ramping scheme enables peptide sequences of all sizes to access the helical state.** **a,** 10,0000 separate simulations of random 11 length peptides were performed for 50 ns in the *NVT* ensemble utilizing a Langevin thermostat with a 100 ps damping constant at 300 K. The potential in Equation 7 was applied with $W = 5.0$ kcal/mol and $B = 1.5$ kcal/mol. **b,** Ramps for the alanine-alanine Wang–Frenkel potential (WF) to the backbone potential (Backbone) are displayed alongside the inter-residue distance for two standard deviations above the mean for the $i$ to $i+2$, $i$ to $i+3$, and $i$ to $i+4$ distributions within $\alpha$-helices (dashed lines). See SI Fig. 13 for where these lines compare to the distributions of $i$ to $i+2$, $i$ to $i+3$, and $i$ to $i+4$ distances in helices from the PDB. The ramps for each interaction are displayed in the solid colored lines. **c,** The same simulations are performed as in **a,** however the ramping scheme is utilized.

study. Thus, the solution we have implemented for Mpipi-Helix is to dynamically shift the intra-helix Wang–Frenkel pair potential over the course of the folding process to accommodate preferred intra-helix residue spacing. This provides us with a potential that is compatible with $i$ to $i+2$, $i$ to $i+3$, and $i$ to $i+4$ interactions, enabling us to apply our helical potential in a modular fashion to regions of the protein chain that we know contain helical domains while leaving the rest totally disordered.

We accomplished this task through what we term 'ramping' the non-bonded interactions. The ramps are continuous and differentiable cubic spline potential functions that go between a non-bonded Wang–Frenkel interaction and a 'backbone' potential which ensures two residues cannot overlap, as shown in SI Fig. 14b. These ramps are placed such that their lower ends are at the distance where the two residue pairings are two standard deviations higher than the average $\alpha$-carbon distance in $\alpha$-helices (see SI Fig. 14b and SI Fig. 13) for their appropriate pairing (e.g., $i$ to $i+2$, $i$ to $i+3$, or $i$ to $i+4$). These ramps have their upper ends on the Wang–Frenkel curve at their lower end plus 0.2 Å. For amino acids where $\sigma$ occurs at a further distance than the two standard deviations plus 0.2 Å criteria, the amino acids are ramped from their $\sigma$ to the backbone potential at $\sigma - 0.2$ Å. Critically, the ramps only turn on when the configuration of four residues in the same helical potential begins encountering significant force from the $\alpha$-helix potential (set to be at a 10% energetic threshold of the helical barrier). This ensures that the ramp is only accessed when the residues are approaching the helical state and are feeling forces from the helical potential. Otherwise, the ramps remain off and the residues experience the Wang–Frenkel potential as normal. Additionally, once a residue is inside the start distance of a ramp but is on the Wang–Frenkel potential, even if it begins to be within the 10% threshold of the helical barrier, it will remain on the Wang–Frenkel potential until it passes to a distance further then the ramp which then enables the ramp to turn on. This means that particles will not 'fall' from the Wang–Frenkel potential to the backbone potential in a non-continuous manner. A Jupyter notebook is provided in the GitHub repository which provides separate plots of every ramp location (akin to SI Fig. 14).

Practically, ramping is performed 'under the hood' in the simulation by communication between the dihedral potential, the Wang–Frankel potential, and an atom style class, each of which was coded specifically for the Mpipi-Helix force field. The atom style class `atom_vec_psproteins.cpp` keeps track of what state a residue is in, both in reference to whether it is within a helical potential that is structured (i.e., within the barrier cutoff) and whether it is currently experiencing 'ramping' with any of its $i+2$ to $i+4$ neighbors. This information is utilized in the `pair_wf_cut_psproteins.cpp` class in order to know whether a pair of residues is experiencing forces from the Wang–Frenkel potential, a ramp, or the backbone potential. The current status of whether a residue is within the barrier cutoff or not is updated in the `dihedral_combinedgaussianpsproteins.cpp` class, which is also where the $\alpha$-helix potential is applied. Finally, there is an additional reverse communication performed via the `pair_wf_cut_psproteins.cpp` within `verlet.cpp` in order to ensure that information is not lost on ghost atoms at the end of a timestep in LAMMPS.

The simulations from SI Fig. 14a were repeated with the same 10,000 sequences using the ramping scheme and the exact same simulation procedure. With the ramping scheme, it is clear that the helical fractions of the sequences does not correlate strongly with the sequences' average residue size as seen in SI Fig. 14d. This is desirable and indicates that, using this ramping scheme, we can make any arbitrary protein sequence helical with Equation 7.

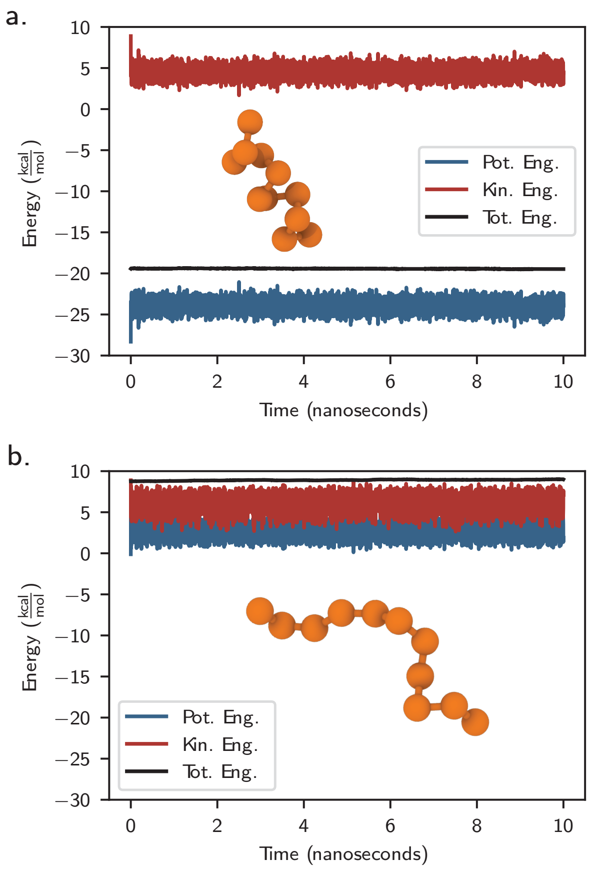


SI Fig. 15. **The Mpipi-Helix $\alpha$-helix potential demonstrates energy conservation within the NVE ensemble. a,** A 10 ns *NVE* ensemble simulation with an 11 length all alanine peptide starting in a helical configuration. **b,** A 10 ns *NVE* ensemble simulation with an 11 length all alanine peptide starting in a coil configuration. Depicted are the potential (Pot Eng.), kinetic (Kin. Eng.), and total energies (Tot. Eng.) within the simulations. Both trajectories demonstrate energy conservation as the drift in total energy is approximately two orders of magnitude smaller than the fluctuations in potential energy.

##### D. Energy Conservation of Mpipi-Helix

As a final check on the implementation of $U_{\text{Mpipi-Helix}}$, including the ramping scheme described above, we tested the energy conservation of the $\alpha$-helix potential in the *NVE* ensemble. Specifically, we simulated an 11 length all alanine peptide for 10 ns in two different starting configurations: a helix (all $\theta_1 = \theta_2 = 92.4°$ and $\alpha = 51.7°$) and a coil (a completely straight peptide chain). Initial velocities were provided according to the velocity command in LAMMPS with a temperature of 298.15 K. The potential was applied with $W = 3.45$ kcal/mol and $B = 4.0$ kcal/mol. Energies were recorded in the simulation every 0.001 ns. The timestep was 10 fs. The results are presented in SI Fig. 15a for the helical configuration and SI Fig. 15b for the coil configuration. The drift in total energy is approximately two orders of magnitude less than the drift in potential energy over the course of the simulation, so we consider our implementation of the potential in LAMMPS to be correct and stable.

##### E. Potts Model

We used a Potts model to produce a mapping from any sequence of four amino acid residues to a thermodynamically accurate well depth that can be simulated in Mpipi-Helix. A trained Potts model describes a learned Boltzmann probability distribution [12, 13]. In this study, the learned Boltzmann distribution adheres to a multiple sequence alignment (MSA) of length four sequences; after successfully training a Potts model, any input sequence of four is mapped to the statistical energy representing the likelihood that the sequence of four is found in the distribution of sequences from the MSA.

Equations 14 and 15 describe a Potts model mathematically.

$$P(a_1, ..., a_L) = \frac{1}{\mathcal{Z}} \exp\{-\beta \mathcal{H}(a_1, ..., a_L)\}. \tag{14}$$

$$\mathcal{H}(a_1, ..., a_L) = -\sum_{1 \le i < j \le L} J_{ij}(a_i, a_j) - \sum_{i=1}^{L} h_i(a_i). \tag{15}$$

In these equations $a$ represents any amino acid, $i$ and $j$ are any position in the protein sequence, and $L$ is the total length of the protein sequence. $\mathcal{Z}$ is the partition function measured over all possible states in the Boltzmann distribution. $J$ is an energetic coupling parameter that assign a statistical energy to any amino acid $a_i$ (appearing in position $i$) and any amino acid $a_j$ (appearing in position $j$) that is in principle unique for any given $a_i$ and $a_j$. $h$ is a field parameter which is the statistical energy for any amino acid $a_k$ (appearing in position $k$). If the parameters $J_{ij}$ and $h_i$ are learned accurately, this constitutes a trained Potts model where sampling from the underlying Boltzmann distribution should reproduce the single (field) and coupled empirical frequencies of amino acids as seen in the MSA. Effectively, this would mean Equation 16 holds.

$$\begin{aligned} f_i(a) &= \langle \delta_{a_i,a} \rangle_P, \\ f_{i,j}(a,b) &= \langle \delta_{a_i,a} \delta_{a_j,b} \rangle_P. \end{aligned} \tag{16}$$

Where $f_i(a)$ is the empirical frequency of amino acid $a$ being in position $i$ and $f_{ij}(a,b)$ is the empirical frequency of amino acid $a$ being in position $i$ while amino acid $b$ is in position $j$. $\langle \delta_{a,a_i} \rangle_P$ and $\langle \delta_{a_i,a} \delta_{a_j,b} \rangle_P$ can be produced from sampling the Potts model (e.g., a Markov Chain Monte Carlo walk). Here, $\langle \delta_{a,a_i} \rangle_P$ is the ensemble average over the Potts model's probability distribution of the observed frequency of amino acid $a$ appearing at position $i$. Similarly, $\langle \delta_{a_i,a} \delta_{a_j,b} \rangle_P$ is the ensemble average over the Potts model's probability distribution of the observed frequency of amino acid $a$ appearing in position $i$ and amino acid $b$ appearing in position $j$.

To generate the empirical frequencies for Potts model training, we built an MSA based on predicted structures from AlphaFold 2.0 [14]. Specifically, we took all predicted AlphaFold 2.0 structures from the SwissProt database [15] and clustered them using mmSeqs2 [16] (`createdb`, `cluster -s 7.5`, `createsubdb`, and `createtsv` commands). From the mmSeqs2 clustering, we took representative sequences (in total 56,474 structures, see GitHub repository). Using the DSSP algorithm [9], we extracted all predicted segments of four consecutive $\alpha$-helix ('H') residues to build the MSA. In total, this produced 5,813,468 sequences of length four. This was used for generating the empirical frequencies for Potts model training.

After the empirical frequencies were collected, amino acid frequencies were adjusted according to the relative background frequency of the amino acids within proteins. To accomplish this, we built an MSA of all length four segments from the clustered representative sequences in the SwissProt database. We extracted single and coupled amino acid empirical frequencies from this MSA and adjusted the empirical frequencies in our helical sequence MSA according to Equations 17 and 18. In these equations, $f^{\text{helix}}$ and $f^{\text{protein}}$ represent amino acid single and coupled frequencies from the MSA containing helical segments of length four and the MSA containing all protein segments of length four, respectively. The sums over $a$ and $b$ are summations over all canonical amino acids. Equation 17 adjusts the empirical frequencies for overrepresentation in helical segments and Equation 18 renormalizes the single and coupled frequencies. The purpose of this adjustment was to account for the fact that certain amino acids are less common in protein sequences in general which should weight their representation in helical segments more highly. The total relative frequency of each amino acid within helical segments compared to its background in the clustered SwissProt Database is presented in SI Fig. 16.

$$\begin{aligned} f_i^*(a) &= f_i^{\text{helix}}(a) \frac{f_i^{\text{helix}}(a)}{f_i^{\text{protein}}(a)}, \\ f_{i,j}^*(a,b) &= f_{i,j}^{\text{helix}}(a,b) \frac{f_{i,j}^{\text{helix}}(a,b)}{f_{i,j}^{\text{protein}}(a,b)}. \end{aligned} \tag{17}$$

$$\begin{aligned} f_i(a) &= \frac{f_i^*(a)}{\sum_a f_i^*(a)}, \\ f_{i,j}(a,b) &= \frac{f_{i,j}^*(a,b)}{\sum_a \sum_b f_{i,j}^*(a,b)}. \end{aligned} \tag{18}$$

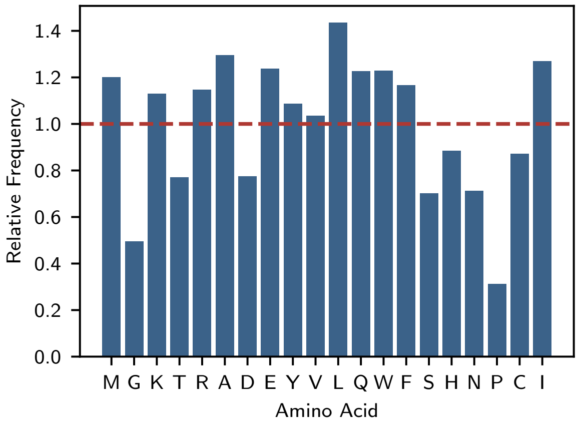


SI Fig. 16. **Relative frequency of amino acids in $\alpha$-helices versus background frequency in proteins**. Displayed is the relative frequency of occurrence for each canonical amino acid in $\alpha$-helices versus their total frequency within proteins. The frequencies were extracted from the 56,474 SwissProt database sequences clustered by mmSeqs2.

After finalizing the empirical training frequencies, we progressed to model training. When training our Potts model, we employed a lattice gas gauge to address overparameterization [12], placing our gauge on alanine, a residue with a very high helical propensity [17]. The equation for the lattice-gas gauge is presented in Equation 19, where we substitute $q$ (the gauged amino acid) with alanine.

$$h_i(q) = J_{ij}(a, q) = J_{ij}(q, a) = 0. \tag{19}$$

With the empirical frequencies from the $\alpha$-helix MSA providing our target distribution, we applied Boltzmann machine learning with direct gradient descent and a basin-hopping approach to train our Potts model. Here, a direct gradient descent approach was feasible given that the protein sequences in the MSA were only of length four. The workflow used Markov chain Monte Carlo (MCMC) to sample the current iteration of Potts model parameters and build observed single and coupled frequencies. Thereafter, the difference between observed frequencies and the empirical frequencies from the underlying MSA was applied using direct gradient descent to update model parameters. Equation 20 describes the direct gradient descent for updating Potts model parameters.

$$\begin{aligned} h_i(a) &\leftarrow h_i(a) + \epsilon(f_i(a) - \langle \delta_{a_i,a} \rangle_P), \\ J_{ij}(a, b) &\leftarrow J_{ij}(a, b) + \epsilon(f_{ij}(a, b) - \langle \delta_{a_i,a} \delta_{a_j,b} \rangle_P). \end{aligned} \tag{20}$$

Where $\epsilon$ is a term set to control the speed of gradient descent.

Our approach was implemented using a custom-built C++/CUDA code. To begin, the starting model parameters for each training run was set to $h_i(a) = -\log f_i(a)$ and all $J_{ij}(a, b) = 0.0$. An iteration of training (i.e., one model parameter set update), constituted 10,240,000 total samples of the Boltzmann distribution built across 100 blocks each with 512 threads running on an NVIDIA A100 GPU. For each iteration, each thread first underwent a 'burning-in' period with 10,000 mutations to a sequence of four amino acids uniquely tracked by each thread. Mutations were accepted based on a Metropolis criterion. Thereafter, each thread produced $\frac{10{,}240{,}000}{512 \times 100} = 200$ sequences to add to the sampled MSA by performing 1,000 mutations between added sequences according to the MCMC scheme. After each iteration, model parameters were updated according to the direct gradient descent approach as outlined previously.

A complete training run for the Potts model consisted of three different phases—optimization phases, search phases, and a final optimization. Optimization phases consisted of 5000 iterations of parameter updates at a temperature of $T = 1.0$, an $\epsilon = 0.25$, and a maximum descent (an enforced upper limit of each parameter's change) equal to 0.1. Between each optimization phase, a search phase (i.e., basin hopping) was undertaken with 100 steps at a temperature of $T = 25.0$, and an $\epsilon = 1.0$ with no maximum parameter change enforced. In total, there were ten optimization phases with search phases between each. Afterwards, a final optimization phase was performed for 5000 steps with a temperature $T = 1.0$, an $\epsilon = 0.025$, and a maximum descent of 0.1. Additionally, at all times in the training process the current best parameter set that provided the greatest agreement with the empirical frequencies was saved. Thus, the final parameters for a given

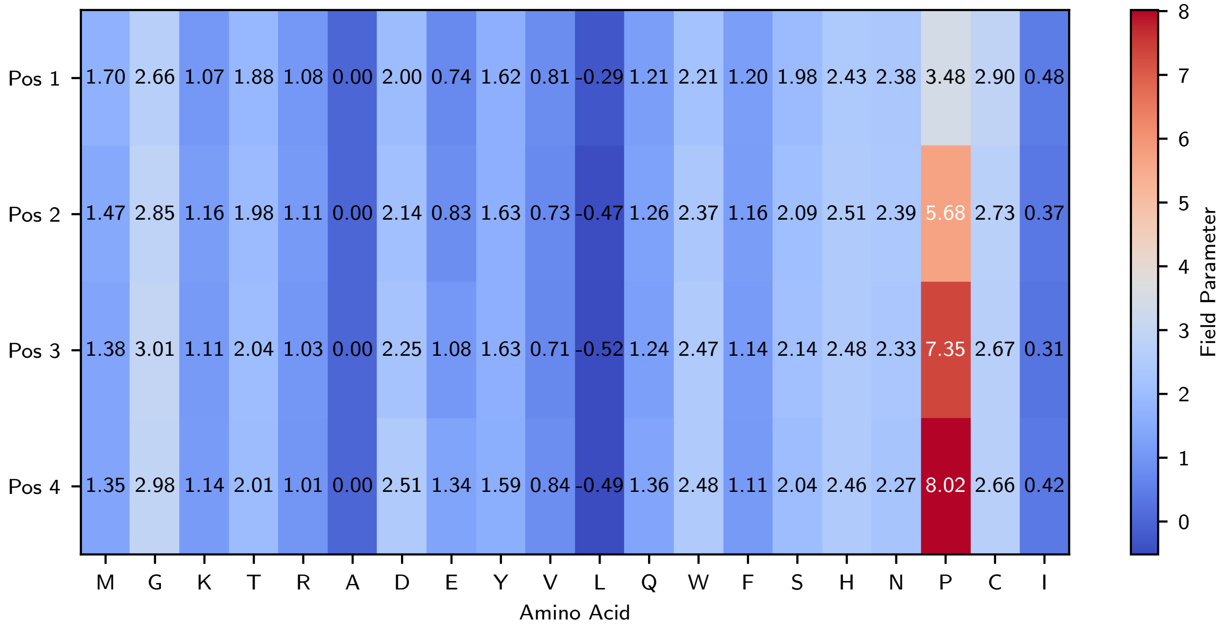


SI Fig. 17. **Field parameters of Potts model.** Amino acid and position specific Potts model parameters.

training run were taken as the best saved parameters. The overall final parameters for the Potts model were taken as the average of ten independent training runs (initialized with unique random number seeds).

These parameters are included as a `.txt` file in the GitHub repository. Additionally, the model parameters are shown in the heatmaps in SI Figs. 17 and 18. The final model parameters appear reasonable for multiple reasons. First, field values for residues such as alanine, leucine, and glutamic acid, which are known helix-forming residues [18], are very low. Conversely, field values for residues such as proline, glycine, and cysteine, known helix-breakers [18], are high. Second, proline pairings which are physically unable to reach the helical state (pos 1 and pos 3, as well as pos 2 and pos 4), have infinite energy, seen by the white squares in SI Fig. 18. Third, charged residues spaced in a manner that allows for salt bridges (pos 1 to pos 3 or 4) have low values. Finally, residues whose helix breaking behaviors are relatively independent of side chains they are paired with within helices (namely proline), have uniformly small magnitudes in their coupling values.

We performed two additional tests to validate our Potts model parameters. The first test explored how well the model could distinguish ‘H’ amino acids and all other types of structural classifications as determined by DSSP. In particular, we took the set of 7,997 protein structures from the PDB extracted via the PISCES PDB culling server [8] and classified each residue in every protein sequence using DSSP. Thereafter, we scored every amino acid in the sequences using the Potts model parameters where the overall score given to a residue was the average Potts energy from the overlapping segments of four amino acids containing that residue. The results are plotted in SI Fig. 19 which shows that the statistical energies for ‘H’ residues from DSSP are lower than any other classification. This provides evidence that our Potts model is trained properly on ‘H’ residues and may even be able to distinguish stretches of $\alpha$-helix residues from other types of secondary structures—although this is not the purpose of our model.

The second test was to explore the trend between Potts energy and the average helical content of segments of four residues from atomistic REST2 [19] simulations of 11 length peptides simulated in the ff19sb [1] force field (see Methods and ‘Dataset from REST2 Enhanced Sampling Atomistic Simulations’ section in the SI for more details on these simulations). The helical content from each segment of four was determined as the average of each constituent residue’s helical fraction via classifications of $\phi$ and $\psi$ backbone angles (following the criterion $-30.0 > \phi > -100.0$ and $-7.0 > \psi > -67.0$). The natural logarithm of the fraction of frames spent in the helical state versus the coil state is then plotted against the Potts energy for each sequence of four. The negative linear trend demonstrates that the Potts model is indeed predicting a correct statistical energy that correlates with helical fraction as simulated in the ff19sb force field. This result can be found in Extended Data Fig. 1f. Further, this linear trend indicates that we should expect a linear relationship between Mpipi-Helix’s well depth and the Potts score for sequences of four amino acids. Both are statistical energies that describe the propensity for a given sequence to be in a helical state as opposed to a coil state. As Boltzmann factors, they will be linearly related, which is also demonstrated by the accuracy of the Mpipi-Helix model following parameterization.

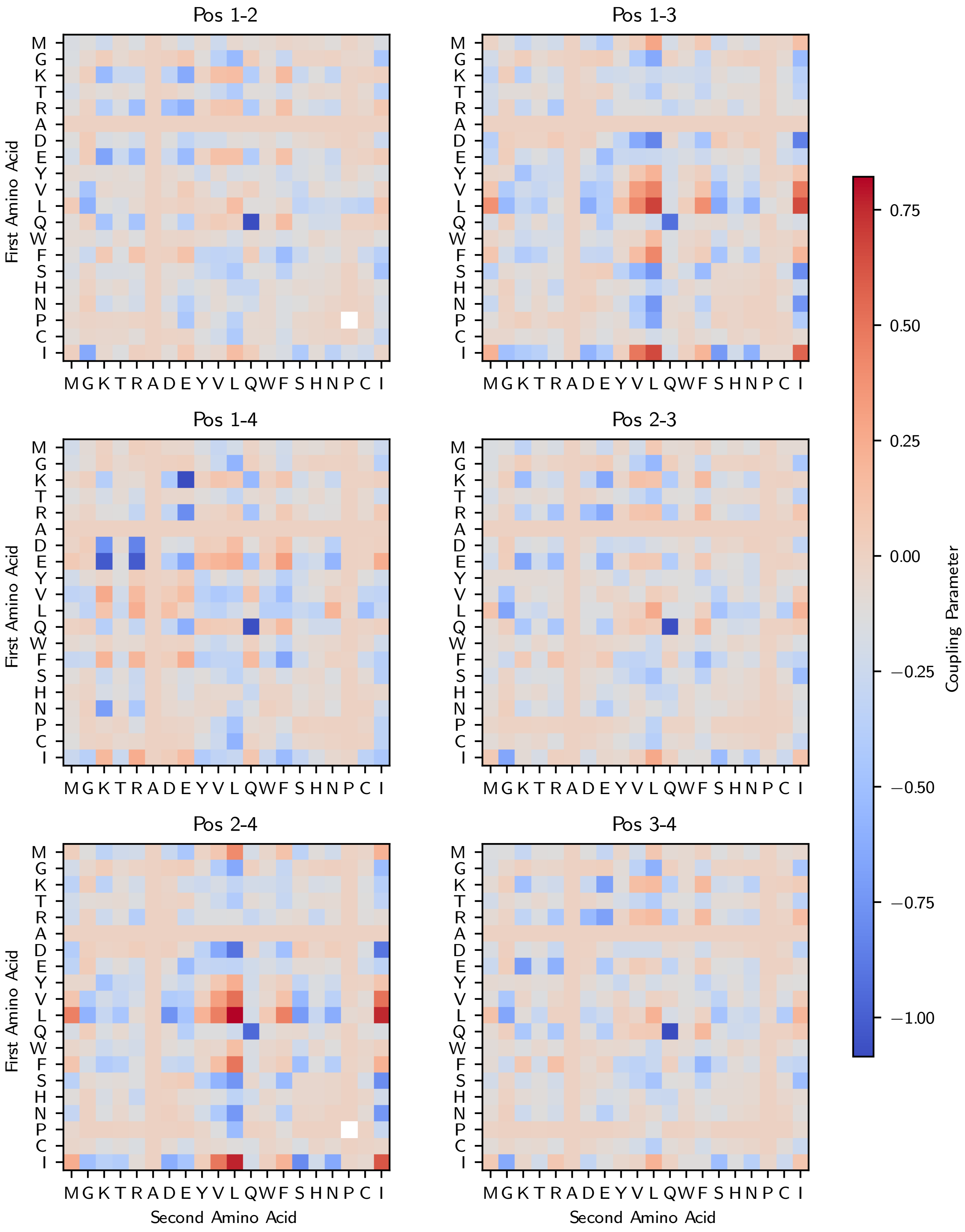


SI Fig. 18. **Coupling parameters of Potts model.** Amino acid pairing and position specific Potts model parameters.

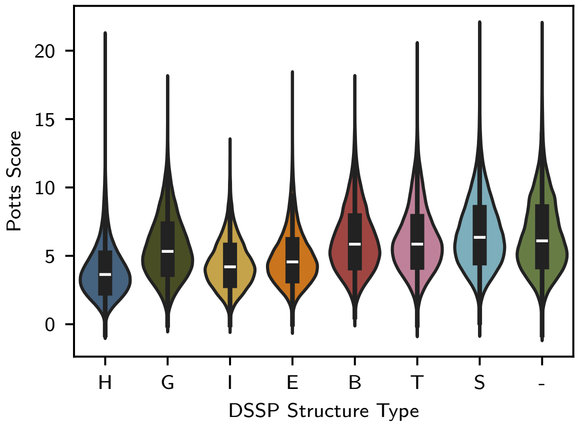


SI Fig. 19. **Potts energies for residues taken from PDB structures and classified with DSSP.** We extracted DSSP classifications [9] from the 7,997 structures we culled from the PISCES PDB culling server. Thereafter, we calculated the average Potts energy for each residue. The distribution of Potts scores from each DSSP structure type demonstrates the Potts energies are lowest for predicted helical 'H' structures.

##### F. Gaussian Process Training

We developed a Gaussian process (GP) workflow to generate our thermodynamic trend between Potts scores and well depths in Mpipi-Helix. The workflow is depicted in Extended Data Fig. 1g-i of the main text. The goal of this workflow was to predict the well depth value in Mpipi-Helix that would reproduce the helical propensities taken from experimental NMR measurements and atomistic simulations. Specifically, we sought to match the helical propensities of alanine-based peptides recorded with NMR measurements at 25 °C in Moreau et al. [18], as well as the helical propensities of peptides simulated in the ff19sb force field [1] using REST2 enhanced sampling [19]. Details on how the helical propensity of each of these datasets was calculated and implemented in the GP workflow are provided in the sections 'Helix–Coil Classification in the GP Workflow', 'Dataset From Experimental NMR Measurements', and 'Dataset from REST2 Enhanced Sampling Atomistic Simulations'. Here, we describe the aspects of the GP workflow that apply to both datasets.

Our GP workflow began by selecting a peptide from one of the two datasets. Five well depth values were provided as inputs (for all sequences parameterized here, we provided a well depth $W = 1.0, 1.5, 2.0, 2.5,$ and 3.0) which launched five separate coarse-grained simulations in Mpipi-Helix. This constituted an initial seed for the GP Bayesian optimization process. In the case of the REST2 enhanced sampling simulations, a single input well depth was applied to the entire peptide. For the NMR sequences, the all alanine peptide had a single well depth applied to the whole peptide, but for the rest of the guest–host peptides the well depth only applied to the segments of four amino acids that included the guest residue with the well depth of the all-alanine caps being fixed. We applied the well depths in such a manner despite the Potts scores deviating along the peptide sequence (e.g., ABCD $\neq$ BCDA in terms of Potts score; similarly, AAXA $\neq$ AXAA in terms of Potts score). This was done for computational efficiency as the cost of GP predictions increases significantly as dimensions are added to the optimization process and more noise is introduced.

The coarse-grained simulations were run in the $NVT$ ensemble at 298.15 K using the Langevin thermostat with a damping of 100 ps and a timestep of 10 fs in the LAMMPS software [6]. The simulations were run with a 1 ns equilibration time and a 1 µs sampling time with simulation frames saved every 0.05 ns. After the simulation was completed, residues in the simulated peptide were assigned helix or coil states according to the Mpipi-Helix coarse-grained helix–coil assignment rule (see section 'Helix–Coil Classification in the GP Workflow'). Using the assignments, a loss function scored each simulation for how well the helical propensity matched the target from the original dataset (discussed in the relevant sections on 'Dataset From Experimental NMR Measurements' and 'Dataset from REST2 Enhanced Sampling Atomistic Simulations').

After the batch of simulations was completed, we fed in the scores for each well depth value tested so far into a GP. The GP was defined by a zero mean function and an RBF kernel with a Gaussian prior on the length scale (mean of 0.5 and standard deviation of 0.1). Inputs into the GP were scaled using scikit-learn's standard scaling function [20]. Training was performed using an Adam optimizer with a learning rate of 0.1 and 50 training iterations. After the GP was trained, an acquisition function was used to select a new batch of training points (i.e., 5 new well depths for further simulations). For the first 100 batches, an expected improvement acquisition function was used (Equation 21 where $W$ is the well depth, $\Phi$ is the cumulative distribution function, $\phi$ is the standard normal probability density distribution, $\mu(W)$ is the mean predicted value at $W$, $\sigma(W)$ is the predicted standard deviation at $W$, $f(W^+)$ is the highest scoring $W$ value predicted so far, and $\mathrm{EI}(W)$ is the expected improvement acquisition function evaluated at $W$). Following this, for the next 100 batches an upper confidence bound acquisition function was used (Equation 22 where $\xi$ is set to 0.1 and $\mathrm{UCB}(W)$ is the upper confidence bound acquisition function evaluated at $W$). During batch selection for each acquisition function, well depth points were chosen sequentially with a negative bias applied to the acquisition function value for all previously chosen points in the batch. This negative bias took the form of a Gaussian with a mean centered at the chosen well depth and a random variance selected between 0.005 and 0.015. Additionally, for computational efficiency we restricted possible well depth values that we could simulate to be an evenly spaced grid from 0.00 to 5.00 kcal/mol in a spacing interval of 0.01 kcal/mol.

$$\begin{aligned} \mathrm{EI}(W) &= \sigma(W)[Z\Phi(Z) + \phi(Z)], \\ Z &= \frac{\mu - f(W^+)}{\sigma(W)}. \end{aligned} \tag{21}$$

$$\mathrm{UCB}(W) = \mu(W) - \xi\sigma(W) \tag{22}$$

Data collection was terminated after 1,000 total simulations were performed (200 batches) for all peptides besides the GTNI, GGKE, and GAAA peptides from the REST2 dataset. These peptides were sampled for an additional 40 batches with the UCB acquisition scheme due to insufficient sampling in the region of the maximum. In these 40 batches, sampling was restricted to the region of 3.0 to 3.21 kcal/mol.

After 1,000 total data points were collected, a GP was used to predict the best overall data point from the entire simulation set. For this final prediction, the input data had no scaling, the GP had a constant mean, and an RBF kernel with a length

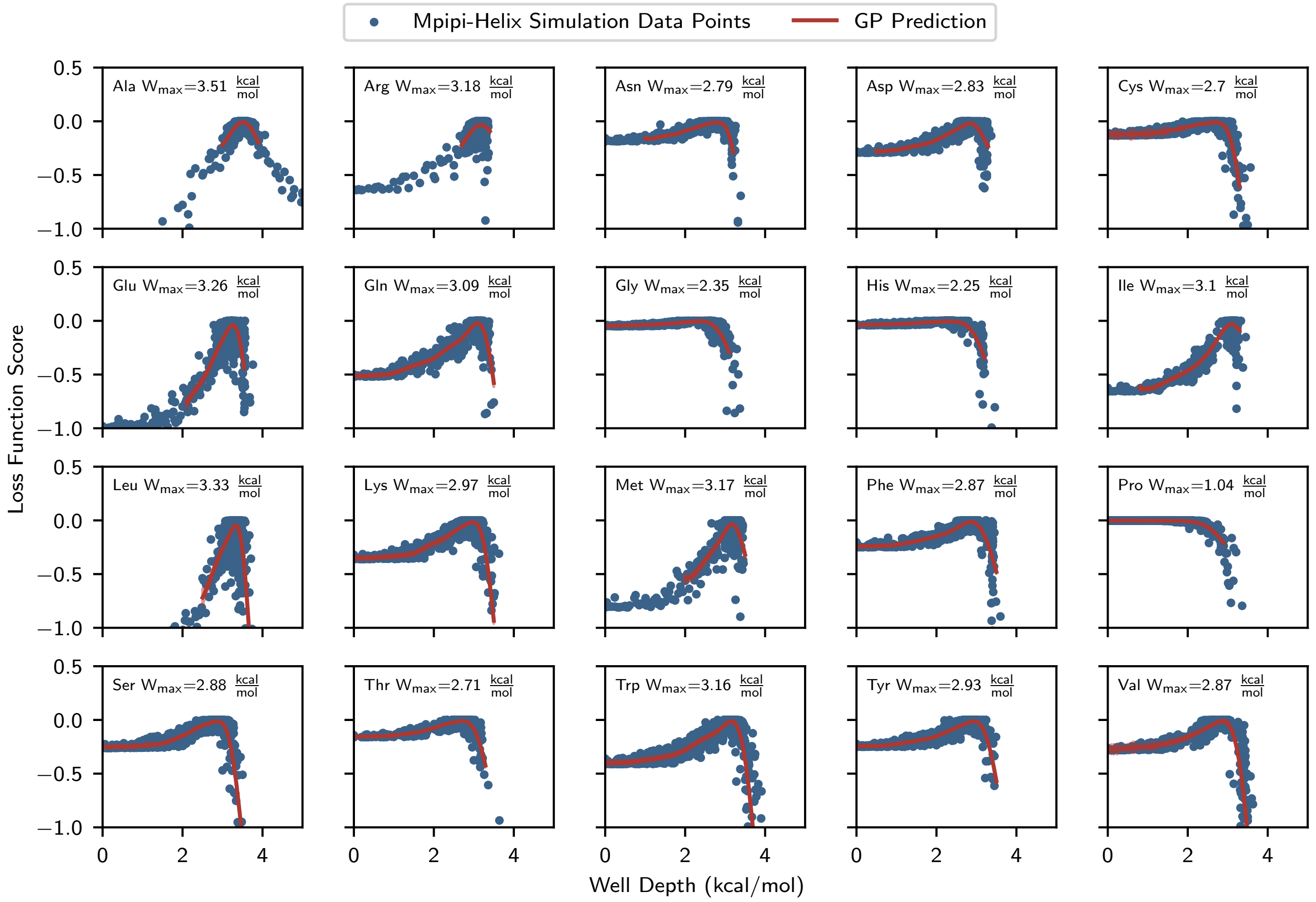


SI Fig. 20. **GP predictions on Mpipi-Helix simulations: Moreau 25°C.** The optimal well depth value for the Moreau 25°C dataset as determined by GP fitting.

scale enforced to be between 0.2 and 1.0 was used. Due to the mean squared error nature of the loss functions we used (see sections 'Dataset From Experimental NMR Measurements' and 'Dataset from REST2 Enhanced Sampling Atomistic Simulations'), the data was particularly noisy far away from the well depth values that maximized the loss function. This caused problems with the stability of the GP prediction due to overfitting of noisy outlier data. To address this, for many of the peptides we restricted the region we were fitting to include only data points which were close to the maximum. All training was performed using an Adam optimizer with a learning rate of 0.1 and 100 iterations. Additionally, we ran every dataset through the GP prediction 100 times with a different seed each time taking the average result as the optimal well depth value. Results for the GP training are show in SI Figs. 20, 21, 22, and 23, where the simulated data points are depicted in blue and the GP prediction is depicted in red. For direct comparisons between the helical fractions and the Lifson–Roig *w* values for the final well depths, the final Mpipi-Helix parameterization, and the training datasets, see the sections 'Dataset From Experimental NMR Measurements' and 'Dataset from REST2 Enhanced Sampling Atomistic Simulations'. All GP/Bayesian optimization scripts for this study were built in-house using the GPyTorch python library [21] and they are available on the GitHub repository. Additionally, the scripts for determining the optimal well depth are also available on the GitHub repository.

Once we determined the optimal well depth values for every sequence, we found our final well depth versus Potts score trend using linear fits to the combined GP datasets. A linear fit is reasonable to describe the overall well depth versus Potts score trend given the linear Potts score to helical fraction relationship (Extended Data Fig. 1f).

Our first step in the fitting was to perform a linear fit on the Moreau et al. data alone to determine the y-intercept of our final linear fit, as seen in SI Fig. 24a. We performed this fitting on the NMR dataset as it does not suffer from the possibility of undersampling and force field biases. Additionally, as alanine has a high helical propensity among amino acids [17], we reasoned that taking the y-intercept of a fitting with alanine-based guest–host peptides would describe helices with low Potts scores well. However, the high helical propensity of alanine also amplifies the effects of helix-breaking residues [17, 22], which is why we turned to a combined fitting of all the training data for the slope of our linear fit.

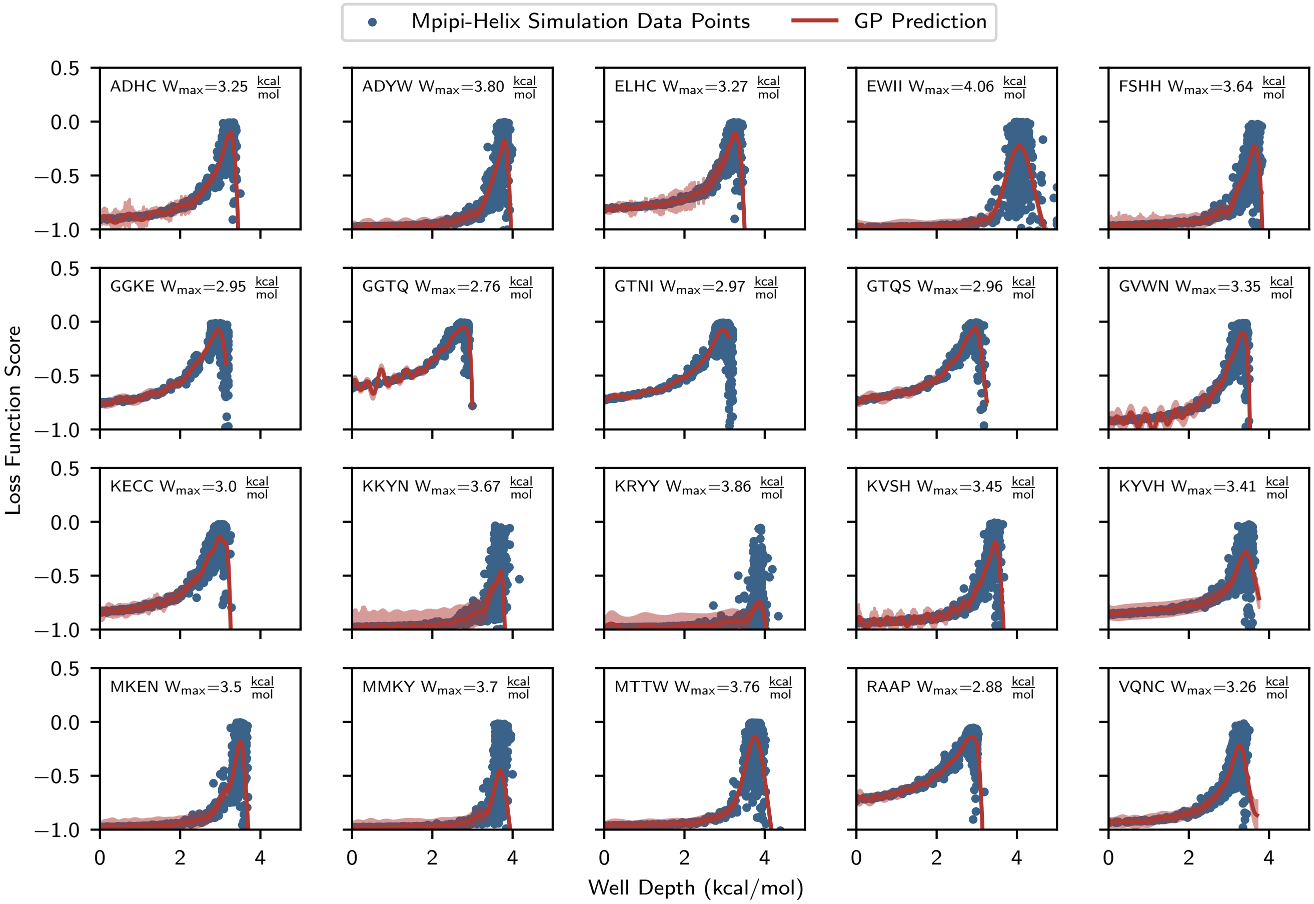


SI Fig. 21. **GP predictions on Mpipi-Helix simulations: REST2 mean.** The optimal well depth value for the REST2 mean Potts score dataset as determined by GP fitting.

Before fitting to all the data to determine our slope, we first shifted the ff19sb REST2 data downwards in well depth to achieve a maximum overlap with the experimental NMR dataset. This was necessary as the ff19sb force field has been shown to overpredict the helical propensity of peptides in comparison with the Moreau et al. dataset [1]. The two datasets before shifting are plotted in SI Fig. 24b. To shift the REST2 sampled dataset, we used the overlap in a kernel density estimation (KDE) of the minus two standard deviations subgroup and the Moreau et al. dataset. As these two groups of sequences have similar Potts scores, in our final trend they should be highly overlapping. Our shifting followed two steps. First, we removed the GAAA sequence from the minus two standard deviations group as well as the H, G, and P guest–host sequences from the Moreau group. These were removed as they are the largest outliers at low well depths within the datasets. Second, we performed a shifting whereby we evaluated the KDE overlap between the two sets of points using scipy's `scipy.stats.gaussian_kde` with a bandwidth of 0.2 and `scipy.integral.simps` functions. The best overlap was selected from shifting values tested between 0.0 and 1.0 in an interval of 0.01. The best value was found to be a downward shift of 0.37 kcal/mol. SI Fig. 24c shows the shifting process with the data points before shifting, as well as the removed GAAA, H, G, and P sequences. SI Fig. 24d shows the maximum overlap from the 0.37 $\frac{\text{kcal}}{\text{mol}}$ with the KDE contour lines for each dataset.

After shifting, we performed a linear fitting for the slope of the well depth versus Potts score trend. First, however, we removed the most strongly helix-breaking residues, including P from the Moreau dataset and TRPC, GTHP, HSPC, TWPC, and GGSP from the REST2 plus two standard deviations dataset. This approach is justified given that the purpose of Mpipi-Helix is to apply a helical potential to sequences that we know should be helical (e.g., from experimental measurements or atomistic simulations). If we want to describe sequences that we know are going to be helical (e.g., helical enough to observe using experimental methods), then we should not bias our parameterization dataset with sequences that have very low helical fractions. Further, validation of the model demonstrates that even after removing these helix-breaking residues, the sequence dependence of Mpipi-Helix is still good enough to describe helix-breaking mutations (see 'Mpipi-Helix Model Validation' below). This is likely the case because sequences with a helix-breaking mutation may be represented in the

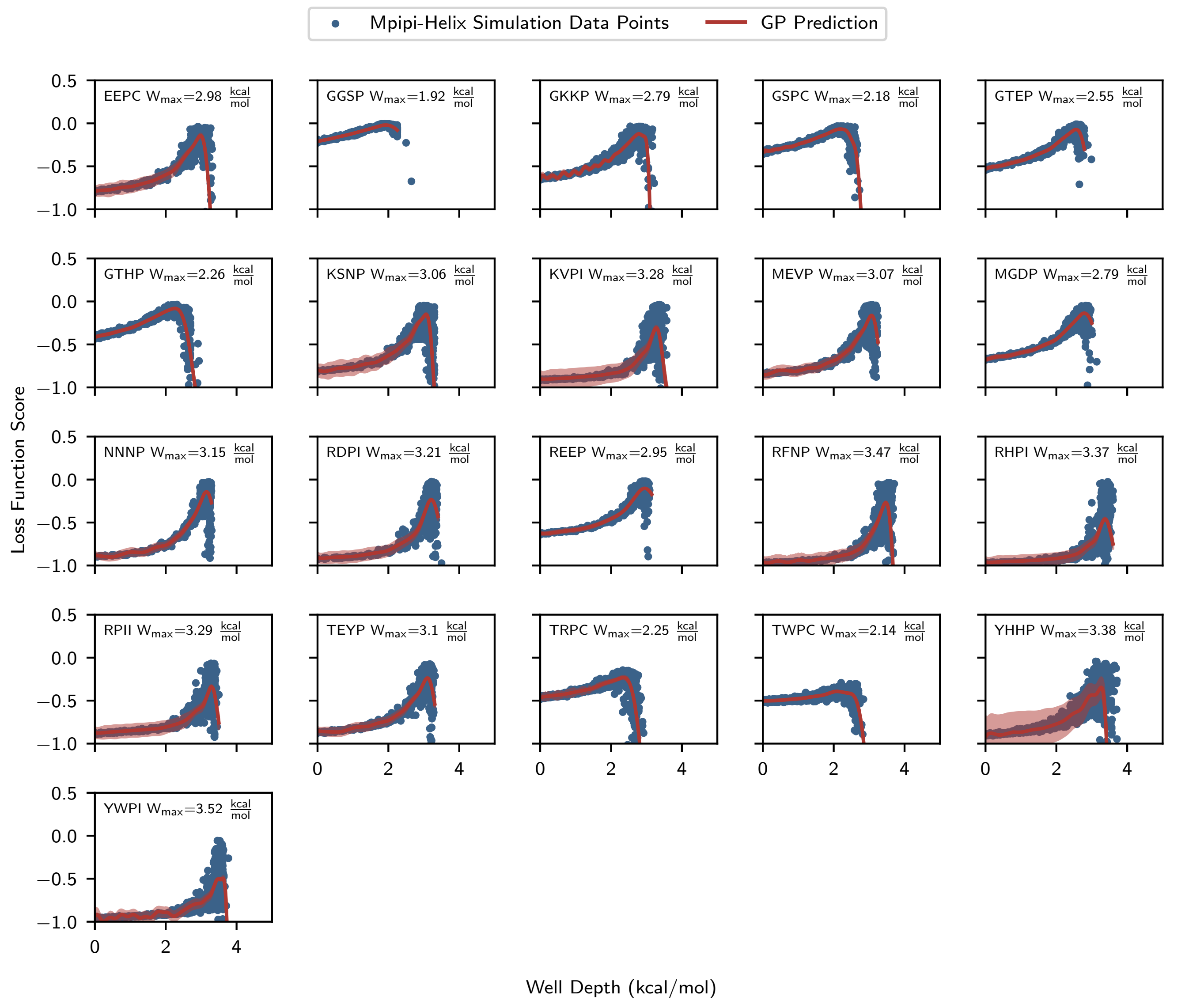


SI Fig. 22. **GP predictions on Mpipi-Helix simulations: REST2 plus two standard deviations.** The optimal well depth value for the REST2 plus two standard deviations Potts score dataset as determined by GP fitting.

underlying $\alpha$-helix MSA that trained the Potts model whereas sequences involving multiple repeated proline or glycine residues (i.e., the sequences removed from the plus two standard deviations dataset) are unlikely to be classified as a helix at all. The residues that were removed from the final fitting are annotated in SI Fig. 24e. SI Fig. 24f shows the linear fit on the combined datasets without the helix-breaking residues. The $R^2$ value for the linear fit likely suffers from the liberties taken in the GP fitting to the REST2 data (e.g., fitting a single well depth value to each sequence) and the fact that we are combining two different datasets, one taken from atomistic simulations and the other taken from NMR experiments. Fitting just the REST2 data alone produces a much stronger trend, as shown in SI Fig. 24g with an $R^2 = 0.48$ value with the helix-breakers included and an $R^2 = 0.32$ value with helix-breakers excluded (not shown). This more closely matches the stronger linear relationship for the atomistic helicity of the REST2 sequences plotted in Extended Data Fig. 1f.

Overall, the combined final trend we achieved is $W = 3.45 - 0.04P$ where $W$ is the well depth and $P$ is the Potts score of the input sequence. The two individual trends are plotted on the same plot in SI Fig. 24h.

Finally, we want to highlight again that our philosophy for this model is to apply the $\alpha$-helix potential to regions of known helical content. We believe there are better tools for predicting helical regions than coarse-grained simulations (e.g., atomistic simulations, NMR measurements, and CD measurements). However, given that we know which regions have helical content in a protein, our coarse-grained model can be effectively used to probe the characteristics of that helix

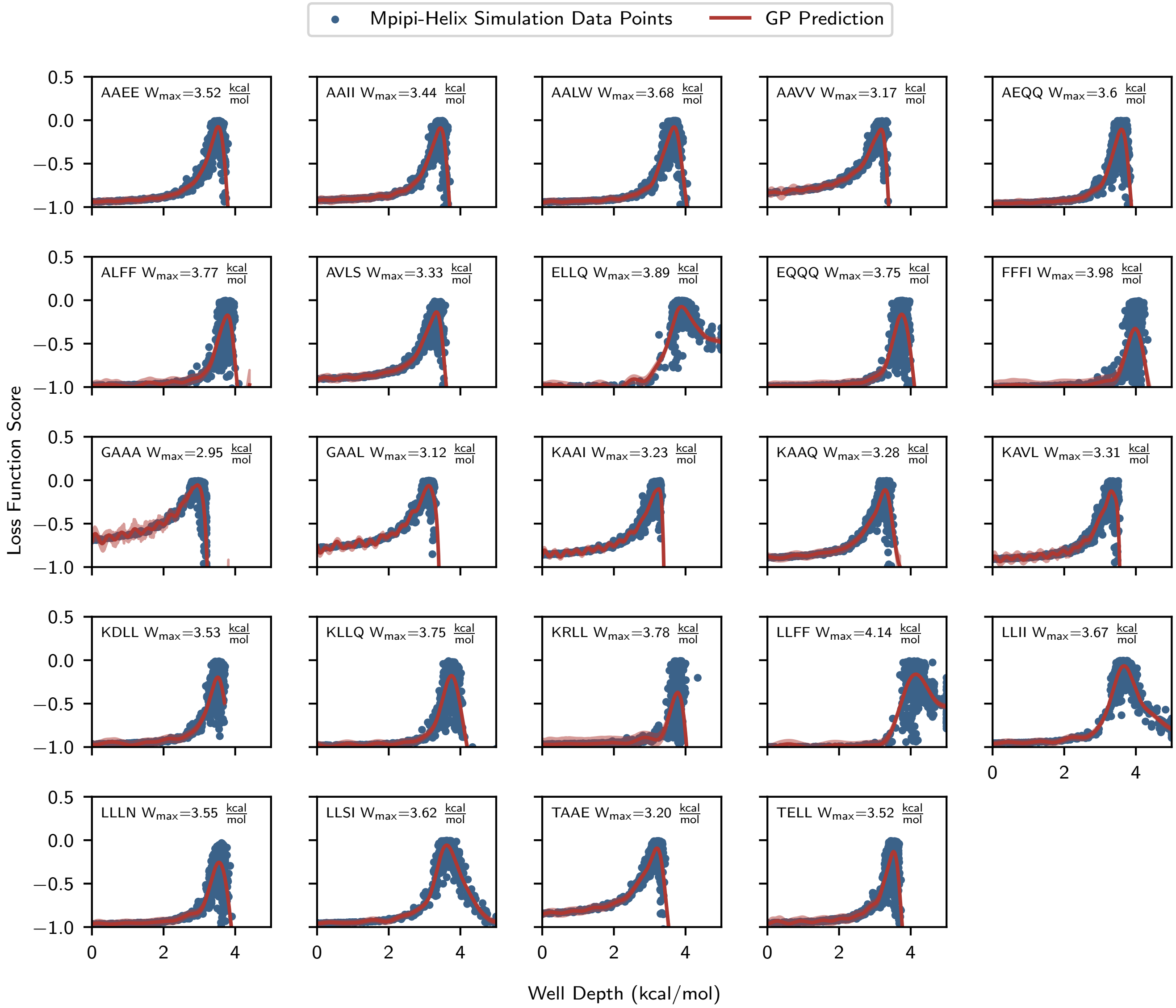


SI Fig. 23. **GP predictions on Mpipi-Helix simulations: REST2 minus two standard deviations.** The optimal well depth value for the REST2 minus two standard deviations Potts score dataset as determined by GP fitting.

within condensates which may be difficult to probe using the aforementioned tools. This is the focus of our model and this manuscript. This also justifies our approach to model validation, where we validate against the helical content of disordered proteins with known helical domains (see 'Mpipi-Helix Model Validation' below).

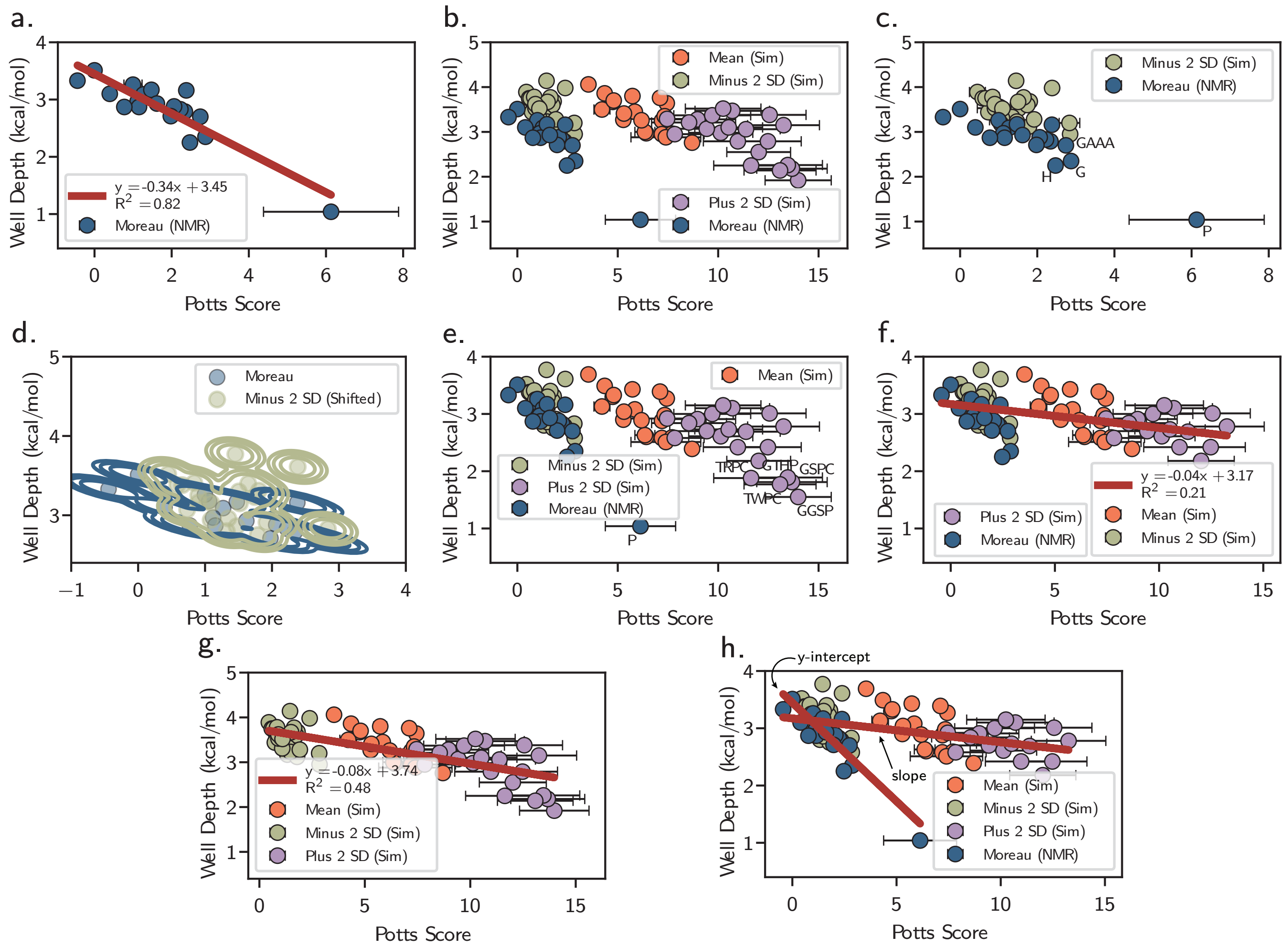


SI Fig. 24. **Trends and shifting from GP results. a,** The y-intercept of the final well depth versus Potts score was taken from a linear fit of the Moreau et al. [18] experimental NMR dataset's well depth values parameterized by our GP. **b,** All GP training optimal well depth values for all sequences. **c,** Minus 2 SD dataset was shifted downwards for maximum overlap with the Moreau dataset using kernel density estimation (KDE). GAAA (from Minus 2 SD), H, G, and P (from Moreau) sequences were removed for the shifting. **d,** Shifting produced from KDE overlap of Minus 2 SD and Moreau. The Minus 2 SD dataset was shifted downwards by 0.37 $\frac{\text{kcal}}{\text{mol}}$ in well depth. **e,** The GP training datasets after shifting all REST2 enhanced sampling downwards by 0.37 kcal/mol. Labeled points were those removed from the linear fitting to achieve the slope of the well depth versus Potts score trend. These included P (Moreau), TRPC, GTHP, GSPC, TWPC, and GGSP (from Plus 2 SD). **f,** The value of the slope for the final well depth versus Potts score trend was achieved from a linear fit. **g,** The linear fit of the REST2 data alone (with helix-breakers). **h,** The two linear fits for the final well depth versus Potts score trend. The lines are annotated with y-intercept and slope to indicate which portion of the combined linear fit we took from each.

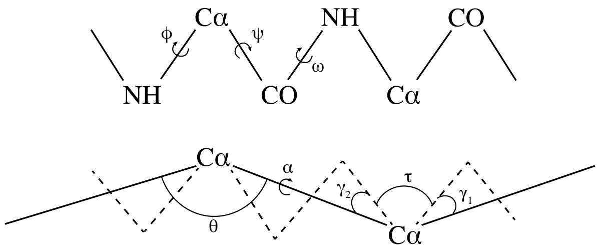


SI Fig. 25. A diagram of the α-carbon to all atom mapping reproduced from Ref. [30]. Back mapping is accomplished through iteratively solving Equations 23 and 24.

##### G. Helix–Coil Classification in the GP Workflow

A critical component of the GP workflow is to calculating helical fractions for every residue in an Mpipi-Helix simulation, which requires labeling whether residues are in a helix or coil state at every simulation frame. This task is challenging to do in an accurate and computationally efficient manner as the $\phi$ and $\psi$ backbone angles which are usually used to determine a protein's secondary structure are hidden at a residue-level resolution. However, the task of predicting secondary structure directly from an α-carbon trace [23–27] or via backmapping techniques [28, 29] has been a topic of much research. Previous studies have provided computational tools to streamline the process of obtaining secondary structures from α-carbon traces.

Despite this, there are convincing reasons why these techniques are not suitable for our parameterization process which requires a large number of simulations to be completed as quickly and accurately as possible. In regard to computational efficiency, backmapping tends to be more expensive than we would like for our GP workflow, given that we need to run on the order of tens of thousands of simulations to parameterize all of our protein sequences. Additionally, the softwares that directly map from an α-carbon trace to a secondary structure are not benchmarked for short peptides, as they were trained on much larger PDB structures and validated on these larger proteins (which are also static structures unlike our dynamic trajectories). This is a by-product of the tools being principally designed to determine the structures of proteins that have stretches of missing side chain and backbone atoms from experimental characterizations. The cg2all software [28] stands out as the best and most efficient software from all of the aforementioned methods. Indeed, we use cg2all to classify helical residues in our validation and subsequent data collection simulations with Mpipi-Helix. However, we were unable to use cg2all in our GP parameterization process as the software became available after our parameterization approach had been finalized and was in the process of running. We provide benchmarks of our classification method against cg2all below.

For the aforementioned reasons, we developed our own technique to map from an Mpipi-Helix α-carbon trace to a helix–coil prediction for every residue in a given peptide (hereafter, the MH method). To accomplish this task, we combined the geometrical criteria of helix formation in our α-helix potential with equations for calculating $\phi$ and $\psi$ angles proposed by Tozzini et al. [30]. For the former criteria, we checked whether a given set of four consecutive amino acids had dihedral and bond angles that reside inside the peak of the helical barrier ($1.33 \leq \theta \leq 1.895$ radians and $0.246 \leq \alpha \leq 1.558$ radians). If this was true, then both of the middle residues would be assigned a helix as they correspond closely to an α-helix geometry (see Extended Data Fig. 1c). The latter criteria, proposed by Tozzini et al. [30], directly maps the $\theta$ and $\alpha$ angles from an α-carbon trace to backbone $\phi$ and $\psi$ angles. The set of equations to accomplish this are shown as Equations 23 and 24 and is depicted pictorially in SI Fig. 25.

$$\cos(\theta_i) = \cos(\tau)[\cos(\gamma_1)\cos(\gamma_2) - \sin(\gamma_1)\sin(\gamma_2)\cos(\phi_i)\cos(\psi_i)] - \sin(\gamma_1)\sin(\gamma_2)\sin(\phi_i)\sin(\psi_i) + \sin(\tau)[\cos(\psi_i)\sin(\gamma_1)\cos(\gamma_2) + \cos(\phi_i)\cos(\gamma_1)\sin(\gamma_2)]. \tag{23}$$

$$\alpha = \phi_i + \psi_i + \pi + \gamma_1\sin(\psi_i) + \gamma_2\sin(\phi_i) + \frac{1}{4}\gamma_1^2\sin(2\psi_i) + \frac{1}{4}\gamma_2^2\sin(2\phi_i) + \gamma_1\gamma_2\sin(\phi_i+\psi_i) - \gamma_1(\tau - \frac{\pi}{2})\sin(\psi_i) - \gamma_2(\tau - \frac{\pi}{2})\sin(\phi_i). \tag{24}$$

These equations assume that $\gamma_1 = 20.7°$, $\gamma_2 = 14.7°$ and, $\tau = 111°$. For Equation 24, it is further assumed that $\phi_i = \phi_{i+1}, \psi_i = \psi_{i+1}$. Because of these assumptions, we used an iterative approach to self-consistently solve for $\phi_i$ and $\psi_i$. If the solutions agreed within 0.1 radians of the original values of $\alpha$ and $\theta$, we considered the previous assumptions to be true and we could accurately assess the $\phi$ and $\psi$ backbone angles. If those were within a helical range on a Ramachandran plot (classifed as $-30.0 > \phi > -100.0$ and $-7.0 > \psi > -67.0$, as used in Ref. [31]) then that residue would be considered helical. If the Equations 23 and 24 failed to converge and match the known values of $\alpha$ and $\theta$ or if the values of $\phi$ and $\psi$ were outside the helical range, this part of the classifier classified residue *i* as being in a coil state. Additionally, for residues in the middle of a peptide chain, there are two dihedral angles associated with the residue. Therefore, we considered that if either $(\alpha, \theta)$ pairing were iteratively solved for and within the helical range, that residue was considered helical. A final benefit of the Tozzini et al. approach to calculating $\phi$ and $\psi$ angles is that residues could be classified into a 'lonely' helical state (i.e., a residue in helical state surrounded on either side by residues in a coil state). This is a critical state to capture for accurately describing helix nucleation and growth, as well as describing helix–coil transitions in standard theories (e.g., Lifson–Roig theory, see section 'Dataset From Experimental NMR Measurements').

The final step in the classification was to combine the classifier that leveraged our helical potential with that predicted from the Tozzini et al. equations. For this task, we used a rule that if either classifier gave a residue a classification of helix, then we considered that residue to be helical, otherwise we would say it was in a coil state.

We validated our MH classification approach using seven different atomistic simulation datasets. We also classified these datasets using the cg2all software for an additional comparison. Three of these datasets were simulations of the 20 guest–host $A_5XA_5$ peptides (where X is replaced with each of the 20 canonical amino acids) in the ff19sb [1], ff14sb [32] and the CHARMM36m [33] force fields. These simulations were performed for 10 µs at 300 K in the *NVT* ensemble (simulated in GROMACS [34] and with input files generated via [35]) at 150 mM NaCl in OPC water (see the 'Mpipi-Helix Model Validation' section for more details on these simulations). Three more datasets were the ff19sb REST2 sampled peptides described in the main text, methods, and the 'Dataset from REST2 Enhanced Sampling Atomistic Simulations' section of this SI. The final dataset was simulations of short helical peptides with initial structures taken from the PDB or generated using PyMol [36]. These peptides were simulated using GROMACS [34] with inputs produced using CHARMM-GUI [35]. The systems were first minimized from their starting configuration using 5000 steepest descent steps with a force tolerance of 1000 kJ/mol/nm and positional restraints on the heavy atoms of the protein. Next, the peptides were equilibrated for 0.125 ns using a 1 fs timestep with a Nosé–Hoover thermostat at a temperature of 300 K and a time constant of 1 ps. Position restraints were maintained on the heavy atoms of the protein and the LINCS algorithm [37] was applied to bonds involving hydrogen atoms. The peptides were then sampled for 5 µs with the same thermostat, a timestep of 2 fs, and frames saved every 0.1 ns. A PME grid was used for electrostatics and the cutoffs for short-ranged neighbor lists, van der Waals interactions, and coulomb cutoff of 1.2 nm for CHARMM36m and 0.9 nm for ff14sb and ff19sb. The PDB files simulated were 2dng, 1kdl, 2a1c, 2pco, and 2dx3, as well as the sequences $A_{21}$ and $A_4(A_4R)_3A_2$ each in the ff19sb [1], ff14sb [32], and CHARMM36m [33] force fields. All peptides were simulated in 150 mM NaCl with TIP3P water for ff14sb and CHARMM36m and OPC water for ff19sb. The residue specific helical fractions for these datasets are displayed in SI Figs. 26, 27, 28, 29, 30, 31, and 32 where the atomistic helical fractions were determined directly from the simulation using a cutoff of $(-30.0 > \phi > -100.0$ and $-7.0 > \psi > -67.0)$ for the helical range. For each peptide, the Mpipi-Helix helical fractions were determined using the aforementioned MH method on the extracted α-carbon trace. Similarly, the cg2all backmapped helical fractions were determined using cg2all to backmap the $\alpha$-carbon coordinates of the atomistic simulation and classify residues using the same $\phi$ and $\psi$ range as above.

From these figures, it is clear that both classification techniques agree very well for the 11 length peptides from the first six datasets. Where cg2all clearly does better than the MH classification method is on the dataset taken from PDB peptides (SI Fig. 32). This seems to indicate that cg2all does better than the MH method for proteins with higher sequence complexity than those we parameterized our the Mpipi-Helix model on. The parity plots for the Mpipi-Helix based classification technique and the cg2all technique are displayed in SI Fig. 33a and b, respectively. While both techniques have a low mean absolute error (3.8% for Mpipi-Helix based classification and 1.4% for cg2all backmapping), it is clear that cg2all is superior. Further, it appears that the MH classification method mostly overpredicts helical content compared to the ground truth. Therefore, outside of the parameterization of Mpipi-Helix, all other classification tasks in this study are performed with the cg2all backmapping strategy.

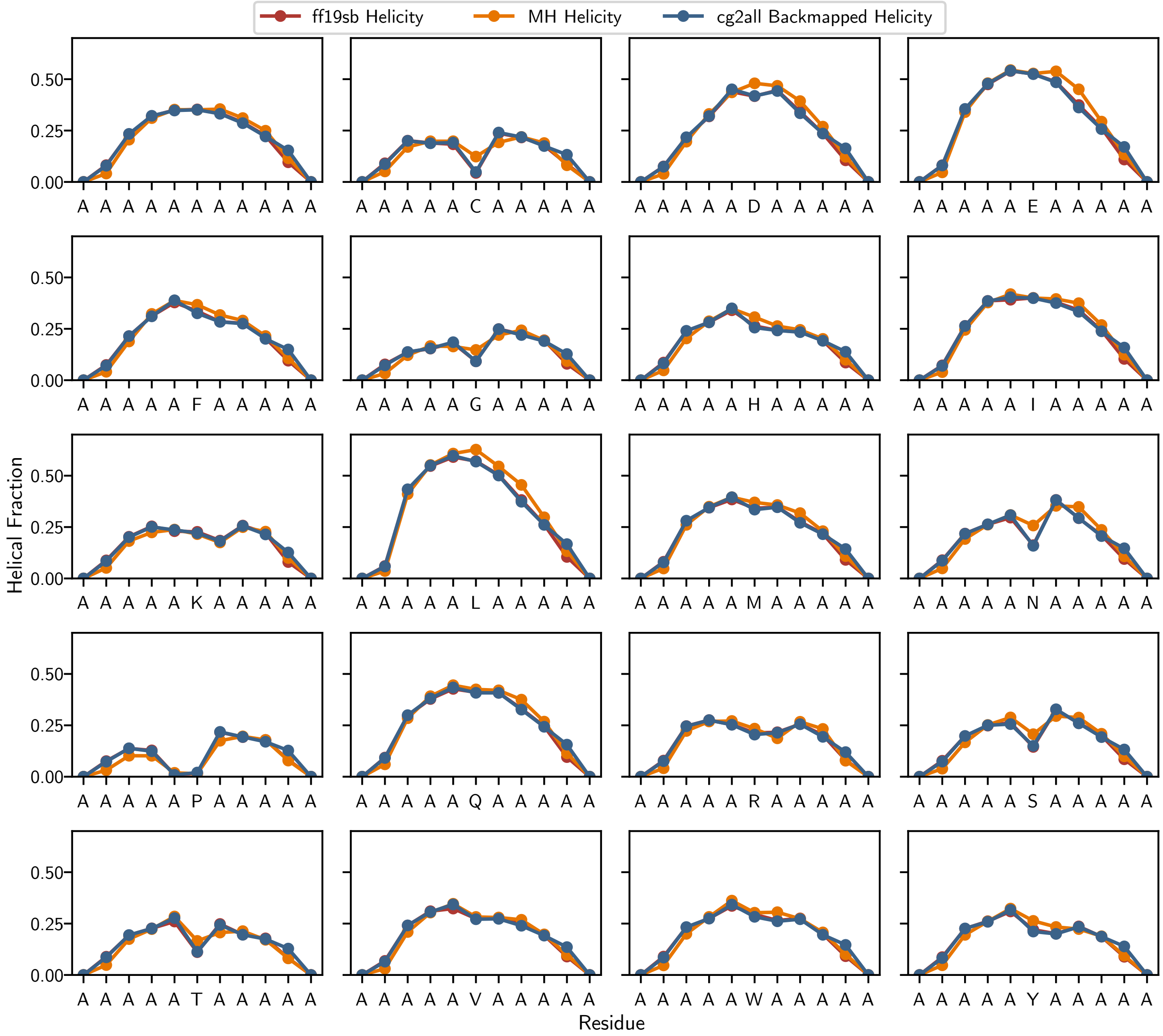


SI Fig. 26. **Helix–coil classifications on $A_5XA_5$ peptides in the ff19sb force field.** The atomistic ground truth helical fractions in the ff19sb force field for the $A_5XA_5$ peptides are shown in red. In orange and blue are the MH classification and cg2all backmapped classification methods, respectively.

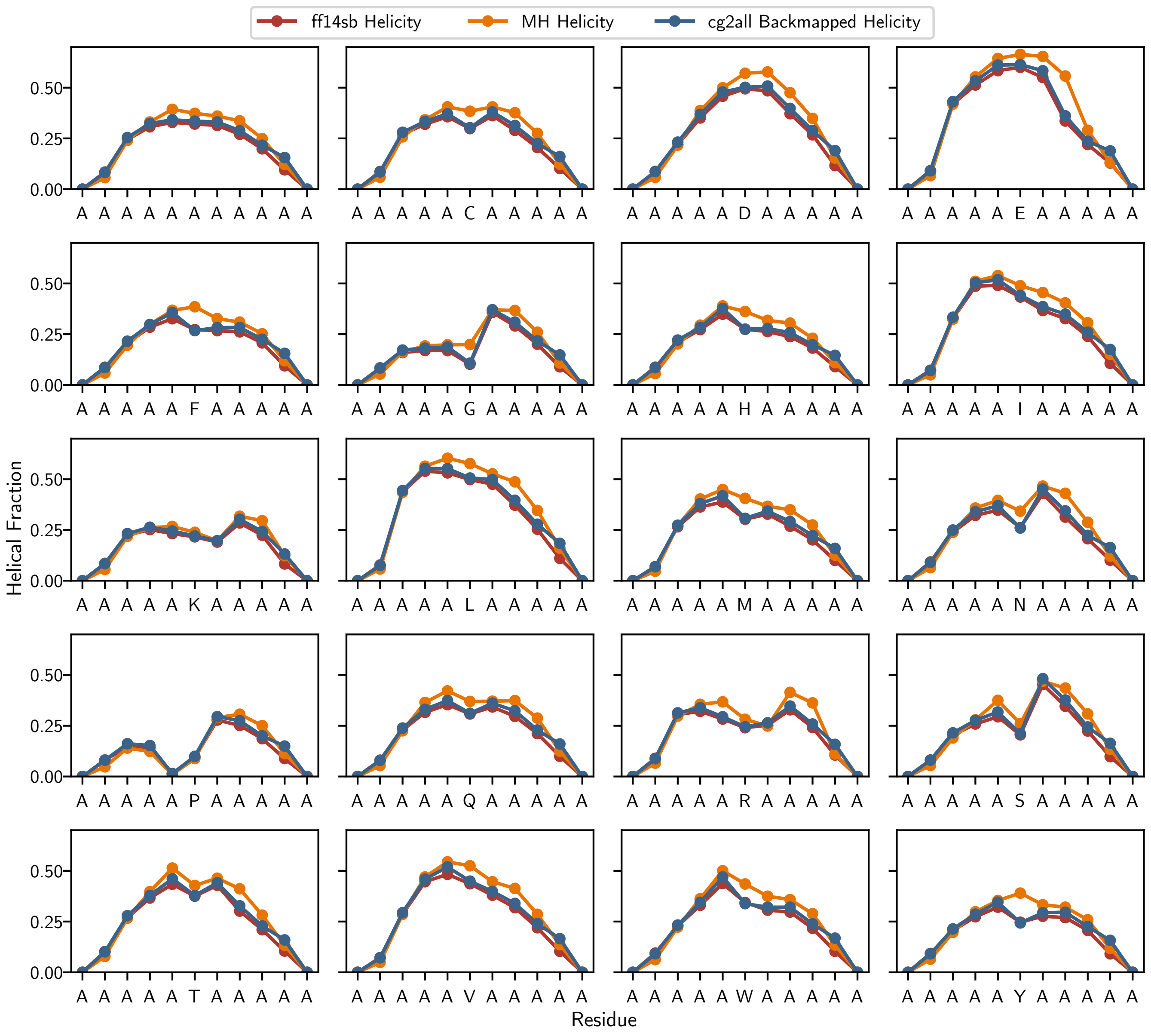


SI Fig. 27. **Helix–coil classifications on $A_5XA_5$ peptides in the ff14sb force field.** The atomistic ground truth helical fractions in the ff14sb force field for the $A_5XA_5$ peptides are shown in red. In orange and blue are the MH classification and cg2all backmapped classification methods, respectively.

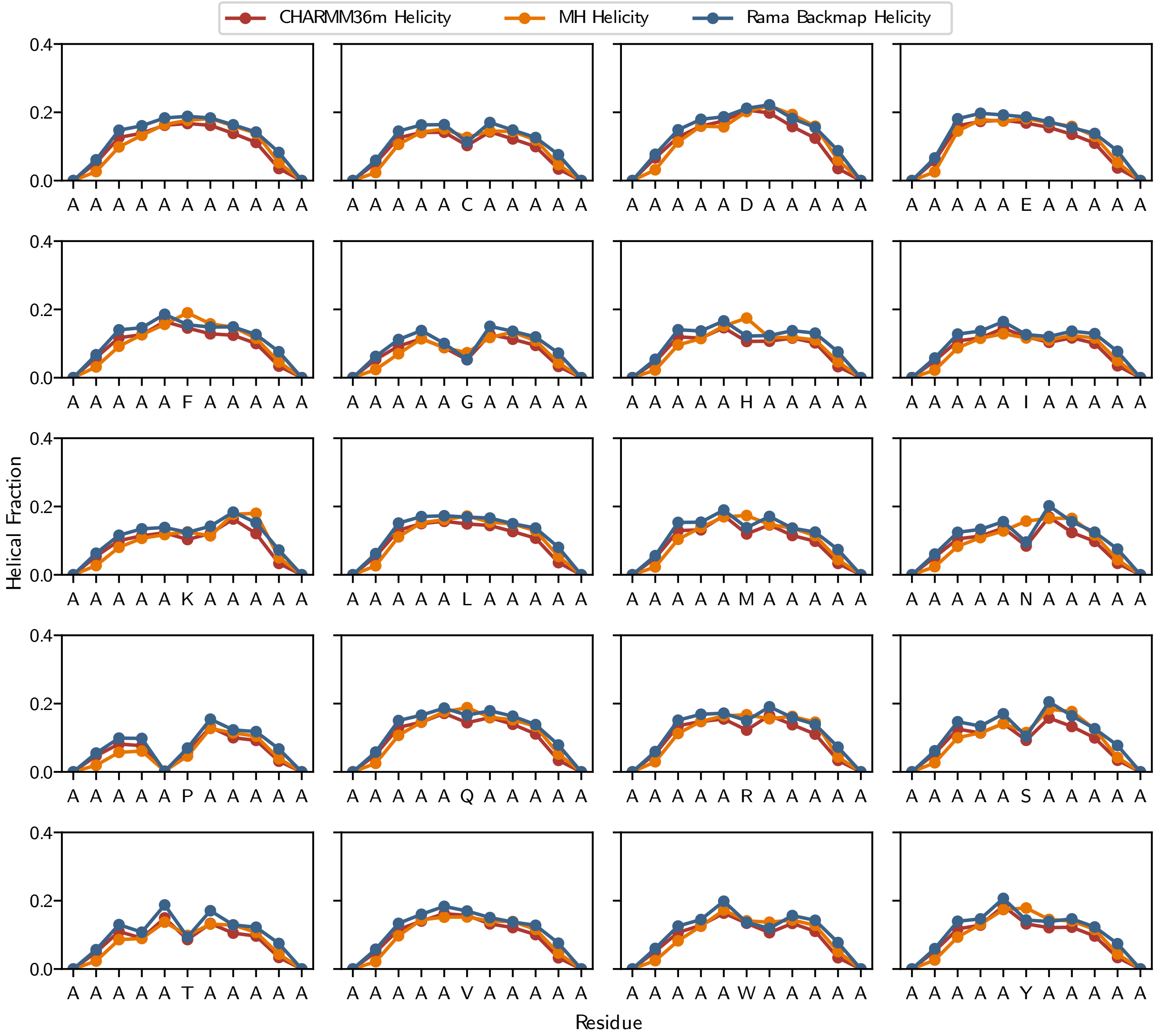


SI Fig. 28. **Helix–coil classifications on $A_5XA_5$ peptides in the CHARMM36m force field.** The atomistic ground truth helical fractions in the CHARMM36m force field for the $A_5XA_5$ peptides are shown in red. In orange and blue are the MH classification and cg2all backmapped classification methods, respectively.

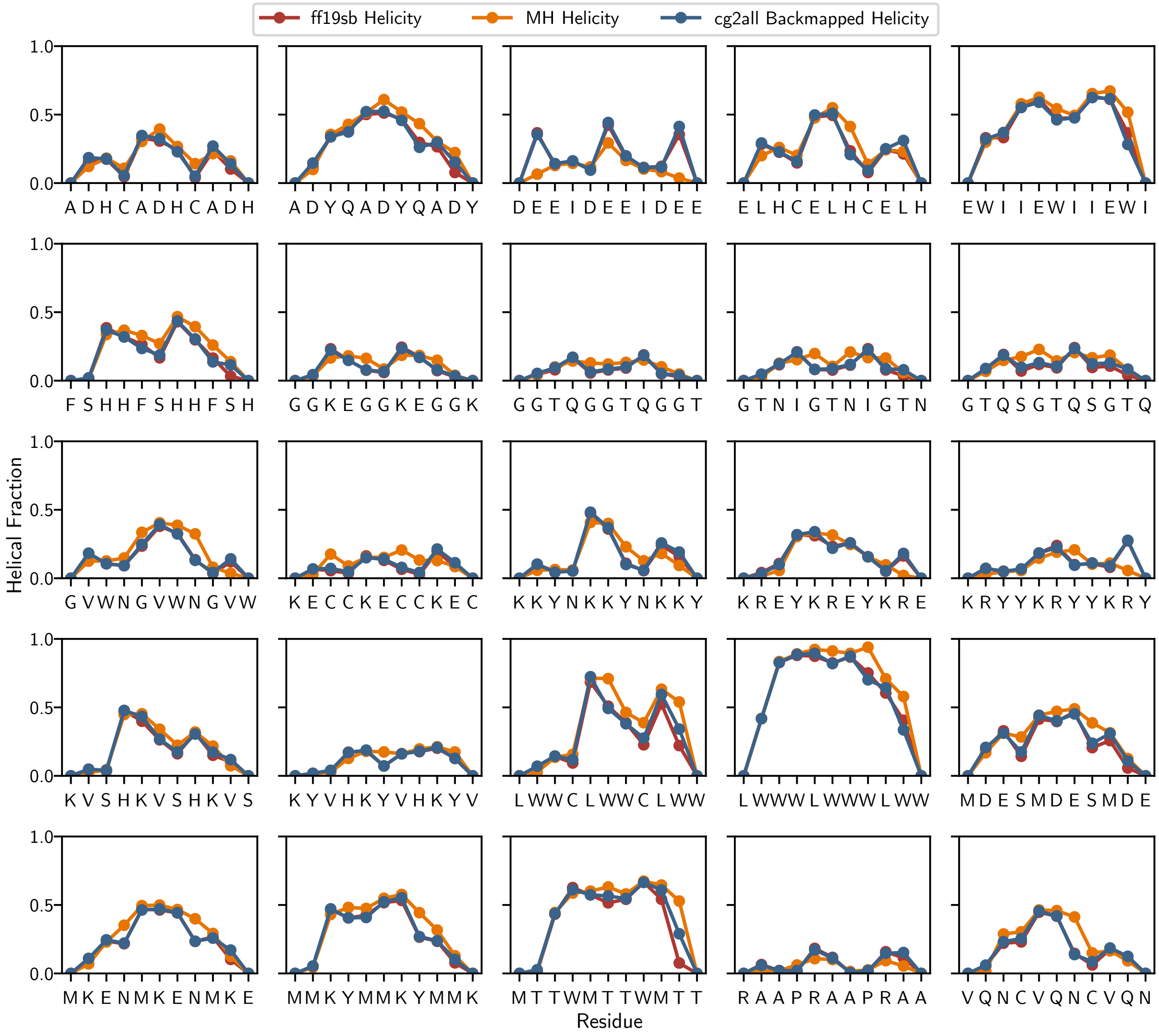


SI Fig. 29. **Helix–coil classifications on the REST2 mean peptides in the ff19sb force field.** The atomistic ground truth helical fractions in the ff19sb force field for the REST2 mean peptides are shown in red. In orange and blue are the MH classification and cg2all backmapped classification methods, respectively.

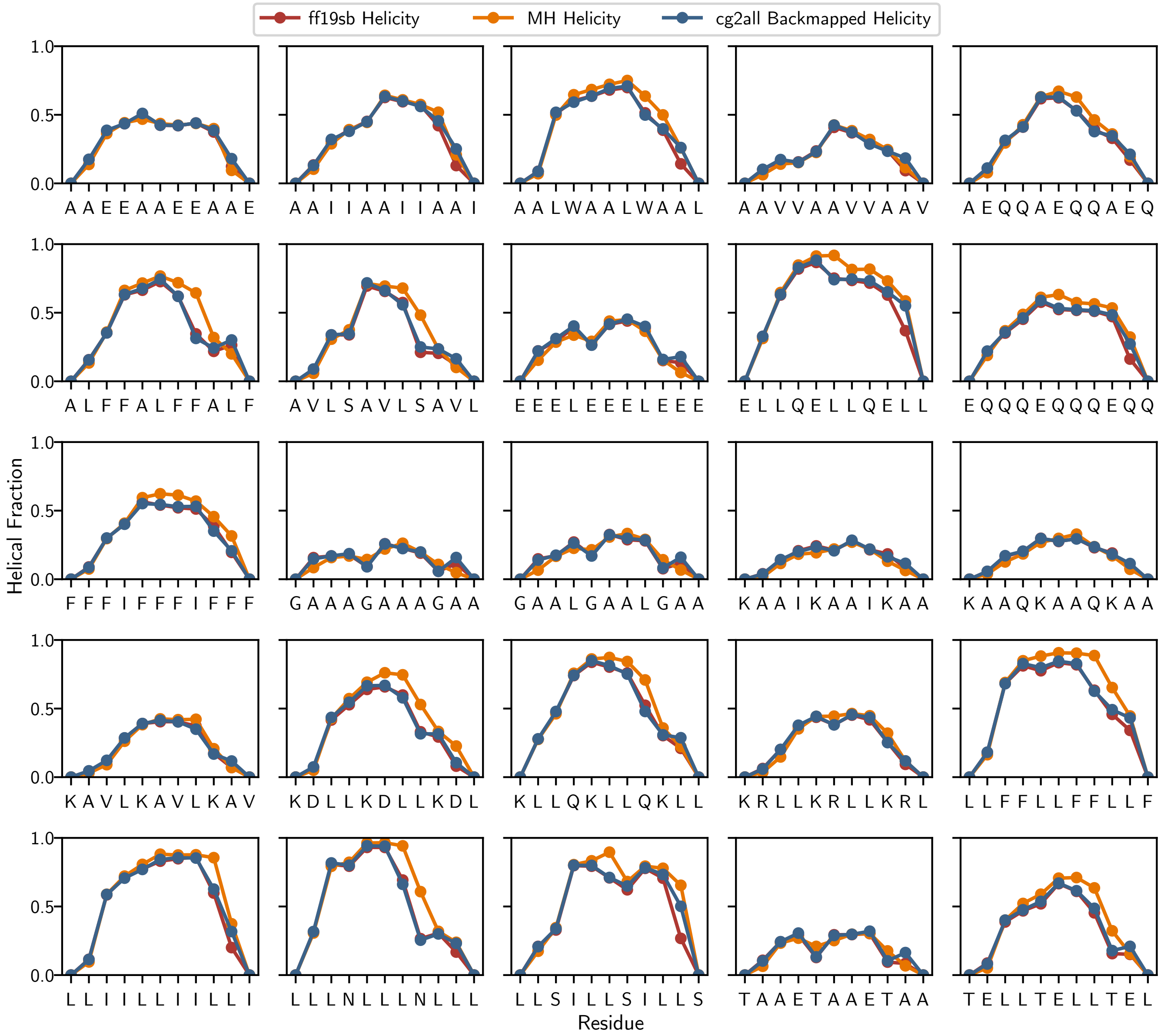


SI Fig. 30. **Helix–coil classifications on the REST2 minus two standard deviations peptides in the ff19sb force field.** The atomistic ground truth helical fractions in the ff19sb force field for the REST2 minus two standard deviations peptides are shown in red. In orange and blue are the MH classification and cg2all backmapped classification methods, respectively.

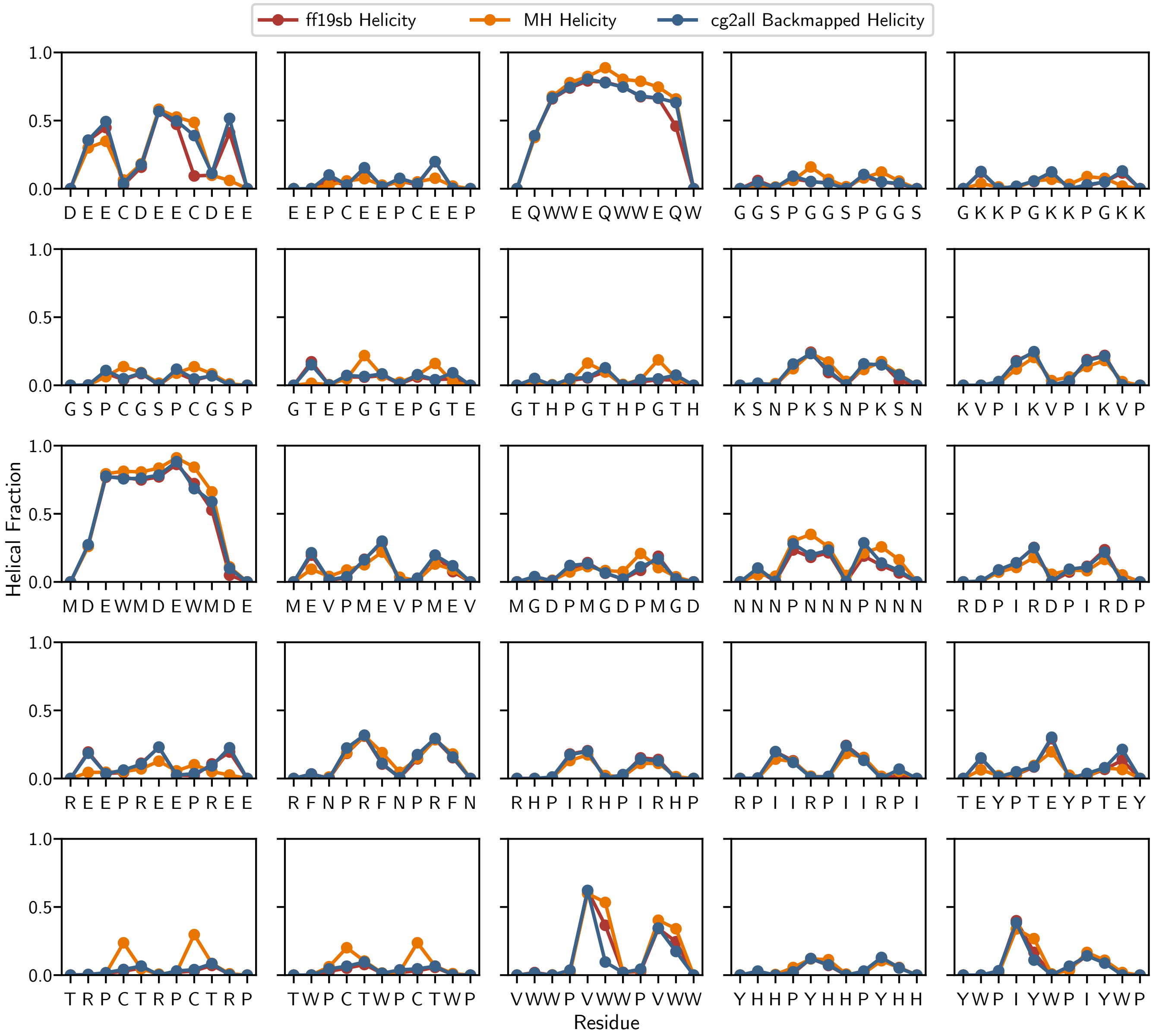


SI Fig. 31. **Helix–coil classifications on the REST2 plus two standard deviations peptides in the ff19sb force field.** The atomistic ground truth helical fractions in the ff19sb force field for the REST2 plus two standard deviations peptides are shown in red. In orange and blue are the MH classification and cg2all backmapped classification methods, respectively.

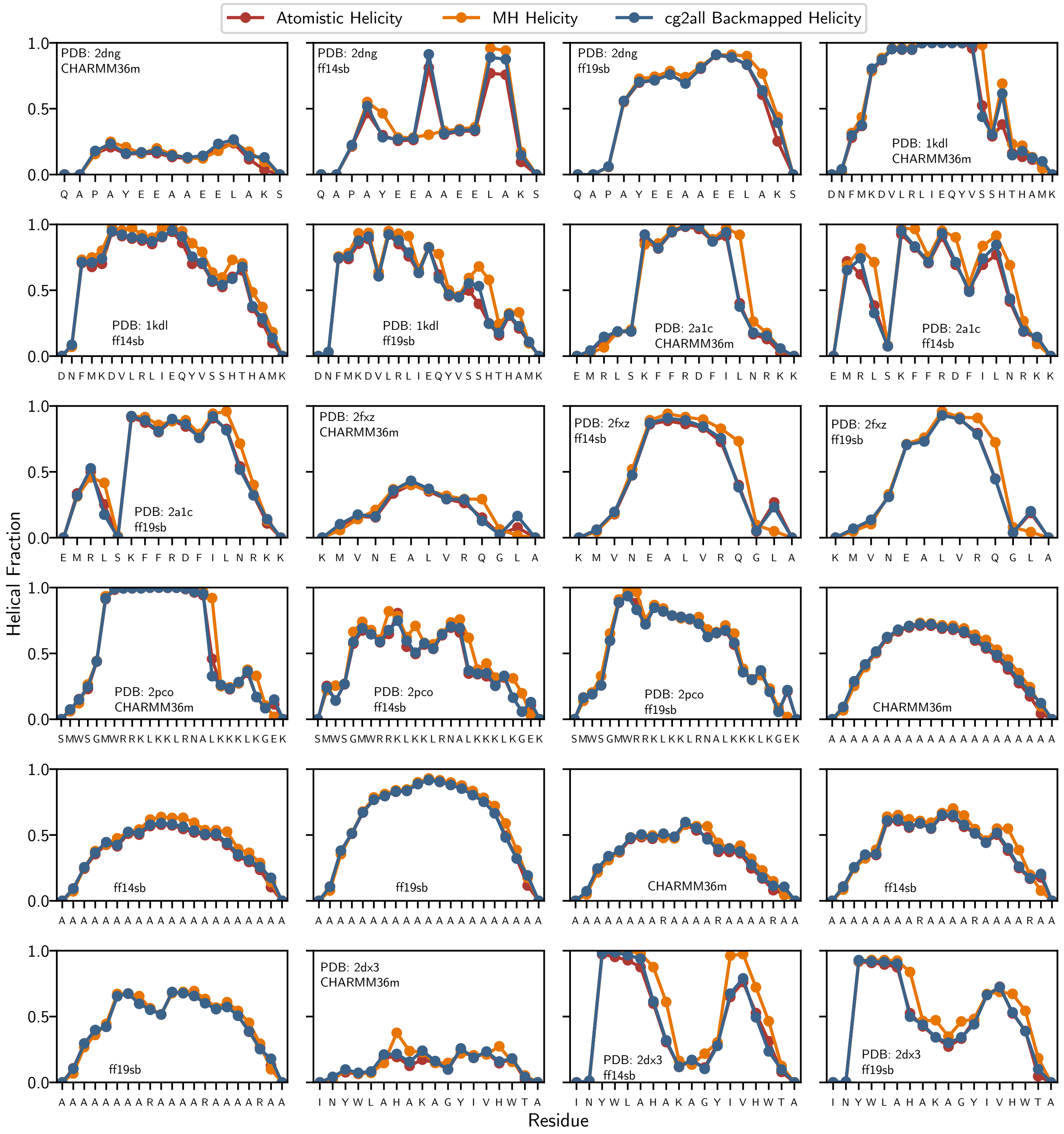


SI Fig. 32. **Helix–coil classifications on PDB and alanine-based peptides in the ff19sb, ff14sb, and CHARMM36m force fields.** The atomistic ground truth helical fractions in the ff19sb, ff14sb, and CHARMM36m force fields for PDB and alanine-based peptides are shown in red. In orange and blue are the MH classification and cg2all backmapped classification methods, respectively.

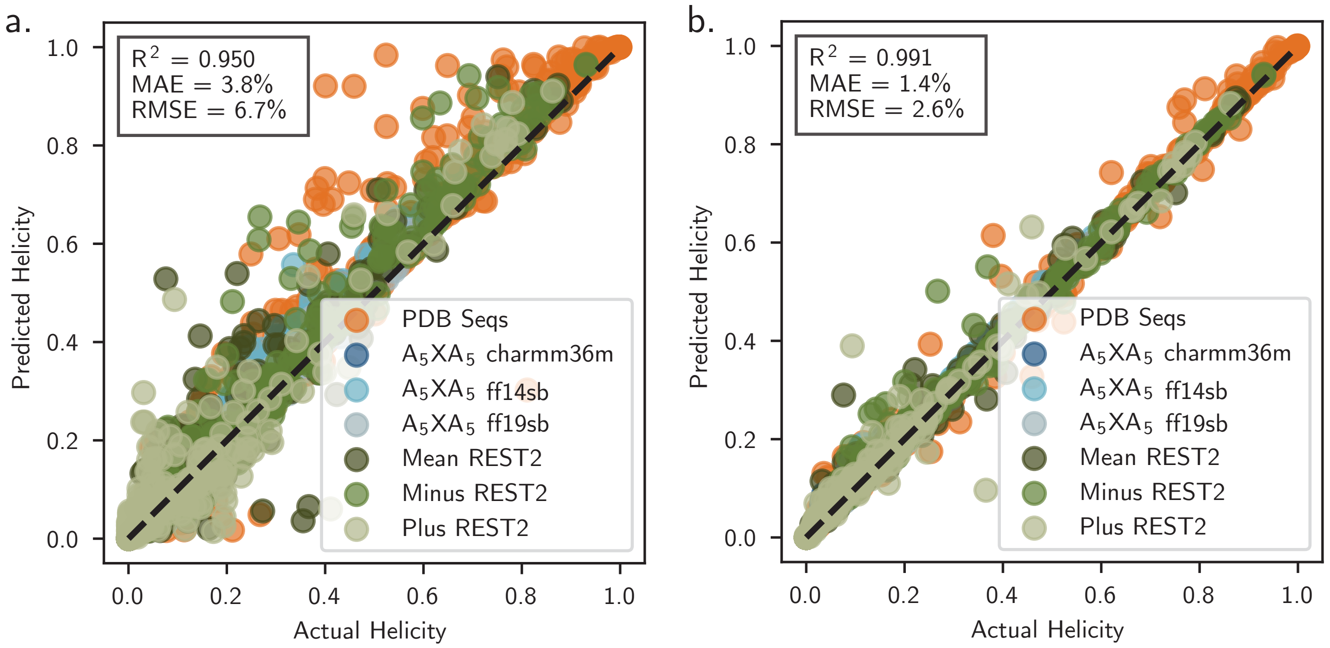


SI Fig. 33. **Parity plots for ground truth helical fractions versus predicted helical fractions.** **a,** The helical fraction for the MH classification method versus the actual atomistic helical fraction for all seven datasets used in validation. **b,** The helical fraction for the cg2all backmapped classification method versus the actual atomistic helical fraction for all seven datasets used in validation.

##### H. Dataset From Experimental NMR Measurements

Moreau et al. used NMR measurements to obtain helical propensities for every canonical amino acid in alanine-based peptides of the form $WK_6L_3A_9XA_9L_3K_6NH2$ [18]. This dataset has been used extensively for parameterizing and validating molecular force fields [1, 10, 38], providing a strong precedent for using its 25°C dataset in our GP parameterization workflow. However, we did not simulate the exact peptide sequences that Moreau et al. studied and instead used peptides of the form $A_5XA_5$. In particular, we removed the lysine caps as peptide solubility and aggregation was not a concern in dilute peptide simulations. We removed the tryptophan residue as we do not need a chromophore for circular dichroism measurements. Finally, we also removed the leucine caps and several alanine residues, reasoning that as long as the alanine end caps in the peptide are long enough, the impact of any distant leucine or alanine residues would be minimal and may be removed for computational efficiency and ease of parameterization. Further, two difficulties that led Moreau et al. to add this large number of leucines and alanines—namely that it is challenging to experimentally resolve helical propensity in peptides with low average helical fractions and the necessity of resolving both helix-formers and breakers across a range of temperatures—are not issues here. In simulations, any helical content can be measured with high accuracy given a long enough simulation trajectory.

As a further justification for our choice of peptide constructs, we note that the ff19sb force field validated the helical propensity of short peptides against Moreau et al.'s dataset by using peptides of the form $Ac - A_4XA_4 - NH_2$ [1]. The authors found that they were able to match the trends in helical propensity from Moreau et al.'s data accurately despite the shorter peptide constructs [1]. Since we do not have explicit capping groups in Mpipi-Helix, we simulated $A_5XA_5$ and similarly expected to reproduce the helical propensities found in Moreau et al. Additionally, it is also noteworthy that several other studies have parameterized or validated force fields against the Moreau et al.'s dataset with no consensus approach for how to mimic Moreau et al.'s peptide sequence computationally. For example, Best et al. compared the helical propensities of the peptide construct $Ac - (A_2XA_2)_3 - NH_2$ [38] to Moreau et al.'s data for atomistic force field parameterization. Additionally, Rizuan et al. [10] used the constructs $A_{20}XA_{20}$ and $A_{20}X_4A_{20}$ to parameterize a coarse-grained force field for describing $\alpha$-helices using Moreau et al.'s data. The discrepancies between the protein sequence used in all of these studies suggests that as long as the sequence contains a significant proportion of alanine residues, Moreau et al.'s data can be successfully used for force field parameterization. For a full list of the protein sequences used, see Table I.

TABLE I. A list of the peptide sequences and abbreviations for the Moreau et al. [18] dataset.

| Full Peptide Sequence | Abbreviation |
|---|---|
| AAAAAAAAAAA | A |
| AAAAACAAAAA | C |
| AAAAADAAAAA | D |
| AAAAAEAAAAA | E |
| AAAAAFAAAAA | F |
| AAAAAGAAAAA | G |
| AAAAAHAAAAA | H |
| AAAAAIAAAAA | I |
| AAAAAKAAAAA | K |
| AAAAALAAAAA | L |
| AAAAAMAAAAA | M |
| AAAAANAAAAA | N |
| AAAAAPAAAAA | P |
| AAAAAQAAAAA | Q |
| AAAAARAAAAA | R |
| AAAAASAAAAA | S |
| AAAAATAAAAA | T |
| AAAAAVAAAAA | V |
| AAAAAWAAAAA | W |
| AAAAAYAAAAA | Y |

Moreau et al. provides the helical propensities of the guest-host peptides in the Lifson–Roig (LR) framework, a thermodynamic framework explaining helix–coil transitions. The helix–coil transition is a structural transition between $\alpha$-helical and coil (disordered) states that has been studied for well over half a century (see LR theory [39] and Zimm-Bragg theory [40]). Thus, in order to use Moreau et al.'s data [18] in the GP training workflow, we needed to generate Lifson–Roig helix–coil thermodynamic parameters from our coarse-grained simulations.

In the LR framework, residues in a peptide are assigned 'H' or 'C' according to whether the backbone $\phi$ and $\psi$ angles of

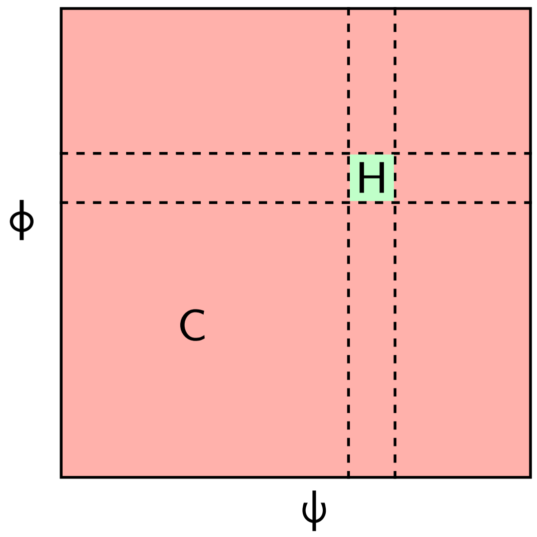


SI Fig. 34. **Helix and coil states as determined by peptide backbone angles.** Under the LR framework, it is necessary to determine which residues in a given simulation snapshot are in a helical ('H') or coil ('C') configuration, which is commonly done by utilizing the protein backbone angles ($\phi$ and $\psi$). In the GP parameterization workflow of Mpipi-Helix, this was done by using the MH method (see 'Helix–Coil Classification in the GP Workflow' section).

the residue are in a helical configuration (SI Fig. 34). From this assignment, each residue in the peptide chain obtains a statistical weight of $u = 1$ for 'C' residues, $w$ for residues in the 'H' state with 'H' residues on either side of them (indicative of a full hydrogen bond), or $v$ for all other types of 'H' assignments. A primary convenience of the LR framework is that the partition function may be solved directly from a matrix equation using these statistical weights (Equations 25 and 26) under the assumption that there are minimal non-local contacts within the peptide.

$$Z = (0\ 0\ 1) \prod_{i=1}^{N} \mathbf{M}_i \begin{pmatrix} 0 \\ 1 \\ 1 \end{pmatrix}. \tag{25}$$

$$\mathbf{M}_i = \begin{pmatrix} w_i & v_i & 0 \\ 0 & 0 & 1 \\ v_i & v_i & 1 \end{pmatrix}. \tag{26}$$

Importantly, Best et al. [31] developed a Bayesian approach using Markov Chain Monte Carlo walks in log-likelihood to extract the parameters $w$ and $v$ from a series of snapshots of a peptide (i.e., a simulation trajectory) where those snapshots have individually classified peptide residues as helical or coil. Details of the approach are provided in Equations 27, 28, and 29. Taking steps in log-likelihood to maximize the probability that the LR parameters explain the observed data allows us to obtain the LR thermodynamic parameters for a given simulation. For this study, we developed a custom python script to employ this technique and acquire the LR values from any simulation that has classified helix and coil frames.

$$p(\text{parameters}|\text{data}) \propto p(\text{data}|\text{parameters})p(\text{parameters}). \tag{27}$$

$$L = p(\text{data}|\text{parameters}) = \prod_{k=1}^{N_k} \rho_k = \prod_{k=1}^{N_k} \frac{1}{Z} \prod_{i=1}^{N} x_{i,k}. \tag{28}$$

$$\ln(L) = \sum_i N_{w,i} \ln(w_i) + \sum_i N_{v,i} \ln(v_i) - N_k \ln(Z). \tag{29}$$

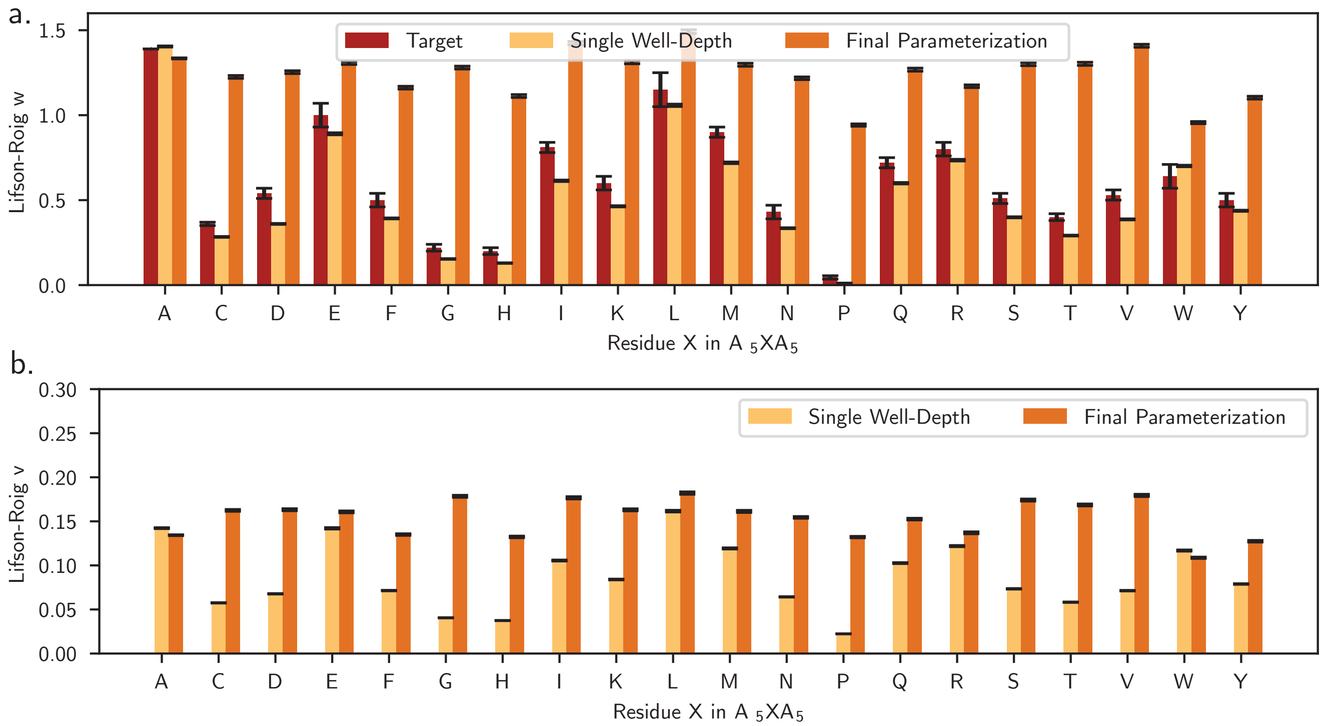


SI Fig. 35. **The LR values for the single well depth results, final parameterization, and target values from Moreau et. al. a,** The $w$ values in the LR framework for each $A_5XA_5$ peptide from the single well depth (i.e., the best well depth from the GP optimization), Mpipi-Helix final parameterization, and target values from Moreau et al. **b,** The $v$ values in the LR framework for each $A_5XA_5$ peptide from the single well depth and Mpipi-Helix final parameterization.

As described above, for Mpipi-Helix's parameterization we simulated $A_5XA_5$ peptides. For GP training, we utilized a loss function of the form Loss $= -(w_{\text{x,Mpipi-Helix}} - w_{\text{x,Moreau}})^2$. Our procedure for determining the $w_{\text{x,Mpipi-Helix}}$ values followed exactly from Best et al. [31] as is described mathematically above. Further details include that we first parameterized to match $w_{\text{A,Mpipi-Helix}}$ based on the entire length of an $A_{11}$ peptide in Mpipi-Helix being represented by a single helical well depth. Then, with $w_{\text{A,Mpipi-Helix}}, v_{\text{A,Mpipi-Helix}}$ and the helical potential describing the protein segments made of all alanine residues fixed, we determined $w_{\text{x,Mpipi-Helix}}$ for every other guest residue. However, to limit the noise in our measurements, we took the middle three residues of the 11 length sequence (i.e., the central AXA) to be solved for as the $w_{\text{x,Mpipi-Helix}}$. This assumption is justified by noting that the helical potential applied to the simulated peptide surrounding the guest residue has significant overlap with the directly neighboring alanine residues. We found that this significantly reduces noise in resolving $w_{\text{x,Mpipi-Helix}}$, which was present due to only having a single point mutation in our protein sequence (which may be why peptides of the form $\text{Ac} - (A_2XA_2)_3 - \text{NH}_2$ were used in Ref. [31, 38] to match Moreau et al.'s data). Finally, in the Bayesian optimization approach, a uniform prior on the parameters was assumed. In the iterative Monte-Carlo steps, an equilibration period of 10,000 moves was performed followed by 100,000 moves in a sampling phase where $w$ and $v$ values were saved every 10 moves. Steps were taken in log-likelihood space with a random value chosen between $-0.05$ and 0.05 accepted based on the Metropolis criterion.

The results of the parameterization are provided in SI Figs. 35 and 36. SI Fig. 35a shows the final $w$ values obtained from the single well depth (i.e., the best well depth from the GP optimization) and the final Mpipi-Helix parameterization applied to the whole peptide compared to the target $w$ value. Figure 35b shows the final $v$ values obtained from the single well depth and the final Mpipi-Helix parameterization. These results were calculated from 300 2.5 µs Mpipi-Helix simulations under each set of parameters for each peptide at 298.15 K in the $NVT$ ensemble with a 100 ps damping constant and 10 fs timestep. Frames were recorded every nanosecond. Additionally, we also provide the residue specific helical fractions for the optimal well depth from the parameterization and from the final Mpipi-Helix parameterization in SI Fig. 36.

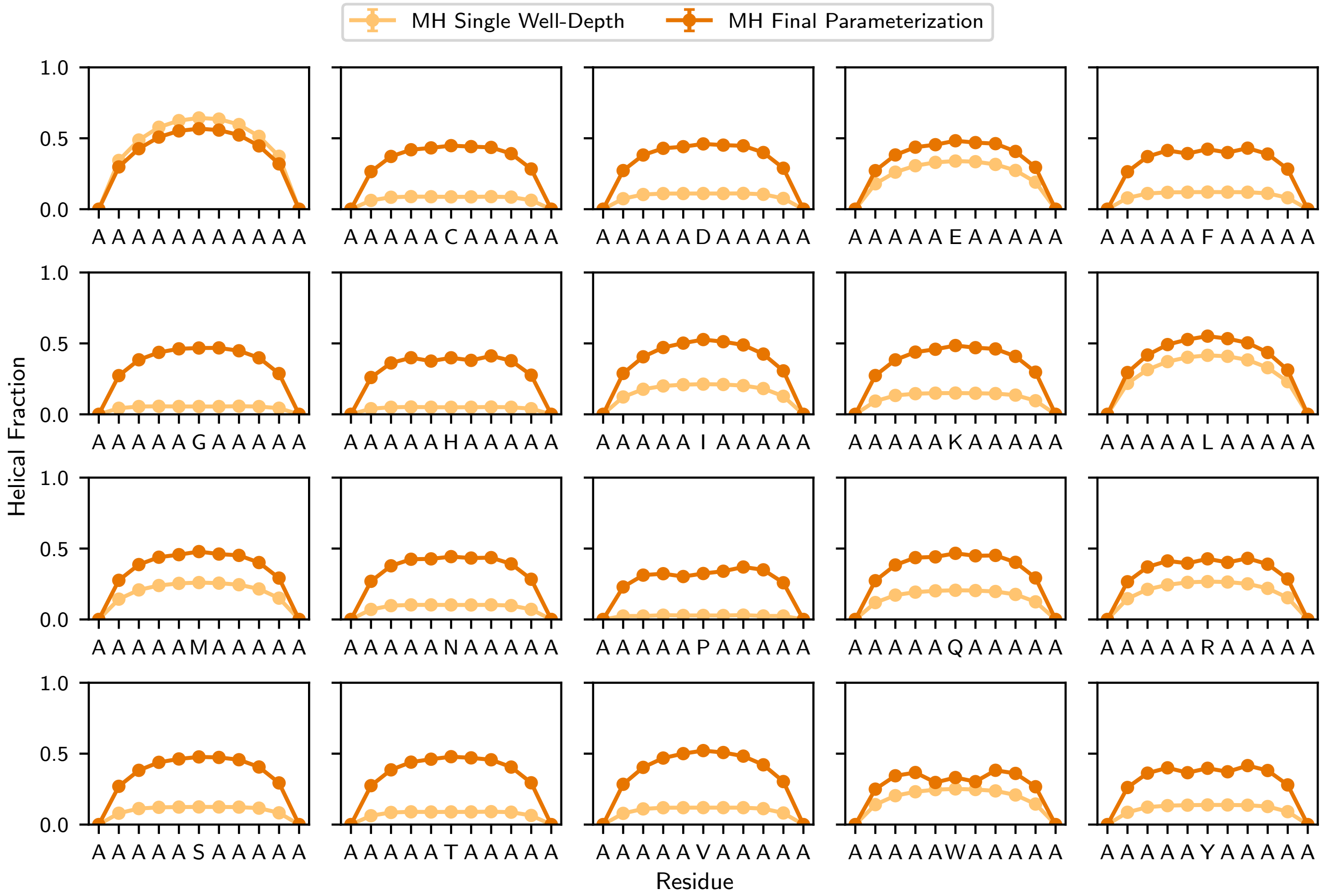


SI Fig. 36. **Helical fraction of $A_5XA_5$ peptides from single well depth values (e.g., the best well depth from the GP optimization) and Mpipi-Helix final parameterization.**

#### I. Dataset from REST2 Enhanced Sampling Atomistic Simulations

The atomistic REST2 simulations of 11 length peptides act as a counterpart to the experimental NMR dataset from Moreau et. al. [18] in our parameterization. In order to minimize the severity of the approximation that a single well depth value could adequately capture the helical propensities of these atomistically simulated peptides in the GP training workflow, we required all peptides to be formed from a repeating ABCDABCDABC pattern (i.e., four different amino acids appearing periodically in the 11 length sequence). From all possible peptides that we could build that adhere to this format (8,855), our goal was to choose a set that spanned a large range of residue sizes and hydrophobicities, as both have been shown to be important in determining residue specific helical propensities in short peptides [17, 41]. To do this we took the unique combinations of four amino acids and measured their average sequence size as determined by the $\sigma$ parameter in Mpipi-Helix, as well as their hydrophobicity as measured by the Urry hydrophobicity scale [42]. In particular, we chose the Urry hydrophobicity scale as it has been demonstrated to be an accurate model for predicting protein phase separation [43]. After we created all 8,855 possible sequences and plotted them on a min-max scaled average size and average hydrophobicity plot, we used scikit-learn's implementation of K-means clustering [20] to create 25 distinct clusters of sequences. From each, we calculated the average Potts score of the full 11 length sequence and fit a Gaussian to the histogram of Potts scores for each cluster. From this Gaussian fit, we took the sequence that was closest to the mean, mean plus two standard deviations, and mean minus two standard deviations from each cluster in order to build a diverse range of sequences predicted by the Potts model to span a large range of helical propensities. A representative histogram with a Gaussian fit and mean, minus two standard deviations, and plus two standard deviations is shown in SI Fig. 37a. The clustered data points and the locations of the chosen mean, plus two standard deviations, and minus two standard deviations points are shown in SI Fig. 37b. Additionally, Tables II, III, and IV show all the selected sequences.

TABLE II. A list of the peptide sequences and abbreviations for the ff19sb REST2 enhanced sampling mean dataset. The converged column indicates whether the simulation was satisfactorily converged. If it was not converged, the sequence was not used in parameterization. Additionally, if the sequence has an $i$ to $i+3$ tryptophan residue (see later in this section) or if it is considered a helix-breaker sequence (see 'GP Training' section), they are annotated with 'Not Used'.

| Full Peptide Sequence | Abbreviation | Converged |
|---|---|---|
| ADHCADHCADH | ADHC | Yes |
| ADYWADYWADY | ADYW | Yes |
| DEEIDEEIDEE | DEEI | No |
| ELHCELHCELH | ELHC | Yes |
| EWIIEWIIEWI | EWII | Yes |
| FSHHFSHHFSH | FSHH | Yes |
| GGKEGGKEGGK | GGKE | Yes |
| GGTQGGTQGGT | GGTQ | Yes |
| GTNIGTNIGTN | GTNI | Yes |
| GTQSGTQSGTQ | GTQS | Yes |
| GVWNGVWNGVW | GVWN | Yes |
| KECCKECCKEC | KECC | Yes |
| KKYNKKYNKKY | KKYN | Yes |
| KREYKREYKRE | KREY | No |
| KRYYKRYYKRY | KRYY | Yes |
| KVSHKVSHKVS | KVSH | Yes |
| KYVHKYVHKYV | KYVH | Yes |
| LWWCLWWCLWW | LWWC | Yes - Not Used |
| LWWWLWWWLWW | LWWW | Yes - Not Used |
| MDESMDESMDE | MDES | No |
| MKENMKENMKE | MKEN | Yes |
| MMKYMMKYMMK | MMKY | Yes |
| MTTWMTTWMTT | MTTW | Yes |
| RAAPRAAPRAA | RAAP | Yes |
| VQNCVQNCVQN | VQNC | Yes |

Once our sequences were chosen, we simulated them in the ff19sb force field [1], which we chose due to its accuracy in predicting helical propensities and because it was benchmarked against the same experimental NMR dataset we used in our GP training workflow [18]. In order to more adequately sample our peptides and have confidence in our prediction of their helical fractions, we utilized the enhanced sampling technique Replica Exchange Solute Tempering 2 [19]. Inputs for these simulations were generated from CHARMM-GUI [35] to run in the GROMACS software [34] using PDB files generated

TABLE III. A list of the peptide sequences and abbreviations for the ff19sb REST2 enhanced sampling minus two standard deviations dataset. The converged column indicates whether the simulation was satisfactorily converged. If it was not converged, the sequence was not used in parameterization. Additionally, if the sequence has an $i$ to $i+3$ tryptophan residue (see later in this section) or if it is considered a helix-breaker sequence (see 'GP Training' section), they are annotated with 'Not Used'.

| Full Peptide Sequence | Abbreviation | Converged |
|---|---|---|
| AAEEAAEEAAE | AAEE | Yes |
| AAIIAAIIAAI | AAII | Yes |
| AALWAALWAAL | AALW | Yes |
| AAVVAAVVAAV | AAVV | Yes |
| AEQQAEQQAEQ | AEQQ | Yes |
| ALFFALFFALF | ALFF | Yes |
| AVLSAVLSAVL | AVLS | Yes |
| EEELEEELEEE | EEEL | No |
| ELLQELLQELL | ELLQ | Yes |
| EQQQEQQQEQQ | EQQQ | Yes |
| FFFIFFFIFFF | FFFI | Yes |
| GAAAGAAAGAA | GAAA | Yes |
| GAALGAALGAA | GAAL | Yes |
| KAAIKAAIKAA | KAAI | Yes |
| KAAQKAAQKAA | KAAQ | Yes |
| KAVLKAVLKAV | KAVL | Yes |
| KDLLKDLLKDL | KDLL | Yes |
| KLLQKLLQKLL | KLLQ | Yes |
| KRLLKRLLKRL | KRLL | Yes |
| LLFFLLFFLLF | LLFF | Yes |
| LLIILLIILLI | LLII | Yes |
| LLLNLLLNLLL | LLLN | Yes |
| LLSILLSILLS | LLSI | Yes |
| TAAETAAETAA | TAAE | Yes |
| TELLTELLTEL | TELL | Yes |

from PyMol [36] that were initialized in an extended conformation. All peptides were uncapped as the Mpipi-Helix model does not have capping groups. These initial simulation configurations were generated with OPC water in 150 mM of NaCl. Our procedure for each sequence was first to run a brief minimization using steepest descent with a force tolerance of 1000 kJ/mol/nm for 5000 steps. Thereafter, a 1 ns $NVT$ equilibration run was performed with a time step of 1 fs at 300 K using the v-rescale thermostat with a time constant of 1 ps and separate groups on the solvent and solute. Position restraints were maintained on the heavy atoms of the protein and the LINCS algorithm [37] was applied to bonds involving hydrogen atoms. After this step, eight temperature replicas were started at 300.00, 322.71, 347.14, 373.42, 401.69, 432.10, 464.81, and 500.00 K. These were simulated in the $NVT$ ensemble using a timestep of 2 fs and the v-rescale thermostat with a time constant of 1 ps and separate groups for the solvent and solute. A PME grid was used for electrostatics and the cutoffs for short-ranged neighbor lists, van der Waals interactions, and coulomb cutoff was set to 0.9 nm. Sampling was run for 1.55 µs (12.4 aggregate µs), with the first 50 ns used for equilibration of the replicas. Exchanges between adjacent replicas were performed every 5000 steps (10 ps).

We confirmed convergence of the replicas by examining the exchange probabilities, the helical fraction per residue, and the free energy surfaces of each peptide. The exchange probabilities for each peptide are plotted in the three groups (mean, minus two standard deviations, and plus two standard deviations) in SI Figs. 38, 39, and 40. Peptides with exchange probabilities below 0.15 between replicates 1 and 2 were considered as not being converged and excluded from the parameterization scheme. This includes the DEEI, MDES, EEEL, DEEC, and MDEW peptides. Additionally, the peptide KREY was excluded from the parameterization despite its 0.17 exchange probability between replicate 1 and 2. This is because of a long, sustained salt-bridge in the simulation that we believe prevented adequate sampling. Further, we also removed peptides that contained $i$ to $i+3$ tryptophan residues. This is because the Mpipi-Helix model has trouble adequately forming helices in this case as tryptophan is the largest residue in Mpipi which makes ramping difficult (see section 'Implementing Mpipi-Helix in the LAMMPS Software'). We thus believe parameterizing on based on $i$ to $i+3$ tryptophans would be difficult and could throw off our parameterization scheme. Additionally, the occurrence of $i$ to $i+3$ tryptophan residues in the helical sequences we took from the SwissProt database (see 'Potts Model' section) is about an order of magnitude lower than what is expected by chance (at 0.032% versus the expected 0.25% by chance). This indicates it is not a very important sequence to parameterize in Mpipi-Helix.

TABLE IV. A list of the peptide sequences and abbreviations for the ff19sb REST2 enhanced sampling plus two standard deviations dataset. The converged column indicates whether the simulation was satisfactorily converged. If it was not converged, the sequence was not used in parameterization. Additionally, if the sequence has an $i$ to $i+3$ tryptophan residue (see later in this section) or if it is considered a helix-breaker sequence (see 'GP Training' section), they are annotated with 'Not Used'.

| Full Peptide Sequence | Abbreviation | Converged |
|---|---|---|
| DEECDEECDEE | DEEC | No |
| EEPCEEPCEEP | EEPC | Yes |
| EQWWEQWWEQW | EQWW | No |
| GGSPGGSPGGS | GGSP | Yes - Not Used |
| GKKPGKKPGKK | GKKP | Yes |
| GSPCGSPCGSP | GSPC | Yes - Not Used |
| GTEPGTEPGTE | GTEP | Yes |
| GTHPGTHPGTH | GTHP | Yes - Not Used |
| KSNPKSNPKSN | KSNP | Yes |
| KVPIKVPIKVP | KVPI | Yes |
| MDEWMDEWMDE | MDEW | No |
| MEVPMEVPMEV | MEVP | Yes |
| MGDPMGDPMGD | MGDP | Yes |
| NNNPNNNPNNN | NNNP | Yes |
| RDPIRDPIRDP | RDPI | Yes |
| REEPREEPREE | REEP | Yes |
| RFNPRFNPRFN | RFNP | Yes |
| RHPIRHPIRHP | RHPI | Yes |
| RPIIRPIIRPI | RPII | Yes |
| TEYPTEYPTEY | TEYP | Yes |
| TRPCTRPCTRP | TRPC | Yes - Not Used |
| TWPCTWPCTWP | TWPC | Yes - Not Used |
| VWWPVWWPVWW | VWWP | Yes - Not Used |
| YHHPYHHPYHH | YHHP | Yes |
| YWPIYWPIYWP | YWPI | Yes |

The helical fractions per residue for every sequence are displayed in SI Figs. 41, 42, and 43. Proper convergence of the sampled peptides is demonstrated by a smooth ordering by temperature of helical fractions with the coldest replica having the highest helical fractions and the hottest replica having the lowest helical fractions. For peptides that are insufficiently sampled, the helical fractions look much more rugged (see DEEI in SI Fig. 41 and DEEC in SI Fig. 43). The only exception for this cold to hot ordering of helical fractions are for sequences that contain proline. For these sequences, the hottest replica typically has the highest helical content for proline and nearby residues. This is due to proline's helix-breaking character being significantly reduced at higher temperatures [18].

Finally, the free energy surfaces which depict the sampling of the root mean squared deviation to a perfect helix and the radius of gyration as order parameters are provided in a Jupyter notebook within the GitHub repository associated with this manuscript. As with the helical fraction plots, convergence on these plots is indicated by a smooth transition of the free energy minimas away from the helical state and towards the coil state as the replicas increase in temperature.

For the GP parameterization scheme, we constructed a loss function that utilized a smoothed version of the atomistic REST2 sampled peptide's helical fraction. Specifically, we calculated this smoothed version of the helical fractions by taking a 5 residue windowed average of the helical fraction for every residue within the peptide. Then, we calculated the loss as the negative average of $(1 - \frac{\text{MH}_i}{\text{REST2}_i})^2$, where $\text{MH}_i$ is the helical fraction of the $i$th residue in the Mpipi-Helix simulation and $\text{REST2}_i$ is the smoothed helical fraction of the $i$th residue from the atomistic REST2 simulations. The smoothing is necessary as we simulate these peptides in Mpipi-Helix using a single well-depth value that applies to the whole protein sequence. This means that it is impossible to recover all the features of the helical fractions from the atomistic simulations as the helical fraction curve for Mpipi-Helix will always be smooth.

SI Figs. 44, 45, and 46 provide the results for the parameterization of the atomistic REST2 simulations within Mpipi-Helix, plotting the helical fractiosn of replica 1 for every sequence, alongside the smoothed helical fractions, the helical fractions of the best well depth value from the Mpipi-Helix parameterization process, and the helical fractions from the final parameterization of Mpipi-Helix. These results are derived from 300 2.5 µs Mpipi-Helix simulations under each set of parameters for each peptide at 298.15 K in the $NVT$ ensemble with a 100 ps damping constant and 10 fs timestep. Frames were recorded every nanosecond. Additionally, we also provide parity plots which plot the target REST2 helical fractions for every set of four consecutive residues in the atomistic peptides against the Mpipi-Helix single well depth values helical

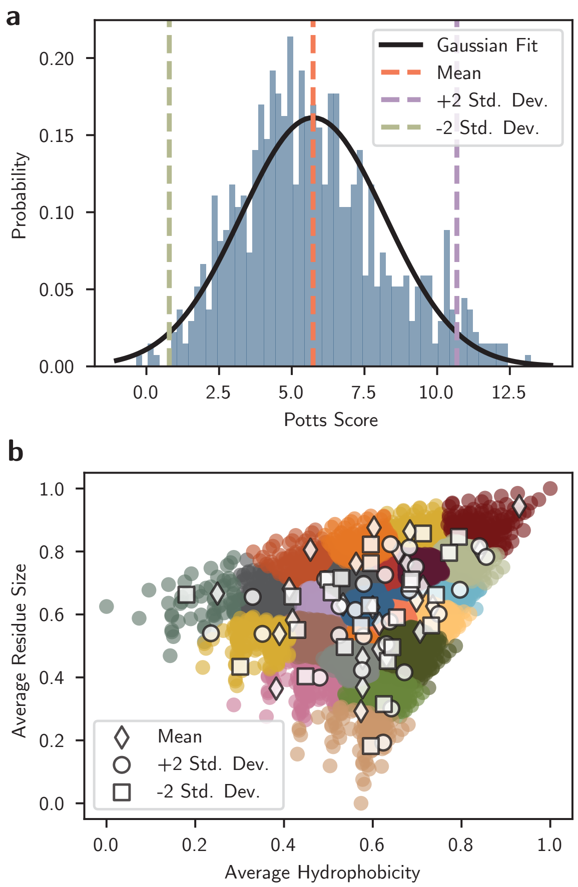


SI Fig. 37. **Sequences were chosen to span average residue size and hydrophobicity for REST2 enhanced sampling simulations to use in the GP parameterization scheme. a,** The histogram for cluster index 1 with a fitted Gaussian. **b,** 8,855 sequences were made as combinations (with replacement) of all possible four length sequences of the 20 canonical amino acids. They were plotted against average residue size ($\sigma$ values in Mpipi) and hydrophobicity (Urry hydrophobicity scale [42]) and their Potts scores were determined for the full 11 length sequence. From each cluster, three sequences were chosen, one that was closest to the mean, mean minus two standard deviations, and mean plus two standard deviations from the Gaussian fit on each cluster.

fraction in SI Fig. 47. The $R^2$ values for these plots ($R^2 = 0.668$ in the mean dataset (a), $R^2 = 0.770$ in the minus dataset (b), and $R^2 = 0.507$ in the plus dataset (c)) indicate that even though the single well depth does not match the target helical fractions for every peptide after parameterization, the trend is still strong and linear. So, while results for certain sequences may be poor due to high helical fractions for large residues being noisy to reproduce in the Mpipi-Helix model (e.g., see ADYW in SI Fig. 44), the overall trends are still adequate for our parameterization. Finally, the Mpipi-Helix final parameterization helical fractions for these peptides tend to differ significantly from the single well depth helical fractions. This is because we used the NMR experimental dataset to get the y-intercept of our Mpipi-Helix well depth trend and due to the shifting of the REST2 datasets when we extracted the slope of our trend.

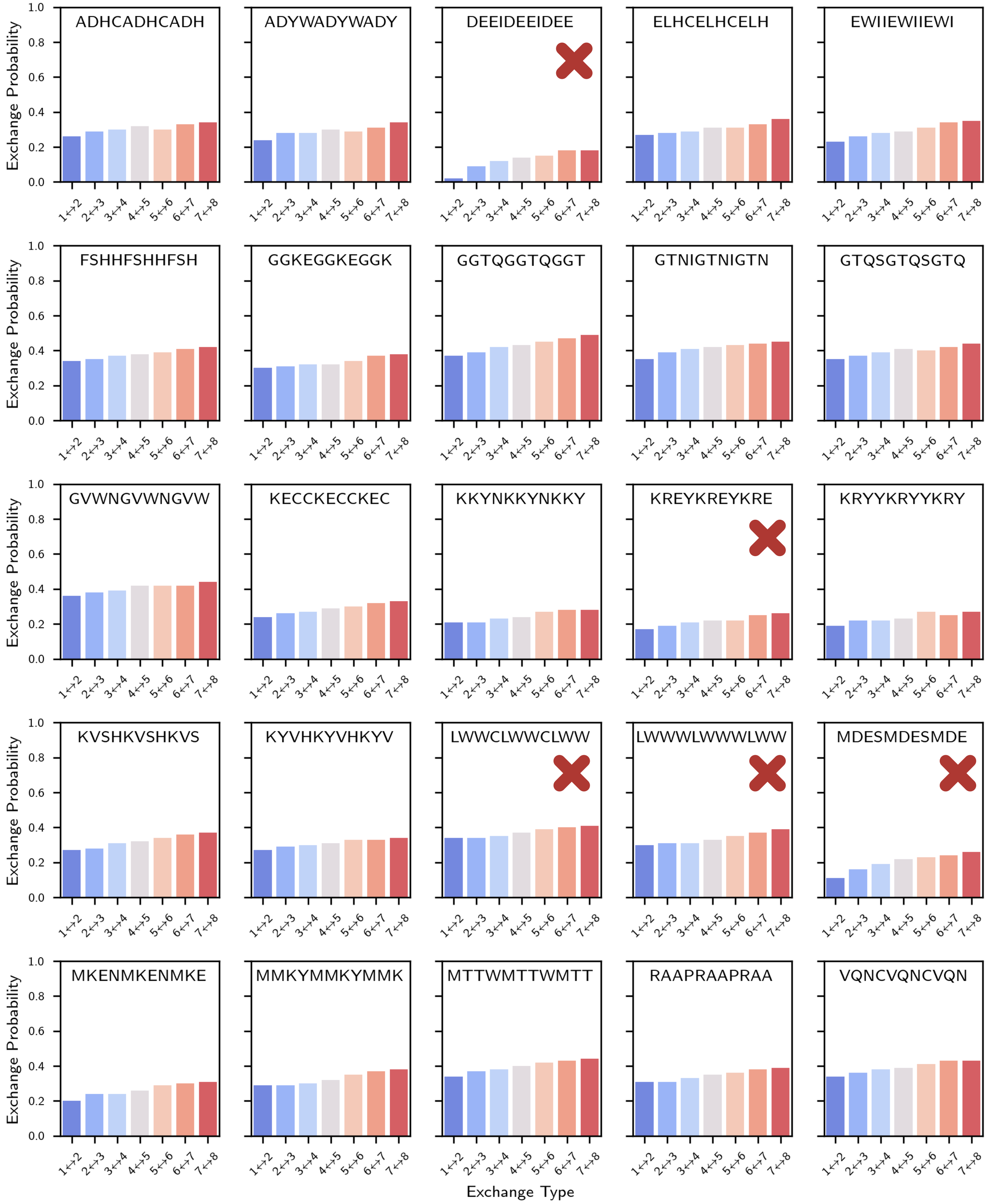


SI Fig. 38. **Exchange probabilities between replicas of mean REST2 sequences.** Red 'X' symbols indicate sequences that were excluded due to insufficient convergence or the presence of $i$ to $i+3$ tryptophan residues.

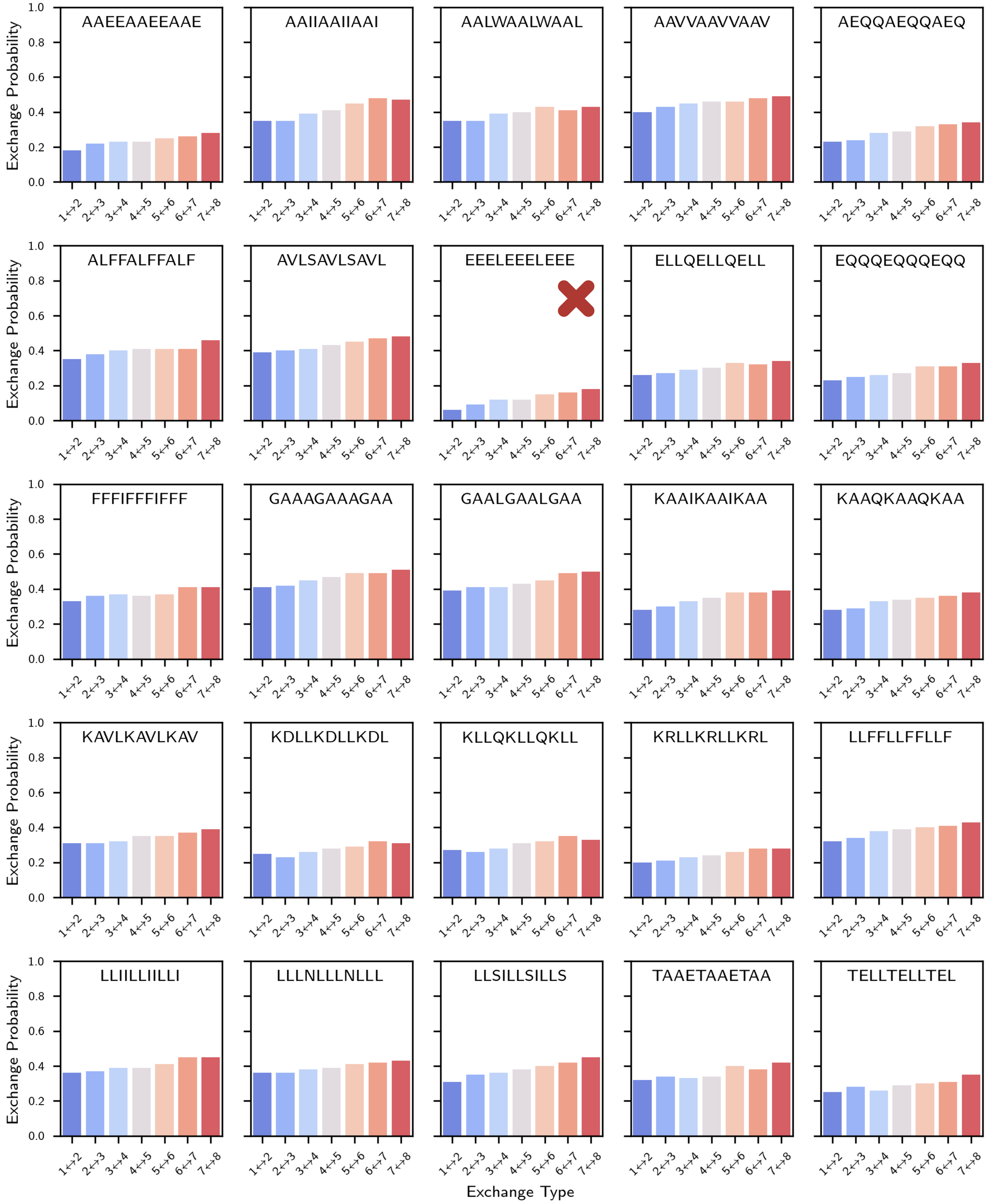


SI Fig. 39. **Exchange probabilities between replicas of minus two standard deviations REST2 sequences.** Red ‘X’ symbols indicate sequences that were excluded due to insufficient convergence or the presence of $i$ to $i+3$ tryptophan residues.

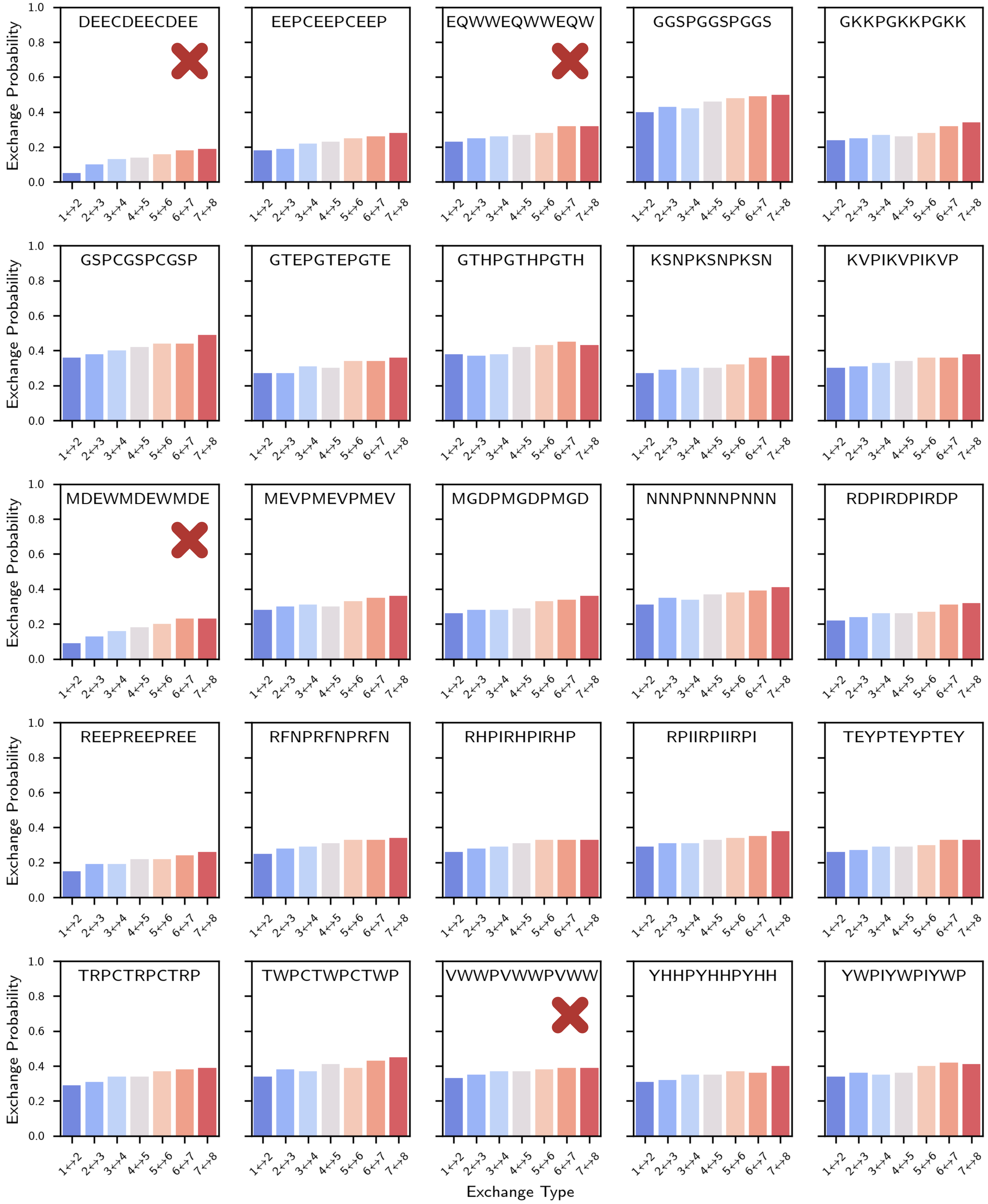


SI Fig. 40. **Exchange probabilities between replicas of plus two standard deviations REST2 sequences.** Red ‘X’ symbols indicate sequences that were excluded due to insufficient convergence or the presence of $i$ to $i+3$ tryptophan residues.

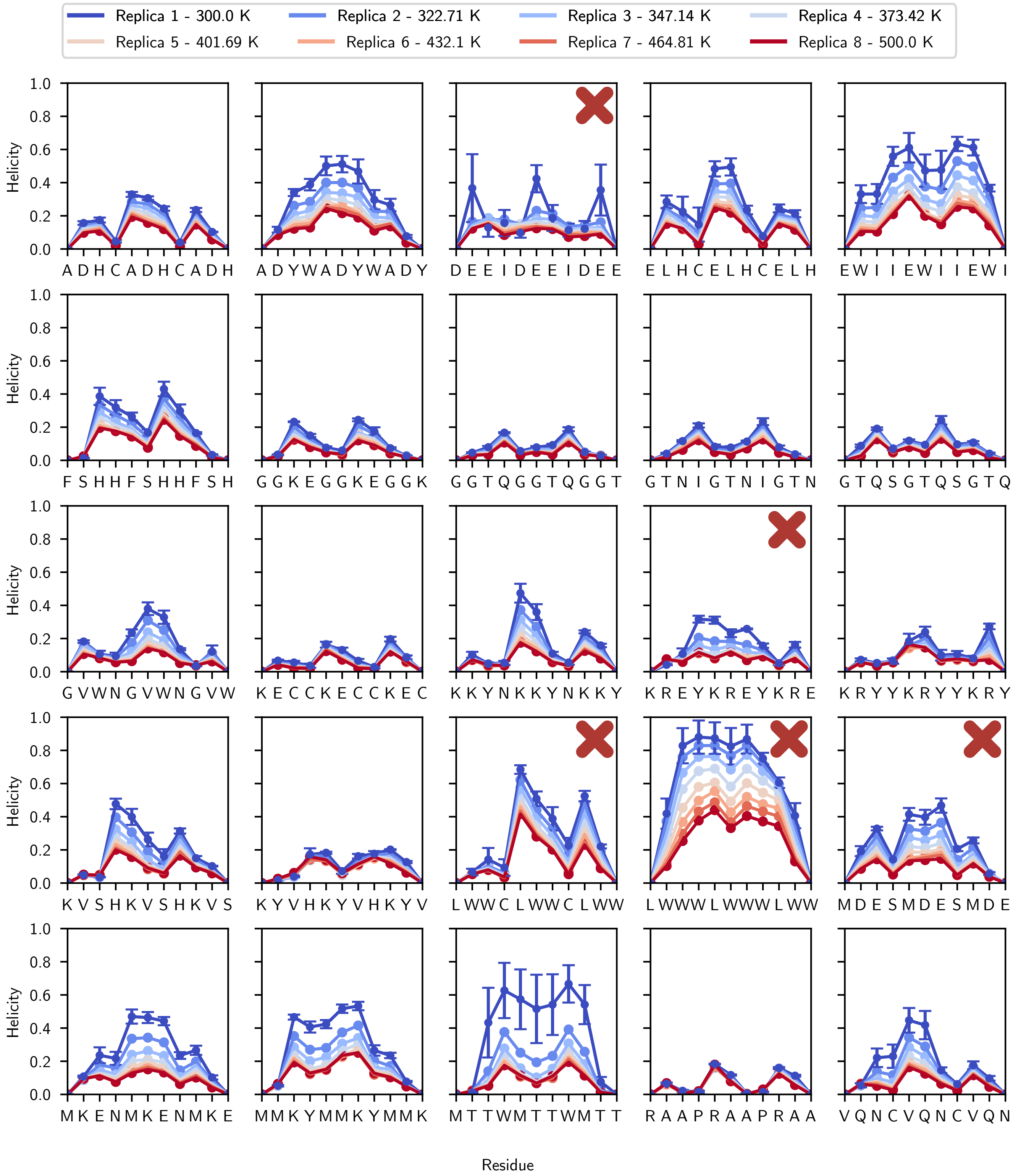


SI Fig. 41. **Residue specific helical fractions for all replicas of mean REST2 sequences.** Red 'X' symbols indicate sequences that were excluded due to insufficient convergence or the presence of $i$ to $i+3$ tryptophan residues.

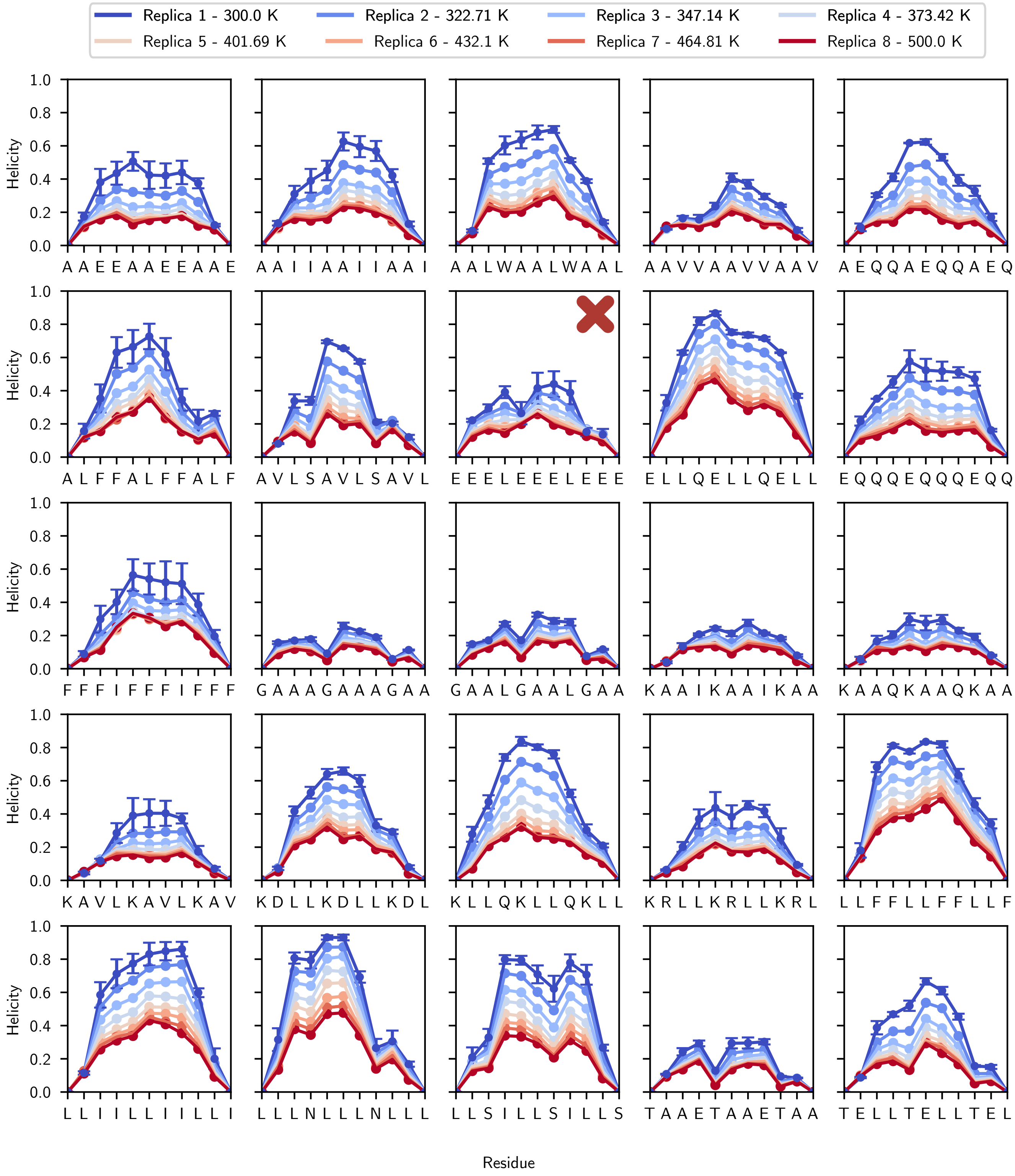


SI Fig. 42. **Residue specific helical fractions for all replicas of minus two standard deviations REST2 sequences.** Red ‘X’ symbols indicate sequences that were excluded due to insufficient convergence or the presence of $i$ to $i+3$ tryptophan residues.

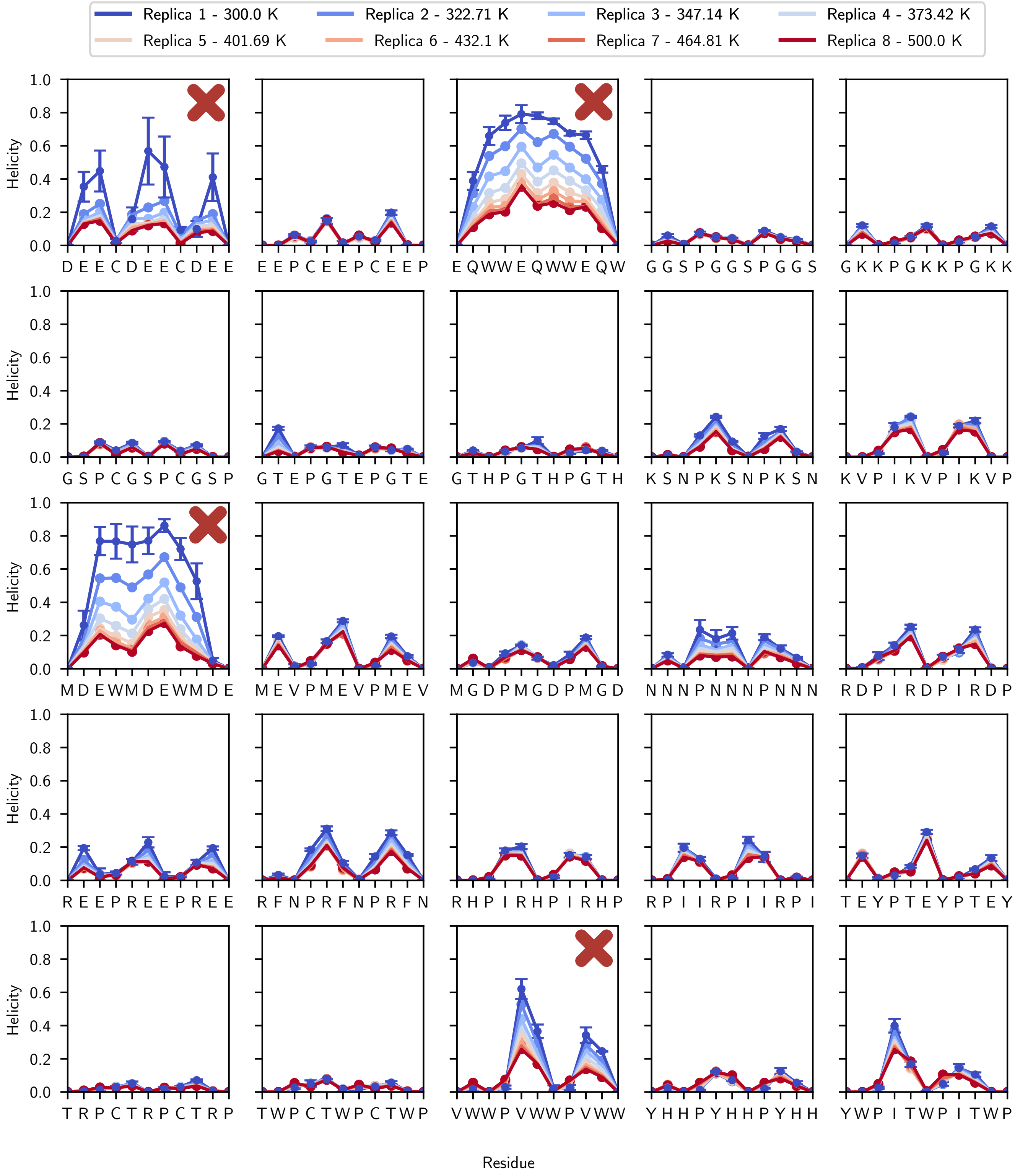


SI Fig. 43. **Residue specific helical fractions for all replicas of plus two standard deviations REST2 sequences.** Red 'X' symbols indicate sequences that were excluded due to insufficient convergence or the presence of $i$ to $i+3$ tryptophan residues.

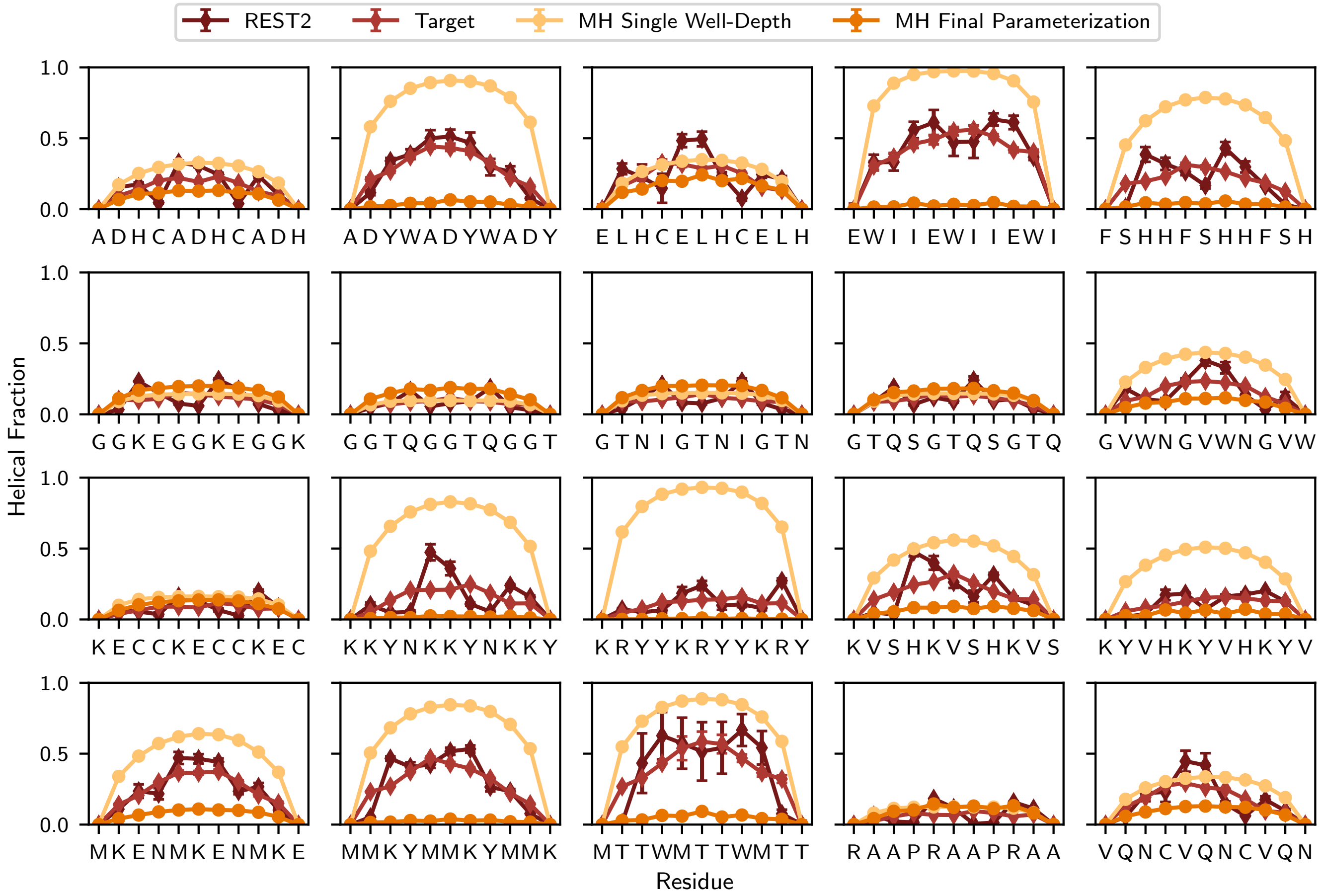


SI Fig. 44. **Parameterization results for the REST2 mean dataset.** The helical fractions for all peptides in the REST2 mean dataset that were included in our parameterization approach. Displayed are the REST2 helical fraction of the first replica taken directly from the atomistic simulation (dark maroon), the smoothed target helical fraction (red), the Mpipi-Helix single well depth helical fraction from the result of our GP parameterization (light yellow), and the result from our final Mpipi-Helix parameterization (orange).

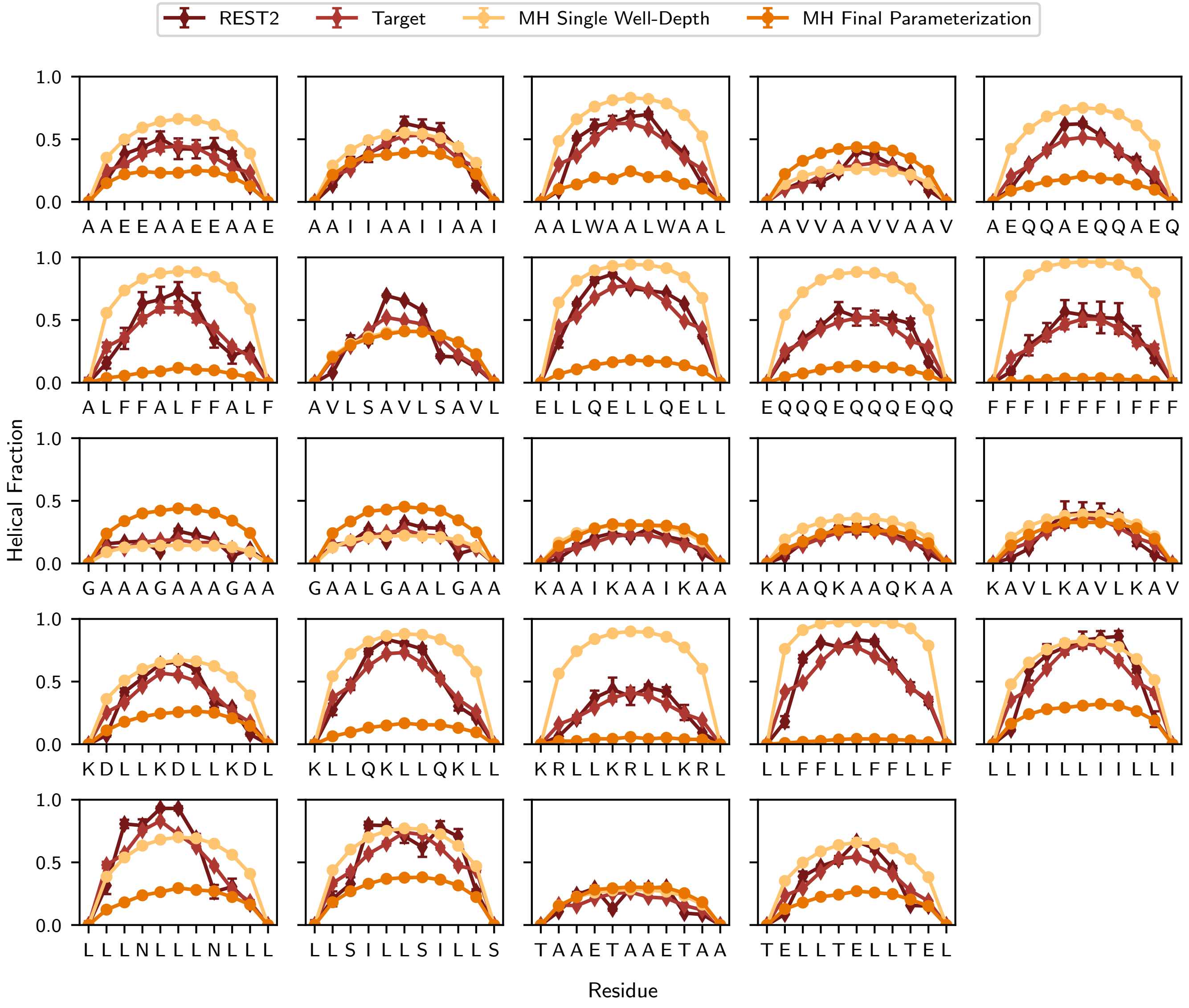


SI Fig. 45. **Parameterization results for the REST2 minus two standard deviations dataset.** The helical fractions for all peptides in the REST2 mean dataset that were included in our parameterization approach. Displayed are the REST2 helical fraction of the first replica taken directly from the atomistic simulation (dark maroon), the smoothed target helical fraction (red), the Mpipi-Helix single well depth helical fraction from the result of our GP parameterization (light yellow), and the result from our final Mpipi-Helix parameterization (orange).

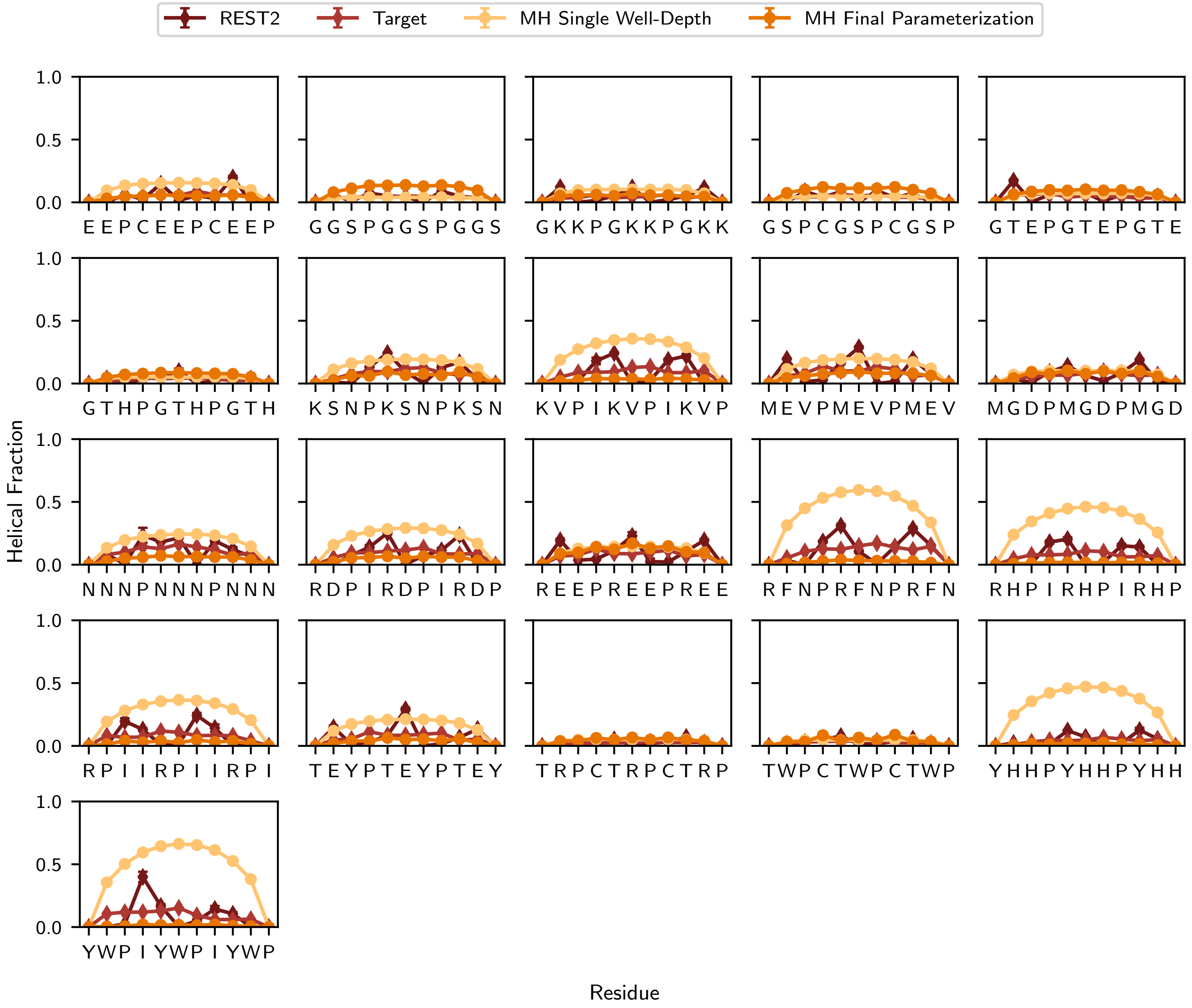


SI Fig. 46. **Parameterization results for the REST2 plus two standard deviations dataset.** The helical fractions for all peptides in the REST2 mean dataset that were included in our parameterization approach. Displayed are the REST2 helical fraction of the first replica taken directly from the atomistic simulation (dark maroon), the smoothed target helical fraction (red), the Mpipi-Helix single well depth helical fraction from the result of our GP parameterization (light yellow), and the result from our final Mpipi-Helix parameterization (orange).

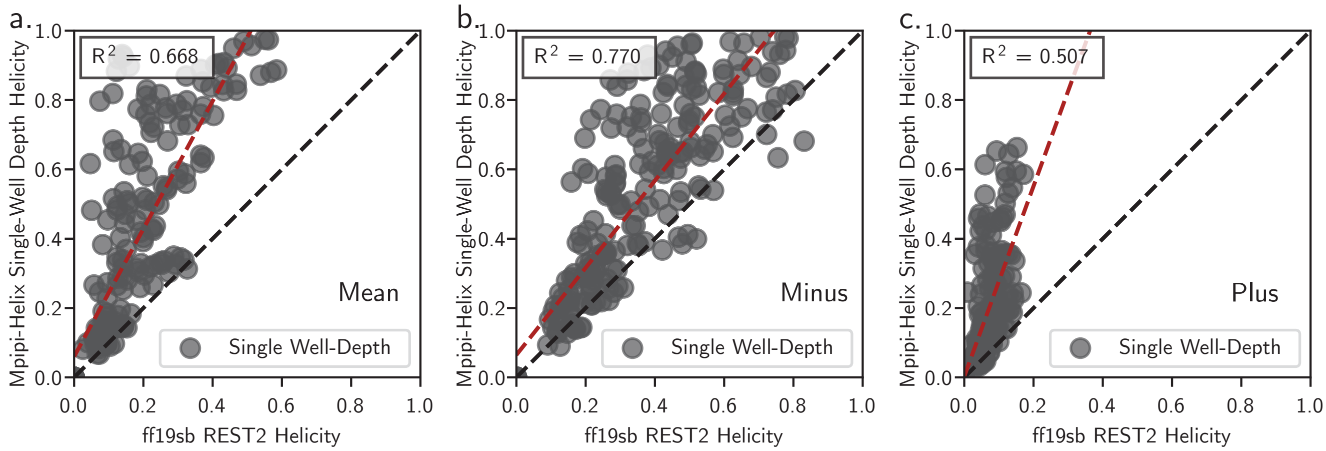


SI Fig. 47. **Comparisons of the target REST2 helical fractions versus single well depth helical fractions in Mpipi-Helix.** The parity plot between the Mpipi-Helix single well depth helical fractions and the target fractions extracted from the REST2 simulations in the **a,** mean, **b,** minus two standard deviations, and **c,** plus two standard deviations datasets.

#### III. MPIPI-HELIX MODEL VALIDATION

##### A. A11 Well-Tempered Metadynamics Simulations

We performed well-tempered metadynamics simulations in order to determine the free energy surfaces of the A11 peptide in three different atomistic force fields: ff19sb [1], ff14sb [32], and CHARMM36m [33]. These simulations were performed in GROMACS [34]. Input configurations were made in an extended configuration using PyMol [36] builder. Simulation inputs were generated using CHARMM-GUI [35] with 150 mM NaCl salt. All systems were first minimized using 5000 steepest descent steps with a force tolerance of 1000 kJ/mol/nm and positional restraints on the heavy atoms of the protein. A subsequent equilibration step was performed for 100 ns at a timestep of 2 fs at 298.15 K with a v-rescale thermostat and a 1 ps damping constant. Cutoffs for van der Waals and Coulomb interactions were set to 0.9 nm. Long-range electrostatics were calculated using a PME mesh grid. Sampling was performed under the same simulation conditions but using well-tempered metadynamics [4] with the PLUMED [44, 45] plugin for GROMACS. Specifically, the biased collective variables were the root-mean-squared deviation to a perfect helix (DRMSD in PLUMED) and the radius of gyration. The bias factor was set to 5, with a hill deposition rate of 10 ps, and hill deposition size of 0.025 kcal/mol. Sampling was performed for 5 µs. Finally, during all steps of the simulations, bonds involving hydrogen were constrained with the LINCS algorithm [37].

To demonstrate the convergence of our metadynamics simulations, we have plotted in SI Fig. 48 three free energy surfaces at 4600, 4800, and 5000 ns of simulation time in f19sb, ff14sb, and CHARMM36m. During these last 400 ns, the FESs do not show any significant changes, demonstrating the convergence of the metadynamics simulations.

##### B. Validation Against Atomistic Simulations

We validated Mpipi-Helix against atomistic simulations of alanine-based peptides. In particular, we first simulated the sequences of $A_5XA_5$ in our Mpipi-Helix force field, where X represents one of the twenty canonical amino acids. Each peptide was simulated at 298.15 K in the *NVT* ensemble using a Langevin thermostat with a damping constant of 100 ps. The aggregate simulation time for each peptide was 750 µs comprised of 300 2.5 µs simulations. Each simulation had an equilibration period of 10 ns with frames saved every 1 ns. Classification of helix and coil residues was determined using the cg2all [28] software paired with classifications of the protein backbone using the criteria that $(-30.0 > \phi > -100.0)$ and $(-7.0 > \psi > -67.0)$ constitutes helical residues.

The atomistic counterpart to these Mpipi-Helix $A_5XA_5$ simulations were done in the ff19sb [1], ff14sb [32], and CHARMM36m [33] force fields. In particular, $A_5XA_5$ peptides were simulated each in the *NVT* ensemble at 298 K using the GROMACS simulation software [34]. All simulations were run with a prior minimization step using 5000 steepest descent steps with a force tolerance of 1000 kJ/mol/nm and positional restraints on the heavy atoms of the protein. Equilibration was performed with positional restraints still maintained for 125 ps at a 1 fs timestep. For the ff19sb simulations, a Nosé–Hoover thermostat was used with a time coupling of 1 ps. For both ff14sb and CHARMM36m, the v-rescale thermostat was used with the same time coupling as ff19sb. In ff19sb and ff14sb, van der Waals and Coulomb cutoffs were set to 0.9 nm with a PME grid for long-ranged electrostatics. CHARMM36m was similar with a 1.2 nanometer cutoff for van der Waals and Coulomb interactions. Finally, all peptides were simulated for a sampling period of 10 µs with a timestep of 2 fs, taking three even blocks for the calculation of errors using block averaging. In all simulations, the LINCS algorithm constrained bonds involving hydrogen atoms [37]. Inputs were built with CHARMM-GUI [35] with input PDB files built in PyMol [36] and peptides started from an extended conformation.

The helical fractions of every $A_5XA_5$ in each force field are reported in SI Fig. 49. There are significant discrepancies between each of the atomistic force fields, with Mpipi-Helix clearly falling within the range of those discrepancies. Generally, CHARMM36m has the lowest helical content out of all the force fields, with Mpipi-Helix having the highest, although it depends on the peptide sequence. Notably, Mpipi-Helix shows less variation to the guest residue identity than the atomistic force fields. As noted in the discussion of our parameterization of Mpipi-Helix, we found that if we matched the helical content of guest–host alanine-based peptides well, our model was to responsive to mutations in more complex protein sequences. This is due to alanine's large response to helix-breaking residues compared to other amino acids, as alanine's low entropic penalty to helix formation is highly modulated by the presence of other amino acids [17].

Despite Mpipi-Helix's inability to respond strongly to mutations in an all alanine-based sequence, its helical fractions still trend linearly with those from the atomistic force fields. In particular, the parity plots which describe how well Mpipi-Helix matches the helical fractions of ff19sb, ff14sb, and CHARMM36m are shown in SI Fig. 50. While points for these parity plots do not fall along the $y = x$ line, the $R^2$ values are 0.461, 0.489, and 0.412 for Mpipi-Helix compared to ff19sb, ff14sb, and CHARMM36m. This indicates a moderate linear trend between the Mpipi-Helix predictions and the atomistic force fields. Again, Mpipi-Helix tends to overpredict helical fractions for alanine-based sequences, but this is by design. As part of our parameterization process, we found that correctly predicting the response to mutations for alanine-based peptides

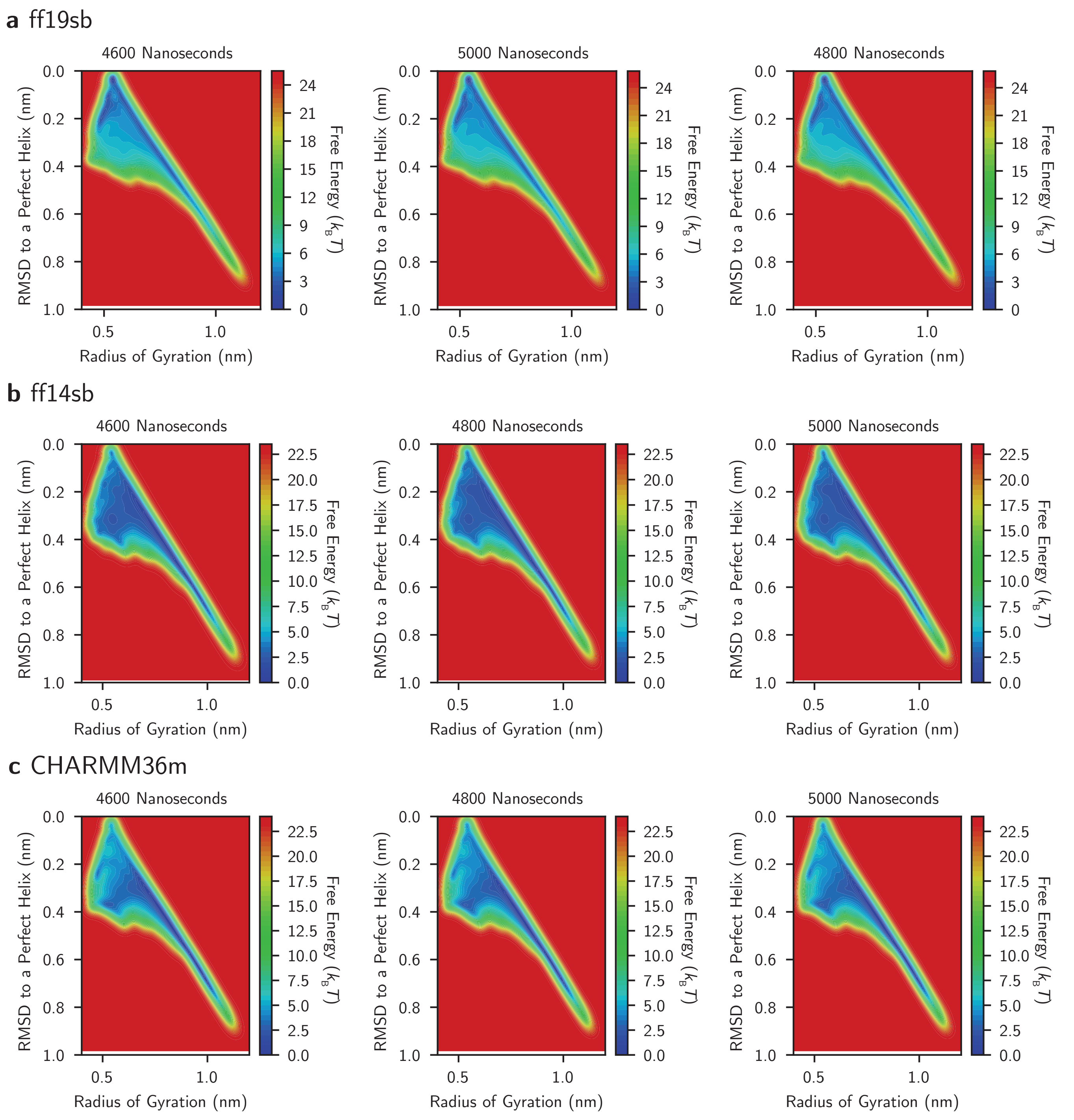


SI Fig. 48. **Convergence of metadynamics simulations of A11 peptide.** The convergence of simulations of A11 peptide in **a,** ff19sb, **b,** ff14sb, and **c,** CHARMM36m as demonstrated by convergence of free energy surfaces during the last 400 ns of sampling time.

meant we could not accurately predict stable peaks in helical fractions which is discussed in the next section on 'Validation Against NMR Measurements and Contemporary Residue-Level Force Fields'.

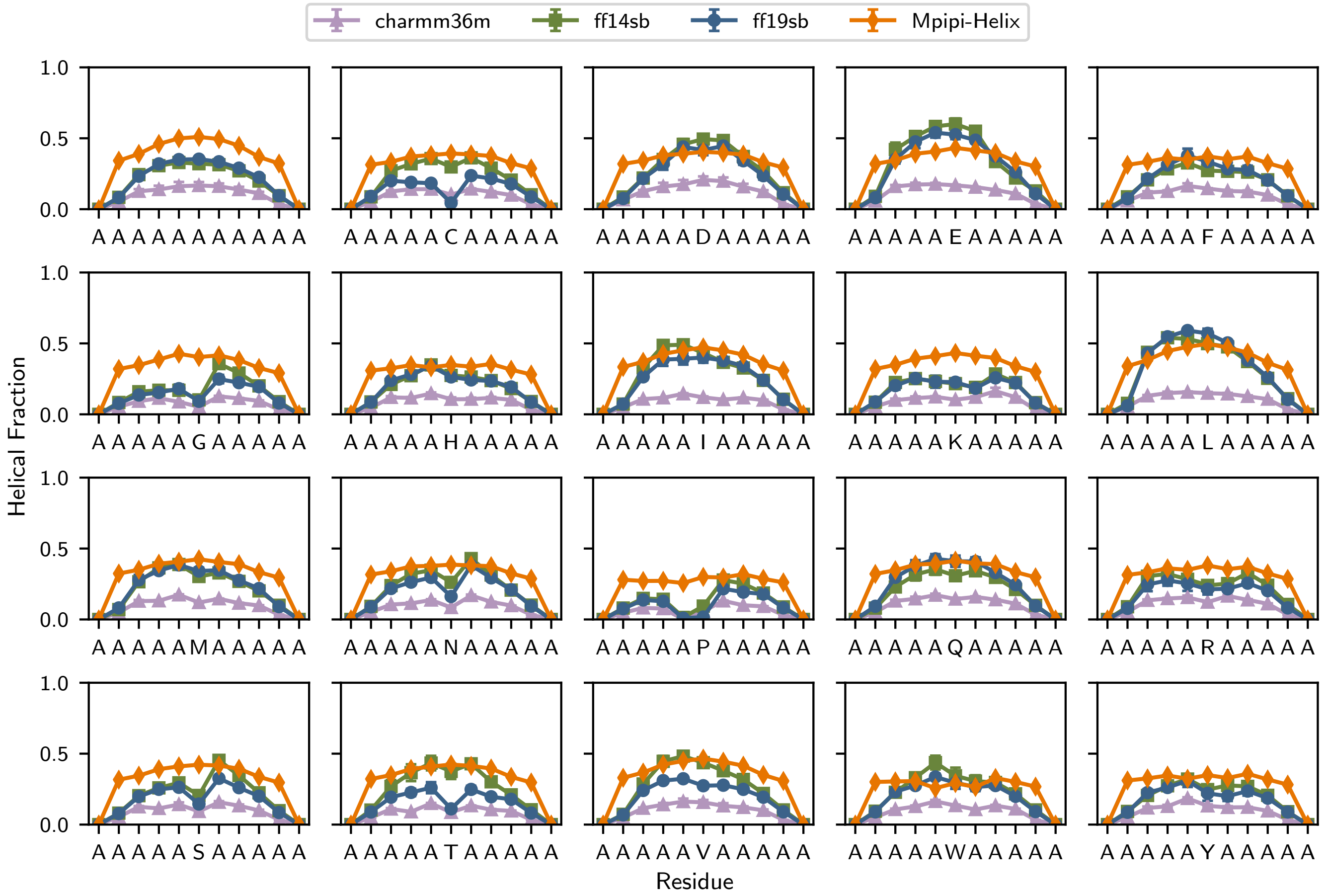


SI Fig. 49. **Helical fractions of $A_5XA_5$ peptides in the Mpipi-Helix, ff19sb, ff14sb, and CHARMM36m force fields.**

##### C. Validation Against NMR Measurements and Contemporary Residue-Level Force Fields

Because atomistic simulations are limited in the length and sequence complexity of peptides they can reasonably sample, and because experimental analogs of the 11 length peptides were in our training dataset, we further validated our Mpipi-Helix force field by simulating IDPs containing helical domains whose helical fractions have been measured using NMR. The proteins we validated against include TDP43 (BMRB 26823), TDP43 with 5 methionine to alanine mutations (TDP43 5M to A) in its helical domain (BMRB 52060), TDP43 with an A326P mutation (BMRB 26828), ANXA11 (BMRB 52944), Androgen Receptor (AR) (BMRB 51479 and 51480), CREB1 (BMRB 27648), FCP1 (BMRB 16296), GAB1 (BMRB 51019), and Hepatitis Core Virus-C (HCV-C) (BMRB 15768). For the AR protein, we combined two chemical shifts, one for the N-terminal domain of the protein and one for the Tau-5 region, as done in Ref. [46] due to difficulties in measuring both regions accurately in the same NMR experiment.

We developed a procedure for extracting helical fractions from the NMR chemical shifts leveraging the d2D software [47] to simulate the IDPs and their associated helical domains in Mpipi-Helix. Our first step was to apply d2D directly to the chemical shifts as extracted from the BMRB. This resulted in the 'experimental NMR' helical fraction which is the depicted in the dashed black line in SI Fig. 51. Next, we used the predicted helical fractions of each residue to determine which residues to assign our Mpipi-Helix helical potential to in our coarse-grained simulations. Specifically, we took the predicted helical fractions and smoothed them in a running window by taking the helical fraction of residue $i$ to be the average of that between residues $i-2$ to $i+2$. We then utilized a cutoff of 5% helical fraction to determine which residues were set to be helical and which were set to be coil in our simulations, noting that d2D predictions tend to be either less than 1% for non-helical regions and greater than 5% for helical regions. Finally, we added a 'helical cap' to each end of the helical domains by classifying three more residues as helical on either side of the domain, thereby ensuring that every residue with $>5\%$ helical content in the helical domain has total inclusion in all possible helical potentials (i.e., dihedral and bond angles) along the protein chain. Using the results from this procedure, we assigned helical potentials to the regions classified as helical using the Mpipi-Helix model.

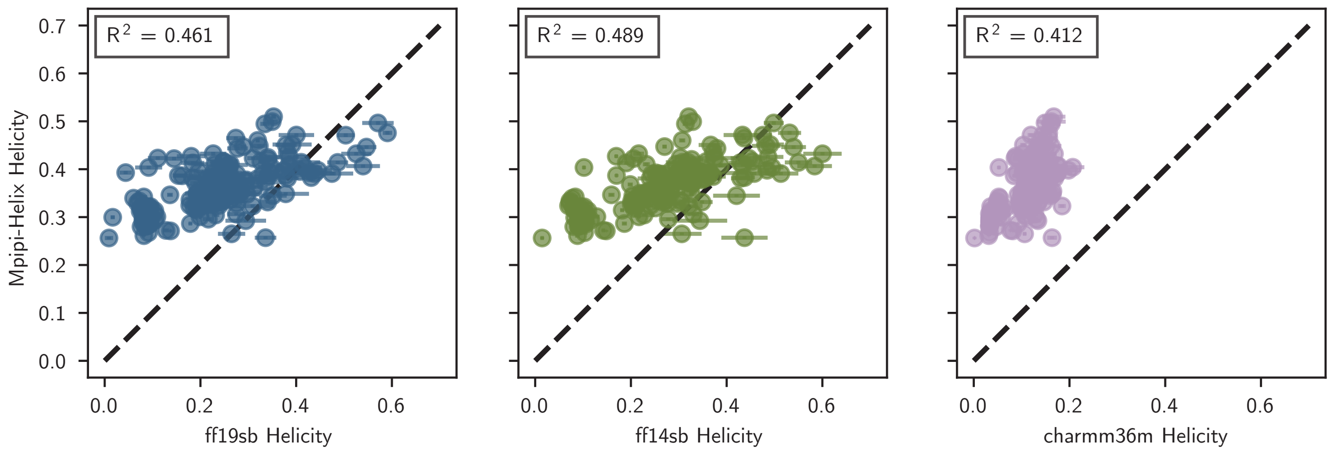


SI Fig. 50. **Parity plots of helical fractions of $A_5XA_5$ peptides.** Parity plots comparing the helical fractions of $A_5XA_5$ peptides simulated in Mpipi-Helix versus ff19sb (left), ff14sb (middle) and CHARMM36m (right).

It is important to note that the NMR experiments were not all performed at the conditions Mpipi-Helix was parameterized for (i.e., 298 K, 150 mM salt, and a neutral pH). In particular, the TDP43 measurements which were performed at a pH of 6.1 and 0 salt. Additionally, ANXA11 and the Androgen Receptor were measured at 278 K, a pH of 7.4, and salt conditions of 250 mM and 20 mM respectively. HCV-C was also measured at 278 K and a pH of 6.6. Thus, while we did adjust the simulation temperature to match the experimental conditions each protein was measured at, we did not adjust the salt concentration or pH of our Mpipi-Helix simulations to prevent straying too far from the parameterized conditions. Further, the helical fractions from the d2D software [47] are only predictions based on the NMR measurements. Despite these caveats, NMR measurements of these helical domains are the best comparison point to validate our Mpipi-Helix model as they are measured from real protein sequences and include cross interactions of disordered regions with $\alpha$-helix domains.

We also applied the same helical assignments to two contemporary residue-level coarse-grained force fields that have recently been published describing $\alpha$-helices in disordered proteins: the HPS-SS [10] and Mpipi+ [11] models. While neither of these models were published with procedures for isolating helical domains along a protein chain (e.g., they apply their helical potentials along the entirety of a protein chain), we used LAMMPS `special_bonds lj/cut 0.0 1.0 1.0` command paired with 'fake' bonds of zero energy to ensure that disordered regions of the chain could be described by FENE-like bonds whereas the helical regions would have the models' helical potential applied and excluded pairwise interactions within the helix. In this way, we could compare Mpipi-Helix with two contemporary coarse-grained models that represent $\alpha$-helix structure in disordered proteins.

With our sequences and helical assignments in place, we simulated each of the BMRB peptides in each force field, using the same cg2all [28] backmapping technique with $\phi$ and $\psi$ backbone angle classifications to determine helical fractions for each model. In particular, we simulated each peptide in each force field using 3 independent simulations each of 10 µs in length. Each simulation was performed with a Langevin thermostat using a 100 ps damping constant and a 10 fs timestep. Frames were recorded every 1 ns. Temperatures reflected exactly what was used in the experimental conditions from the NMR measurements (26823 - 298 K, 52060 - 298 K, 26828 - 298 K, 52944 - 278 K, 51479 + 51480 - 278 K, 27648 - 298 K, 16296 - 298 K, 51019 - 298 K, 15768 - 278 K).

After classifying the helical fractions from each peptide and model, we see that Mpipi-Helix performs better at reproducing the experimental NMR derived helical fractions. In particular, the primary deficiency of the HPS-SS and Mpipi+ models appears to be the inability of generating defined and sustained peaks in helical fractions, instead exhibiting fluctuations in the helical fraction along the sequence. Examples of this can be seen in the TDP43 and ANXA11 proteins where HPS-SS and Mpipi+ have several sharp peaks whereas the NMR predictions only predict one or two peaks within the same helical domain.

Overall, given how well Mpipi-Helix matches the peaks in the predicted helical fractions from NMR measurements validates its use in exploring helical domains within disordered proteins. Further, as mentioned in the Methods, the ability to represent disordered and $\alpha$-helical segments within the same simulation and reproduce experimental observables provides validation for using the Mpipi-Helix model to simulate helical domains within condensates and probing folding landscapes.

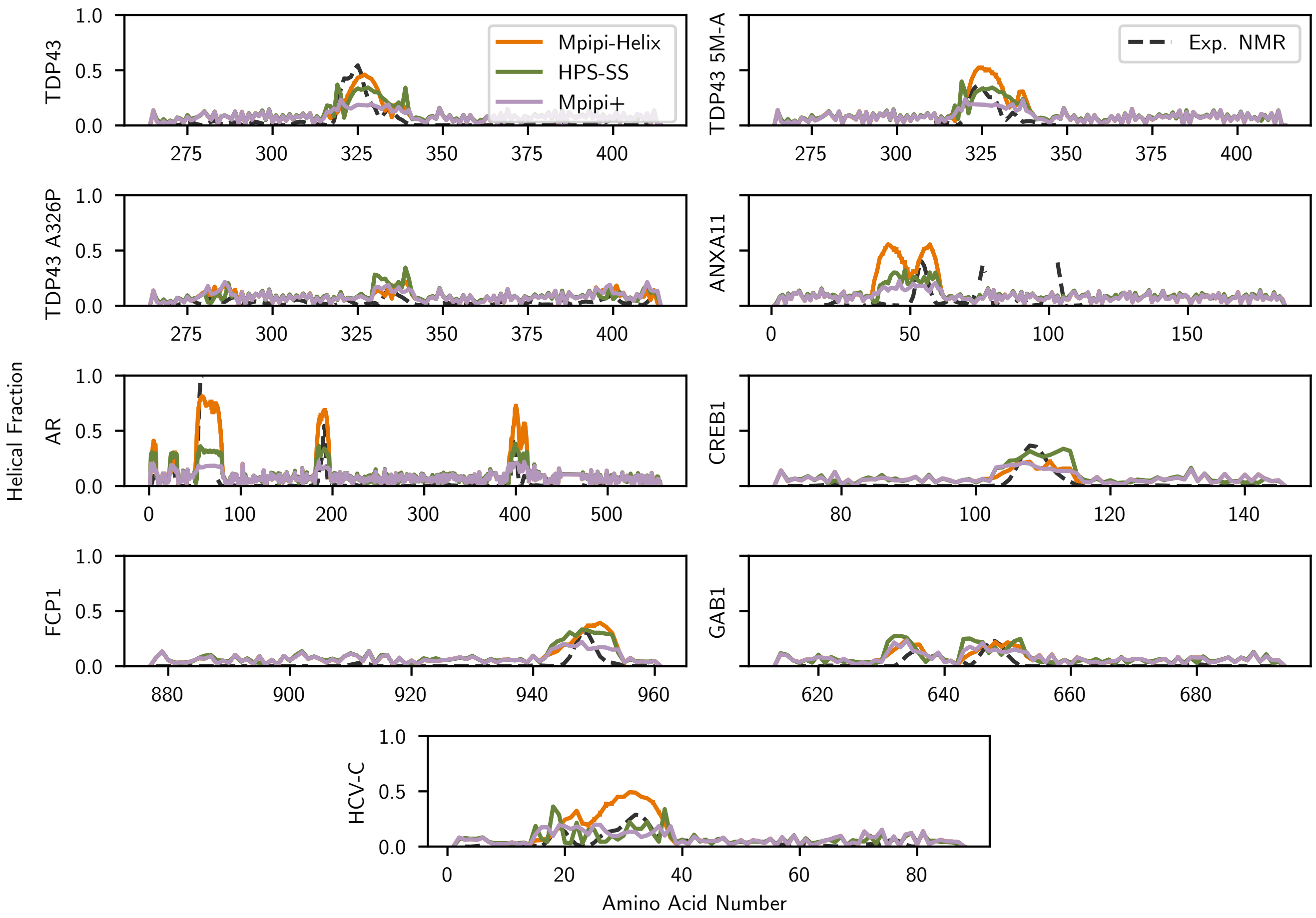


SI Fig. 51. **Experimental NMR derived helical fractions versus Mpipi-Helix, HPS-SS, and Mpipi+ residue-level coarse-grained force fields.** The experimental NMR derived helical fractions using the d2D software [47] compared to three residue-level coarse-grained force fields: Mpipi-Helix, HPS-SS, and Mpipi+.

### IV. HELICAL PEPTIDES IN MODEL CONDENSATES

#### A. Phase Diagrams of Co-Condensate Peptides and Determining Molecular Densities, Co-Condensate Peptide Number, and Crowder Pressure

We designed three co-condensate peptides to make up our model condensates: FYAAYF, NYQQYN, and RYAAYE. Each of these peptides have diverse sequence characteristics (hydrophobic, polar, and charged), yet contain the requisite multivalent residues (e.g., tyrosine) to enable condensate formation. We validated that each sequence could form condensates by calculating each peptide's phase diagram. As an initial step to calculating the phase diagrams, we performed *NPT* simulations to estimate critical temperatures for each peptide. Specifically, we first simulated $12 \times 12 \times 12$ chains in an *NPT* simulation with a Berendsen barostat at 100 atms and a 100 ps damping constant to compress the peptides. This simulation was run for 0.3 ns using a Langevin thermostat at 100 K and a 100 ps damping constant. Afterwards, a binary search algorithm was deployed with simulations run for 1.5 ns at a pressure of 0 atms with the same thermostats as above, but beginning from the original compressed simulation. The binary search used initial bounds of 200 and 400 K (i.e., two initial simulations were performed, one at 200 K and the other at 400 K). Thereafter, simulations were performed at the middle temperature between the highest temperature simulation that produced a condensate (which we classified as a simulation with a density of $> 0.3\text{g/cm}^3$) and the lowest temperature simulation that did not condense. This lead to critical point estimations of 372 K for FYAAYF, 347 K for NYQQYN, and 342 K for RYAAYE.

Afterwards, we chose 4 temperatures below the critical temperature in a 5 K spacing to run direct coexistence simulations in a slab geometry for each peptide. For these simulations, we performed an initial compression of $12 \times 12 \times 12$ peptide chains using a Berendsen barostat at 2 atms of pressure and a 10 ps damping constant. This was alongside a Langevin thermostat with a 100 ps damping constant and a 200 K temperature. We compressed until the density stopped changing for each sequence. Using the compressed output configuration, we seeded direct coexistence simulations in the slab geometry with box dimensions of $150 \times 150 \times 750$ Å$^3$ for all three sequences. These direct coexistence simulations were then run for 1.25 μs in the *NVT* ensemble with a Langevin thermostat and a 100 ps damping constant. Densities were recorded every nanosecond using 100 equally spaced bins along the z-axis.

We then extracted density profiles from our slab simulations. After aligning every profile from the simulation, we extracted the dense phase density as the central 7 bins of each profile and the dilute phase density as the outer 25 bins on either side of the dense phase (see shaded grey regions in SI Fig. 52). Using the density of the dilute and dense phases, we calculated phase diagrams following the laws of rectilinear diameter and coexisting densities, as has been done in other works [7, 48]. The final phase diagrams are provided in SI Fig. 52.

We also performed direct coexistence simulations in the slab geometry at 298.15 K for each of our peptides to determine dense phase densities at room temperature. The slab simulation density profiles are shown in SI Fig. 53. All simulation parameters besides the temperature were identical to those described above and we extracted dense phase densities in the same manner. From the estimation of the dense phase density and the desired box size of $60 \times 60 \times 60$ Å$^3$—which is chosen to be large enough to avoid finite-sized effect—we calculated the number of co-condensate peptide chains required to simulate alongside our helical peptides and form a condensate. In particular, we calculated that we would require 144 chains for FYAAYF, 141 chains for NYQQYN, and 145 chains for RYAAYE.

Finally, for the crowded solution, we decided to match the number of co-condensate peptide chains as is in the FYAAYF condensate (e.g., use 144 chains of 6-length coarse-grained PEG). We then calculated the pressure that would match the box size of FYAAYF ($\sim 60$Å$^3$) using *NPT* simulations (SI Fig. 54). For these simulations, we simulated 144 6-length PEG chains at various pressures using a Berendsen barostat with a 100 ps damping constant and a Langevin thermostat at 298.15 K and a 100 ps damping constant. These simulations lasted either 350 ns (points with larger error bars in SI Fig. 54 where the error is calculated from block averaging 3 blocks of data) or 600 ns (points with smaller error bars in SI Fig. 54) with 100 ns being discarded as equilibration. The final pressure that matched the FYAAYF box size was 71.3 atms.

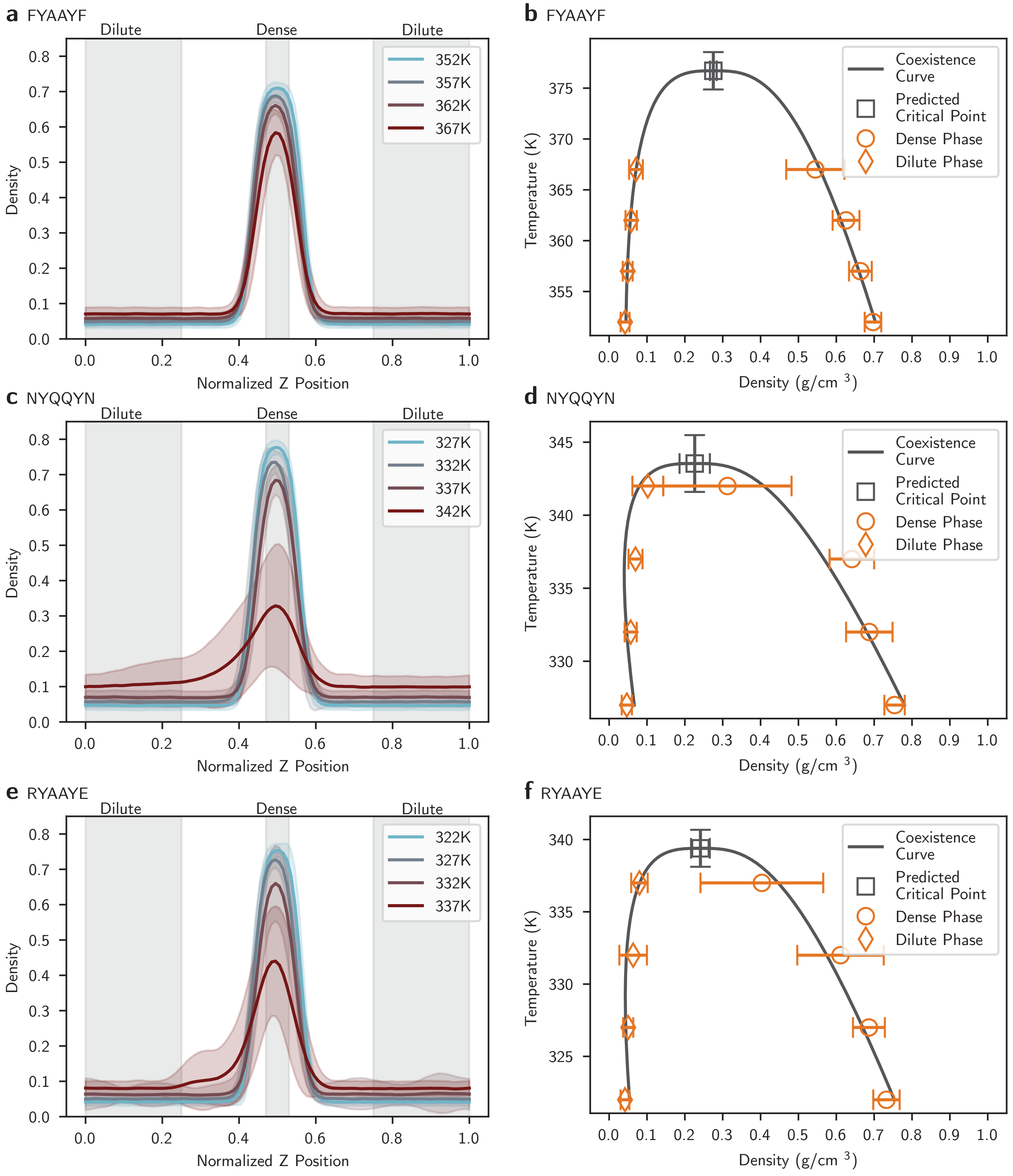


SI Fig. 52. **Density profiles and phase diagrams of co-condensate peptides.** The density profiles of the **a,** FYAAYF, **c,** NYQQYN, and **e,** RYAAYE peptides in Mpipi at temperatures close to their critical temperatures. The phase diagrams of **b,** FYAAYF **d,** NYQQYN, and **f,** RYAAYE from the extracted dense and dilute phase densities in **a**, **c**, and **e**. Phase diagrams were fitted using the laws of rectilinear diameter and coexisting densities. Errors were calculated using the standard deviation of the protein densities and propagated into the calculation of the critical point.

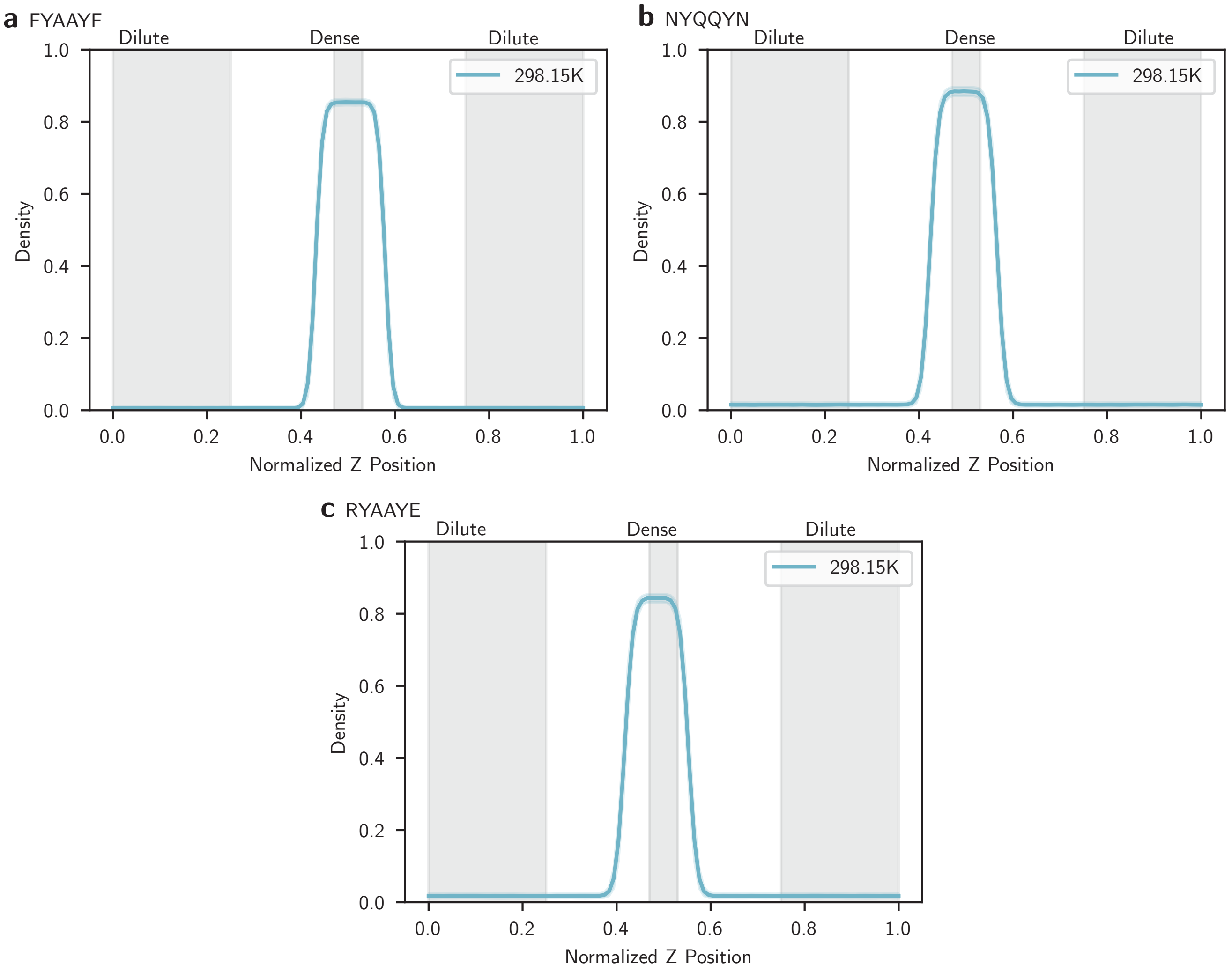


SI Fig. 53. **Density profiles of co-condensate peptides at 298.15 K.** The density profiles of the **a,** FYAAYF, **b,** NYQQYN, and **c,** RYAAYE peptides in Mpipi at 298.15 K. Densities were extracted from direct coexistence simulations in a slab geometry and error is the standard deviation over the course of the simulation.

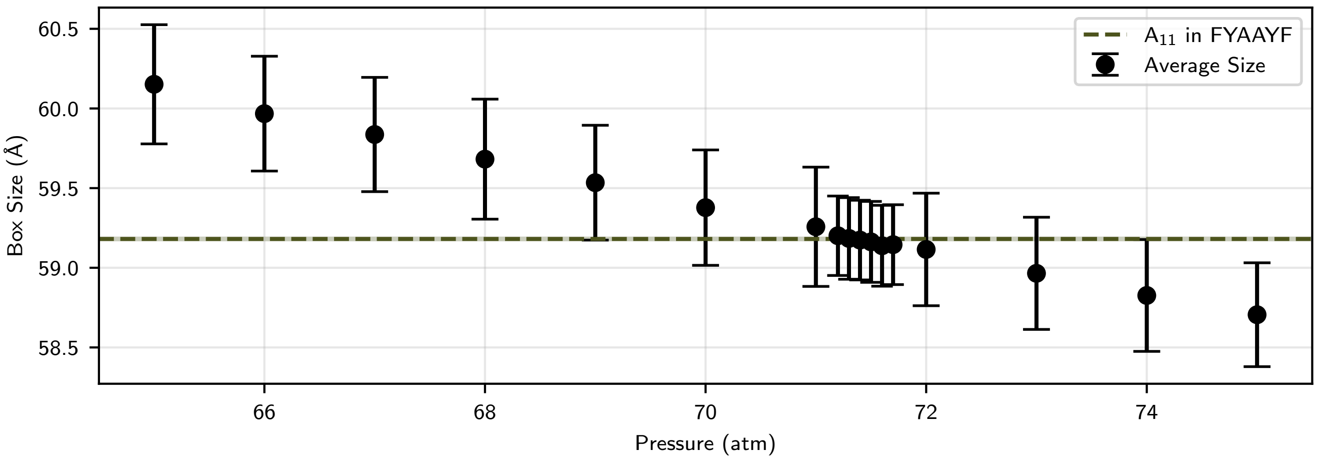


SI Fig. 54. **Pressure calculations to match FYAAYF molecular densities in crowder.** Results of simulations that measured box size versus the pressure applied to a system of A11 in PEG. The final pressure of 71.3 atms reproduces the box size of A11 in FYAAYF, matching FYAAYF's molecular density.

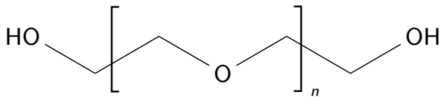

SI Fig. 55. The chemical formula for polyethylene glycol (PEG).

##### B. PEG Model - Excluded Volume Polymer

We parameterized a model for polyethylene glycol (PEG) that is compatible with Mpipi-Helix. Here, our main goal was to capture PEG's behavior as an entropic crowder. As such, we parameterized PEG as disordered polymer with excluded volume interactions, ignoring any specific chemical interactions with amino acids in polypeptides. Accordingly, the Hamiltonian that describes PEG interactions includes only an excluded volume interaction, the Weeks-Chandler-Anderson (WCA) potential, and a harmonic bond potential. These terms are describe in Equation 30.

$$U_{\text{total}} = U_{\text{WCA}} + U_{\text{Bond}}. \tag{30}$$

The WCA potential is a purely repulsive potential described by $r_{ij}$, which is the distance between interacting monomers, $\sigma$, which is the effective size of a monomer, and $\epsilon$, which is the repulsive strength between the monomers. The form of the WCA potential is described in Equation 31.

$$U_{\text{WCA}} = 4\epsilon\left[(\frac{\sigma}{r_{ij}})^{12} - (\frac{\sigma}{r_{ij}})^{6}\right] + \epsilon,\ r_{ij} \leq 2^{1/6}\sigma. \tag{31}$$

The harmonic bond potential is a bonded potential between consecutive polymers on the PEG chain. It is described by $r_{ij}$, which is the distance between consecutive monomers on a PEG polymer, $K$, which is the bond stiffness, and $r_0$, which is the equilibrium bond length. The form of the harmonic bond potential is described in Equation 32.

$$U_{\text{Bond}} = K(r_{ij} - r_0)^2. \tag{32}$$

There are thus four parameters that need to be specified in order to simulate PEG polymers. These are $\epsilon$, $\sigma$, $K$, and $r_0$.

We began by specifying $\epsilon$ and $\sigma$ of the WCA potential. First, we set $\epsilon = k_{\text{B}}T$ for $T = 22°\text{C}$ ($\approx 0.586\frac{\text{kcal}}{\text{mol}}$) to be on the level of thermal energy at room temperature. This $\epsilon$ term does not depend on the partner the PEG monomer is interacting with. In this way, PEG represents a general entropic crowder that imparts volume exclusion towards all components in a system.

In order to determine the bead size for a PEG monomer, we considered a representation of PEG with a central oxygen site in the repeating polymer formula (displayed in SI Fig. 55). ProteinVolume 1.3 was used to determine the effective van der Waals volume of a PEG monomer [49]. Specifically, a PDB file of the PEG monomer was uploaded to the ProteinVolume 1.3 webserver which calculated an estimated volume of 52.441 Å$^3$. Approximating the monomer to be a sphere, the radius of a PEG mononmer is $r = 2.322$ Å. Given this radius for the PEG monomer, we wished for the PEG WCA potential to become repulsive at a distance of $r = 4.644$ Å for PEG–PEG interactions. Due to the truncation and shifting of the WCA potential, this results in a $\sigma = 4.137$ Å for PEG–PEG interactions. For all other interactions between PEG monomers and coarse-grained residues represented in Mpipi-Helix, the size of interaction follows an arithmetic mixing rule for $\sigma$, with the $\epsilon$ value remaining unchanged. Additionally, from the representation of PEG centered on the oxygen site, we calculated that the molar mass of a PEG monomer as approximately 45.601 g/mol. Finally, to accurately describe the bond distance distribution of PEG, we carried out atomistic simulations of single PEG polymers in OPLS [50] with TIP4P-2005 water [51] and a 150 mM concentration of NaCl [52]. We then used this data to fit the bonded interactions of our coarse grained PEG model.

To prepare the simulations with PEG, LAMMPS [6] configuration and potential files were generated for a 20-length PEG polymer according to the OPLS force field. Next, using a custom code the PEG configuration file was solvated with TIP4P water molecules and the potential file was updated with TIP4P-2005 water interaction parameters. This initial solvated configuration was in a $35.0 \times 35.0 \times 35.0$ Å$^3$ box. From here, an $NPT$ simulation was performed using LAMMPS to relax the initial configuration of PEG and water molecules. Specifically, the simulation was carried out with a time step of 0.5 fs for a duration of 1 ns. Long-ranged Coulombic interactions were calculated using a PPPM grid with a 1e−4 tolerance on force errors. Additionally, the TIP4P water molecules were integrated using LAMMPS `rigid/npt/small` command and the PEG polymer was integrated with a Nosé-Hoover thermostat. The pressure on the water molecules was set to be 1 atmosphere with a damping constant of 100 fs. The temperature was set to be 295.15 K with a damping constant of 100 fs for both the water molecules and PEG polymer. The velocity-Verlet algorithm was used to integrate the system in time.

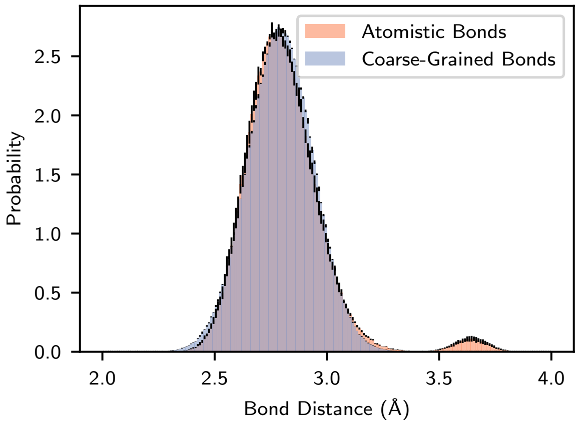


SI Fig. 56. **Bond distance distributions for single-chain simulations of PEG at atomistic and coarse-grained resolutions.** Atomistic simulations were performed in the *NVT* ensemble with the OPLS force field, TIP4P-2005 water, and 150 mM of NaCl. Coarse-grained simulations were performed in implicit solvent using a Langevin thermostat and the Hamiltonian in Equation 30. Error bars on the distributions indicate the standard error from three simulation replicates at each resolution.

After relaxation, sodium and chloride ions were place into the configuration manually to achieve a 150 mM concentration of NaCl. An additional 1 ns *NPT* simulation was performed in order to relax the system. The conditions for this simulation were identical to the previous relaxation run described above, with the ions present also under the Nosé-Hoover thermostat. From the solvated configuration with ions, three simulation replicates were run to sample the bond distance distribution of the PEG polymer. Simulations were run in the *NVT* ensemble using a Nosé-Hoover thermostat on the PEG polymer and ions. The TIP4P water was run with a `rigid/nvt/small` thermostat. For both, the temperature was set to be 295.15 K with a 100 fs damping constant. All simulation settings were the same as described above. Each replicate simulation was run for 10 ns with configurations of the PEG polymer saved every picosecond. The bond distance distribution was calculated using a custom code.

A custom Monte Carlo code was written to guess what parameters would produce an adequate match for the bond distance distribution between the coarse-grained and atomistic simulations. After a series of guesses using the Monte Carlo code, single chain coarse-grained simulations of the PEG polymers were run in LAMMPS to determine how well the distributions matched atomistic simulations. Specifically, single-chain PEG simulations were run at 295.15 K in implicit solvent using a Langevin thermostat with a damping constant of 100 ps integrated with the velocity-Verlet algorithm in LAMMPS. The timestep was 10 fs. The length of the PEG polymer in these coarse-grained simulations were also 20, matching the atomistic simulations. The coarse-grained Hamiltonian is described as Equation 30 above. A value of $K = 13.5\,\text{kcal/mol}$ and $r_0 = 2.76$ Å was used as the final bonded parameters after manual tuning following the guesses with the Monte Carlo code. The atomistic and coarse-grained bond distance distributions are plotted in SI Fig. 56. SI Fig. 56 indicates a very accurate match between the two resolutions.

We thank Shannon Zhang and Dr. Michael Webb for helpful code in setting up OPLS configurations and force field parameters for PEG. We also thank Dr. Alina Emelianova for helpful discussions regarding the coarse-graining of PEG.

##### C. Density and Volume Fraction Calculations

Five independent simulations were run separately for FYAAYF, NYQQYN, RYAAYE, and PEG crowder to determine the densities and volume fractions of the co-condensate proteins. Besides the absence of the helical peptide, they were run in an analogous manner as the model helical peptides in condensates simulations, with 2.5 µs of sampling time (see Methods). Thereafter, volume fractions were calculated based on the average self-interaction $\sigma$ value of the constituent co-condensate monomers and the volume of the simulation box for each frame.

##### D. Markov State Models and PCCA+ Spectral Clustering

We utilized Markov State Models (MSMs) to analyze the thermodynamics and kinetics of our helical peptides within disparate environments (e.g., dilute solutions, crowded solutions, and condensates). All MSMs were built with the pyEMMA toolkit [53]. In the construction of our MSMs, we assumed that the degrees of freedom associated with the co-condensate peptides relaxed on timescales faster than the lag time of our MSM. This allowed us to not include the degrees of freedom of the co-condensate peptides within our MSMs (e.g., FYAAYF, NYQQYN, RYAAYE, or the PEG crowder). This assumption is reasonable because the peptides are of a short length and are entirely disordered while the helical peptides undergo a much longer folding process (Extended Data Fig. 1d). In other words, the co-condensate peptide kinetic transitions are on a much faster timescale than the kinetic transitions associated with helix folding/unfolding. Thus, the effects of the co-condensate peptides are implicitly included in the MSMs by their impact on the helical peptide's degrees of freedom.

For all MSMs, we first went through a parameter scanning process to determine model hyperparameters (e.g., input features, lag time, number of clusters). All of the relevant parameters are shown in SI Fig. 57 for each of our helical peptides in FYAAYF, with the dilute, crowded, NYQQYN, and RYAAYE following very similar behavior (not shown). In particular, we utilized the VAMP2 score for to select the best input features (SI Fig. 57a), the VAMP2 score to determine the optimal number of clusters (SI Fig. 57b), and a sweep of lag times to determine which optimally preserved kinetic timescales (SI Fig. 57c). For the input features, we considered whether to include calculated bond and dihedral angles along the 11 length peptide (Angles), pairwise distances between non-bonded neighbors (Pairwise Distances), or both (All) (SI Fig. 57a). As including both the bonded information and the pairwise distances maximized the VAMP2 score, we proceeded with that for all MSMs. We found that the number of clusters that maximized VAMP2 score without overfitting in each condition was approximately 1000 cluster centers (SI Fig. 57b), although marginal improvements are predicted for the AL and AQ systems with slightly larger cluster numbers. For a lag time, we settled on 2 ns as there was approximately no change in the implied timescales of the kinetic processes for larger times (SI Fig. 57c). Thus, for all MSMs, our construction utilized time-lagged independent component analysis for input features of both bond and dihedral angles and pairwise distances, 1000 cluster centers, and a lag time of 2 ns. All plots for SI Fig. 57 were generated using PyEMMA's software.

We further validated our MSMs using the Chapman–Kolmogorov test (SI Fig. 58). We tested all of our helical peptides in the FYAAYF condensate with pyEMMA's built-in CK test function. As we later used the PCCA+ algorithm with four metastable states for interpreting our MSMs, we also used four states to test our models against actual simulation data. SI Fig. 58 demonstrates that for each peptide in the FYAAYF condensate, the prediction of transitions between each of the metastable states using the model matches the empirical statistics from the simulation data. This validates our choice of hyperparameters from the above analysis. The CK tests for the peptides in the other condensates agrees with these plots (not shown).

Additionally, we used the PCCA+ spectral clustering algorithm provided by pyEMMA to determine the timescales of transitions between metastable states in our MSM. Critically, PCCA+ clustering preserves the original timescales of the Markov process which allowed us to interpret the underlying kinetic transitions of our $\alpha$-helices in a coarse-grained manner [54]. SI Fig. 59 shows the details of PCCA+ mapping using A11 in FYAAYF as an example. Here, SI Fig. 59a-b shows the stationary probability density and free energy of the A11 peptide projected onto the first two time-lagged independent components from our MSM analysis. SI Fig. 59c then shows the distinct clusters that were used to build the data for SI Fig. 59a-b. From these three panels, it is impossible to distinguish coarse-grained states by eye. However, after applying PCCA+ and requesting four states, we can impose sharp boundaries to PCCA+ and observe four distinct states (SI Fig. 59d). From this point, we can show why PCCA+ is an excellent choice for our application. In particular, the decomposition into metastable states is intuitive for our helix–coil transition process with four states that map to a coil, a C-terminal helix state, an N-terminal helix state, and a full helix. These states are depicted in SI Fig. 59d, but their interpretation is quite clear in SI Fig. 60 and SI Fig. 61. For SI Fig. 60, we plot the probability density of each state in our PCCA+ clustering onto the order parameters of radius of gyration and root-mean-squared deviation to a perfect helix, which clearly shows a helical state (top left panel), two intermediate states (top right and bottom left panels), and a coil state (bottom right panel). This evidence is further supplemented by analyzing the helical content of each state, which we do for all peptides in all environments in SI Fig. 61. Here, the left most column of plots clearly correspond to full helices given their high helical contents. The next two columns show N-terminal and C-terminal helices, respectively, as seen in

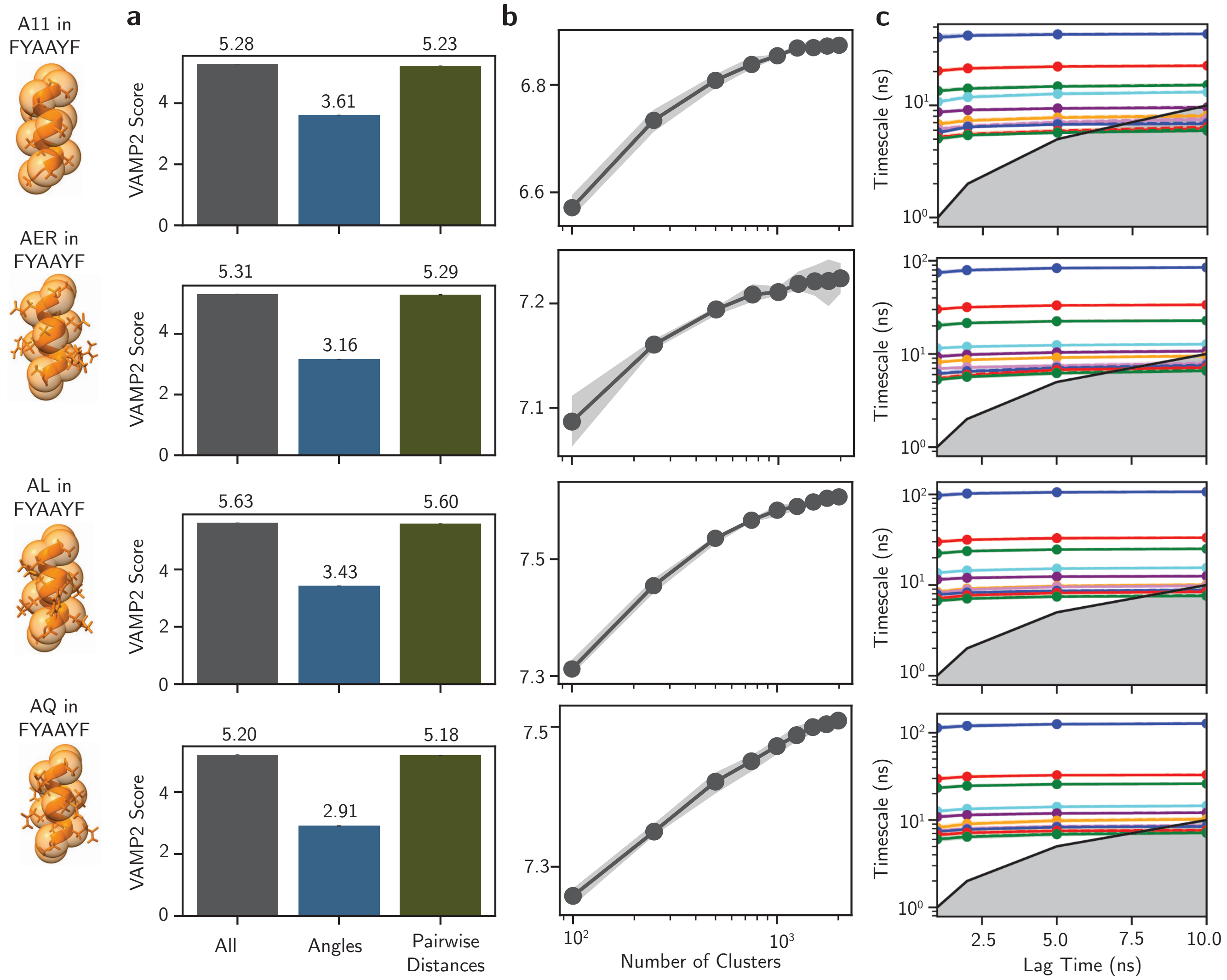


SI Fig. 57. **MSM hyperparameter validation. a,** Validation of input features for MSMs was performed using VAMP2 analysis on input bond/dihedral angles (Angles), all non-bonded pairwise distances (Pairwise Distances), and both feature sets together (All). **b,** Validation of the number of cluster centers for kmeans++ clustering was performed using VAMP2 and a lag time of 2 ns (y-axis is VAMP2 score). **c,** Validation of the lag time was performed using an implied timescale analysis. From this analysis, we chose to use all input features paired with time-lagged independent component analysis with 1000 cluster centers at a lag time of 2 ns.

the relative positions of their helical fraction peaks. Finally, the low helical content of the last column (especially compared to the prior three) demonstrates a coil state. Overall, we find PCCA+ is a highly interpretable and very useful framework for extracting thermodynamic and kinetic information from our MSMs.

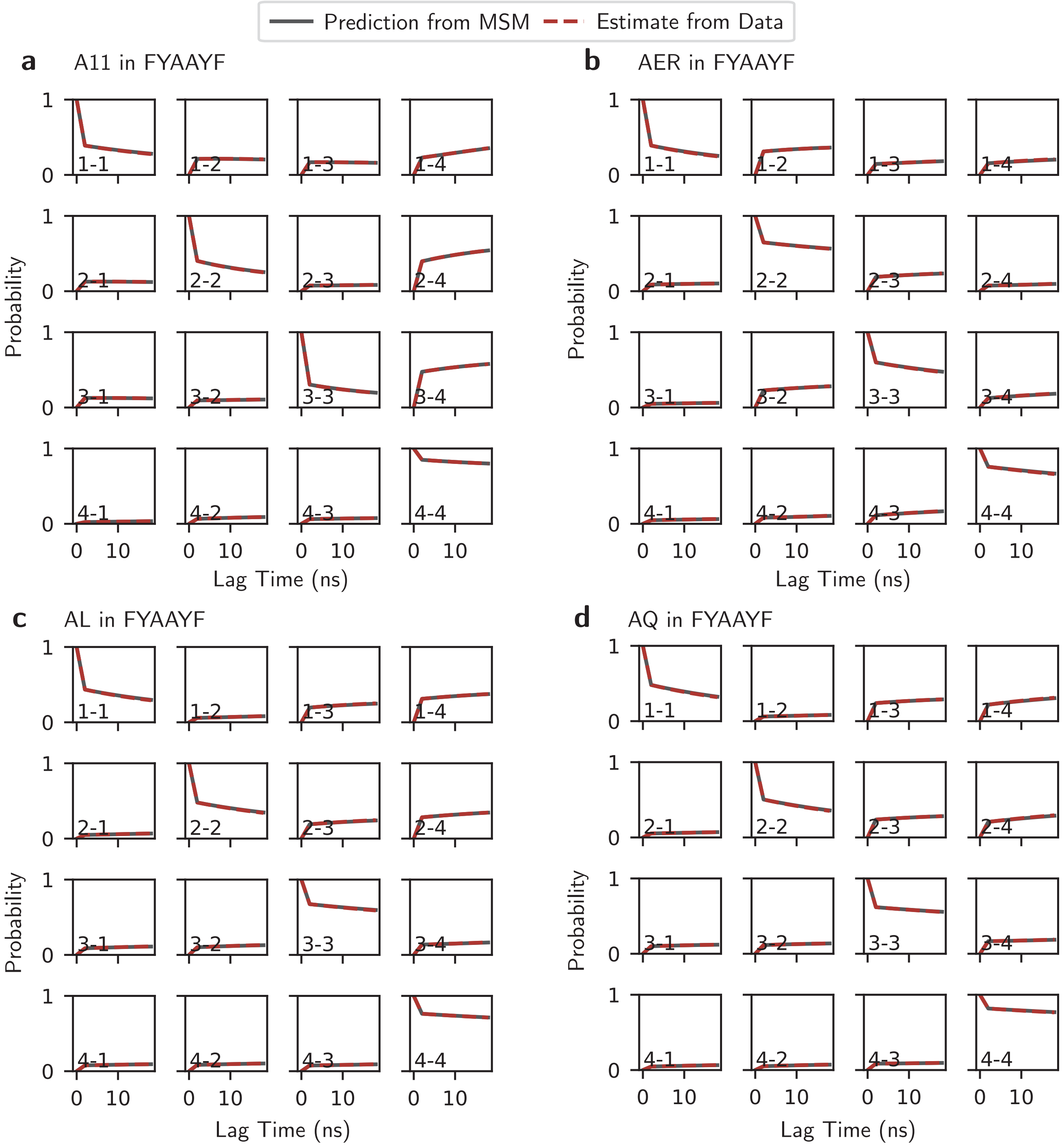


SI Fig. 58. **Chapman–Kolmogorov test for helical peptides in FYAAYF.** Validation of MSM construction using the Chapman–Kolmogorov test on **a,** A11, **b,** AER, **c,** AL, and **d,** AQ in the FYAAYF condensate.

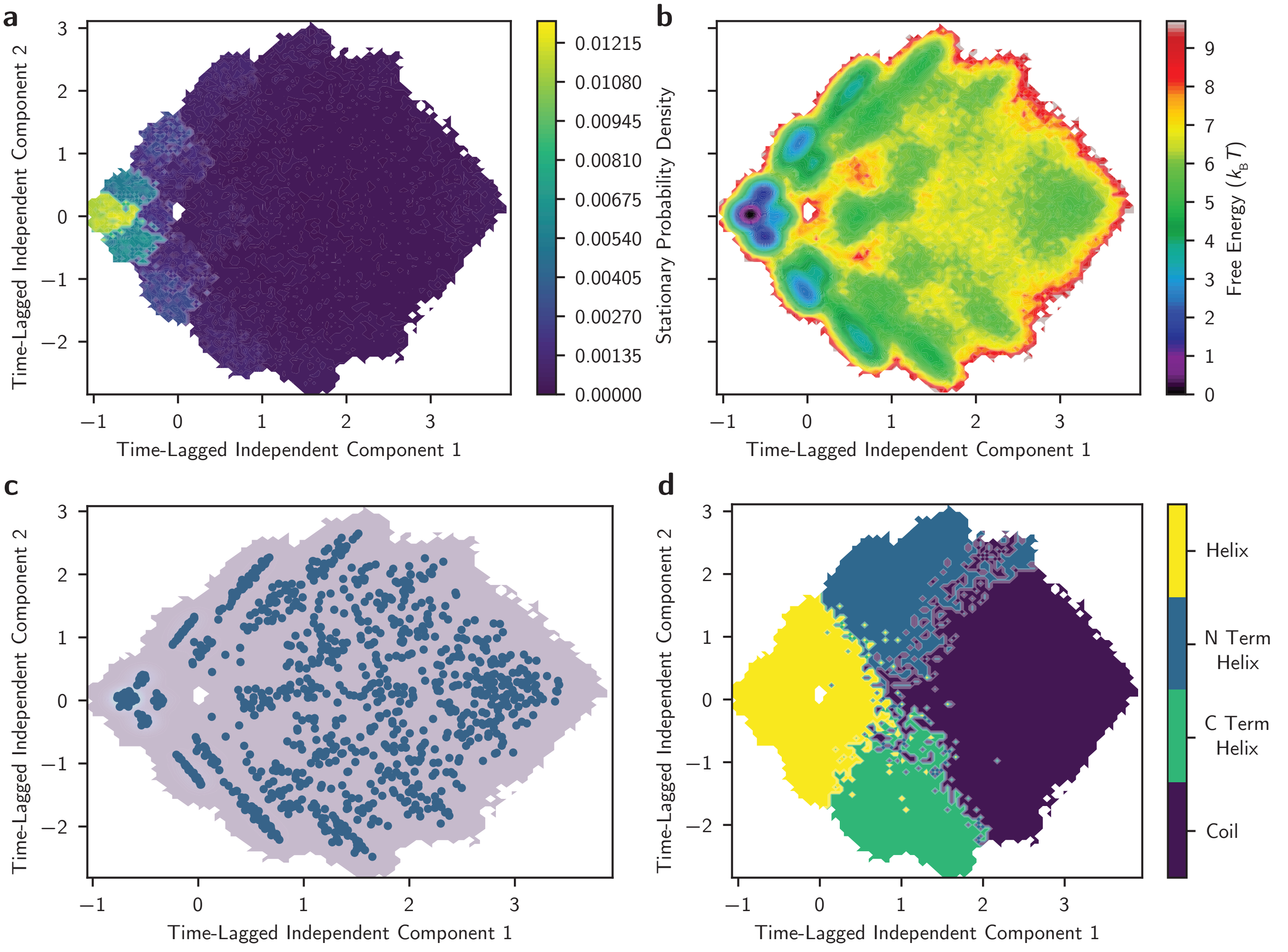


SI Fig. 59. **Projecting the MSM onto PCCA+ metastable states. a,** The stationary probability and **b,** free energy from A11 in FYAAYF after MSM construction. **c,** Cluster centers used in MSM construction. **d,** Metastable states from PCCA+ with sharp boundaries.

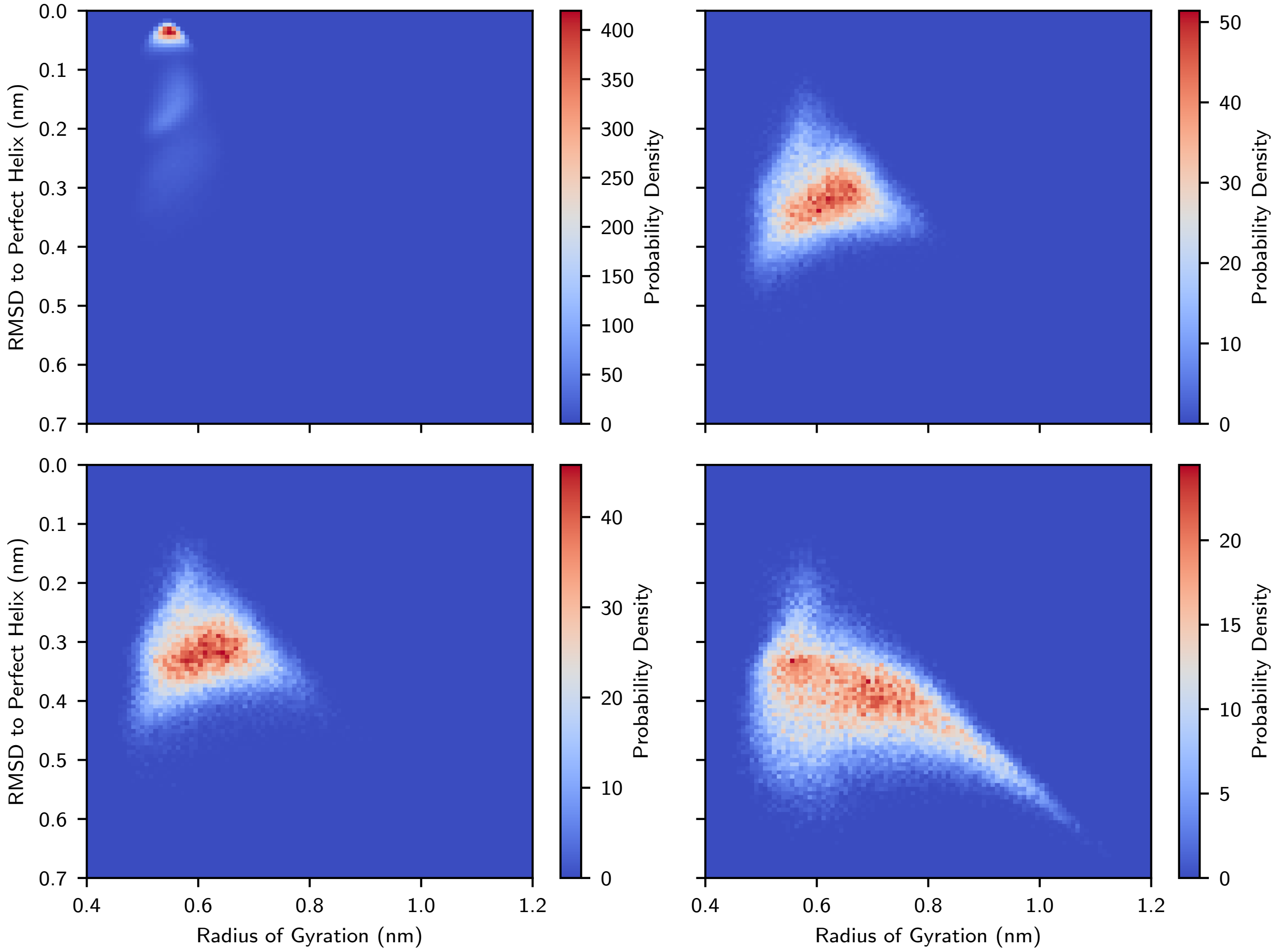


SI Fig. 60. **Probability densities from the four PCCA+ metastable states of A11 in FYAAYF.** The probability density of four metastable states from PCCA+ spectral clustering which demonstrates a state corresponding to a full helix (top left panel), partial helices (top right and bottom left panels), and a coil state (bottom right panel).

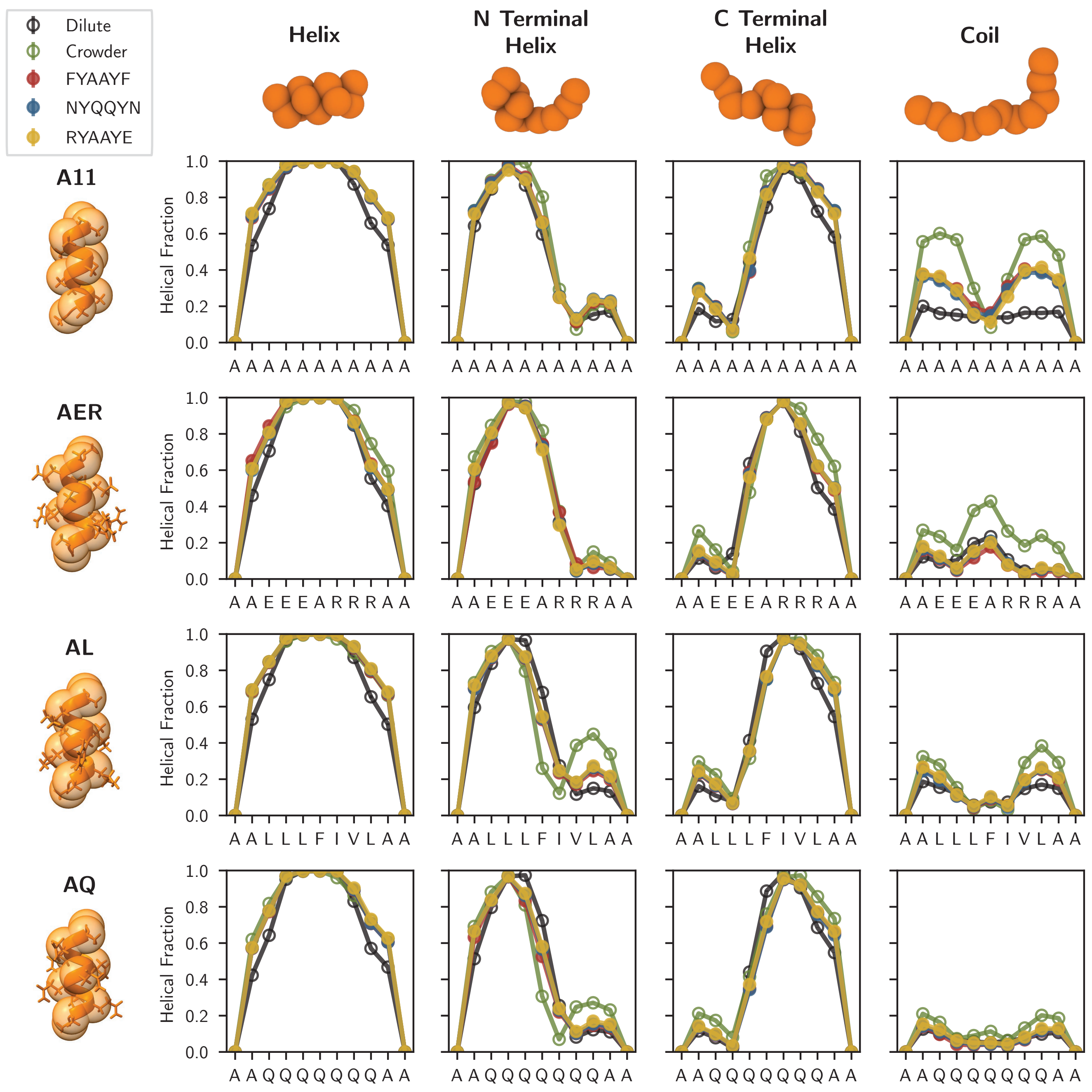


SI Fig. 61. **Helical fractions for helical peptides in each of the four PCCA+ metastable states.** The helical fraction for A11, AER, AL, and AQ in the dilute and crowded solutions as well as the FYAAYF, NYQQYN, and RYAAYE condensates. The left column of plots corresponds to a full helix state. The middle two columns correspond to an N-terminal and C-terminal helix. The rightmost column of plots corresponds to a coil state.

##### E. Additional Results of Helical Peptides in Model Condensates

The free energy surfaces of each helical peptide are plotted within each environment in SI Fig. 62 against the order parameters of radius of gyration and root-mean-squared deviation to a perfect helix. The free energy surfaces depict stark differences between dilute and condensed phases (FYAAYF, NYQQYN, RYAAYE, and crowder) with condensed phases displaying a much more favorable helical state. Additionally, there are variations between peptides, with A11 displaying a relatively similar free energy surface between all the condensed phases whereas the others have free energy surfaces for FYAAYF, NYQQYN, and RYAAYE that fall between the behavior of the dilute and crowded environments. All surfaces are produced from 750 μs of aggregate simulation time for each peptide–environment pairing.

The mean first passage time (MFPT) between all metastable states from the PCCA+ spectral clustering built from our MSMs is shown in SI Fig. 63 for all helical peptides in every environment. Additionally, the unnormalized values for the slowest and second slowest implied timescales from the MSMs are shown in SI Fig. 64.

SI Fig. 65 displays the number of close contacts between each helical peptide and all the types of residues (e.g., alanine included) in the co-condensate peptides of the FYAAYF (left panel), NYQQYN (center panel), and RYAAYE (right panel) condensates. Additionally, we show the full radial distribution functions (RDFs) for every helical peptide in the condensates and crowded solution in SI Figs. 66-69. The higher coordination of the co-condensate peptide residues when the peptide is unfolded, which is seen by the higher first peaks of the RDF in black, is indicative of the enthalpic driving force for protein unfolding in condensates.

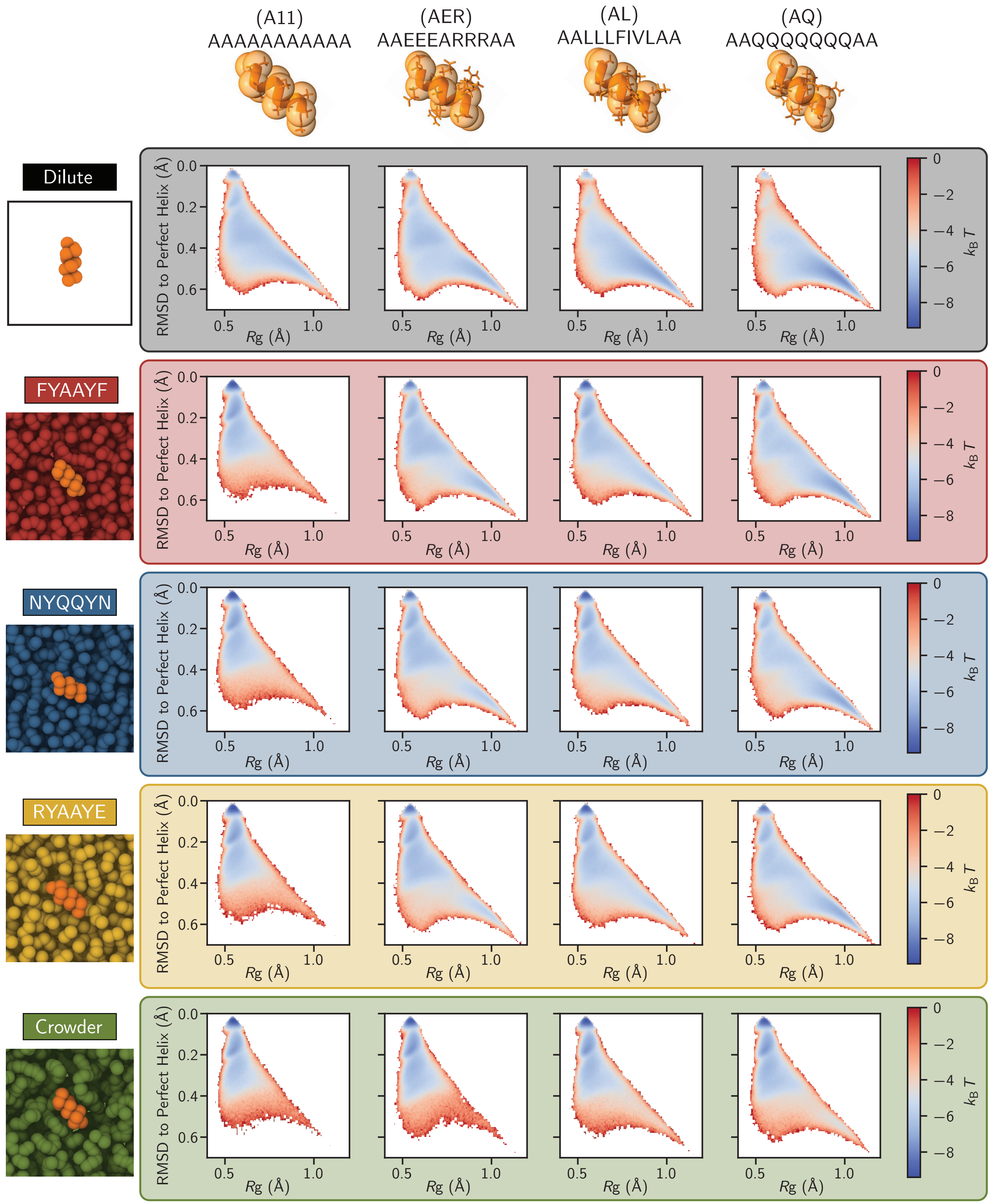


SI Fig. 62. **Free energy surfaces of helical peptides in each environment.** The free energy surfaces for each helical peptide (A11, AER, AL, and AQ) in each chemical environment (dilute, FYAAYF, NYQQYN, RYAAYE, and crowder).

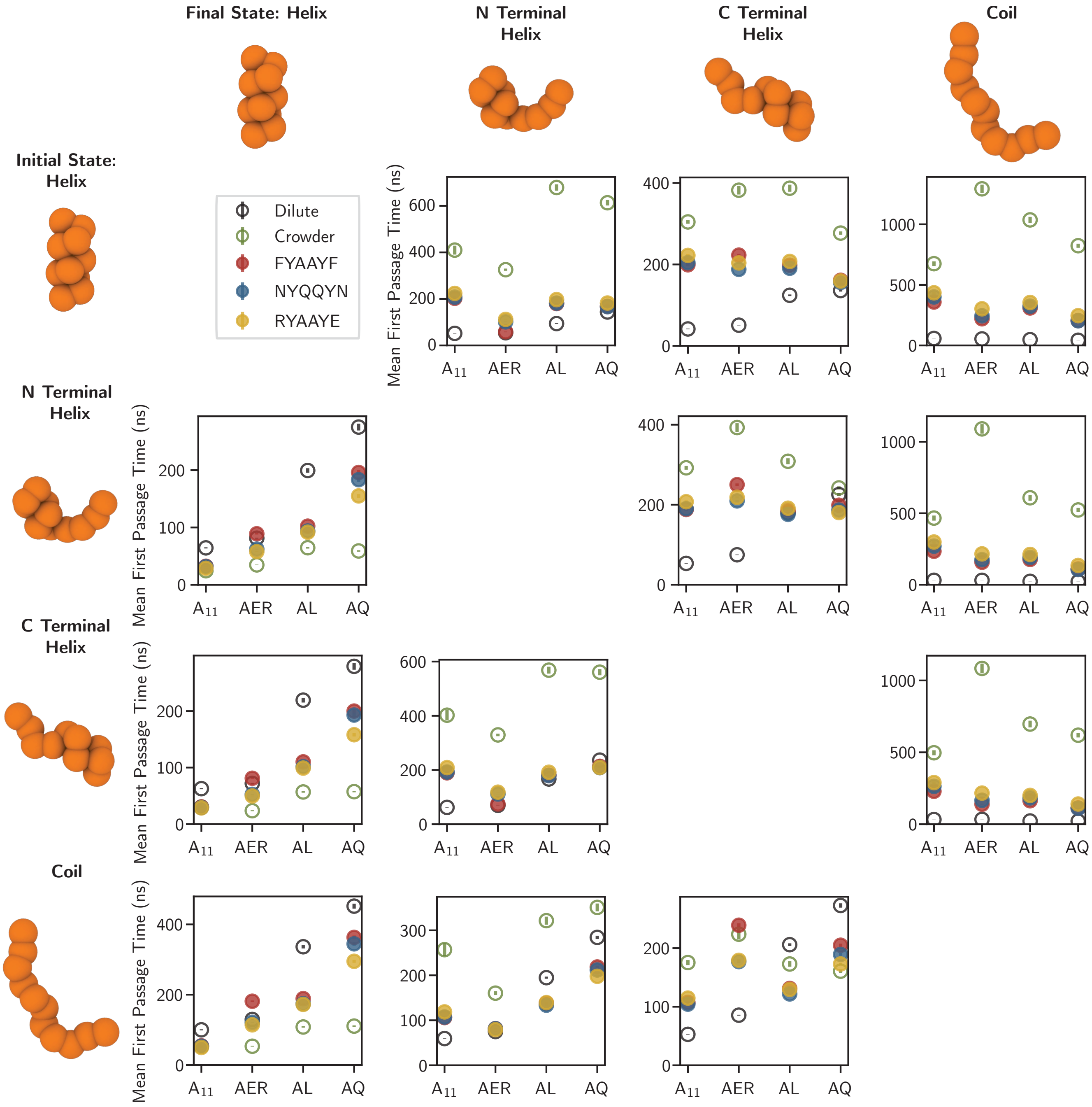


SI Fig. 63. **The mean first passage time for each metastable state transition.** The mean first passage time for transitions among all metastable states (Helix, N-terminal Helix, C-terminal Helix, and Coil) produced via PCCA+ spectral clustering on the MSMs generated for all combinations of helical peptides (A11, AER, AL, and AQ) and environments (dilute, FYAAYF, NYQQYN, RYAAYE, and crowder).

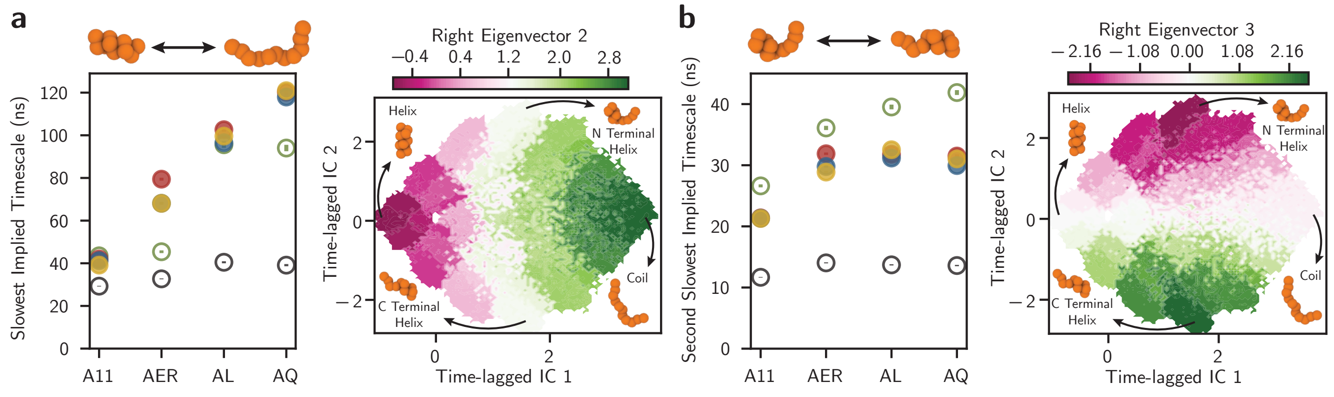


SI Fig. 64. **The implied timescales for the relaxation of the slowest kinetic modes of the $\alpha$-helix folding process. a,** The slowest implied kinetic relaxation timescale (left), corresponding to the transition between the helix and coil states (right). Standard errors are from 100 different instantiations of the MSMs combined with Bayesian uncertainty within the models. The right eigenvector plot is from an MSM applied to the A11–FYAAYF condensate pairing. **b,** The second slowest implied kinetic relaxation timescale (left), corresponding to the transition between the N and C-terminal helix. Errors are calculated and the right eigenvector plot is created as in **a**.

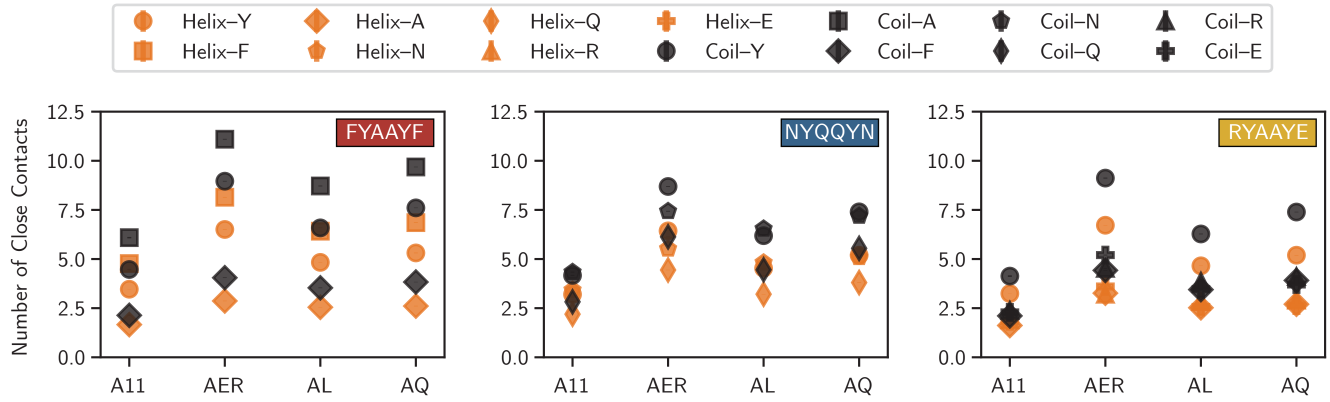


SI Fig. 65. **Close contact measurements between helical peptides and co-condensate peptide residues.** Average number of close contacts between the central 7 residues of each helical peptide (A11, AER, AL, and AQ) and co-condensate peptide residues in the FYAAYF (left), NYQQYN (center), and RYAAYE (right) condensates.

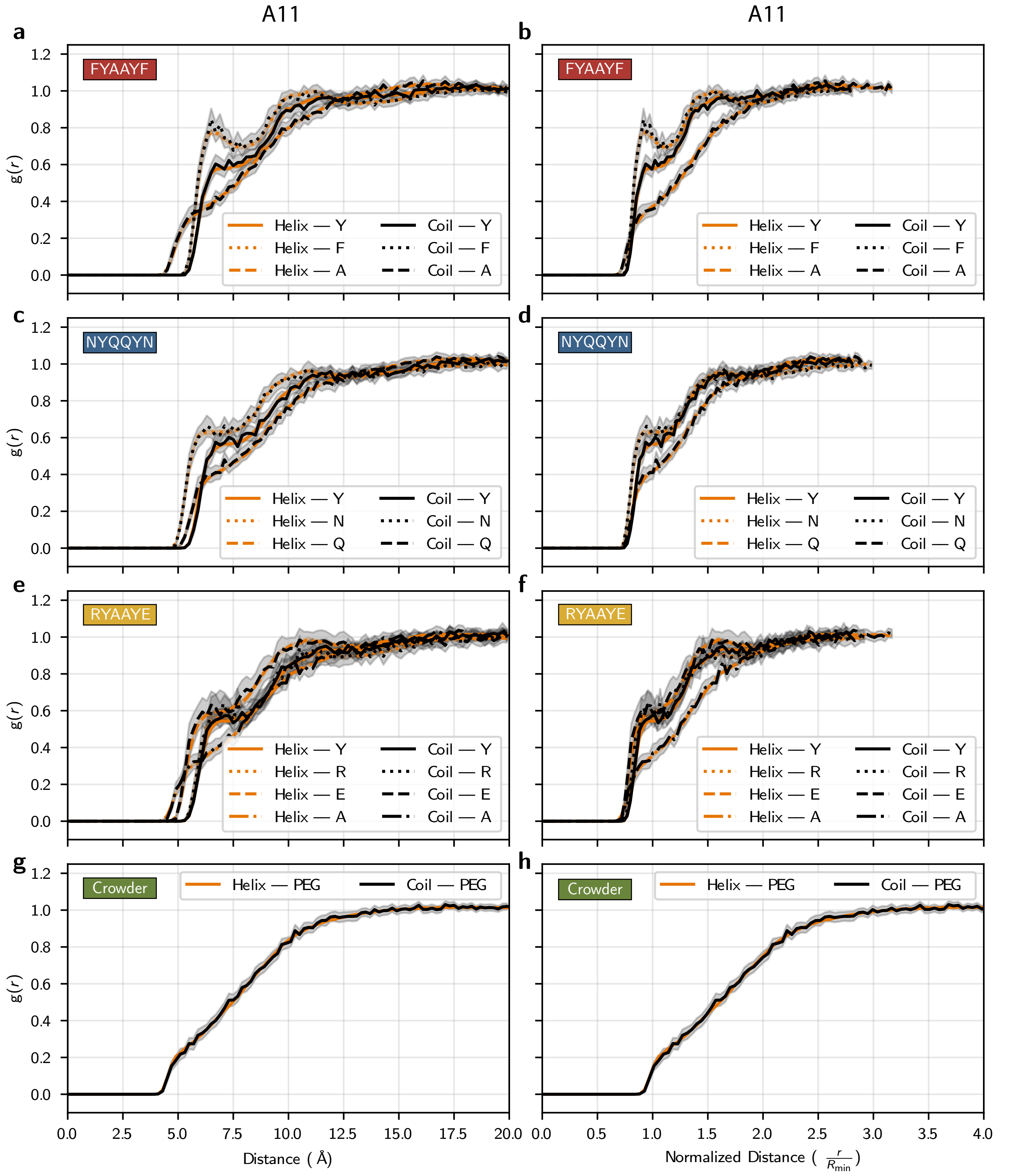


SI Fig. 66. **RDFs and distance normalized RDFs for A11 in every condensed chemical environment (FYAAYF, NYQQYN, RYAAYE, and Crowder).**

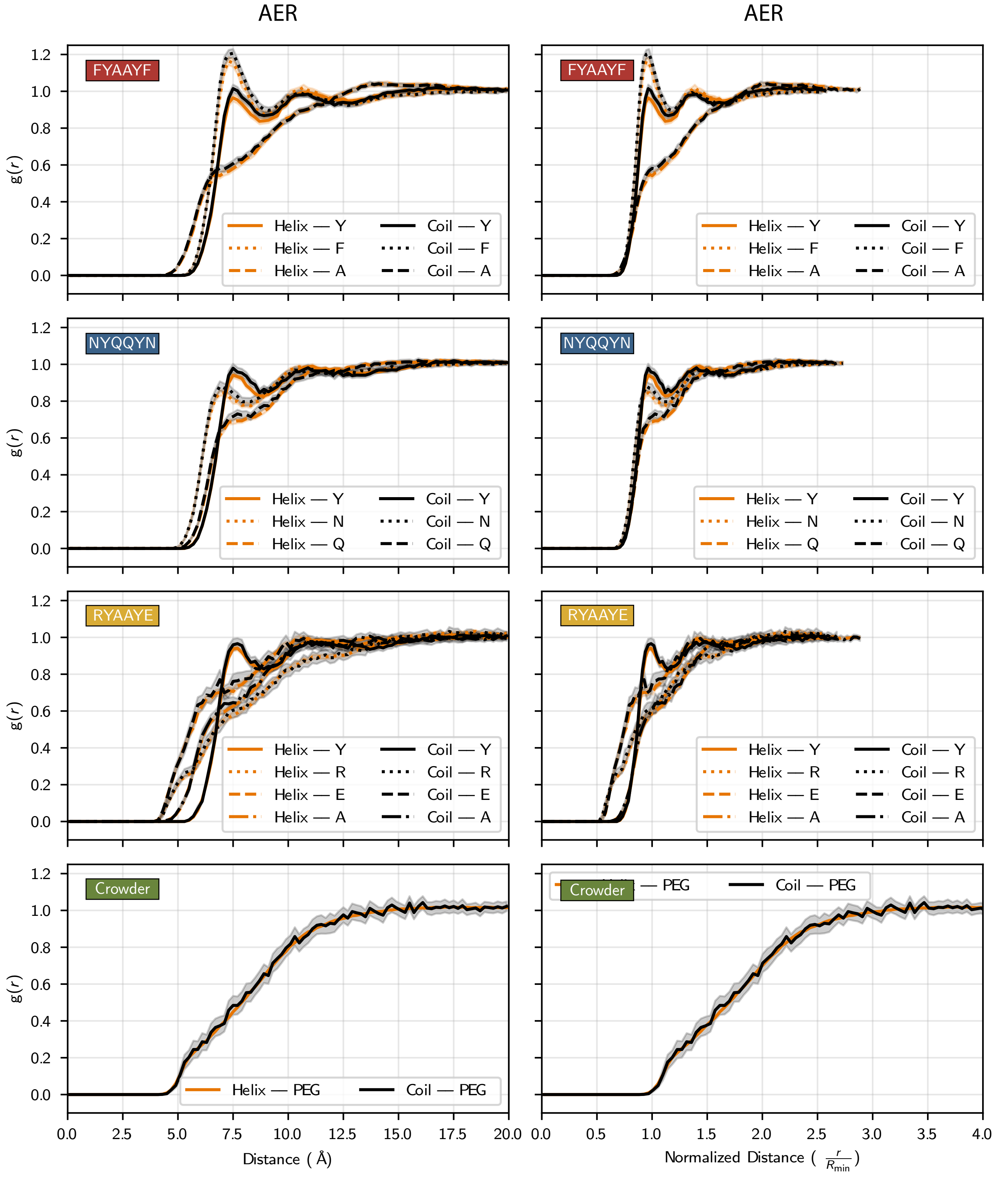


SI Fig. 67. **RDFs and distance normalized RDFs for AER in every condensed chemical environment (FYAAYF, NYQQYN, RYAAYE, and Crowder).**

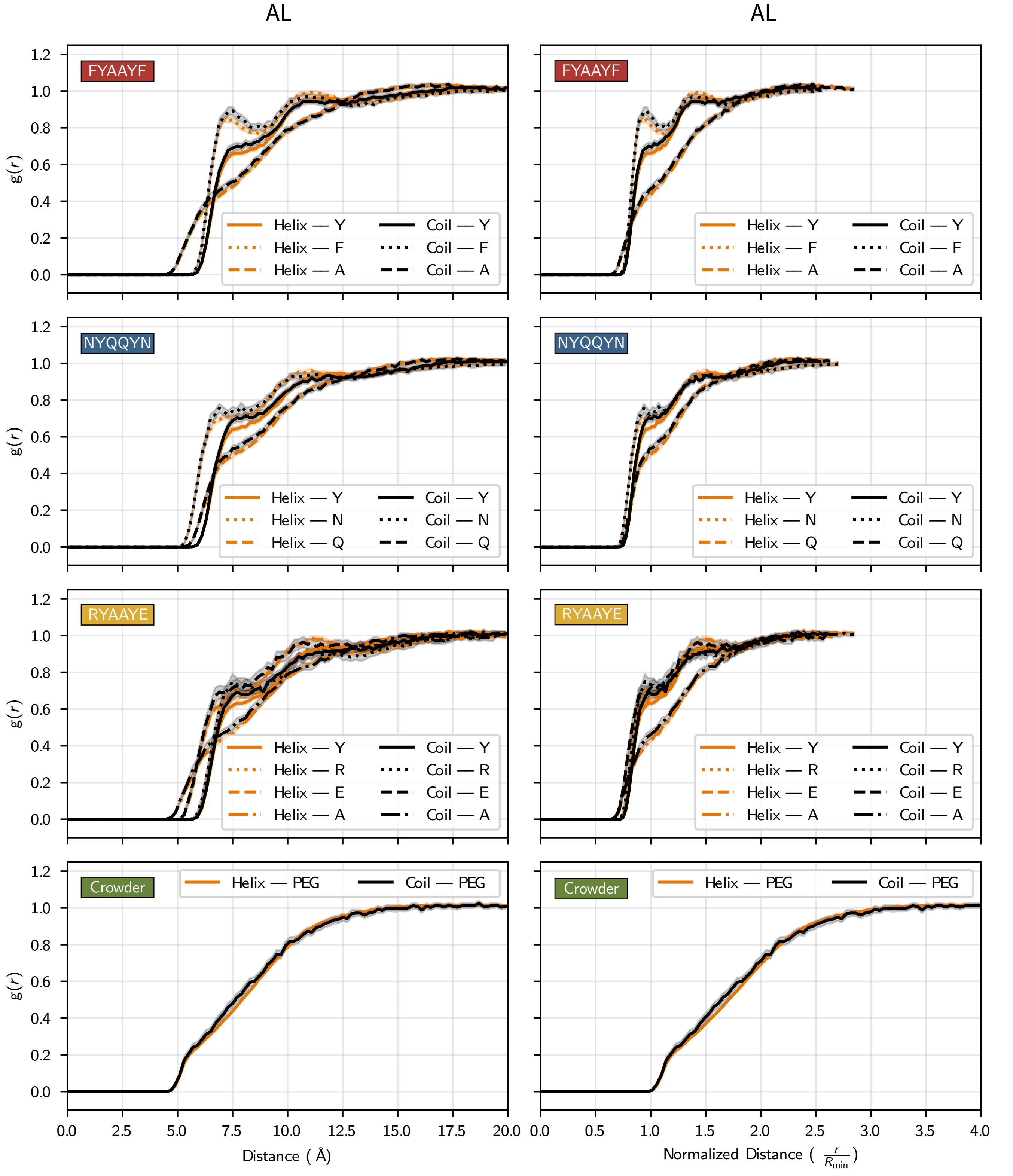


SI Fig. 68. **RDFs and distance normalized RDFs for AL in every condensed chemical environment (FYAAYF, NYQQYN, RYAAYE, and Crowder).**

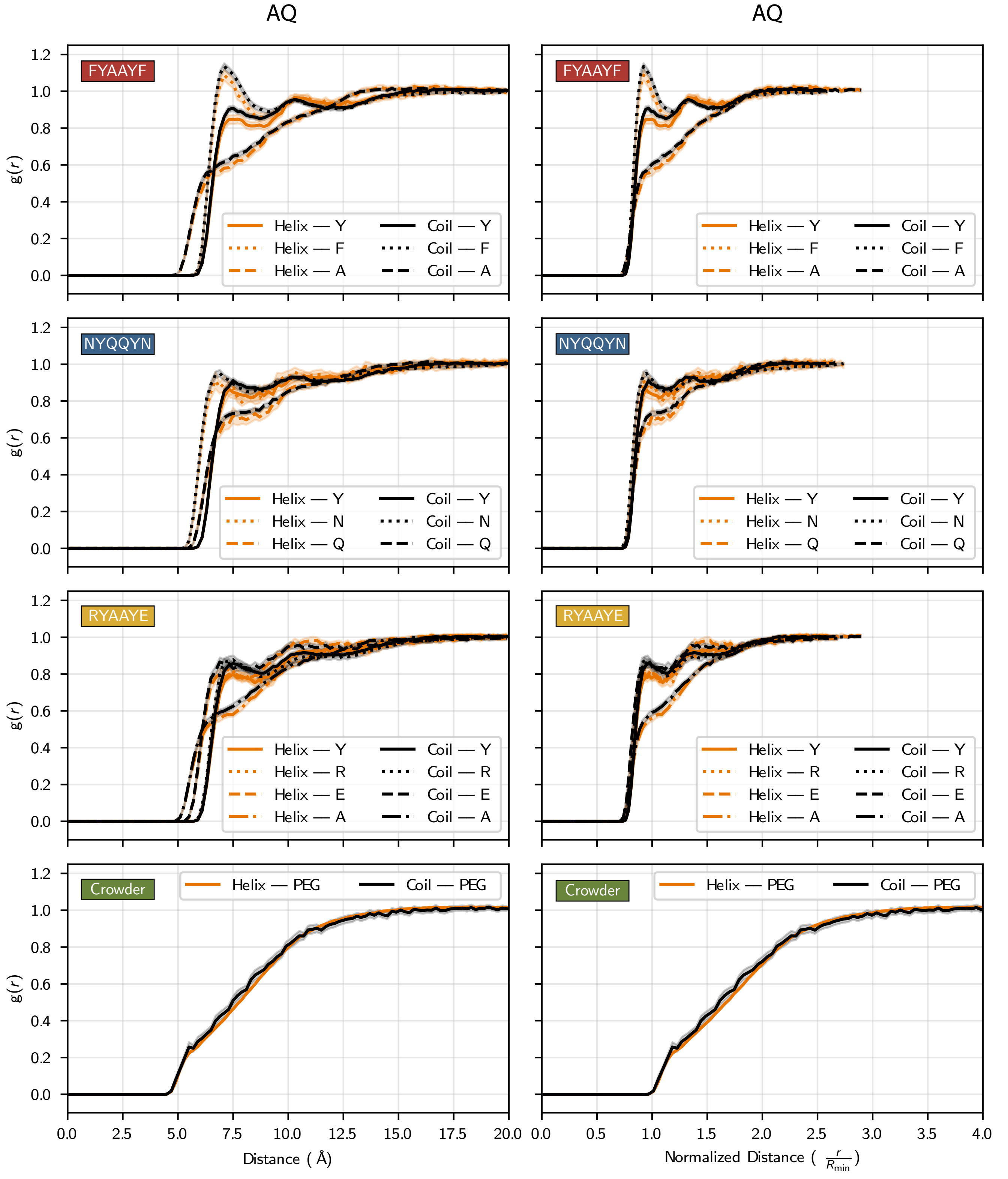


SI Fig. 69. **RDFs and distance normalized RDFs for AQ in every condensed chemical environment (FYAAYF, NYQQYN, RYAAYE, and Crowder).**

### V. IDP HELICAL DOMAINS IN CONDENSATES

#### A. Co-Condensate IDP Densities and Pressures

In order to simulate TDP43, ANXA11, and AR in condensates, we needed to ensure that the co-condensate proteins were able to phase separate on their own. To test this, we simulated FUS, hnRNPA1, TIA1, RPB1, MED1, and BRD4 using direct coexistence simulations in a slab geometry (see Methods for details). As shown by the density profiles in SI Fig. 70, FUS, hnRNPA1, and TIA1 are able to phase separate at 298.15 K on their own. However, RPB1, MED1, and BRD4 all require crowder to phase separate, as has been demonstrated experimentally [55, 56]. For these systems, we were able to phase separate these proteins with 10% PEG-8000 (e.g., PEG chains of 176 monomers in length).

We next determined the pressure in the *NPT* ensemble that would recapitulate the densities of the RPB1, MED1, and BRD4 condensates from the direct coexistence simulations (SI Fig. 71, Methods).

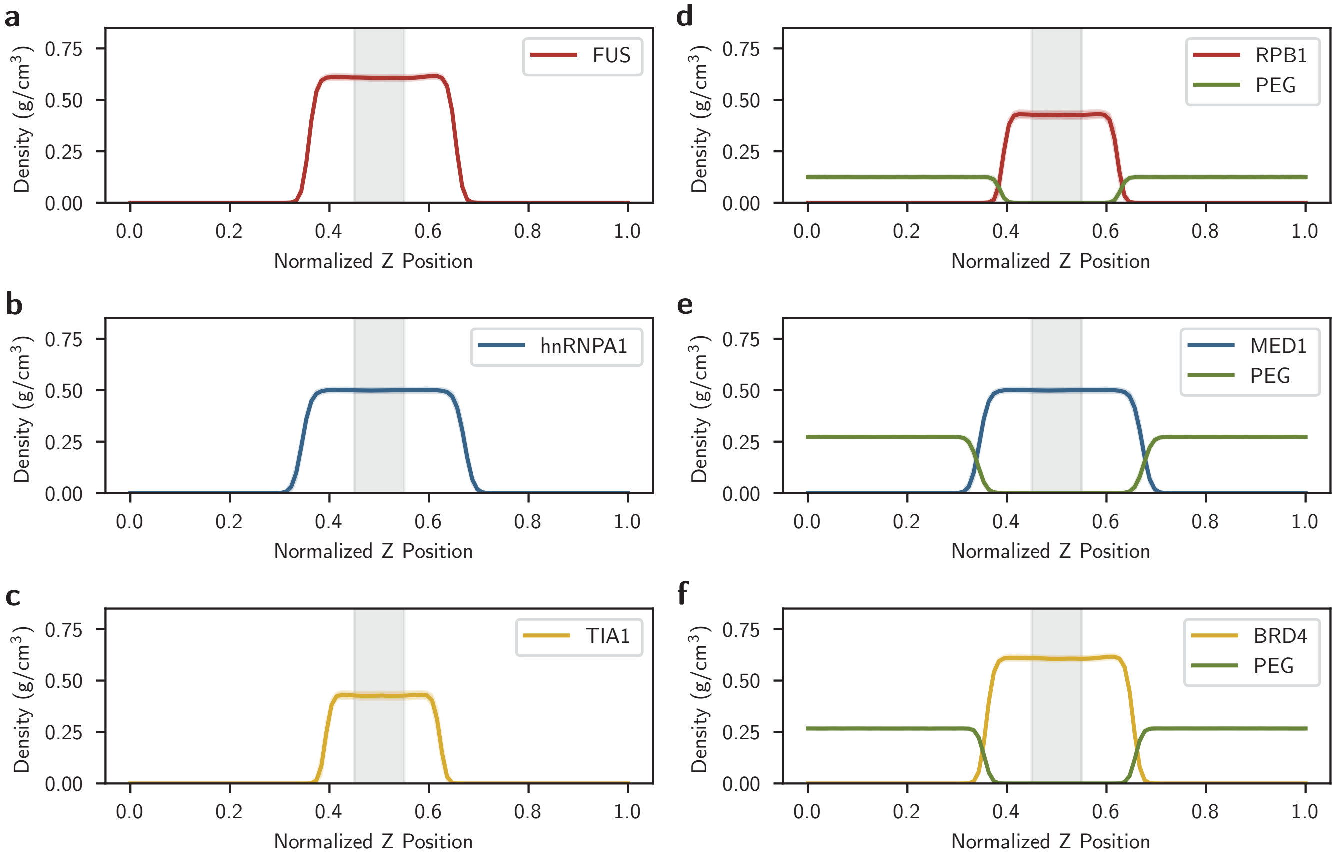


SI Fig. 70. **Density profiles from direct coexistence simulations of co-condensate proteins.** The density profiles from direct coexistence simulations in the slab geometry using the Mpipi-Helix force field for **a,** FUS, **b,** hnRNPA1, **c,** TIA1, **d,** RPB1, **e,** MED1, and **f,** BRD4. The simulations in **d**-**f** were run with 10% PEG-8000. Simulations were performed for 1.25 µs with the first 250 ns discarded as equilibration. Error is displayed as the standard deviation across the sampling period of 1 µs.

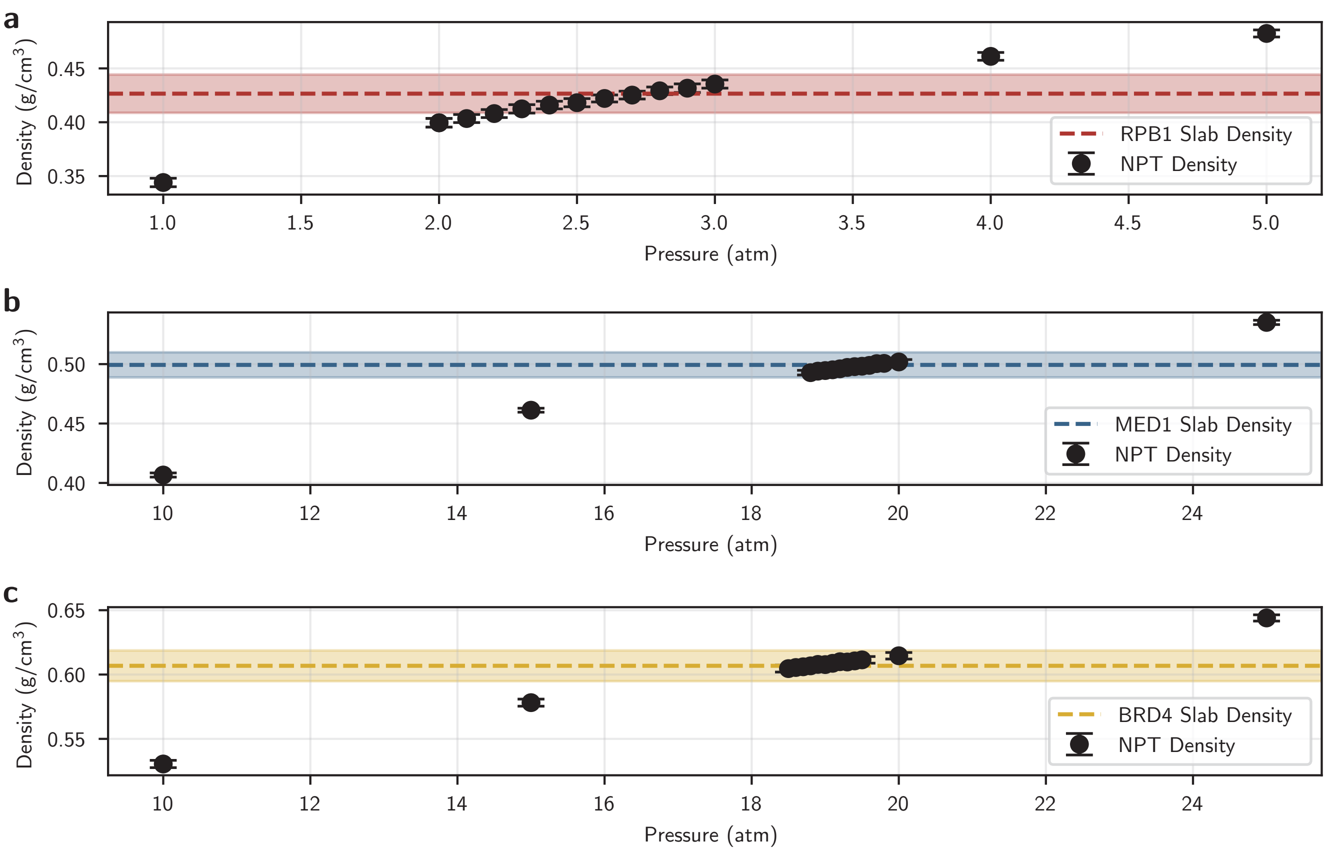


SI Fig. 71. **Pressures in $NPT$ ensemble that recapitulate direct coexistence densities of co-condensate proteins.** The results (black circles) of $NPT$ simulations which were performed to determine the pressure that would recapitulate the density of the co-condensate proteins (**a,** RPB1, **b,** MED1, and **c,** BRD4) from the direct coexistence simulations with 10% PEG. Simulations were performed in the $NPT$ ensemble (using a Langevin thermostat at 298.15 K with a 100 ps damping constant and a Berendsen barostat with a 100 ps damping constant) for 200 ns with the first 100 ns removed for equilibration. The simulation density was recorded every 0.5 ns. Error is plotted as the standard deviation across the sampling period.

##### B. MSM Validation

We built MSMs for the helical domains in dilute solutions and condensates using the pyEMMA toolkit [53]. A key assumption for these MSMs is that the degrees of freedom associated with the co-condensate proteins relax faster than the lag time of our MSMs, thus allowing us to not include the co-condensate proteins in our model construction. As such, the effects of the co-condensate proteins are implicitly incorporated into the MSM through their impact on the degrees of freedom of the helical domains.

A parameter scan was used to determine the hyperparameters (i.e., the input features, lag time, and number of clusters) for our MSMs (SI Figs. 72, 73, and 74) in an identical fashion as the helical peptides (see 'Markov State Models and PCCA+ Spectral Clustering' section above). From this analysis, we determined that it was reasonable to use input features of bond angles, dihedral angles, and non-bonded pairwise distances between residues in the helical domain, because this combination of features consistently had the highest VAMP2 score. We chose 1500 cluster centers for model construction as this is the largest number of clusters before the MSMs show dips in VAMP2 score or high errors (e.g., TDP43 + TIA1 in SI Fig. 72). Finally, we chose a lag time of 5 ns because the implied timescales change only modestly for longer lag times. Our choices are validated by the Chapman–Kolmogorov test (SI Figs. 75, 76, and 77). Here, the agreement between the MSM prediction and the estimate from the MD data suggests that the models can successfully describe the folding landscape of our helical domains in each environment.

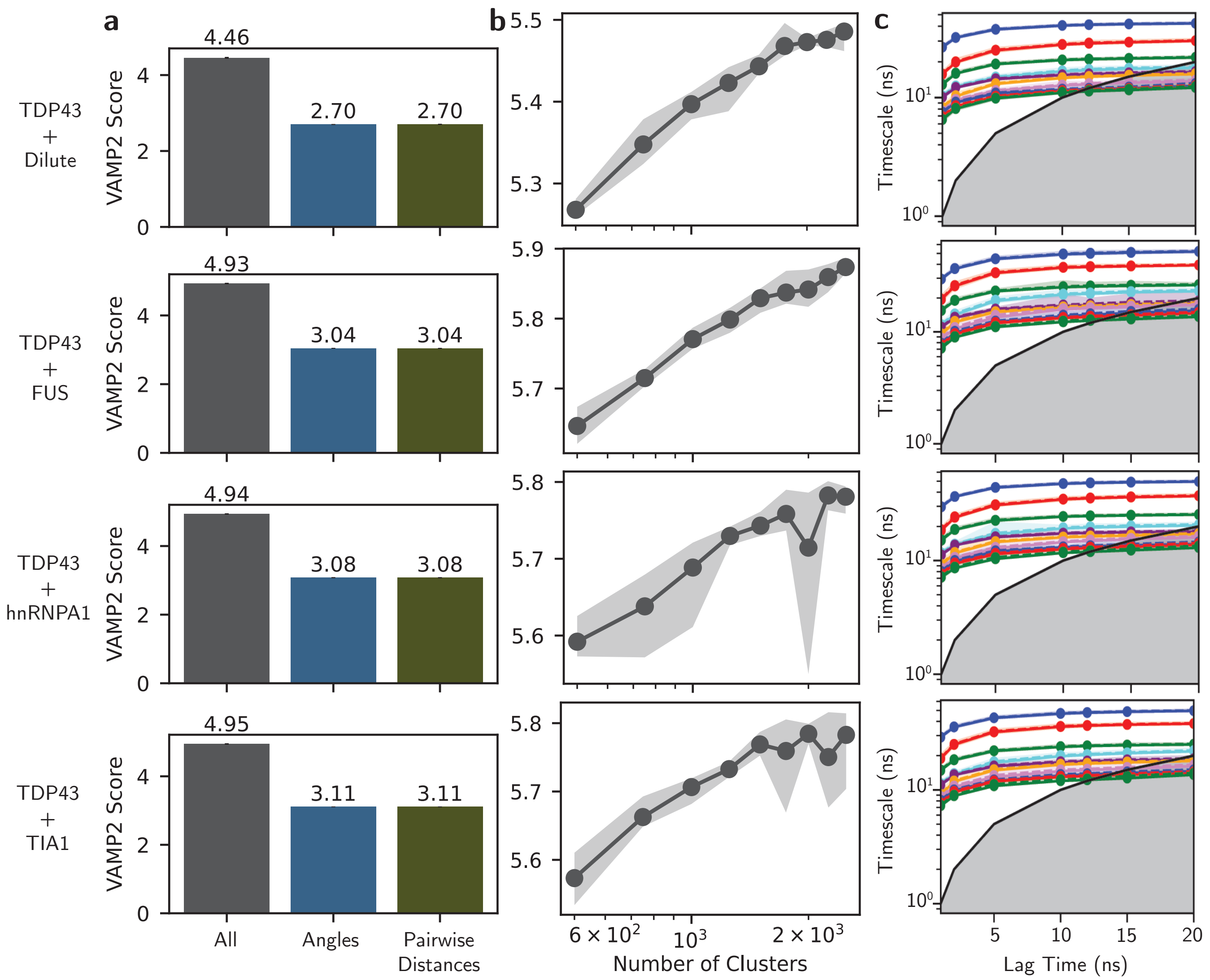


SI Fig. 72. **MSM hyperparameter validation for TDP43 in a dilute solution and condensates.** **a,** Validation of input features for MSMs was performed using VAMP2 analysis on input bond/dihedral angles (Angles), all non-bonded pairwise distances (Pairwise Distances), and both feature sets together (All). **b,** Validation of the number of cluster centers for kmeans++ clustering was performed using VAMP2 and a lag time of 5 ns (y-axis is VAMP2 score). **c,** Validation of the lag time was performed using an implied timescale analysis. From this analysis, we chose to use all input features paired with time-lagged independent component analysis with 1500 cluster centers at a lag time of 5 ns as the final MSM hyperparameters.

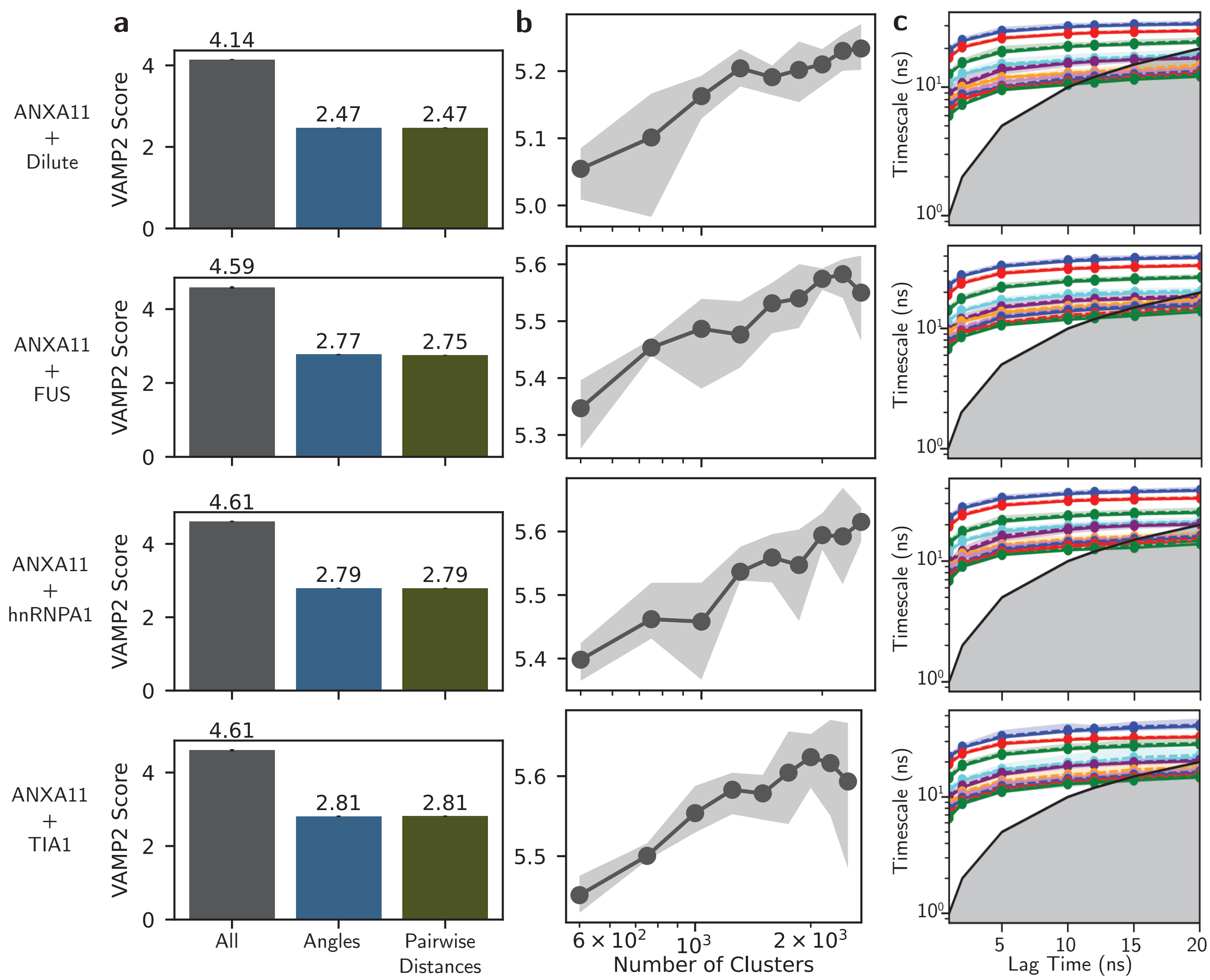


SI Fig. 73. **MSM hyperparameter validation for ANXA11 in a dilute solution and condensates.** **a,** Validation of input features for MSMs was performed using VAMP2 analysis on input bond/dihedral angles (Angles), all non-bonded pairwise distances (Pairwise Distances), and both feature sets together (All). **b,** Validation of the number of cluster centers for kmeans++ clustering was performed using VAMP2 and a lag time of 5 ns (y-axis is VAMP2 score). **c,** Validation of the lag time was performed using an implied timescale analysis. From this analysis, we chose to use all input features paired with time-lagged independent component analysis with 1500 cluster centers at a lag time of 5 ns as the final MSM hyperparameters.

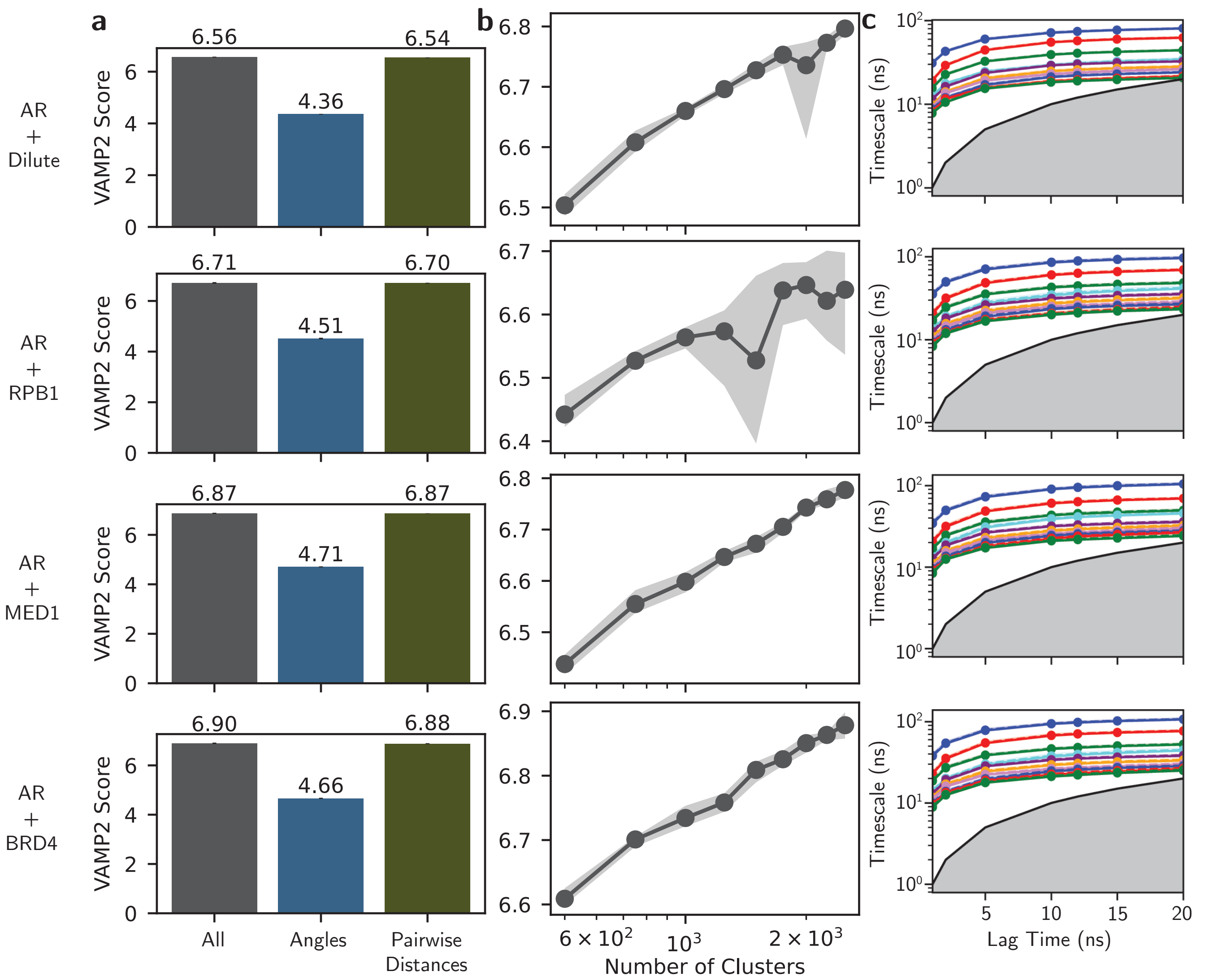


SI Fig. 74. **MSM hyperparameter validation for AR in a dilute solution and condensates. a,** Validation of input features for MSMs was performed using VAMP2 analysis on input bond/dihedral angles (Angles), all non-bonded pairwise distances (Pairwise Distances), and both feature sets together (All). **b,** Validation of the number of cluster centers for kmeans++ clustering was performed using VAMP2 and a lag time of 5 ns (y-axis is VAMP2 score). **c,** Validation of the lag time was performed using an implied timescale analysis. From this analysis, we chose to use all input features paired with time-lagged independent component analysis with 1500 cluster centers at a lag time of 5 ns as the final MSM hyperparameters.

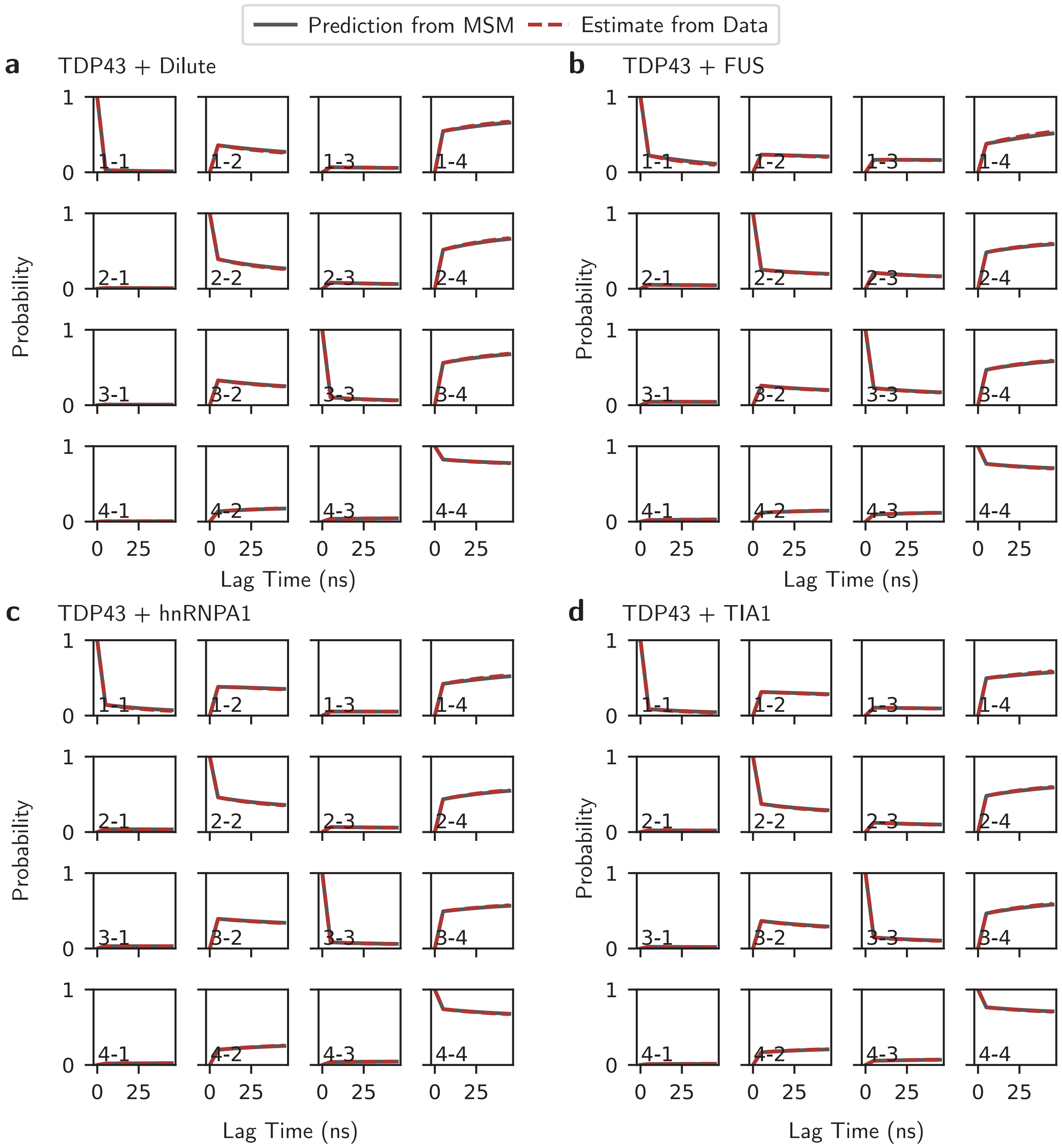


SI Fig. 75. **Chapman–Kolmogorov test for TDP43.** Validation of MSM construction using the Chapman–Kolmogorov test for TDP43 in **a,** dilute solution, **b,** FUS condensate, **c,** hnRNPA1 condensate, and **d,** TIA1 condensate.

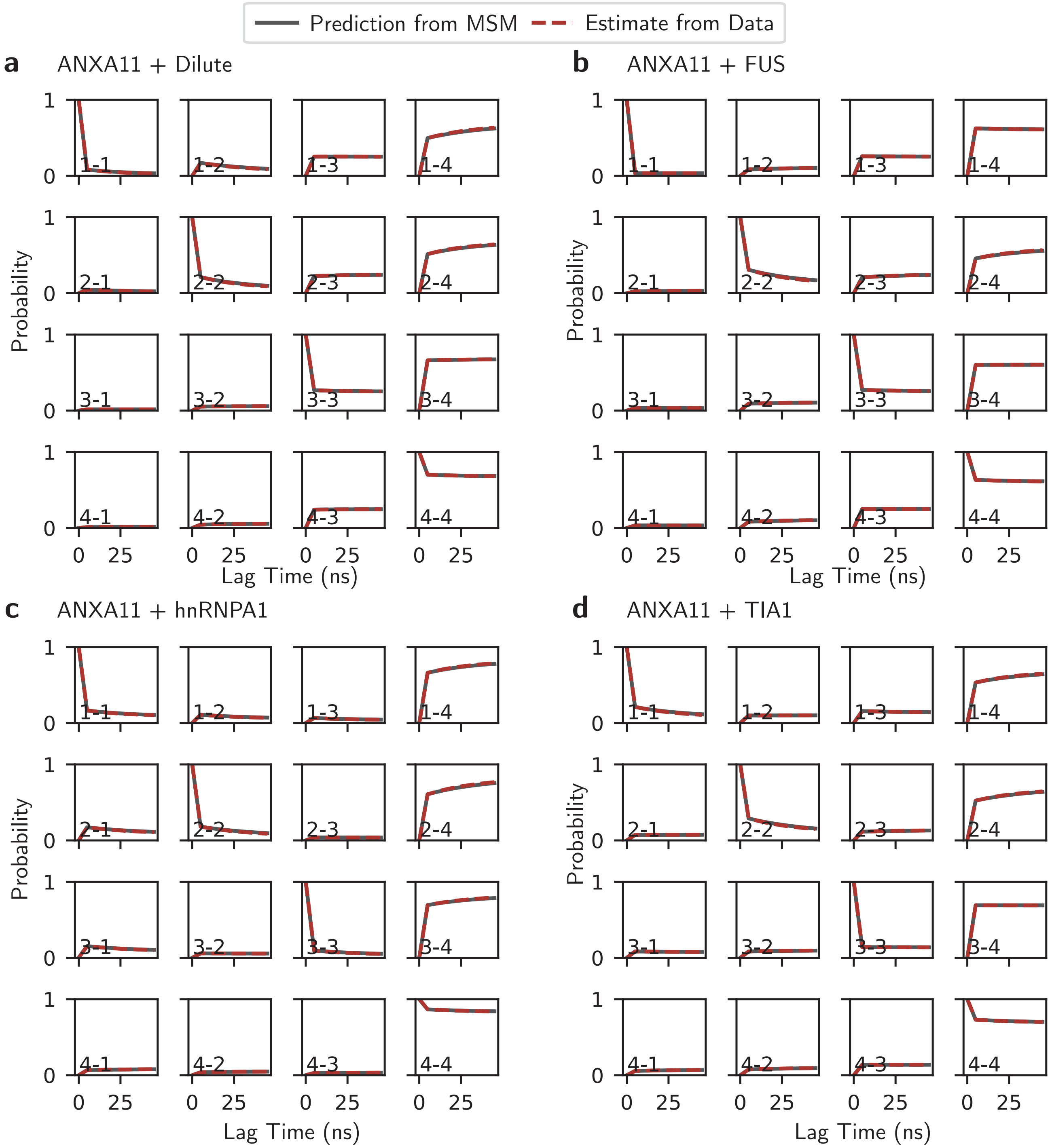


SI Fig. 76. **Chapman–Kolmogorov test for ANXA11.** Validation of MSM construction using the Chapman–Kolmogorov test for ANXA11 in **a,** dilute solution, **b,** FUS condensate, **c,** hnRNPA1 condensate, and **d,** TIA1 condensate.

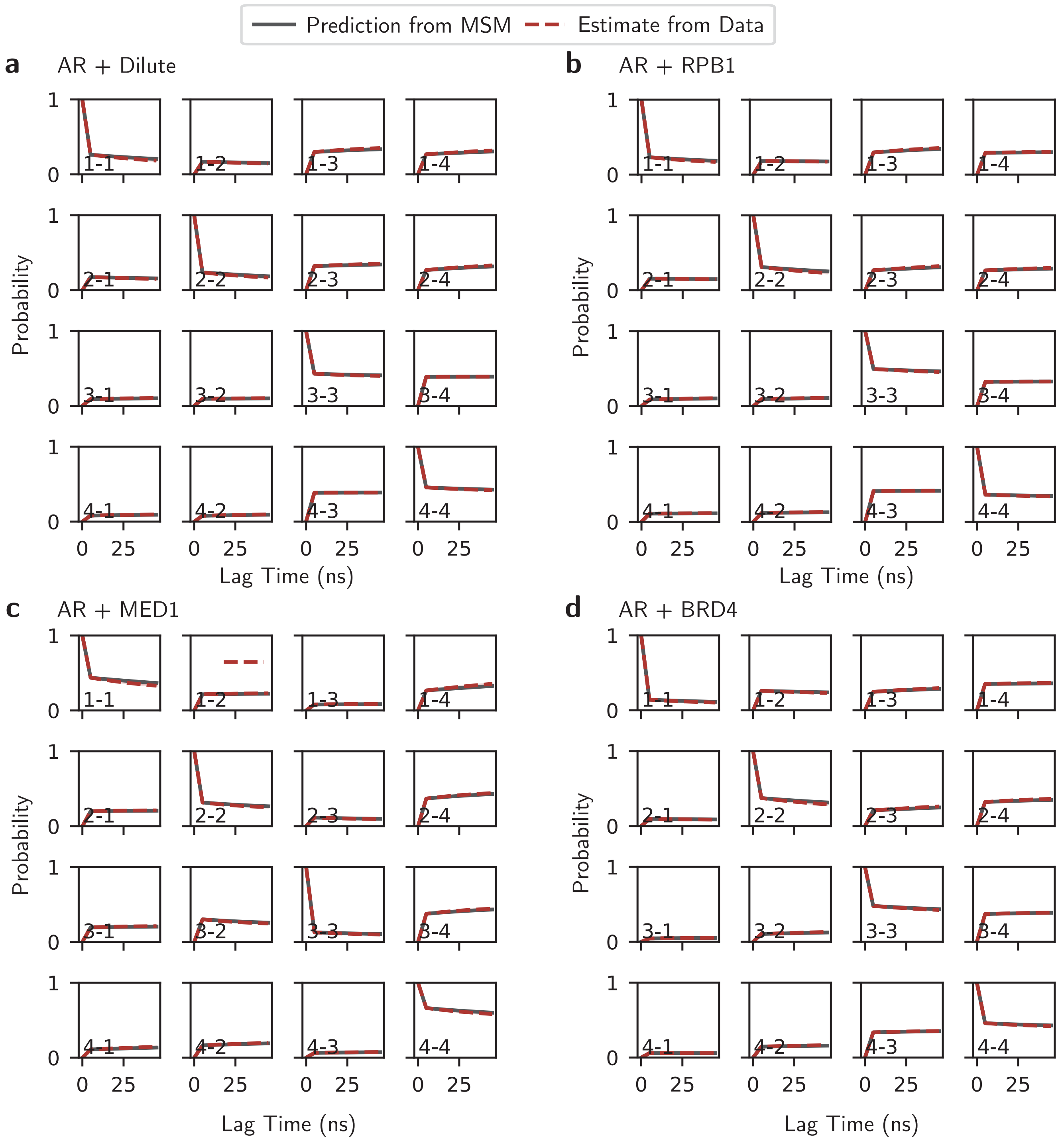


SI Fig. 77. **Chapman–Kolmogorov test for AR.** Validation of MSM construction using the Chapman–Kolmogorov test for AR in **a,** dilute solution, **b,** RPB1 condensate, **c,** MED1 condensate, and **d,** BRD4 condensate.

##### C. Additional Results for IDP Helical Domains in Condensates

SI Fig. 78 plots the unnormalized values of the slowest implied relaxation timescale for each helical domain within the condensates and dilute solution. SI Fig. 79 depicts the contacts per residue between the helical domains and the co-condensate proteins for all residue type categories. Similarly, SI Fig. 80 shows the residue contact probability differences between the helical domains and the co-condensate proteins for all 20 canonical amino acids. See Methods for details on these calculations.

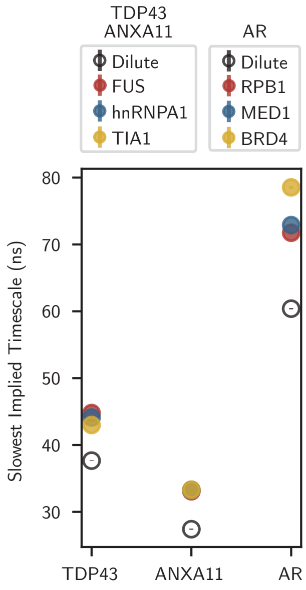


SI Fig. 78. **The implied timescales for the relaxation of the slowest kinetic mode of the $\alpha$-helix domains.** The slowest implied kinetic relaxation timescale for the $\alpha$-helix domains in each environment. This timescale corresponds to a folding transition (see Extended Data Fig. 5c). Standard error is calculated across 25 different instantiations of our MSMs combined with combined with Bayesian uncertainty within the models.

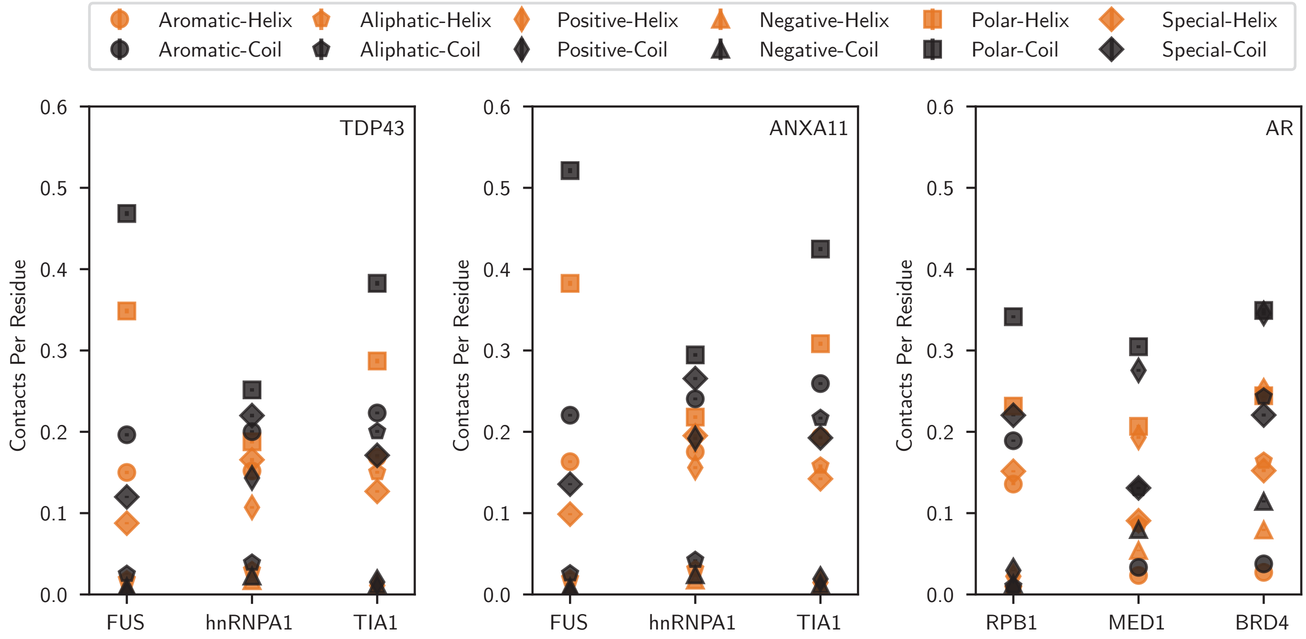


SI Fig. 79. **Close contact measurements between TDP43, ANXA11, and AR and co-condensate proteins.** Per-residue close contact numbers between TDP43 (left), ANXA11 (middle), and AR (right) with each co-condensate protein for every residue category type. Contacts made with residues in the helical state within TDP43, ANXA11, and AR are shown in orange and contacts with residues in the coil state are shown in black.

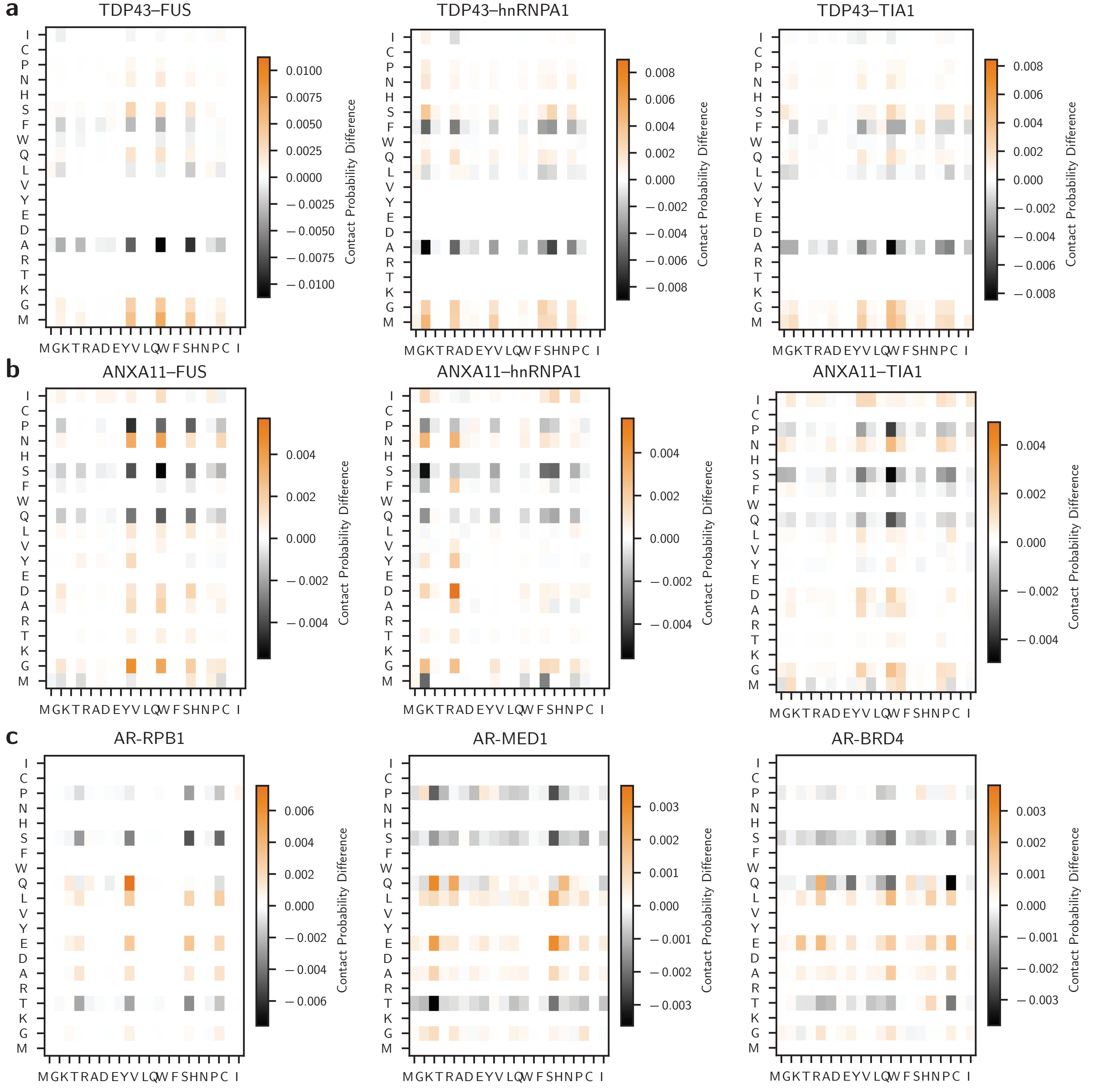


SI Fig. 80. **Full residue contact probability differences for TDP43, ANXA11, and AR in each condensate.** The residue contact probability differences for **a**, TDP43, **b**, ANXA11, and **c**, AR in each condensate.

#### D. Thermodynamic and Kinetic Data on the Additional Helices in AR

The AR protein contains five stretches of helical residues (see BMRB 51479 and 51480 as well as [46]). The first two of helical segments are very short (aa 1-9 and aa 22-32). The next helical segment is what we focus on in the main text and it is the longest polyglutamine region of the AR protein (aa 50-81). We call this region polyQ1. The other two helical segments are a shorter polyglutamine segment (aa 181-198), which we call polyQ2, and the helix within the Tau5 segment (aa 390-414). For these two helical domains, we calculate the helical fraction, slowest implied timescale, close contact numbers in folded and unfolded states on a per residue basis, and the residue type contact probability differences (SI Fig. 81). From these calculations, we see qualitatively similar behavior to the polyQ1 helix in AR, but as expected the exact details are dually informed by the folded domain and co-condensate protein sequences. Please note that in all depictions of the helical domains (SI Fig. 81 and Main Text Fig. 4c), we include an additional 3 residues on either side of the sequence as the Mpipi-Helix helical potential includes those residues (to provide full coverage of helical potentials on the residues that are helical in the NMR chemical shifts—see Methods).

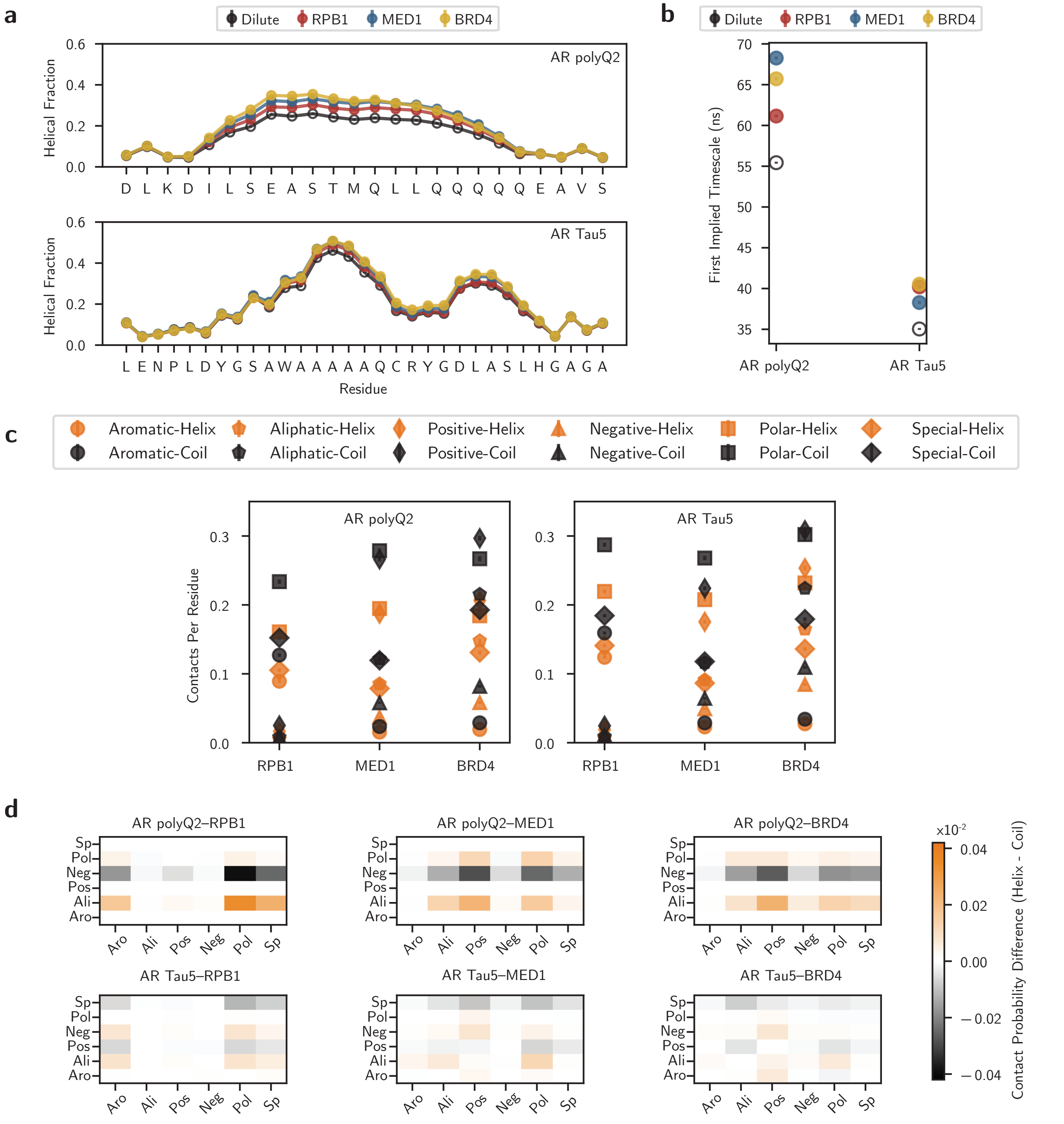


SI Fig. 81. **Thermodynamics and kinetics of the polyQ2 and Tau5 helices in AR.** The **a,** helical fractions, **b,** first (slowest) implied timescale, **c,** close contacts per residue, and **d,** residue type contact probability differences for the polyQ2 helix (aa 181-198) and Tau5 helix (aa 390-414). All calculations and errors are reported as in Main Text Fig. 4.

[1] C. Tian, K. Kasavajhala, K. A. A. Belfon, L. Raguette, H. Huang, A. N. Migues, J. Bickel, Y. Wang, J. Pincay, Q. Wu, and C. Simmerling, ff19SB: Amino-Acid-Specific Protein Backbone Parameters Trained against Quantum Mechanics Energy Surfaces in Solution, Journal of Chemical Theory and Computation **16**, 528 (2020).
[2] P. Robustelli, S. Piana, and D. E. Shaw, Developing a molecular dynamics force field for both folded and disordered protein states, Proceedings of the National Academy of Sciences **115**, E4758 (2018).
[3] J. Zhu, X. Salvatella, and P. Robustelli, Small molecules targeting the disordered transactivation domain of the androgen receptor induce the formation of collapsed helical states, Nature Communications **13**, 6390 (2022).
[4] A. Barducci, G. Bussi, and M. Parrinello, Well-Tempered Metadynamics: A Smoothly Converging and Tunable Free-Energy Method, Physical Review Letters **100**, 020603 (2008).
[5] X. Wang, S. Ramírez-Hinestrosa, J. Dobnikar, and D. Frenkel, The Lennard-Jones potential: When (not) to use it, Physical Chemistry Chemical Physics **22**, 10624 (2020).
[6] A. P. Thompson, H. M. Aktulga, R. Berger, D. S. Bolintineanu, W. M. Brown, P. S. Crozier, P. J. in 't Veld, A. Kohlmeyer, S. G. Moore, T. D. Nguyen, R. Shan, M. J. Stevens, J. Tranchida, C. Trott, and S. J. Plimpton, LAMMPS - a flexible simulation tool for particle-based materials modeling at the atomic, meso, and continuum scales, Computer Physics Communications **271**, 108171 (2022).
[7] J. A. Joseph, A. Reinhardt, A. Aguirre, P. Y. Chew, K. O. Russell, J. R. Espinosa, A. Garaizar, and R. Collepardo-Guevara, Physics-driven coarse-grained model for biomolecular phase separation with near-quantitative accuracy, Nature Computational Science **1**, 732 (2021).
[8] G. Wang and R. L. Dunbrack, PISCES: A protein sequence culling server, Bioinformatics **19**, 1589 (2003).
[9] W. Kabsch and C. Sander, Dictionary of protein secondary structure: Pattern recognition of hydrogen-bonded and geometrical features, Biopolymers **22**, 2577 (1983).
[10] A. Rizuan, N. Jovic, T. M. Phan, Y. C. Kim, and J. Mittal, Developing Bonded Potentials for a Coarse-Grained Model of Intrinsically Disordered Proteins, Journal of Chemical Information and Modeling **62**, 4474 (2022).
[11] Z. Hu, T. Sun, W. Chen, L. Nordenskiöld, and L. Lu, Refined Bonded Terms in Coarse-Grained Models for Intrinsically Disordered Proteins Improve Backbone Conformations, The Journal of Physical Chemistry B **128**, 6492 (2024).
[12] S. Cocco, C. Feinauer, M. Figliuzzi, R. Monasson, and M. Weigt, Inverse statistical physics of protein sequences: A key issues review, Reports on Progress in Physics **81**, 032601 (2018).
[13] M. Ekeberg, C. Lövkvist, Y. Lan, M. Weigt, and E. Aurell, Improved contact prediction in proteins: Using pseudolikelihoods to infer Potts models, Physical Review E **87**, 012707 (2013).
[14] J. Jumper, R. Evans, A. Pritzel, T. Green, M. Figurnov, O. Ronneberger, K. Tunyasuvunakool, R. Bates, A. Žídek, A. Potapenko, A. Bridgland, C. Meyer, S. A. A. Kohl, A. J. Ballard, A. Cowie, B. Romera-Paredes, S. Nikolov, R. Jain, J. Adler, T. Back, S. Petersen, D. Reiman, E. Clancy, M. Zielinski, M. Steinegger, M. Pacholska, T. Berghammer, S. Bodenstein, D. Silver, O. Vinyals, A. W. Senior, K. Kavukcuoglu, P. Kohli, and D. Hassabis, Highly accurate protein structure prediction with AlphaFold, Nature **596**, 583 (2021).
[15] The UniProt Consortium, UniProt: The Universal Protein Knowledgebase in 2025, Nucleic Acids Research **53**, D609 (2025).
[16] M. Steinegger and J. Söding, MMseqs2 enables sensitive protein sequence searching for the analysis of massive data sets, Nature Biotechnology **35**, 1026 (2017).
[17] T. P. Creamer and G. D. Rose, Side-chain entropy opposes alpha-helix formation but rationalizes experimentally determined helix-forming propensities., Proceedings of the National Academy of Sciences **89**, 5937 (1992).
[18] R. J. Moreau, C. R. Schubert, K. A. Nasr, M. Török, J. S. Miller, R. J. Kennedy, and D. S. Kemp, Context-Independent, Temperature-Dependent Helical Propensities for Amino Acid Residues, Journal of the American Chemical Society **131**, 13107 (2009).
[19] L. Wang, R. A. Friesner, and B. J. Berne, Replica Exchange with Solute Scaling: A More Efficient Version of Replica Exchange with Solute Tempering (REST2), The Journal of Physical Chemistry B **115**, 9431 (2011).
[20] F. Pedregosa, G. Varoquaux, A. Gramfort, V. Michel, B. Thirion, O. Grisel, M. Blondel, P. Prettenhofer, R. Weiss, V. Dubourg, J. Vanderplas, A. Passos, D. Cournapeau, M. Brucher, M. Perrot, and É. Duchesnay, Scikit-learn: Machine Learning in Python, Journal of Machine Learning Research **12**, 2825 (2011).
[21] J. R. Gardner, G. Pleiss, D. Bindel, K. Q. Weinberger, and A. G. Wilson, GPyTorch: Blackbox Matrix-Matrix Gaussian Process Inference with GPU Acceleration (2021), arXiv:1809.11165 [cs].
[22] P. Luo and R. L. Baldwin, Interaction between water and polar groups of the helix backbone: An important determinant of helix propensities, Proceedings of the National Academy of Sciences **96**, 4930 (1999).
[23] G. Labesse, N. Colloc'h, J. Pothier, and J. P. Mornon, P-SEA: A new efficient assignment of secondary structure from C alpha trace of proteins, Computer applications in the biosciences: CABIOS **13**, 291 (1997).
[24] S.-Y. Park, M.-J. Yoo, J.-M. Shin, and K.-H. Cho, SABA (secondary structure assignment program based on only alpha carbons): A novel pseudo center geometrical criterion for accurate assignment of protein secondary structures, BMB Reports **44**, 118 (2011).
[25] S. M. Law, A. T. Frank, and C. L. Brooks III, PCASSO: A fast and efficient C$\alpha$-based method for accurately assigning protein secondary structure elements, Journal of Computational Chemistry **35**, 1757 (2014).
[26] M. A. Sallal, W. Chen, and K. A. Nasr, Machine Learning Approach to Assign Protein Secondary Structure Elements from C$\alpha$ Trace, in *2020 IEEE International Conference on Bioinformatics and Biomedicine (BIBM)* (2020) pp. 35–41.
[27] M. N. Saqib, J. D. Kryś, and D. Gront, Automated Protein Secondary Structure Assignment from C$\alpha$ Positions Using Neural Networks, Biomolecules **12**, 841 (2022).

[28] L. Heo and M. Feig, One bead per residue can describe all-atom protein structures, Structure **32**, 97 (2024).
[29] P. Rotkiewicz and J. Skolnick, Fast procedure for reconstruction of full-atom protein models from reduced representations, Journal of Computational Chemistry **29**, 1460 (2008).
[30] V. Tozzini, W. Rocchia, and J. A. McCammon, Mapping All-Atom Models onto One-Bead Coarse-Grained Models: General Properties and Applications to a Minimal Polypeptide Model, Journal of Chemical Theory and Computation **2**, 667 (2006).
[31] R. B. Best and G. Hummer, Optimized Molecular Dynamics Force Fields Applied to the Helix-Coil Transition of Polypeptides, The Journal of Physical Chemistry B **113**, 9004 (2009).
[32] J. A. Maier, C. Martinez, K. Kasavajhala, L. Wickstrom, K. E. Hauser, and C. Simmerling, ff14SB: Improving the Accuracy of Protein Side Chain and Backbone Parameters from ff99SB, Journal of Chemical Theory and Computation **11**, 3696 (2015).
[33] J. Huang, S. Rauscher, G. Nawrocki, T. Ran, M. Feig, B. L. de Groot, H. Grubmüller, and A. D. MacKerell, CHARMM36m: An improved force field for folded and intrinsically disordered proteins, Nature Methods **14**, 71 (2017).
[34] M. J. Abraham, T. Murtola, R. Schulz, S. Páll, J. C. Smith, B. Hess, and E. Lindahl, GROMACS: High performance molecular simulations through multi-level parallelism from laptops to supercomputers, SoftwareX **1–2**, 19 (2015).
[35] S. Jo, T. Kim, V. G. Iyer, and W. Im, CHARMM-GUI: A web-based graphical user interface for CHARMM, Journal of Computational Chemistry 10.1002/jcc.20945 (2008).
[36] Schrödinger, LLC, The PyMOL molecular graphics system, version 1.8 (2015).
[37] B. Hess, H. Bekker, H. J. C. Berendsen, and J. G. E. M. Fraaije, LINCS: A linear constraint solver for molecular simulations, Journal of Computational Chemistry **18**, 1463 (1997).
[38] R. B. Best, D. de Sancho, and J. Mittal, Residue-Specific $\alpha$-Helix Propensities from Molecular Simulation, Biophysical Journal **102**, 1462 (2012).
[39] S. Lifson and A. Roig, On the Theory of Helix—Coil Transition in Polypeptides, The Journal of Chemical Physics **34**, 1963 (1961).
[40] B. H. Zimm and J. K. Bragg, Theory of the Phase Transition between Helix and Random Coil in Polypeptide Chains, The Journal of Chemical Physics **31**, 526 (1959).
[41] T. P. Creamer and G. D. Rose, Interactions between hydrophobic side chains within $\alpha$-helices, Protein Science **4**, 1305 (1995).
[42] D. W. Urry, D. C. Gowda, T. M. Parker, C.-H. Luan, M. C. Reid, C. M. Harris, A. Pattanaik, and R. D. Harris, Hydrophobicity scale for proteins based on inverse temperature transitions, Biopolymers **32**, 1243 (1992).
[43] R. M. Regy, J. Thompson, Y. C. Kim, and J. Mittal, Improved coarse-grained model for studying sequence dependent phase separation of disordered proteins, Protein Science **30**, 1371 (2021).
[44] M. Bonomi, G. Bussi, C. Camilloni, G. A. Tribello, P. Banáš, A. Barducci, M. Bernetti, P. G. Bolhuis, S. Bottaro, D. Branduardi, R. Capelli, P. Carloni, M. Ceriotti, A. Cesari, H. Chen, W. Chen, F. Colizzi, S. De, M. De La Pierre, D. Donadio, V. Drobot, B. Ensing, A. L. Ferguson, M. Filizola, J. S. Fraser, H. Fu, P. Gasparotto, F. L. Gervasio, F. Giberti, A. Gil-Ley, T. Giorgino, G. T. Heller, G. M. Hocky, M. Iannuzzi, M. Invernizzi, K. E. Jelfs, A. Jussupow, E. Kirilin, A. Laio, V. Limongelli, K. Lindorff-Larsen, T. Löhr, F. Marinelli, L. Martin-Samos, M. Masetti, R. Meyer, A. Michaelides, C. Molteni, T. Morishita, M. Nava, C. Paissoni, E. Papaleo, M. Parrinello, J. Pfaendtner, P. Piaggi, G. Piccini, A. Pietropaolo, F. Pietrucci, S. Pipolo, D. Provasi, D. Quigley, P. Raiteri, S. Raniolo, J. Rydzewski, M. Salvalaglio, G. C. Sosso, V. Spiwok, J. Šponer, D. W. H. Swenson, P. Tiwary, O. Valsson, M. Vendruscolo, G. A. Voth, A. White, and The PLUMED consortium, Promoting transparency and reproducibility in enhanced molecular simulations, Nature Methods **16**, 670 (2019).
[45] G. A. Tribello, M. Bonomi, D. Branduardi, C. Camilloni, and G. Bussi, PLUMED 2: New feathers for an old bird, Computer Physics Communications **185**, 604 (2014).
[46] S. Basu, P. Martínez-Cristóbal, M. Frigolé-Vivas, M. Pesarrodona, M. Lewis, E. Szulc, C. A. Bañuelos, C. Sánchez-Zarzalejo, S. Bielskutė, J. Zhu, K. Pombo-García, C. Garcia-Cabau, L. Zodi, H. Dockx, J. Smak, H. Kaur, C. Batlle, B. Mateos, M. Biesaga, A. Escobedo, L. Bardia, X. Verdaguer, A. Ruffoni, N. R. Mawji, J. Wang, J. K. Obst, T. Tam, I. Brun-Heath, S. Ventura, D. Meierhofer, J. García, P. Robustelli, T. H. Stracker, M. D. Sadar, A. Riera, D. Hnisz, and X. Salvatella, Rational optimization of a transcription factor activation domain inhibitor, Nature Structural & Molecular Biology **30**, 1958 (2023).
[47] C. Camilloni, A. De Simone, W. F. Vranken, and M. Vendruscolo, Determination of Secondary Structure Populations in Disordered States of Proteins Using Nuclear Magnetic Resonance Chemical Shifts, Biochemistry **51**, 2224 (2012).
[48] G. L. Dignon, W. Zheng, Y. C. Kim, R. B. Best, and J. Mittal, Sequence determinants of protein phase behavior from a coarse-grained model, PLOS Computational Biology **14**, e1005941 (2018).
[49] C. R. Chen and G. I. Makhatadze, ProteinVolume: Calculating molecular van der Waals and void volumes in proteins, BMC Bioinformatics **16**, 101 (2015).
[50] W. L. Jorgensen, D. S. Maxwell, and J. Tirado-Rives, Development and Testing of the OPLS All-Atom Force Field on Conformational Energetics and Properties of Organic Liquids, Journal of the American Chemical Society **118**, 11225 (1996).
[51] J. L. F. Abascal and C. Vega, A general purpose model for the condensed phases of water: TIP4P/2005, The Journal of Chemical Physics **123**, 234505 (2005).
[52] I. M. Zeron, J. L. F. Abascal, and C. Vega, A force field of Li+, Na+, K+, Mg2+, Ca2+, Cl-, and SO42- in aqueous solution based on the TIP4P/2005 water model and scaled charges for the ions, The Journal of Chemical Physics **151**, 134504 (2019).
[53] M. K. Scherer, B. Trendelkamp-Schroer, F. Paul, G. Pérez-Hernández, M. Hoffmann, N. Plattner, C. Wehmeyer, J.-H. Prinz, and F. Noé, PyEMMA 2: A Software Package for Estimation, Validation, and Analysis of Markov Models, Journal of Chemical Theory and Computation **11**, 5525 (2015).
[54] S. Röblitz and M. Weber, Fuzzy spectral clustering by PCCA+: Application to Markov state models and data classification, Advances in Data Analysis and Classification **7**, 147 (2013).
[55] M. Boehning, C. Dugast-Darzacq, M. Rankovic, A. S. Hansen, T. Yu, H. Marie-Nelly, D. T. McSwiggen, G. Kokic, G. M. Dailey, P. Cramer, X. Darzacq, and M. Zweckstetter, RNA polymerase II clustering through carboxy-terminal domain phase separation,

Nature Structural & Molecular Biology **25**, 833 (2018).

[56] B. R. Sabari, A. Dall'Agnese, A. Boija, I. A. Klein, E. L. Coffey, K. Shrinivas, B. J. Abraham, N. M. Hannett, A. V. Zamudio, J. C. Manteiga, C. H. Li, Y. E. Guo, D. S. Day, J. Schuijers, E. Vasile, S. Malik, D. Hnisz, T. I. Lee, I. I. Cisse, R. G. Roeder, P. A. Sharp, A. K. Chakraborty, and R. A. Young, Coactivator condensation at super-enhancers links phase separation and gene control, Science **361**, eaar3958 (2018).
